# A recurrent pathogenic *BRCA2* truncating variant reveals a role for BRCA2-PCAF complex in modulating NF-κB-driven transcription

**DOI:** 10.1101/2025.05.12.652853

**Authors:** Anna Minello, Jesus Gomez-Escudero, Sreerama Chaitanya Sridhara, Charlotte Martin, Elodie Girard, Juan C. Cañas, Yasin Memari, Maria Rose Bustos, Antonio Galarreta, Virginie Boucherit, Evgeny Imyanitov, Sigridur Klara Bodvarsdottir, Sylvain Baulande, Anne Vincent-Salomon, Nicolas Servant, Matthias Altmeyer, Stefan Sigurdsson, Dominique Stoppa-Lyonnet, Serena Nik-Zainal, Aura Carreira

**Affiliations:** Institut Curie, Université PSL, CNRS UMR3348, 91400 Orsay, France; Université Paris-Saclay, CNRS UMR3348, 91400 Orsay, France; Genome Instability and Cancer Predisposition Laboratory, Centro de Biologia Molecular Severo Ochoa (CBM), CSIC-UAM, Madrid 28049, Spain; CBIO-Centre for Computational Biology, INSERM U900, Mines ParisTech, Paris, France; Academic Department of Medical Genetics, School of Clinical Medicine, University of Cambridge, Cambridge CB2 0QQ, UK; Early Cancer Institute, University of Cambridge, Cambridge CB2 0XZ, UK; Cancer Research Laboratory, Faculty of Medicine, School of Health Sciences, University of Iceland, Reykjavik, Iceland; BioMedical Center, School of Health Sciences, University of Iceland, Reykjavik, Iceland; Department of Molecular Mechanisms of Disease, University of Zurich, Zurich 8057, Switzerland; N.N. Petrov National Medical Research Center of Oncology, Saint Petersburg, Russia; Institut Curie, PSL University, ICGex Next-Generation Sequencing Platform, 75005 Paris, France; Diagnostic and Theragnostic Medicine Division, Institut Curie, Paris, France, Institut Curie, Paris, France; Department of Genetics, Institut Curie, Paris 75005, France; Paris-Cité University, Paris, France; INSERM U830, Institut Curie, Paris 75005, France

**Author notes:** Equal contribution (Anna Minello, Jesus Gomez-Escudero, Sreerama Chaitanya Sridhara).

**Keywords:** BRCA2, breast cancer, haploinsufficiency, LOH, genome instability, HRD, PARP inhibitors, pathogenic variants, PCAF, NF-κB

## Abstract

Germline monoallelic truncating mutations in BRCA2, a critical mediator of homologous recombination (HR), predispose individuals to breast and ovarian cancer. While tumorigenesis is usually attributed to biallelic inactivation, emerging evidence suggests that haploinsufficiency may suffice in certain contexts. To investigate this, we recreated two *BRCA2* pathogenic truncating variants in heterozygosis in non-tumorigenic breast epithelial cells. Cells carrying a truncating mutation that was not produced prompted sensitivity to PARP inhibitors (PARPi) and reduced the HR capacity indicating haploinsufficiency. Surprisingly, the other variant was expressed as a truncated product and prompted a transcriptional rewiring. Mechanistically, the truncated BRCA2 product formed abnormal oligomers with full-length BRCA2 and bound to the PCAF acetyltransferase, sequestering it. This led to reduced global histone H4 acetylation and decreased NF-κB transcriptional activity, ultimately impairing epithelial cell migration—a process also altered in tumors. Our findings uncover a previously unrecognized BRCA2–PCAF axis that modulates NF-κB-driven transcriptional program, a process that is co-opted by a recurrent BRCA2 pathogenic variant.

## Introduction

Inactivating germline mutations affecting a single copy of *BRCA2* are associated with an increased risk of breast and ovarian cancer. However, the precise sequence of events that take place from the inherited *BRCA2* mono-allelic mutation to tumorigenesis remains poorly defined. A wealth of evidence suggests that BRCA2 exerts its tumor-suppressive activity through its role in DNA repair by homologous recombination (HR)^1^. At DNA double-strand breaks (DSBs) or replicative DNA lesions, BRCA2 binds and loads RAD51 onto the resected ssDNA to promote pairing and strand exchange with the sister chromatid template for high-fidelity repair. At DSBs occurring at highly transcribed regions, BRCA2 supports DDX5-mediated unwinding of DNA-RNA hybrids for HR repair^2^. BRCA2 also protects stalled replication forks from aberrant nuclease degradation^3,4^, and it prevents and promotes the repair of replication stress-associated ssDNA gaps via HR^5–8^. Other less canonical functions include a scaffolding role in cytokinesis^9,10^, stabilizing kinetochore-microtubules attachments for proper chromosome segregation^11,^ and a poorly characterized transactivation function for transcription associated with the histone acetyltransferase p300/CBP-associated factor (PCAF), also known as KAT2B^12^. Whether these functions are linked to its tumor suppressor activity is still unclear.

As a tumor suppressor, the inactivation of the wild-type allele is believed to drive tumor formation; however, this event is not always found in BRCA2-associated tumors^13–17^ suggesting that the mutated allele *per se* may trigger tumorigenesis. This idea entails an impact of the mutated allele in heterozygosis, which has been reported in different model systems although with variable outcomes^1,18–28^. Interestingly, it has been reported that depending on the location of the variant in the sequence, the predisposition to breast or ovarian cancer is different^29^ suggesting that for example, depending on whether the variant is recognized by the nonsense-mediated mRNA decay machinery, the way tumorigenesis is triggered might differ. However, earlier studies found no evidence for this^30^. Here, to shed light on this question and the early steps of tumorigenesis, we recreated two well-known BRCA2 pathogenic truncating variants, c.771_775del (also known as c.999del5, p.Asn257Lysfs*17, hereafter del5), or c. 5946delT (p.Ser1982Argfs*22, hereafter delT) in heterozygosis in the non-malignant breast epithelial cell line MCF10A. We revealed that these two truncating variants lead to very different outcomes in heterozygosis: notably, cells carrying the del5 variant showed reduced levels of the BRCA2 full-length protein, high sensitivity to genotoxic stress, and reduced HR capacity. These results confirm haploinsufficiency. In contrast, cells carrying the delT variant showed normal levels of BRCA2, resistance to genotoxic stress, and normal HR capacity. Strikingly, the delT variant exerted a dominant negative effect resulting in epigenetic and transcriptomic changes related to epithelial cell proliferation and the NF-κB signaling pathway which were also found in a subset of tumors carrying the same variant. In cells, the delT variant conferred reduced cell growth and cell migration in 2D while increasing their invasive potential in 3D. We identify PCAF acetyltransferase, an interacting partner of BRCA2, as responsible for at least some of the NF-κB-related expression changes.

Overall, we uncovered an NF-κB transcriptional regulation axis that involves PCAF association with BRCA2 impacting cell migration and invasion potential that is rewired in tumors expressing a common pathogenic variant^31,32^. Moreover, we demonstrate that the phenotypic outcome of BRCA2 truncating variants is not equivalent and, we show that haploinsufficiency is conferred by a truncating variant of BRCA2 that is not produced. The differences in chemotherapy sensitivity in cells bearing distinct pathogenic variants in heterozygosis as observed in this work may represent an opportunity for cancer prevention.

## Results

### +/delT and +/del5 cells exhibit different sensitivity to PARPi and mitomycin C

To test haploinsufficiency associated with BRCA2 we chose to recreate two common germline pathogenic truncating variants identified in breast cancer patients: 1. 5946delT (c.5946del), a prevalent variant in individuals with an Ashkenazi Jewish ancestry^34^ resulting in an early stop codon in exon 11 and a mislocalized truncated protein^35^. 2. 999del5 (c.771_775del), a founder mutation in Iceland responsible for ∼70% of the *BRCA1/2* breast cancer predisposition in that population^36^, resulting in a premature stop codon in exon 9. The corresponding truncated protein is unstable, not detected^37^, and can be considered null.

Using transcription-activator-like effector nuclease (TALEn) and CRISPR-Cas9 methodology, we generated two independently derived cell lines bearing each mono-allelic truncating variant of BRCA2 in MCF10A, a non-transformed human mammary epithelial cell line ^33^ (Fig. 1a). The gene-edited population was sorted, the clones were expanded, and the mutation was verified by PCR amplification of the targeted genomic locus by Sanger sequencing (Fig. 1a). For 5946delT, the mutation obtained was 5947delG (thereafter +/delT); however, given that it results in the same premature stop codon (at aa 2002), we decided to pursue our experiments with these cells.

**Figure 1:**
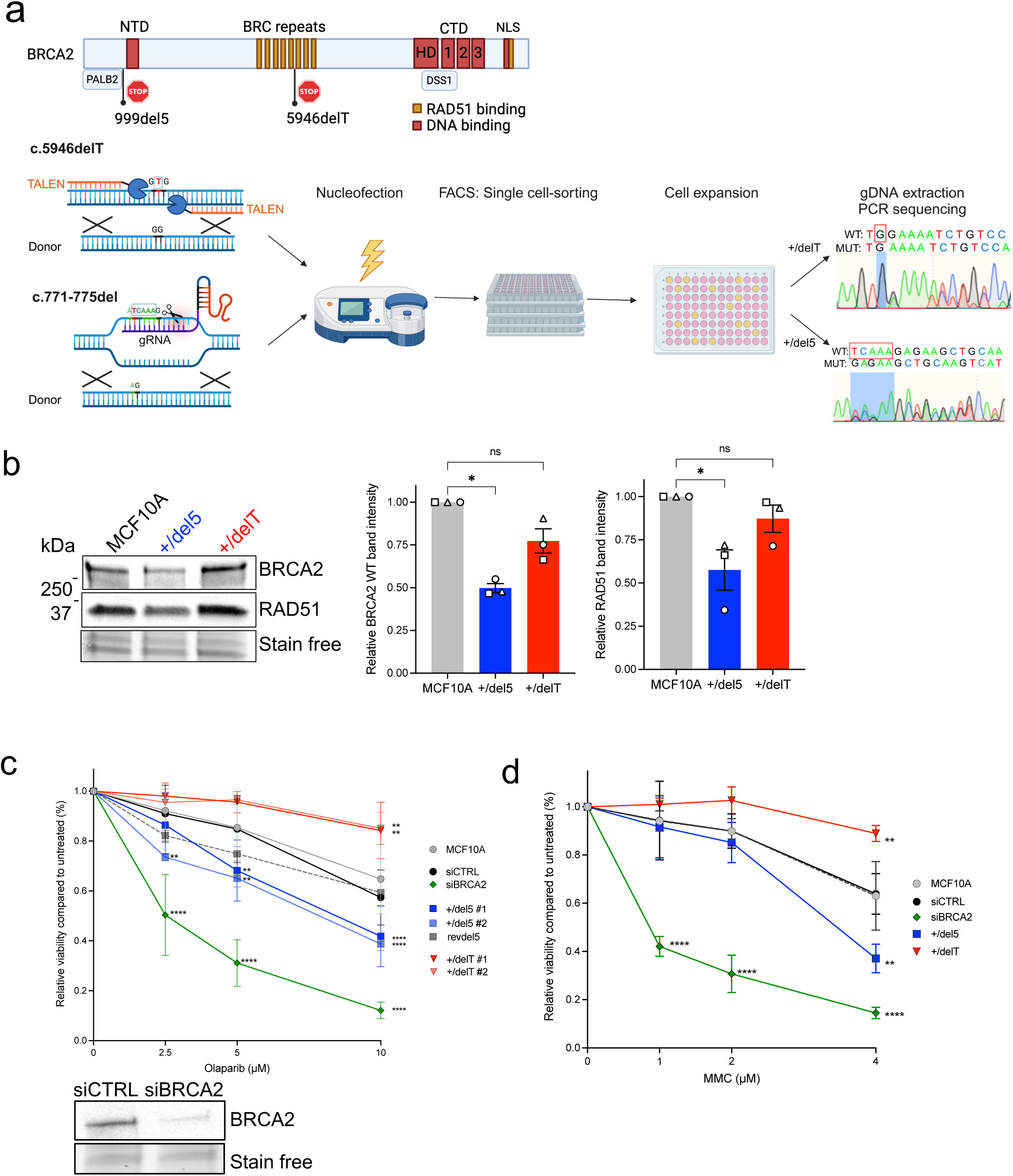
+/delT and +/del5 cells exhibit different sensitivity to PARPi and mitomycin C. **a,** (**Top**) BRCA2 schematic structure including the functional domains highlighting the two truncating mutations object of this study (stop sign) and the main interacting partners (below). Figure created with Biorender.com. DNA and RAD51 binding domains are represented in red and blue, respectively. Abbreviations: NTD, N-terminal DNA binding domain; CTD, C-terminal DNA binding domain; HD, helical domain; OB folds, oligonucleotide/oligosaccharide-binding folds; NLS, nuclear localization signal. **(Bottom left)** Schematic representation of the workflow to generate the stable heterozygote cell lines in MCF10A cells by TALEN (Pacman) (top) or CRISPR (scissors) (bottom) as indicated. (**Bottom right**) Sanger sequencing results. **b**, (**Left**) Representative Western Blot from whole cell extracts of the indicated cell lines showing BRCA2 and RAD51 protein levels detected with specific antibodies. (**Right**) Western Blot quantification of BRCA2 and RAD51 protein levels. Bands were normalized to the stain-free (loading control) and BRCA2 or RAD51 levels of the control cell line (MCF10A). The error bars represent the mean ± SD of three independent experiments, each represented with a different symbol shape. Statistical difference was determined on the mean by a Kruskal-Wallis test followed by Dunn’s multiple comparison test. ns, not significant, * p = 0.0127 (MCF10A vs +/del5 in BRCA2 levels), * p = 0.0199 (MCF10A vs +/del5 in RAD51 levels). **c**, Quantification of cell viability assays using MCF10A, two different clones of +/del5, two clones of +/delT, +/revdel5 and MCF10A cells depleted of BRCA2 (siBRCA2) or treated with control siRNA (siCTRL) and monitored by MTT assay upon treatment with increasing doses of the PARP inhibitor Olaparib at 6 days post-treatment, as indicated. The data represent the mean ± SD of the following number of independent experiments: siCTRL and +/del5 (n=6), MCF10A (n=5), +/delT #1 and +/del5 #2 (n=4), siBRCA2, +/delT #2 and +/revdel5 (n=3). **d,** Quantification of cell viability assays in the indicated cell lines and monitored by MTT assay upon 1h treatment with increasing doses of MMC, as indicated. After seeding, viability was monitored on day 6. The data represent the mean ± SD of three independent experiments. Statistical difference in **c-d** was determined by a two-way ANOVA test with Dunnett’s multiple comparisons test. The p-values show significant differences compared to the MCF10A cells. ** p <0.01, **** p<0.00001. (**Bottom left**) Representative Western Blot showing the depletion of BRCA2 protein levels as detected with a BRCA2 Ab (OP95) in cells transfected with siRNA against BRCA2 or CTRL 48h before the survival assay in **c** and **d**.

Given the role of BRCA2 in maintaining genome stability we evaluated the impact of the two pathogenic variants in heterozygosis in this function. We first determined the levels of BRCA2 protein in these cells. As expected, cells expressing +/del5 exhibited a reduction of BRCA2 protein levels to half compared to the MCF10A parental cells (Fig. 1b). In contrast, +/delT cells showed similar BRCA2 protein levels to that of the MCF10A parental cells. Interestingly, a similar trend was observed for RAD51 (Fig. 1b). This was despite similar levels of mRNA and protein of BRCA2 and RAD51 in the different clones bearing both variants (Extended Data Fig. 1a, b).

To evaluate the function of BRCA2 in these cells, we first used the PARP inhibitor Olaparib, a chemotherapeutic agent used in the clinic for HR-deficient *BRCA1/2*-mutated tumors^38,39^. As expected, cells depleted of BRCA2 showed high sensitivity to Olaparib whereas this was not the case for MCF10A cells (Fig. 1c). Remarkably, two +/del5 independent clones (Extended Data Fig. 1b) displayed moderate sensitivity to PARPi (Fig. 1c), consistent with the reduced levels of BRCA2 and RAD51 observed in these cells (Fig. 1b). Importantly, using gene-editing to correct del5 mutation to WT in one of these clones (Extended Data Fig. 1c) restored the resistance to levels comparable to that of the MCF10A parental cells (Fig. 1c) confirming that the sensitivity originated from the mutation in BRCA2 (Fig. 1c). A similar trend was observed for the DNA crosslinking agent mitomycin C (MMC), +/del5 showing increased sensitivity at 4μM unlike the +/delT cells (Fig. 1d). Intriguingly, the resistance of +/delT was higher than that of the MCF10A cells in PARPi conditions, which when correcting the mutation (hereafter +/revdelT; Extended Data Fig. 1e) was restored to MCF10A levels (Extended Data Fig. 1f). +/delT cells also showed hyper-resistance to MMC treatment (Fig. 1d).

To get more insights into the dysfunction of +/del5 cells, we monitored the number of γH2AX foci, a well-established marker of DSBs, and RAD51 foci using quantitative imaging-based cytometry (QIBC) in cells treated either with MMC and PARPi or left untreated. In these settings, MMC (3 μM, 24h) elicited a much stronger increase in DNA damage than PARPi (10 μM, 24h) as manifested by the overall increased intensity in γH2AX in all cells (Extended Data Fig. 2a). Interestingly, +/delT cells showed decreased levels of γH2AX in MMC conditions (Extended Data Fig. 2a) correlating with its increased resistance to MMC in the MTT assay (Fig. 1d). Cells treated with Olaparib showed only a slight increase in γH2AX foci and RAD51 foci in all cells; however, in the presence of MMC, +/del5 and +/delT cells displayed a reduced number of RAD51 foci compared to the MCF10A parental cell line (Extended Data Fig. 2a).

As both Olaparib and MMC cause replicative lesions in addition to other types of DNA damage, we also tested cell sensitivity to nucleotide depletion by hydroxyurea (HU) treatment. In contrast to MMC and PARPi, neither clone showed sensitivity to HU, exhibiting comparable resistance to that of MCF10A cells (Extended Data Fig. 2b). Given that the HU elicited only mild sensitivity even at a high dose (5 mM) in these cells, we conducted clonogenic survival under the same conditions to confirm these results. Consistent with the MTT assay, the clonogenic survival assay showed the same outcome (Extended Data Fig. 2c) concluding that these cells do not display higher sensitivity to nucleotide depletion compared to the MCF10A controls.

Altogether, only +/del5 cells showed reduced protein levels of BRCA2 and RAD51 and exhibited increased sensitivity to PARPi and MMC which was restored with the reverted mutation whereas +/delT displayed increased resistance to both treatments. However, both cell lines moderately reduced the number of RAD51 foci in MMC conditions whereas only +/delT decreased the number of γH2AX foci. Moreover, neither +/delT nor +/del5 cells changed sensitivity to nucleotide depletion-induced replication stress compared to the MCF10A controls.

### +/del5 cells show reduced HR capacity and accumulate PARPi-induced ssDNA gaps

Given the sensitivity to different genotoxic agents, we directly measured the DSB repair capacity of +/del5 cells using a modified version of our previous cell-based HR reporter assay^40^. This test measures DSB-mediated gene targeting activity at a specific locus (AAVS1 site) within the endogenous *PPP1R12C* gene. In this case, we co-transfected the cells with three plasmids, one expressing GFP-Cas9, one expressing a gRNA designed to cut at the AAVS1 site, and a promoter-less mCherry donor plasmid flanked by two homology sequences to the targeted locus. DSB-mediated gene targeting at the AAVS1 locus would result in mCherry expression from the endogenous PPP1R12C promoter, which is proportional to the HR repair capacity of the cells. Cells transfected with the mCherry plasmid and the GFP-Cas9 but without the gRNA were used as controls. To enrich the population of transfected cells, we sorted the GFP-Cas9 positive cells 2 days after transfection. After 6 days in culture, 1.5 % of MCF10A cells showed mCherry positive signal as compared to 0.3% of the control cells without gRNA. Similarly, 1.2 % of +/delT cells expressed mCherry. Importantly, only 0.5% of +/del5 cells showed mCherry positive signal (Fig. 2a) suggesting a reduced capacity of these cells to repair DSBs by HR.

**Figure 2:**
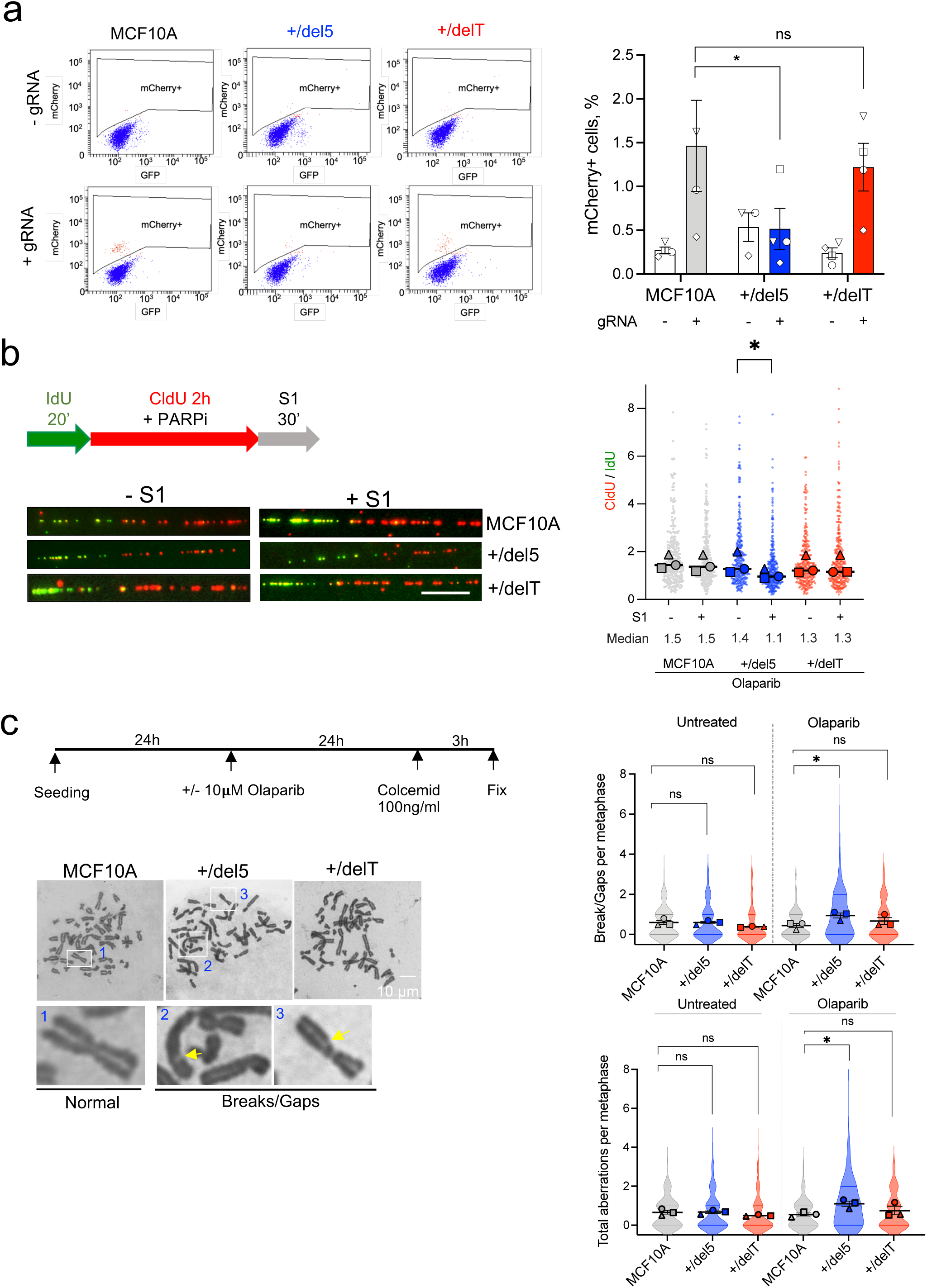
+/del5 cells show reduced HR capacity and accumulate PARPi-induced ssDNA gaps. **a, (Left)** Representative example of the flow cytometry gating and analysis of mCherry positive cells as a readout of the HR assay^40^ at 8 days post-transfection in MCF10A control and BRCA2-mutated cells. (**Right**) Frequency of mCherry positive cells in cells transfected with the promoter-less donor plasmid (AAVS1-2A-mCherry) without (empty bars) or with the gRNA, as indicated. The error bars represent the mean ± SEM of four independent experiments, each represented with a different symbol shape. Statistical difference was determined by a two-way ANOVA test followed by Dunnet’s multiple comparison test. The p-values show differences compared to the MCF10A cells. ns, not significant, * p = 0.0447. **b,** (**Top**) Labeling scheme and representative images of the replication tracks labeled as indicated from MCF10A control and BRCA2-mutated cells, in 30 μM Olaparib treated condition followed by 30 min of S1 nuclease (or S1 buffer only) treatment, as indicated. The scale bar indicates 15 μm. (**Right**) Quantification of CldU/IdU ratio in the indicated cells. The black line represents the median. Each dot represents a DNA fiber, and the means of each experiment are superimposed on the data. N=3 with 100 fibers analyzed for each condition. Statistical difference was determined by a two-tailed ratio paired t-test of the means for each cell line (-S1 vs +S1). *p= 0.0376. **c,** (**Left**) Experimental setup and representative images of chromatid breaks/gaps from the same cells either left untreated or upon treatment with Olaparib (10 μM for 24 h) as indicated. The scale bar indicates 10 μm. Quantification of breaks/gaps (**right, top**) or total chromosomal aberrations (**right, bottom**) in the cell lines indicated left untreated or treated with Olaparib. The means are superimposed on the individual data points represented in a violin plot. Statistical difference was determined by the two-way ANOVA test followed by Dunnett’s multiple comparison test. Breaks/gaps plot: ns, not significant, * p=0.0117. n=3 independent experiments with 40 metaphases analyzed per condition. Total aberrations plot: ns, not significant, * p=0.0108. n=3 independent experiments with 40 metaphases analyzed per condition.

BRCA2 deficient cells accumulate ssDNA gaps in conditions of replication stress such as PARPi or HU treatment^7,20,41^. We recently showed that PARPi- and HU-induced replication stress require distinct DNA binding domains of BRCA2^8^. Given that the del5 variant conferred sensitivity to PARPi but not to HU, we tested whether +/del5 cells accumulated PARPi-induced ssDNA gaps using single-molecule DNA combing^42^. Following a first pulse with IdU, cells were pulsed with CldU in the presence of 30 µM PARPi for 2h. After release, the cells were further incubated with or without S1 nuclease for 30 min; this enzyme creates nicks only in ssDNA regions resulting in a shorter tract length^25^. We then measured the ratio of CldU/IdU in both conditions for each cell line. As expected, MCF10A did not show sensitivity to the S1 enzyme as manifested by an equal median CldU/IdU ratio of 1.5 before and after treatment with S1 (Fig. 2b). This was also the case for +/delT cells (median CldU/IdU ratio of 1.3 before and after S1 treatment). In stark contrast, +/del5 cells displayed high sensitivity to S1 nuclease as revealed by a consistent reduction in the CldU/IdU ratio (median CldU/IdU ratio 1.4 (-S1) to 1 (+S1)), suggesting that +/del5 cells accumulate PARPi-induced ssDNA gaps (Fig. 2b). These results agree with the observed sensitivity of +/del5 cells to PARPi (Fig. 2b).

We further performed metaphase spreads after 24h treatment with PARPi (10 μM) to detect the presence of ssDNA gaps or breaks in mitosis. Only +/del5 cells exhibited an increased frequency of aberrations, in particular chromatid breaks/gaps, under these conditions whereas none of the cells showed this phenotype in untreated conditions (Fig. 2c).

Collectively, +/del5 cells exhibited a moderate decrease in HR capacity and accumulated PARPi-induced ssDNA gaps as revealed by S1-nuclease DNA combing and metaphase spreads. In contrast, +/delT cells showed normal DSB repair capacity and did not present detectable PARPi-induced ssDNA gaps.

### Breast tumors from del5 and delT-mutated patients display similar mutational HRD signature, percentage of LOH, and age at diagnosis

The phenotypes observed in our heterozygous cells suggested that a single mutation in BRCA2 might be sufficient to trigger genome instability and hence promote tumorigenesis. To shed light on this, we collected BRCA2 locus-specific LOH data from breast tumors bearing either one or the other variant, as LOH is the predominant way by which the second allele is inactivated^16^. We recovered data from previously published work^13,15,17,29,43^ (67), CRB of Institut Curie (9), Penn. University (2) and N.N. Petrov National Medical Research Center of Oncology (8) (index cases) gathering a total of 27 c.5946delT (delT) and 59 c.999del5 (del5) tumors. We found that the percentage of LOH in breast cancer patients was 63% for c.5946delT and 51% for c.999del5 cases (Fig. 3a) suggesting that LOH is not a pre-requisite for cancer (Table S1). Moreover, the age at diagnosis was equivalent among the patients with tumors that have undergone LOH versus those that do not (Fig. 3b, del5 median age: 48.5 (LOH) vs 49 (no LOH); delT median age: 47 (LOH) vs 46 (no LOH)).

**Figure 3:**
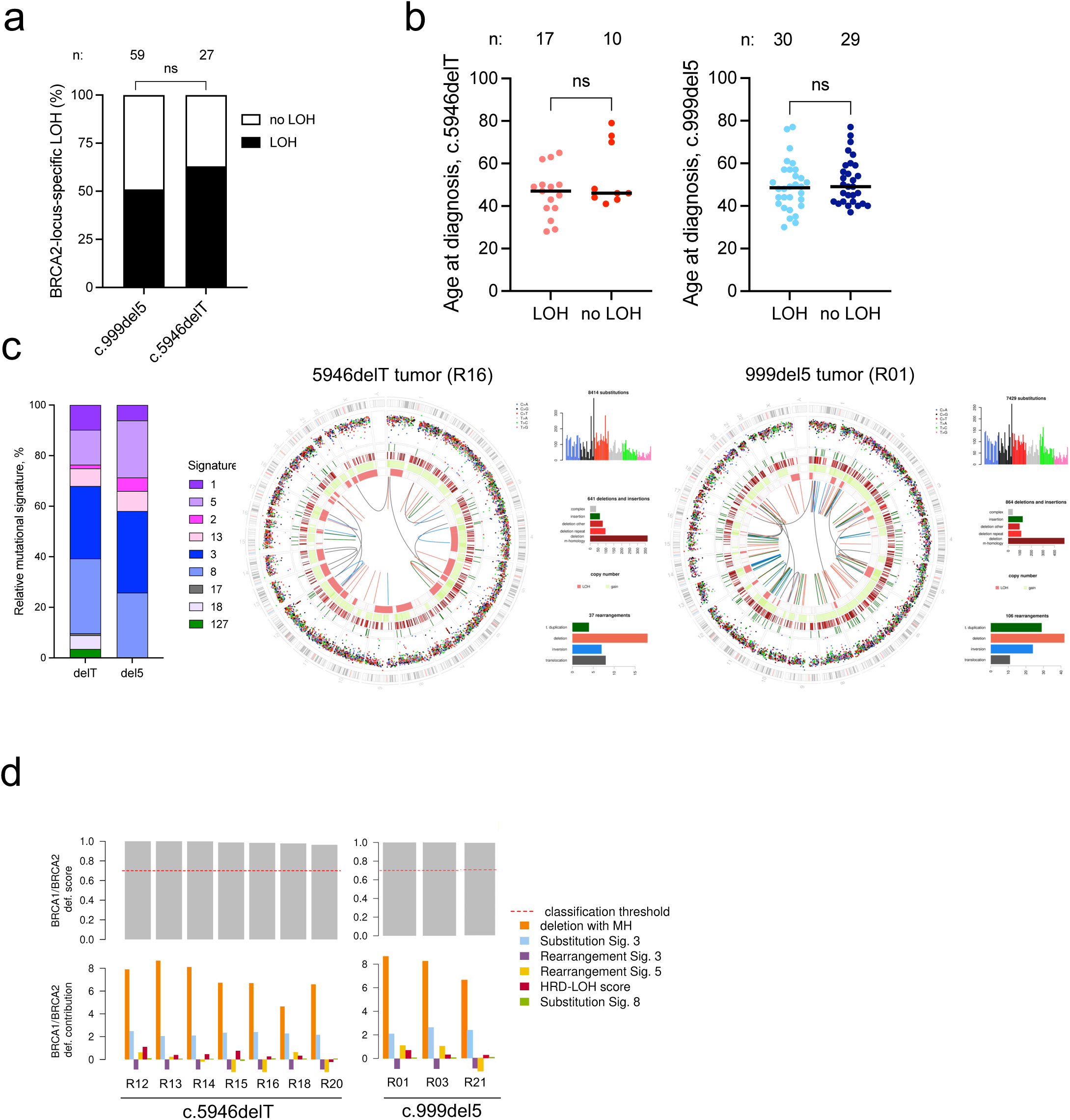
Breast tumors from del5 and delT-mutated patients display similar mutational HRD signature, percentage of LOH, and age at diagnosis. **a,** Bar graph representing the frequency of LOH at BRCA2 genomic locus in tumors bearing c.5946delT (27) or 999del5 (59) variant, as indicated. Statistical difference was determined by Fisher’s exact test. ns, not significant. **b,** (**Left**) Scatter plot representing the age at diagnosis of patients carrying c.5946delT (n=27) or (**right**) c.999del5 (n=59) variant, whose tumor has undergone LOH or not, as indicated. Statistical difference was determined using a two-tailed unpaired t-test. ns, not significant. **c,** (**Left**) Bar graph representing the proportion of mutational signatures (legend on the right) in a set of 7 primary tumors bearing the BRCA2 delT mutation and 11 with the BRCA2 del5 mutation, as indicated. (**Right**) Representative examples of the genome profile of a BRCA2 delT tumor and a del5 tumor, as indicated. The features depicted in the circos plots from the outermost rings moving inwards are (I) the karyotypic ideogram; (II) base substitutions; dot color: blue, C>A; black, C>G; red, C>T; gray, T>A; green, T>C; pink, T>G); (III) insertions shown as short green lines; (IV) deletions shown as short red lines; (V) major (green blocks, gain) and minor (red blocks, loss); and (VI) rearrangements shown as central lines (green, duplications; red, deletions; blue, inversions; grey, translocation). Next to each circos plot are histograms showing the: (top) number of mutations contributing to each substitution signature; (middle) deletion and insertions; and (bottom) the number of rearrangements contributing to each rearrangement signature. **d,** (**Top**) Bar plot of HRDetect scores for each sample. The threshold is fixed at 0.7 (red dotted line). (**Bottom**) Quantification of the contribution of each mutational signature per sample explaining the total HRD score (legend on the right).

Breast tumors defective in *BRCA1/2* generally display a specific set of genetic patterns of mutations (mutational signatures) that is collectively referred to as homologous recombination deficiency (HRD) signature. This pattern is thought to arise from the error-prone compensatory mechanisms that take place to repair DSB or collapsed forks when BRCA1/2 is not functional^44,45^. Given the differences we observed in the phenotype of breast epithelial cell lines bearing the delT and del5 mono-allelic pathogenic variants and the defective HR observed in the latter, we asked whether this phenotype correlated with a specific mutational pattern in tumors. To do so, we extracted DNA from a small cohort of breast cancer patients bearing the c.5946delT or c.999del5 variant. When possible, we recovered frozen tumor samples and matched adjacent healthy tissue, or else we compared the pattern with DNA extracted from blood from the same patient. As we could only recover 3 tumors from carriers of the c.999del5 variant, we also used previously published WGS data from 8 tumors bearing the same variant^44^. For the original samples, we performed WGS at 30X sequence depth, called the somatic mutations, and extracted the mutational signatures. In total, 7 delT tumors and 11 del5 tumors (8 previously reported ^44,^ and 3 new) were analyzed. We found that all tumors exhibited one or more of the following signatures: base-substitution signature 3 (SBS3), SBS8, rearrangement signature 5, and copy number profile showing widespread LOH and deletion with microhomology (Fig. 3c, Extended Data Fig. 3) as generally found in *BRCA1/2* deficient tumors^45^. We used HRDetect ^44^, a WGS-based predictor of HR deficiency based on 5 different signatures. According to the genomic profiles, all tumors analyzed *de novo* (7 delT and 3 del5) scored above the 0.7 cutoff indicating HRD signature (Fig. 3d) independently of their subtype or hormonal status (Table S2).

Together, 50-60% of tumors bearing c.5946delT and c.999del5 truncating pathogenic variants showed LOH suggesting a single mutation in *BRCA2* is sufficient for tumorigenesis. Moreover, BRCA2-locus LOH did not accelerate cancer onset in the 86 cases analyzed. Interestingly, despite the different genome instability features, both c.5946delT and c.999del5 truncating pathogenic variants gave rise to a similar HRD mutational signature in breast tumors.

### *BRCA2* delT pathogenic variant results in transcriptomic changes related to epithelial cell proliferation in cells and tumors

Given the apparent genomically stable phenotype of cells expressing the delT variant, we examined the transcriptome of these cells by RNAseq. We took 3 sets of independent clones arising from the same screen (Fig. 1a, Extended Data Fig. 1a) to interrogate whether the two mono-allelic variants resulted in any gene expression changes. As controls, we used the parental cell line (MCF10A, Ctrl #2) and two clones from either the del5 screen (Ctrl #1) or the delT screen (Ctrl #3) that did not result in gene-edited BRCA2. After selecting protein-coding genes and filtering out the genes with low expression, we pursued the analysis with 13,842 genes. Principal component analysis (PCA) discriminated +/delT clones from the MCF10A and +/del5 clones, explaining 35% of the variance by PC1 (Extended Data Fig. 4a). The comparison among the 3 biological replicates resulted in 5% of the transcriptome rewired: 722 differentially expressed genes (DEG) (300 up and 422 down) (log2 fold change > 1.5 and adj. p-value 0.05) compared to the wild type counterparts (Fig. 4a, Table S3). In contrast, +/del5 cells showed no DEG. Interestingly, Gene Ontology (GO) enrichment analysis of biological processes (BP) indicated that the +/delT cells exhibited deregulated pathways pertaining to epithelial cell migration and epithelial proliferation processes (Fig. 4b).

**Figure 4:**
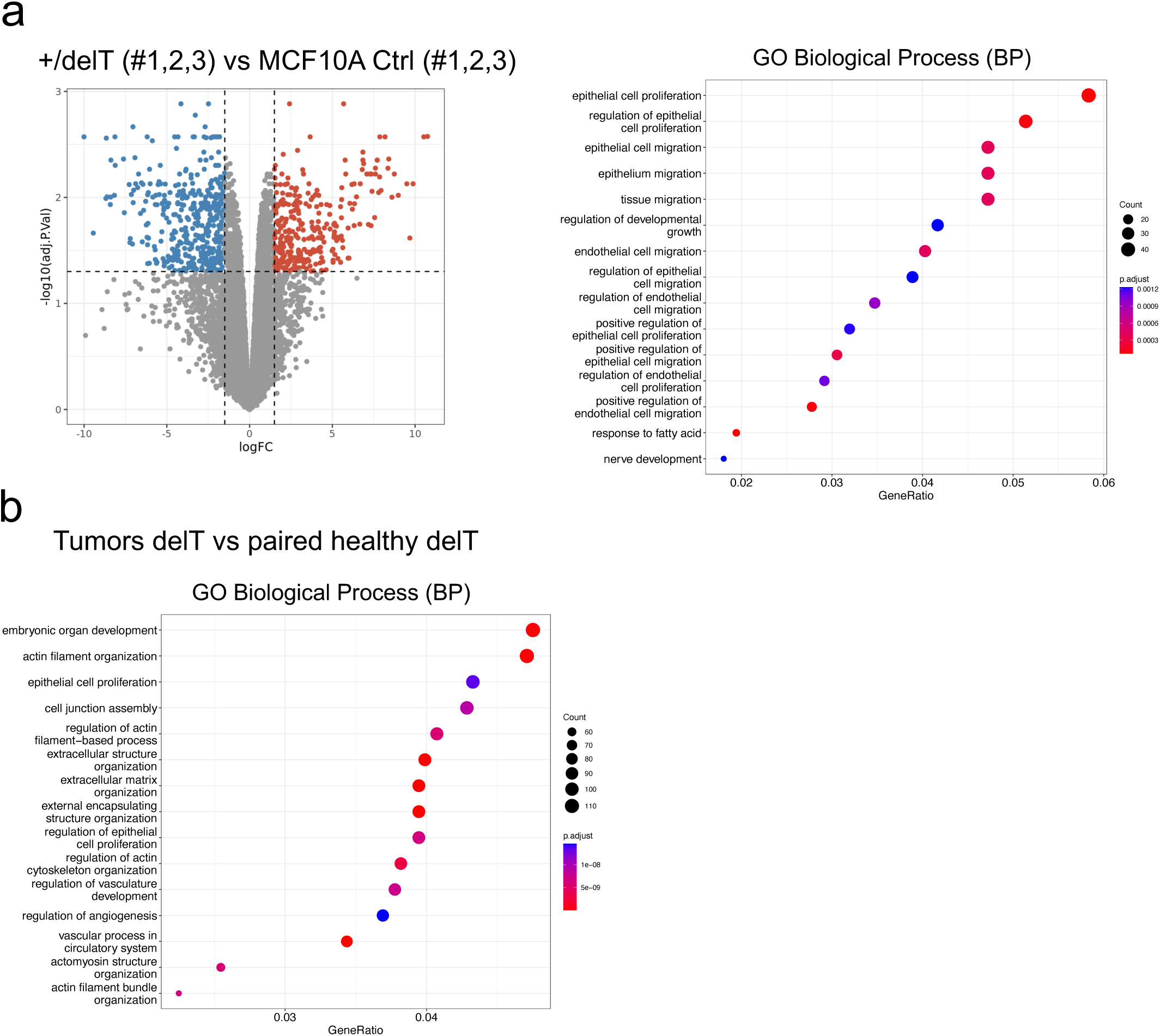
*BRCA2* delT pathogenic variant results in transcriptomic changes related to epithelial cell proliferation in cells and tumors. **a, (Left)** Volcano plot representing the differentially up-regulated (red) and down-regulated genes (blue) of 3 biological replicates of +/delT clones (from separate isolated clones) when compared to 3 biological replicates (from separate isolated clones) of MCF10A controls. DEGs were obtained by applying a log2 fold change > 1.5 and an adjusted p-value <0.05. (**Right**) Bubble plot showing the differentially expressed genes (DEG) categorized into biological processes (BP) from the same data, ranked by adjusted p-value (red to blue: high to low significance) using clusterProfiler (v4.4.4). **b,** Bubble plot showing the top DEG in GO terms for BP in a cohort of 9 delT tumors compared to their matched healthy adjacent tissue controls (Table S2) ranked by adjusted p-value using clusterProfiler (v4.4.4).

Next, to find out whether the DEG observed in +/delT MCF10A cells were relevant in tumors, we collected RNA from 9 primary tumors and matched non-neoplastic adjacent tissue of breast cancer patients carrying the same germline mutation (delT), 7 of which were used for the mutational signature analysis (Fig. 3c, d), and performed RNAseq. We selected protein-coding genes and discarded those with low expression (n=14,479). A log2 fold change (LFC) of 1.5 (adj. p-value <0.05) was set to consider DEG compared to the matched adjacent tissue. In delT tumors, 1079 genes appeared upregulated and 1521 downregulated (Table S4). Interestingly, the top 3 deregulated genes pertained to the embryonic organ development, actin filament organization, and epithelial cell proliferation processes (Fig. 3b). Unfortunately, RNA from del5 tumors was not available to perform the same analysis.

Overall, cells expressing the delT variant resulted in substantial global transcriptomic changes in MCF10A cells whereas the del5 variant did not. The deregulated pathways in +/delT cells were related to the epithelial cell proliferation category which was also deregulated in a cohort of delT tumors.

### +/delT cells display a slower proliferation rate and reduced migration capacity in 2D while increasing the invasion capacity in 3D

Given that the delT mutation altered genes involved in cell migration and proliferation we interrogated the capacity of +/delT cells to grow and form colonies. We monitored the proliferation of these cells over 7 days. Interestingly, the proliferation rate of +/delT cells measured by an Incucyte system was reduced (Fig. 5a, Extended Data Fig. 5a) compared to the +/del5 and MCF10A parental cells despite a cell cycle profile unchanged (Extended Data Fig. 5b). Moreover, +/delT cells adopted a ‘fibroblast-like’ shape in contrast to the cuboidal shape of MCF10A and +/del5, typical of epithelial cells. This manifested in a different pattern of actin staining and reduced circularity in +/delT cells compared to the other cell lines (Fig. 5b).

**Figure 5:**
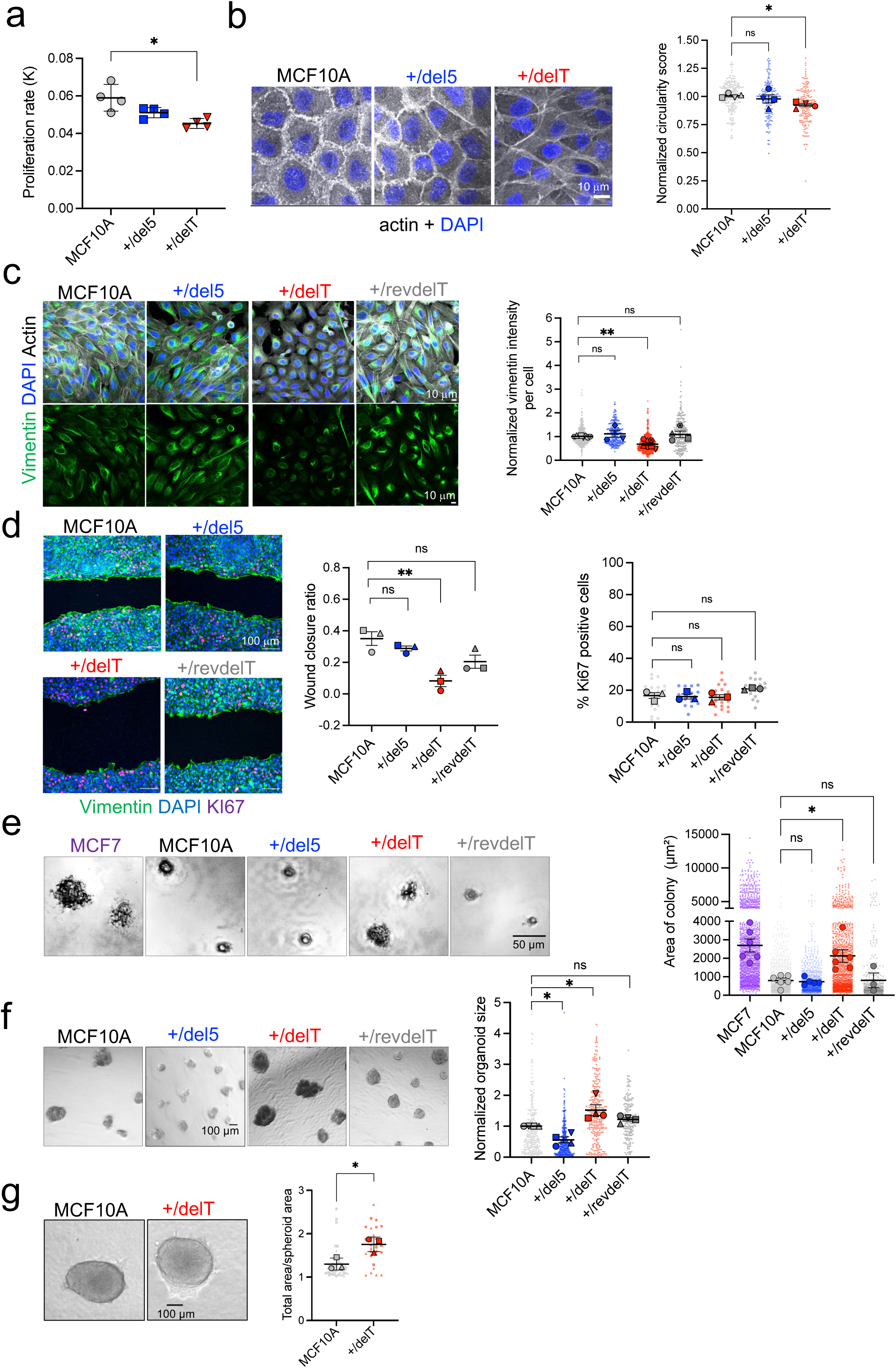
+/delT cells display a slower proliferation rate and reduced migration capacity in 2D while increasing the invasion capacity in 3D. **a,** Proliferation rate from the cell lines indicated. 5×10^3^ cells were seeded and monitored over 7 days (n=4). Statistical difference was determined by the Kruskal-Wallis test with Dunn’s multiple comparisons tests. * p = 0.0134. Only significant p-values are shown. **b,** (**Left**) Representative immunofluorescence images of MCF10A cells stained for F-actin (white); nuclei were counterstained with DAPI (blue). For each condition, 1×10^5^ cells were seeded. Scale bar: 10 μm. (**Right**) Quantification of the circularity score based on the actin staining at the junctions and calculated using the “cell shape” descriptor in ImageJ software. Each dot represents a cell from 4 independent experiments. 46 cells were measured in each experiment, and the circularity score was normalized to the MCF10A control. The means of each experiment are represented as different symbols superimposed on the data. Statistical difference was determined by a Kruskal-Wallis test with Dunn’s multiple comparisons test on the mean of each experiment. * p = 0.0285; ns, not significant. **c,** (**Left**) Representative immunofluorescence images of MCF10A cells stained for vimentin (green), actin (white), and DAPI (blue). Scale bar: 10 µm. (**Right**) Quantification of vimentin means intensity value per cell using actin staining to define the boundaries of the cells. 50-80 cells were analyzed per experiment and condition, (n=4). The means of each experiment are represented as different symbols superimposed on the data. Statistical difference was determined by Kruskal-Wallis with Dunn’s multiple comparisons test on the mean of each experiment. ns, not significant. ** p=0.0030. **d,** (**Left**) Representative immunofluorescence images of scratch wound healing assay of MCF10A cells stained for vimentin (green), Ki67 (magenta), and DAPI (blue) 16h after the wound. (**Right**) Quantification of wound closure ratio measuring the distance between the two fronts of the wound at t=0 and t=16h. The means of each experiment are represented as different symbols superimposed on the data. Statistical difference was determined by one-way ANOVA with Dunnett’s multiple comparisons test performed on the mean of each experiment. 6-10 wounds analyzed per condition (n=3). ns, not significant, **p= 0.02. (**Far right**) Quantification of the percentage of Ki67-positive cells in the Scratch-wound healing assay. 6-10 wounds analyzed per condition (n=3). The means of each experiment are represented as different symbols superimposed on the data. One-way ANOVA with Dunnet’s multiple comparison test was performed on the mean of each experiment. ns, not significant 0.98, 0.90, 0.17. **e,** (**Left**) Representative images of anchorage-independent growth of the MCF10A clones measured by soft agar assay as compared to MCF7 breast tumoral cells. 1×10^4^ cells were seeded and colonies were stained with crystal violet after 14 days. (**Right**) Quantification of the colony area. For each condition, 20 fields were analyzed; colonies were measured using ImageJ software. The lines represent the mean of 6 replicates except for the +/revdelT (n=3). The means of each experiment are superimposed on the data. Statistical difference was determined with the Kruskal-Wallis test with Dunn’s multiple comparisons test (the p-values show significant differences compared to MCF10A): * p = 0.0385. **f, (Left)** Representative 4X brightfield images of MCF10A, +/delT, and +/revdelT organoids. (**Right**) Quantification of the organoid area of the indicated cell lines 12 days after seeding 10,000 cells in 24-well plates in BME medium, (n=4). The means of each experiment are superimposed on the data. Statistical difference was determined by one-way ANOVA with Dunnet’s multiple comparison test on the means of each experiment. ns, not significant, * p = 0.0317 (MCF10A vs +/del5), 0.0119 (MCF10A vs +/delT). **g, (Left)** Representative brightfield images of spheroids formed by hanging drop method embedded in collagen gels imaged after 24h in culture. **(Right)** Quantification of the images on the left. n=3. Paired-t-test. * p=0.0492.

Given the changes in cell shape and the gene expression related to epithelial cell proliferation and migration observed in the +/delT cells, we monitored the vimentin levels, a cytoskeletal protein that regulates cell migration and shape and has been associated with epithelial-to-mesenchymal transition (EMT)^46^. Remarkably, +/delT displayed reduced vimentin cellular intensity as compared to MCF10A and the +/del5 cells (Fig. 5c). Correcting the mutation (+/revdelT) (Extended Data Fig. 1c, e) restored the levels of vimentin (Fig. 5c) suggesting the changes observed were specifically due to the delT variant.

To confirm the changes in epithelial cell migration directly, we performed a wound-healing assay^47^. First, a confluent monolayer of cells was serum-starved for 24h and released into a medium with 2% serum to limit cell growth. Under these conditions, a linear scratch was created using a 10 μl pipette tip, after which the closure of the wound was monitored at times 0h and 16h after incubation at 37°C (Extended Data Fig. 5c).

+/delT cells showed reduced wound closure capacity compared to the parental MCF10A cells or the +/del5 cells (Fig. 5d, Extended Data Fig. 5c). This phenotype was partially restored in the +/revdelT cells and was not due to differences in cell proliferation as the levels of the cell proliferation marker ki67 were comparable under these growth conditions (i.e., 2% serum) (Fig. 5d). We then monitored the capacity of the cells to individually form colonies as these features could be relevant for transformation. Despite its slower proliferation rate and reduced migration, +/delT cells showed normal colony formation potential or even slightly higher than the controls (Extended Data Fig. 5d). This prompted us to test their anchorage-independent growth capacity in soft-agar, a well-established method to evaluate the transformation potential of cells^48^. We used breast epithelial cancer cells MCF7 as a reference as they are known to grow under these conditions. MCF7 cells showed a median colony size of 2080 μm^2^ whereas MCF10A cells showed much smaller colonies, as expected (median of 537 μm^2^). Similar to MCF10A cells, +/del5 cells showed a median colony size of 474 μm^2^. Remarkably, +/delT showed a 2-fold increased colony size (1143 μm^2^) compared to MCF10A or +/del5 (Fig. 5e). Importantly, +/revdelT cells restored the size of the colonies to MCF10A levels (Fig. 5e) confirming the effect was due to the delT variant.

To mimic cell growth in the mammary gland, we plated MCF10A clones and adherent BME drops to generate organoids^49^. In these settings, all three cell lines formed multicellular organoids embedded in the medium (Fig. 5f). Interestingly, +/delT organoids grew bigger colonies compared to the parental MCF10A counterparts, and this phenotype was restored in the +/rev delT organoids (Fig. 5f), recapitulating the results obtained in soft-agar (Fig. 5e). In contrast, +/del5 organoids grew smaller colonies than the MCF10A cells (Fig. 5f). Consistently, spheroids from +/delT cells showed higher invasion capacity in collagen gel than MCF10A parental cells (Fig. 5g). Together, these results indicate that +/delT cells show decreased vimentin levels, reduced cell proliferation and circularity, and decreased cell migration capacity in 2D while increasing colony formation capacity of MCF10A in two anchorage-independent conditions in 3D and higher invasion potential in spheroid setting. In contrast, +/del5 cells displayed similar levels of vimentin and similar shape and migration capacity to MCF10A cells while reducing the organoid size.

### The delT variant induces epigenetic changes in the NF-**κ**B signaling pathway by sequestering the PCAF acetyltransferase

BRCA2 comprises a transactivation domain in its N-terminus^50^ whose activity requires its interaction with the acetyl-transferase PCAF^12^. Given the large transcriptomic changes observed in +/delT cells, we hypothesized that the truncated product of BRCA2 resulting from the expression of the delT variant, which comprises both the predicted transactivation domain and the PCAF interacting region, could be interfering with the transactivation activity of the wt allele of BRCA2 in the nucleus in a dominant negative fashion. To test this idea, we first set out to detect the truncated BRCA2 protein in +/delT cells with a predicted molecular weight of 224 kDa. Using an antibody raised against the N-terminal region of BRCA2 (validation in Extended Data Fig. 6b), we detected a weak band around 250 KDa marker (Fig. 6a) at the position at which this product runs in CAPAN-1 cells, a pancreatic cancer cell line expressing 3 copies of the same delT polypeptide^51^. We also performed a nuclear/cytoplasmic fractionation (Extended Data Fig. 6a) and found the same band in the nucleus of +/delT cells but not in +/del5 or MCF10A cells supporting the presence of the delT product in the nucleus of +/delT cells. Consistently, the delT product was located almost exclusively in the nucleus in CAPAN-1 cells, as it was PCAF, as manifested by a nuclear/cytoplasmic fractionation performed in these cells (Fig. 6b).

**Figure 6.**
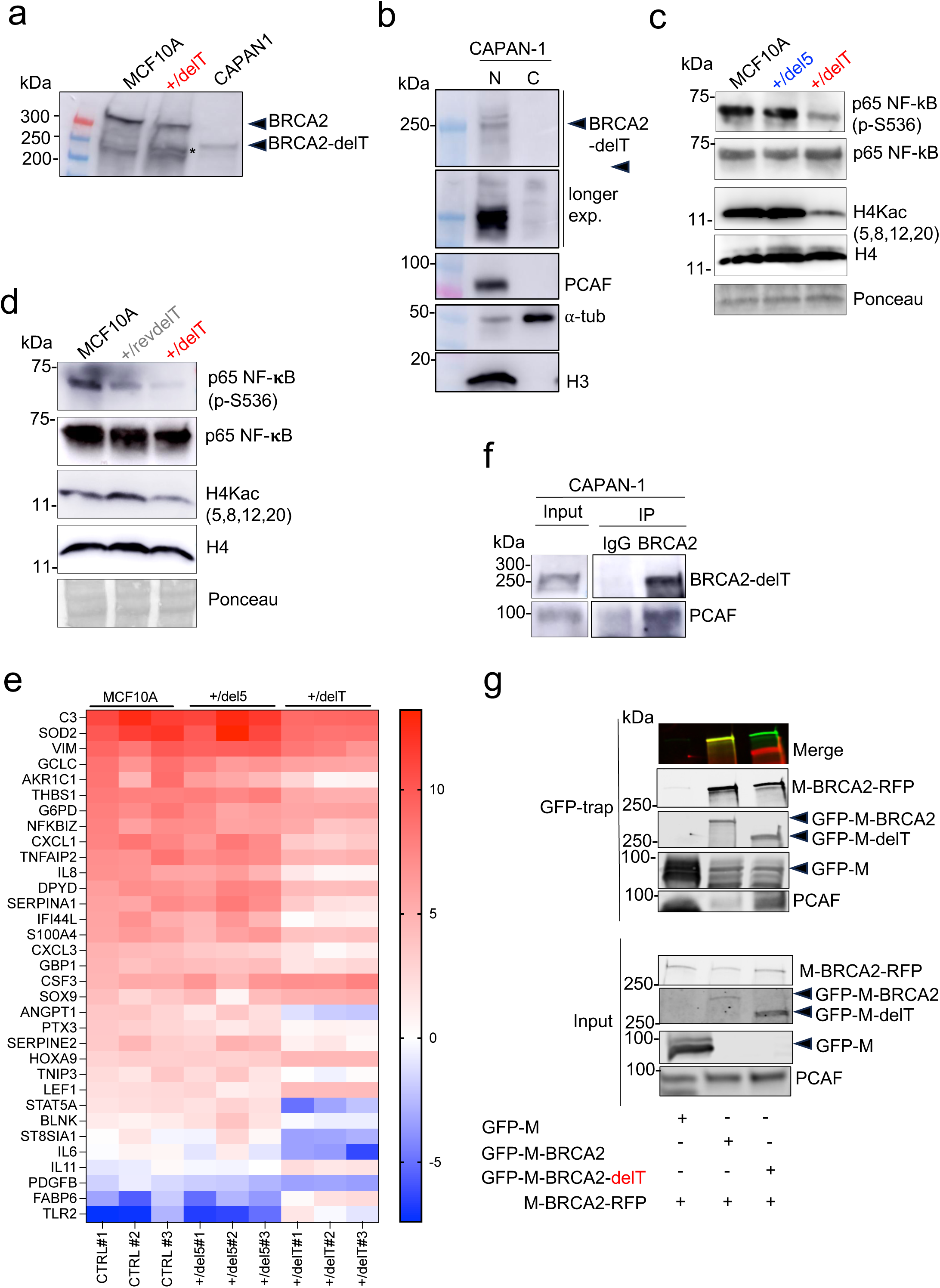
The delT variant induces epigenetic changes in the NF-κB signaling pathway by sequestering the PCAF acetyltransferase. **a,** Western blot showing the levels of the truncated delT product in the indicated cell lines using CAPAN1 cells expressing only the delT product as control. The protein was detected with an antibody raised against the N-terminal region of BRCA2 (Genscript). The antibody used for BRCA2 recognizes an epitope in the N-terminal region (Genscript, validation in Extended Data Fig. 6b). **b,** Immunoblot showing the delT BRCA2 (BRCA2 N-ter antibody), and PCAF levels in the cytoplasmic (C) and nuclear (N) fractions in CAPAN-1 cells. α-tubulin and H3 immunoblots served as controls of cytoplasmic and nuclear fractions, respectively. Data are representative of two independent experiments. **c,** Western blot showing acetylated lysines (K5, K8, K12, K16) on histone 4 (H4Kac) or S536 phosphorylation of the p65 subunit of NF-κB in MCF10A cells or cells bearing +/del5, or +/delT variant, as indicated. Below each blot is represented the mean ± SD of 3 experiments quantified relative to the total levels of H4 or NF-κB. Results are presented as percentages compared to the MCF10A parental clone. **d,** Western blot showing acetylated lysines (K5, K8 K12, K16) on histone 4 (H4Kac) or S536 phosphorylation of the p65 subunit of NF-κB in MCF10A, +/revdelT, or +/delT cells, as indicated. (n=2). **e,** Heatmap showing the expression of 33 known NF-κB target genes in MCF10A, +/del5, and +/delT clones, as indicated and assessed by RNAseq. **f,** IP of BRCA2 and detection of PCAF in complex with BRCA2-delT product present in human pancreatic cell line CAPAN-1. The BRCA2 antibody used is the same as in (A). Rabbit IgG was used as a control (n=3). **g,** GFP-trap pull-down of GFP-MBP-BRCA2 WT from HEK293T whole cell extracts co-transfected with GFP-MBP-BRCA2 WT (positive control), GFP-MBP-NLS (negative control) or GFP-MBP-delT with MBP-BRCA2-WT-RFP, in the presence of Benzonase. Complexes containing MBP-BRCA2-WT-RFP were run on 4–15% SDS-PAGE followed by western blot using anti-RFP and anti-GFP specific antibodies, as indicated. (n=3).

Previous work reported that the transient depletion of BRCA2 in breast epithelial cells supports survival in epidermal growth factor (EGF)-depleted conditions of the same cells once the expression of BRCA2 is recovered. This phenomenon was linked to the activation of the nuclear factor κB (NF-κB) pathway and an increased histone H4 acetylation^19^. Given that EGF induces NF-κB activation^52^ and that this pathway is related to migration and proliferation altered in +/delT cells and delT tumors, we tested the proliferation in EGF-depleted conditions and the levels of H4 acetylation and NF-κB activation. Using an MTT assay, we measured the fraction of surviving cells in the presence or absence of EGF. Consistent with the proliferation results (Fig. 5a), in normal media (+EGF), the survival was reduced in +/delT compared to the MCF10A cells; in contrast, the absence of EGF did not affect further their survival whereas it reduced that of the MCF10A cells (Extended Data Fig. 6c). We therefore performed western blot analysis to detect the levels of acetylation of histone H4 and the activation status of NF-κB monitoring the phosphorylation of the p65 (RelA) subunit at S536. Strikingly, +/delT cells showed reduced levels of both H4 acetylation (K5, K8, K12, K20) and NF-κB activation (p65-pS536) whereas +/del5 showed similar levels compared to the MCF10A cells (Fig. 6c). Importantly, these changes were partially restored in the +/revdelT cells (Fig. 6d) indicating they are linked to the delT variant. Given the known transcriptional activity of NF-κB, we looked at the mRNA levels of NF-κB target genes from our RNAseq data (from https://www.bu.edu/nf-kb/gene-resources/target-genes) in +/delT cells compared to the MCF10A and +/del5. Consistent with the reduced levels of p65-pS536, we found 33 genes known as target genes of NF-κB deregulated in +/delT cells with an overall downregulation of the genes compared to +/del5 and MCF10A cells (Fig 6e). An overall downregulation of NF-κB target genes was also observed in tumors delT vs their healthy controls (Extended Data Fig. 6d). We discarded a possible defect in NF-κB translocation to the nucleus as the signal of nuclear p65 was similar in MCF10A and +/delT cells (Extended Data Fig. 6e).

To test whether these transcriptional and epigenetic changes could be related to PCAF being sequestered by the delT product we next performed an immunoprecipitation (IP) of BRCA2 in nuclear cell extracts of both MCF10A and +/delT cells using the N-terminal BRCA2 antibody. As previously reported^12^, PCAF co-immunoprecipitated with BRCA2 this was in both MCF10A and +/delT cells, showing a slight increase in PCAF signal in the latter (Extended Data Fig. 6f). However, to determine whether the BRCA2 delT product alone formed a complex with PCAF, we performed a BRCA2 IP in CAPAN-1 cells, which lack the full-length BRCA2 protein. PCAF readily co-immunoprecipitated with the BRCA2 delT product in CAPAN-1 cells indicating that this truncated product of BRCA2 associates with PCAF in cells (Fig. 6f).

Given the low levels of delT product in +/delT cells, we hypothesized it may sequester PCAF by forming aberrant oligomers with full-length BRCA2. To test this, we co-expressed GFP-MBP–tagged delT and MBP-BRCA2-RFP in HEK293T cells and performed a GFP-trap. As expected, the GFP-MBP-BRCA2 formed a complex with MBP-BRCA2-RFP, unlike the GFP-MBP tag-only control, as we previously showed^53^. Importantly, the GFP-MBP-delT product was able to associate with MBP-BRCA2-RFP and this complex also associated with PCAF (Fig. 6g).

Together, the BRCA2 delT product was expressed in the nucleus in MCF10A +/delT and CAPAN-1 cells. Unlike the +/del5 cells, the +/delT cells showed reduced levels of histone H4 acetylation and NF-κB pS635 which was partly rescued when correcting the mutation. Consistently, EGF conferred a growth advantage to MCF10A cells over +/delT cells probably through NF-κB activation. Importantly, an overall downregulation of NF-κB target genes was observed in +/delT cells and tumors from patients expressing the delT variant. Finally, the BRCA2 delT product associated with PCAF as observed in CAPAN-1 cells and formed aberrant oligomers with the full-length BRCA2 and PCAF when overexpressed in HEK293T cells.

### Overexpression of PCAF partly rescues the NF-**κ**B target gene expression and the phenotype of +/delT cells

To further support the link between PCAF acetyltransferase and the transcriptomic changes observed in the +/delT cells, in particular of NF-κB-related genes, we overexpressed either PCAF-GFP or GFP as a control in +/delT cells and performed qRT-PCR for 6 known NF-κB target genes that were downregulated in the +/delT cells by RNAseq (Fig. 6d). We noticed that the transfection itself induced NF-κB activity so we analyzed the mRNA levels 6 days post-transfection.

We first analyzed the expression of these genes in +/delT vs MCF10A and confirmed by qRT-PCR the downregulation of *CXCL1,* IL6, *VIM*, *CXCL3*, *CXCL8,* and *NFKBIZ* (Extended Data Fig. 7a). We also confirmed that the mRNA levels of *RELA* and *PCAF* remained unchanged. Interestingly, transfecting +/delT cells with PCAF-GFP was sufficient to increase the mRNA levels of *CXCL1* and *VIM* without altering the mRNA levels of *RELA* compared to the same cells transfected with the empty vector (GFP) (Fig. 7a). A general increasing trend was also observed for *IL6*, *CXCL3*, *CXCL8* and *NFKBIZ* although it was not statistically significant probably due to the variability of transfection efficiency revealed by the different *PCAF* expression levels among the experiments (Extended Data Fig. 7b). Consistent with the overall recovery of the expression of these genes, overexpression of PCAF-GFP restored two of the phenotypes observed in the +/delT cells: it increased the vimentin protein intensity levels as detected by immunofluorescence (Fig. 7b) and it reduced the organoid growth of +/delT cells (Fig. 7c) compared to the GFP-alone transfected cells.

**Figure 7.**
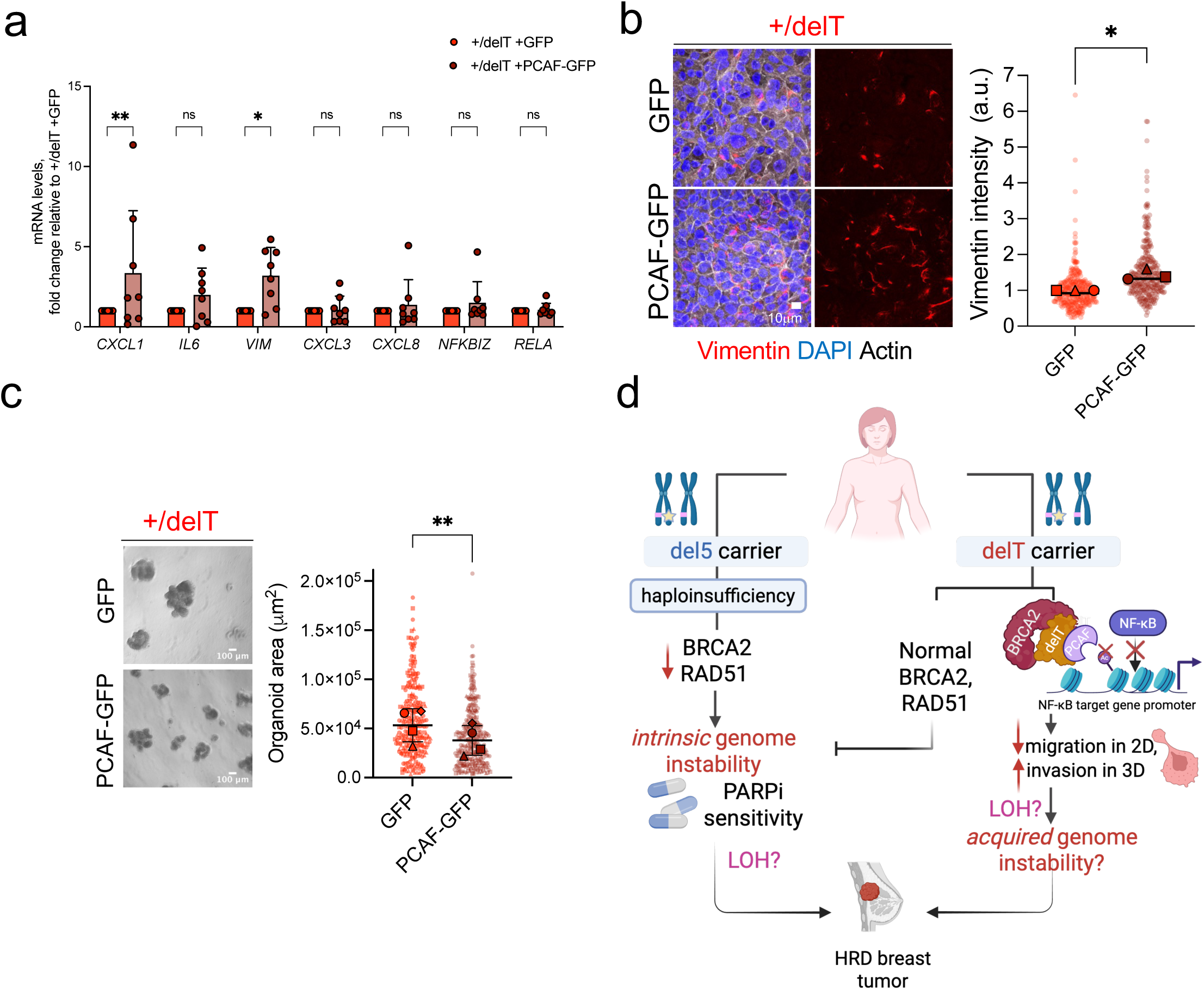
Overexpression of PCAF partly rescues the NF-kB target gene expression and the phenotype of +/delT cells. **a,** RT-qPCR of NF-κB regulated genes (*CXCL1*, *IL6, VIM, CXCL3, CXCL8, NFKBIZ*) and *NF-κB* (*RelA*) in GFP- or PCAF-GFP-transfected +/delT cells. Samples were collected six days after a single round of transfection with the indicated plasmids. The data was first normalized using *PPIA* as a reference gene and then represented as the fold-change expression of the indicated genes relative to GFP-transfected +/delT cells (n=8). Statistical difference was determined by two-way ANOVA with Šídák’s multiple comparisons test. **p=0.006, *p=0.012, ns: not significant, p>0.05. **b,** (**Left**) Representative immunofluorescence images of GFP or PCAF-GFP transfected +/delT cells stained for vimentin (red), actin (white), and DAPI (blue). Scale bar: 10 µm. (**Right**) Quantification of vimentin mean intensity value per cell using actin staining to define the boundaries of the cells. 30 cells were analyzed per field (n=3). The means of each experiment are represented as different symbols superimposed on the data. Statistical difference was determined by a t-student test with Welch correction. *p=0.037. **c,** (**Left**) Representative 4X brightfield images of organoids derived from +/delT MCF10A cells transfected with GFP or PCAF-GFP. (**Right**) Quantification of the organoid area at 12 days after seeding 10,000 cells in 24-well plates in BME media (n=4). The means of each experiment are superimposed on the data. Statistical difference was determined by paired t-test comparison on the means. ** p = 0.0083. **d,** Working model of the early processes of tumor formation in *BRCA2*-mutated carriers depending on the mutation (del5 or delT). Left arm: according to our results, cells in +/del5 carriers would show reduced BRCA2 and RAD51 levels leading to *intrinsic* genomic instability manifested in PARPi sensitivity, reduced HR, and accumulation of replication-associated ssDNA gaps. Right arm: Cells in BRCA2 +/delT carriers, while exhibiting normal BRCA2 and RAD51 levels that suppress genome instability, they would display alterations in cell-migration pathways and promote cell invasion in 3D, a phenotype linked to NF-κB transcriptional regulation. This rewiring is associated with the sequestering of its coactivator PCAF by the delT truncated product which forms aberrant oligomers with full-length BRCA2; this in turn reduces histone H4 acetylation limiting NF-κB-access to the chromatin and its related gene expression. Tumor formation in this scenario might involve epigenetic and transcriptomic rewiring of genes related to epithelial cell proliferation including NF-κB-regulated genes as observed in tumors expressing this variant. In both scenarios, LOH might occur as a second hit, take place at a later stage (or be induced), or be absent, without altering the age of onset. In the case of delT carriers, genome instability may be *acquired* as a fitness advantage for transformation. Both scenarios would lead to HRD as the predominant mutational signature in the primary breast tumor. Figure created with Biorender.com.

Together, overexpression of PCAF partially rescued the mRNA levels of NF-κB target genes tested, especially *CXCL1* and *VIM*. Accordingly, this was also sufficient to revert two of the phenotypes observed in +/delT cells: the reduced vimentin intensity levels and the increased organoid growth.

## Discussion

In this work, we interrogated whether BRCA2 heterozygosity confers haploinsufficiency by recreating two of the most common truncating pathogenic variants in non-malignant breast epithelial cells. Surprisingly, we found that the two variants lead to very different outcomes in heterozygosis: +/del5 cells displayed reduced levels of BRCA2 and RAD51, and genomic instability features such as increased sensitivity to MMC and PARPi, reduced HR capacity, and the presence of PARPi-induced ssDNA gaps. Accordingly, tumors bearing this variant showed an HRD signature. These results support the idea of haploinsufficiency conferred by BRCA2 truncating variants that are not produced. In stark contrast, +/delT cells showed similar levels of full-length BRCA2 and RAD51, hyper-resistance to PARPi and MMC, normal HR capacity, and did not accumulate PARPi-induced ssDNA gaps. Interestingly, +/delT cells showed a large deregulation of genes (5% of the protein-coding genes) related to epithelial cell migration and proliferation pathways that were commonly deregulated in breast tumors bearing the same pathogenic variant. This transcriptome rewiring correlated with overall reduced levels of acetylation of histone H4 and decreased activation of NF-κB p65 subunit. In line with this, +/delT cells showed reduced proliferation, decreased levels of vimentin, and poor collective cell migration in 2D while an increased capacity to grow both in anchorage-independent conditions, in organoid culture, and higher invasion in collagen gels, indicative of an invasive potential in 3D.

The delT polypeptide lacks the nuclear localization signals located at the C-terminus of BRCA2 and was shown to be mainly located in the cytoplasm in human pancreatic cancer cells CAPAN-1 when overexpressed with a GFP tag^35^. Using the same cell line, we detected the delT product almost exclusively in the nucleus. This discrepancy might arise from the overexpression of the GFP-delT product in the previous work.

How is the delT polypeptide located in the nucleus? PALB2 is a protein described to localize BRCA2 to the nucleus^54^; the region of interaction with PALB2 is encompassed in the delT product which could promote delT nuclear localization.

How does the delT product, which is expressed in low amounts, produce such a transcriptional and acetylation rewiring in the +/delT cells? BRCA2 has been reported to form self-interacting complexes that preferentially engage the N-terminal and C-terminal regions of the protein ^53,55,56^. We show here that the delT product associates with the full-length BRCA2 protein in the form of aberrant oligomers that contain PCAF, supporting the idea that the delT product “sequesters” PCAF.

What is the link between PCAF, acetylation of H4, and NF-κB activation? A positive feedback loop has been described for H4 acetylation and p65 phosphorylation: p65 phosphorylation promotes the binding of p65 to the acetyltransferase P300/CBP and the acetylation of H4. H4 acetylation enhances NF-κB gene transcription, further sustaining NF-κB activation^57^. Although this type of regulation has not been described for PCAF, PCAF acetylates both NF-κB^58^ and histones H3 and H4 and binds directly to P300/CBP^59^ . Thus, either as a partner of P300/CBP or independently, we propose that BRCA2-bound PCAF gets recruited to phosphorylated p65, promoting H4 acetylation and NF-κB binding to its target genes to activate transcription. In the +/delT cells, delT-sequestered PCAF cannot acetylate H4 precluding full access of NF-κB to its target genes, explaining the general downregulation of NF-κB target genes observed in +/delT cells and tumors (Fig. 7d). This model is supported by the following results: 1. H4 acetylation and NF-κB phosphorylation are partially restored in the CRISPR-corrected +/revdelT cells. 2. Overexpression of PCAF in +/delT cells rescues some of NF-κB target gene expression. 3. Overexpression of PCAF in +/delT cells rescues the vimentin levels and reduces organoid growth.

Our results strongly suggest that the BRCA2-PCAF complex modulates epithelial cell migration genes through the NF-κB signaling pathway and reveal this activity as a possible contributor to its tumor suppressor function. Intriguingly, this pathway and H4 acetylation were shown to be upregulated in conditions of transient depletion of BRCA2 conferring proliferating advantage in EGF-deprived medium^19^ in a model of non-transformed human breast epithelial cells. In contrast, another report showed the NF-κB signaling pathway was downregulated in prophylactic mastectomy-derived BRCA2^mut/+^ progenitor cells and non-transformed mammary epithelial cells knock-down for BRCA2^18^. These discrepancies might arise from the use of different model systems or reflect the possible dissimilarities between distinct variants as we observe in this study. Nevertheless, these reports and our work reinforce the idea that the NF-κB signaling pathway plays a role in *BRCA2*-linked tumorigenesis and we attribute this function, at least in part, to the association of BRCA2 with PCAF.

+/delT cells showed hyper-resistance to MMC and PARPi compared to MCF10A cells. These findings were surprising given the unchanged levels of BRCA2 and RAD51 protein in these cells. One possibility is that the decreased H4 acetylation observed in +/delT cells protects the DNA of these cells from damage consistent with the reduced number of γH2AX foci observed by QIBC upon MMC treatment in these cells. Moreover, PCAF acetyltransferase activity on histone H4 has been shown to promote the recruitment of MRE11 and EXO1 nucleases to resect DNA at replication forks, an activity that is controlled by ATR phosphorylation^61^. Consistently, depletion of PCAF in BRCA2 deficient cells, that cannot protect stalled replication forks, promotes PARPi resistance^61^. We speculate that by sequestering PCAF, the delT product could be promoting the PARPi hyper-resistance observed in +/delT cells, a mechanism that could be acting in tumors.

The fact that +/delT cells maintain similar BRCA2 WT protein levels to the MCF10A suggests a mechanism of homeostatic regulation that requires further investigation. From the number of RNAseq reads, we estimate that 10-20% of the total BRCA2 mRNA observed corresponds to the mutated allele. The maintained levels of full-length BRCA2 and RAD51 would explain why the DNA repair activity of BRCA2 and DNA damage sensitivity in these cells is not reduced. Importantly, the results obtained with +/del5 cells imply that the HR function of BRCA2 is dosage sensitive, suggesting that any HR hypomorphic mutation in BRCA2 could potentially trigger tumor formation. This idea agrees with the moderate breast cancer susceptibility we reported due to missense hypomorphic variants of *BRCA2*^62^. In contrast, BRCA2 transcriptional regulation activity does not seem to be dosage sensitive as the +/del5 cells did not show a perturbed transcriptional program.

Despite the lack of HR deficiency in heterozygosity, tumors expressing the delT variant also showed HRD signature suggesting that HRD is the predominant driver of tumor progression, as previously described for BRCA1/2-associated tumors^26^. Nonetheless, given that only 2 out of 10 tumors analyzed *de novo* in this study retained the BRCA2 WT allele, we cannot rule out the possibility that other mutational signatures might arise in other tumors that express the *BRCA2* WT allele. However, a recent report suggests that the accumulation of certain metabolites derived from glycolysis transiently depletes BRCA2 levels; this so-called *induced* haploinsufficiency in cells bearing a monoallelic truncating variant of BRCA2 resulted in mutational patterns related to HRD tumors^28^, which could play a role in the HRD signature observed in delT tumors that did not show LOH. Remarkably, LOH status did not change the age at diagnosis in patients bearing either variant strongly suggesting that inactivation of a single allele is sufficient to promote tumorigenesis as previously suggested^14^.

We propose that two different selective processes might be at play in the del5 vs delT scenarios to drive tumorigenesis (Fig. 7d). In c.5946delT, inactivating the WT allele would be an advantage for the tumor to instigate genome instability. This *acquired* genome instability contrasts with the *intrinsic* genome instability present in the case of 999del5, in which the levels of genome instability may be already sufficient to promote tumor formation. In this context, the loss of the second allele might occur as a consequence of genome instability rather than as a requisite for tumorigenesis. Consistent with this idea, only 51% of del5 tumors analyzed presented LOH.

In summary, a common pathogenic truncating variant of BRCA2 uncovers a role for the BRCA2-PCAF complex in NF-κB transcriptional control which could act as a driver of transformation and invasiveness in *BRCA2*-mutated tumors. Moreover, the collected data on LOH in tumors and the age at diagnosis from patients carrying both variants reinforce the idea that LOH is not a prerequisite for tumorigenesis consistent with the revised two-hit hypothesis, past and recent work^13,14,28,63^. In support of that idea, a recent study reported epigenetic effects of *Brca1* haploinsufficiency in accelerating tumorigenesis in mice models suggesting a single allele primes tumor formation in *BRCA1*-related tumors^64^.

Finally, our findings demonstrate that the phenotypic outcome of different *BRCA2* pathogenic variants, as exemplified with c.5946delT and c.999del5, is not equivalent and confirms that haploinsufficiency occurs in *BRCA2*-mutated context at least when the protein is not produced. These results may have important clinical implications as pathogenic variants are treated equally in the clinic. Based on our results, we predict that carriers of the del5 variant might suffer from higher toxicity to the healthy tissue when treated with PARPi. In contrast, delT carriers might benefit from PARPi as a prophylactic measure as the +/delT healthy tissue would not be sensitive to the drug.

## ACKNOWLEDGMENTS

We thank all the members of the Carreira lab for their insightful comments on the manuscript. We thank the CRB of Institut Curie (FR) for making available tissue samples used in this work and gratefully acknowledge the patients who donated. We thank Dr. Sandrine Caputo (Institut Curie, FR) for the clinical information on some of the tumor data presented in Table S1. We thank MD Katherine L. Nathanson and MD Susan Domchek (Penn. University) for insightful discussions and for providing LOH status information on some of the c.5946delT tumors in Table S1. We thank Miguel ngel Quintela lab (CNIO, SP) for their help generating organoids. We also thank Maaike Vreeswijk (Leiden Univ.) for helping find information on tumors through the CIMBA consortium and for insightful comments and Stéphan Vagner (Institut Curie, FR) for insightful comments on the manuscript. We are grateful to Prof. Kyle M. Miller (University of Texas, US) for the plasmid expressing PCAF-GFP and GFP control vector.

This work was supported by funding from ICGex (Institut Curie), Gray Foundation Basser Initiative “Innovation Award” (Penn. University), the French Breast Cancer Association Ruban Rose “Avenir Prize”, the Worldwide Cancer Research and the Spanish Association for Cancer Research (AECC) grant 22-0222 to A.C. A. M. was supported by a French governmental fellowship and a 4th year PhD Fellowship from the French Medical Research Foundation (FRM). SRC was supported by Agencia Española de Investigacion (MCIN/AEI) [PID2020115977RB-I00] grant to AC. EI was supported by the Russian Science Foundation (grant 17-75-30027). MB was supported by Göngum Saman. High-throughput sequencing was performed by the ICGex NGS platform of the Institut Curie supported by the grants ANR-10-EQPX-03 (Equipex) and ANR-10-INBS-09-08 (France Génomique Consortium) from the Agence Nationale de la Recherche (“Investissements d’Avenir” program), by the ITMO-Cancer Aviesan (Plan Cancer III) and by the SiRIC-Curie program (SiRIC Grant INCa-DGOS-465 and INCa-DGOS-Inserm_12554). Data management, quality control, and primary analysis were performed by the Bioinformatics platform of the Institut Curie.

## AUTHOR CONTRIBUTIONS

AM and AC conceived the study. AM, JCE, SRC, CM, and JCC performed the experiments with assistance from VB. MB extracted the DNA from some of the tumors under the supervision of SS and provided clinical information on tumors. SKB provided LOH status results and clinical information on 999del5 tumors. EG and YM performed the bioinformatic analysis with the supervision of NS and SNZ. AG performed the QIBC experiments under the supervision of MA. SB supervised next-generation sequencing analysis. AVS and her group at the CRB provided DNA, RNA, and clinical information on tumors from I. Curie. DSL provided insightful ideas and comments on the project and the manuscript. All authors analyzed the data; A.C. wrote the manuscript with input from all the authors.

## Methods

### Cell lines

MCF10A cells, kindly provided by Marie Dutreix Lab (Institut Curie, FR), were grown in DMEM-HG/F-12 (Biowest) supplemented with 5% horse serum (Thermo Fisher Scientific), 20 ng/ml human epidermal growth factor (Sigma-Aldrich), 0.5 mg/ml hydrocortisone (Sigma-Aldrich), 100 ng/ ml cholera toxin (Sigma-Aldrich), 10 µg/ ml insulin (Sigma-Aldrich), and 1% penicillin-streptomycin (Thermo Fisher). MCF7 (kindly provided by Patricia Uguen, Institut Curie, FR) were cultured using DMEM (HIMEDIA) supplemented with 10% FBS (EuroBio Abcys) and 1% of L-Glutamine (EuroBio Abcys). CAPAN1 cells were obtained from Dr. Ana M. Pizarro, (IMDEA Nanociencia, SP) and were grown in the same conditions as MCF7 supplemented with 20% FBS. HEK293T cells were a kind gift from Dr. Mounira Amor-Gueret’s lab. All cells were regularly tested for mycoplasma contamination.

### Generation of BRCA2 gene-edited clones

+/del5 and +/revdel5 were obtained using CRISPR/Cas9-gene editing. 2 µg for each of the following plasmids were used: pCas9-GFP (Addgene # 44719), target sequences cloned in pAC326 containing the pBS U6 sgRNA CRISPR (Addgene # 43860) expressing a reporter mCherry. The homology sequence was introduced by transfecting a single-strand oligonucleotide (oAC513 or oAC994, Table S5) containing the mutation flanked by homology arms to the endogenous BRCA2 region where the mutated base is located, modified with a phosphorothioate linkage (Sigma-Aldrich) and PAGE-purified. From the same screen, we obtained two other clones with 999del2 and 999del4 mutations that, although not equal, the first leads to the same premature stop codon only keeping one amino acid more, and the second results in a premature stop codon 40 aa downstream of the mutation instead of 17 (Extended Data Fig. 1a). +/delT was obtained using 5 µg of each TALEN encoding plasmid (TALEN-5′ #V35620 and TALEN-3′ #V35820, Life Technologies) and 1 µg of double-strand oligonucleotide comprising the delT mutation flanked with homology arms obtained by PCR from CAPAN-1 genomic DNA. Two more clones were obtained from the same screen bearing the same mutation (Extended Data Fig. 1a +/revdelT was obtained using CRISPR by transfecting 2 µg of the following plasmids: Cas9-GFP (Addgene #44719) and pU6-pegRNA-GG-acceptor (Addgene#132777), containing the target sequence and the homology sequence. The oligonucleotides and plasmids used for the generation of these clones are listed in Tables S5 and S6, respectively.

Cells were transfected using nucleofection (AMAXA technology, Lonza) using nucleofector kit T and the program T024 according to manufacturer instructions. The day after transfection the media was changed and 48 h post-transfection the cells were trypsinized. Individual GFP-mCherry double-positive cells were sorted using a BD FACSAria III (BD Bioscience) into 96 well-plates containing a complete culture medium. Single-cell-derived colonies were gradually expanded and screened for mutations in the two alleles using targeted PCR and the primers listed in Table S5.

### RNA extraction, RNAseq and analysis

2×10^6^ cells were harvested and RNA was isolated using the NucleoSpin Triprep kit (Macherey-Nagel). Before sequencing, RNA integrity and purity were verified through BioAnalyzer RNA 6000 Nano (Agilent) and Nanodrop. RNA libraries were prepared using standard Illumina protocols suitable for strand-specific RNA sequencing. Briefly, a first step of polyA selection using magnetic beads is done to focus sequencing on polyadenylated transcripts. After fragmentation, cDNA synthesis was performed, and resulting fragments were used for dA-tailing and ligation to indexed adapters (barcodes associated with each sample are included in Table S7). PCR amplification was finally achieved to create the final cDNA library. After qPCR quantification of the equimolar library pool using the KAPA library quantification kit (Roche), sequencing was carried out using 2*100 cycle mode (paired-end reads, 100 nucleotides) to get around 200M paired-end reads per sample.

Raw sequences were generated per sample using either the TruSeq Stranded mRNA kit on the Illumina HiSeq 2000 instrument (run B255) or the Stranded mRNA prep Ligation-Illumina on the NovaSeq 6000 (runs D610, D1342 and D1471) at the NGS platform of the Institut Curie, resulting in the production of 100bp paired-end fastq files. Raw data were then processed using the Institut Curie RNAseq pipeline https://zenodo.org/records/7446922. The overall quality of the reads was first checked using FastQC (respectively v0.11.9 for the run D1471, v0.11.8 otherwise), and the sequencing orientation, also known as strandness, was assessed using RseQC (respectively v4.0.0 for the run D1471, v2.6.4 otherwise). For the quality control (QC), those reads were then aligned on both a ribosomal RNAs database using bowtie (respectively v1.3.0 for the run D1471, v1.2.3 otherwise) and subsequently on the human reference genome (hg19 assembly) using STAR (respectively v2.7.6a for the run D1471, v2.6.1a otherwise).

Raw counts were generated by the bioinformatics platform pipeline using Star gene quantification mode (–quantMode GeneCounts) on the Gencode v19 gene database. On the 58720 Ensembl genes available, only 18699 protein-coding genes on autosomal and X chromosomes were kept in further analysis. Raw counts were normalized using the TMM method using the R packages EdgeR (v3.38.2) and Limma’s zoom function (v3.52.2). Genes with almost null expression were filtered out (n=13842). Principal Component Analysis (PCA) was performed on normalized data using the R package FactoMineR (v2.6). Heatmaps were generated using the R package pheatmap (v1.0.12). A linear model considering the annotation (Ctrl, +/del5, +/delT) was fitted to the counts. An adjusted p-value of 5% and a log fold change of 1.5 were set to consider a gene differentially expressed (DEGs). The R package clusterProfiler v4.4.4 was used to perform gene set enrichment analysis for the previous lists of DEGs.

Tumor samples were collected retrospectively after the informed consent of patients and are available through the CRB (Biological Resources Center) of Institut Curie. The RNA from the c.5946delT tumors and adjacent tissue was obtained following standard protocols; briefly, DNA was extracted by ethanol/chloroform after tissue dissociation, and the RNA was extracted using Qiazol followed by myRNeasy kit (Qiagen). The RNAseq from the tumors was analyzed using the same pipeline as mentioned for the cell clones.

### RT-qPCR

Approximately 1×10^6^ cells were harvested, and RNA was isolated using the NucleoSpin Triprep kit (Macherey-Nagel). After quantifying the total RNA using Nanodrop, equal amounts of RNA were reverse-transcribed to cDNA using iScript™ Advanced cDNA Synthesis kit (Bio-Rad). Then, the diluted cDNA was used to measure the expression of the respective genes in the CFX Opus Real-Time PCR system (Bio-Rad) using FastStart Universal SYBR Green Master (Merck). The relative mRNA levels were estimated as follows: 2^(Ct reference – Ct sample)^, where Ct reference and Ct sample are mean threshold cycles of RT-qPCR performed in duplicate or triplicate of the *PPIA* (reference) and the gene of interest (sample). The primers used for amplification of respective loci are included in Table S5.

### Cell proliferation assay with Incucyte System

5×10^3^ cells of MCF10A WT and BRCA2-mutated cells were seeded in triplicate in 24 well culture plates (Falcon/Corning). Cells were then incubated for 8 h at 37 °C in a humidified chamber to adhere to the culture plate. Cell growth was imaged every 3 hours over 7 days using the Incucyte System (Sartorius). The software calculates the% of confluence covered by cells at each time point. The proliferation rate was calculated from the interpolation of the logistic growth curve obtained in four independent experiments.

### Actin staining and circularity score

Confluent MCF10A cells seeded in coverslips inside of P24 plates were fixed with Paraformaldehyde 4% (Electron Microscopy Science) for 15 min at RT. After a wash with PBS, they were permeabilized with 0.1% Triton X (Fisher Scientific) in PBS for 10 min and then blocked in 2% BSA (Sigma-Aldrich) in PBS for 30 min. Primary antibodies were diluted into a blocking medium (2% BSA in PBS) and incubated overnight. The day after, coverslips were washed trice with PBS, and incubated with the appropriate 2^ary^ antibodies in a humidity chamber at 37 °C for 1 h. The coverslips were washed 3 times with PBS, and mounted with glass prolong mounting media (Thermo Fisher). mages were acquired with a 63x objective in an LSM800 Inverted Confocal Microscope (Zeiss) and analyzed using ImageJ software. To quantify actin intensity, a region of interest (ROI) was delineated around the perimeter of each cell and the mean fluorescent pixel intensity was measured. With the same ROIs, the circularity score (4π*area/perimeter^2) was calculated using the shape descriptor tool from ImageJ. Values are from 0 to 1, with a value of 1.0 indicating a perfect circle.

### Wound-healing assay

MCF10A cells were grown in P96 wells until confluence and then serum-starved for 24h. Then a scratch was performed in the middle of the well with a 10 µl pipette tip. Cell debris was removed by two washes with PBS 1X, and then cells were released in DMEM/F12 media supplemented with 2% horse serum to limit proliferation and incubated further at 37 °C. Bright-field images (Extended Data Fig. 5d) were taken at that moment (time 0) and after 16 h with a Leica Flexicam C1 camera. Wound size was calculated in the imageJ drawing 3 lines at the top, middle, and bottom of the wound image. Migration was calculated as the ratio of the average size of the wound at the endpoint (t=16h) compared to the initial time point (t=0h). Immunofluorescence images were taken at the endpoint with an LSM800 Inverted Confocal Microscope (Zeiss) and processed with ImageJ software to generate a maximal intensity projection.

### Transformation by soft-agar assay

1×10^4^ MCF10A cells were suspended in a top layer of DMEM-F12 supplemented media containing 0.30 % agar (Sigma) and plated on a bottom layer of DMEM-F12 supplemented media containing 0.6 % agar in 35 mm plates. The cells were additionally supplied with a complete medium every 3 days. After 3 weeks, colonies were stained and fixed with 0.01 % crystal violet and 10 % ethanol in H_2_O. Colonies were imaged with a 10X objective in a DM6000B upright widefield Microscope (Leica). For each condition, 20 fields were acquired. Images were quantified using ImageJ. The plugin extended depth of field was used to convert 3D acquisitions to 2D images; subsequently, colony size and circularity were measured using ImageJ version 1.53t.

### Organoid handling and growth assessment

1×10^4^ MCF10A cells were seeded on top of 250 µl of Cultrex Reduced Growth Factor Basement Membrane Gel (RyD) in P24 well plates in DMEM/F12 media (Gibco) supplemented with L-Glutamine, 1X B27 (Gibco), 10 µg/ml insulin (Merck 11376497001), 20 µg/ml hEGF (Sigma-Aldrich) and 1 µg/ml Hydrocortisone (Sigma-Aldrich). In Figure 6E, +/delT cells were transfected with GFP or PCAF-GFP plasmids (kind gift from Prof. Kyle Miller) using JetPrime according to the manufacturer’s specifications. The day after, cells were detached and 1×10^4^ were seeded on top of the Cultrex gel. The media was changed every 2-3 days, images were acquired on day 12 post-seeding, and the organoid area was measured by drawing a line delineating the perimeter of each organoid in ImageJ.

### Western blot

Cellular pellet was lysed in lysis buffer (50mM HEPES pH 7.5, 250mM NaCl, 5mM EDTA, 1% NP-40,1mM DTT, 1mM PMSF, 1X protease inhibitor cocktail (Roche)) and cells were incubated on ice for 60 min, vortexed every 10 minutes. Lysates were sonicated (3 times x 5 seconds) and then centrifuged at 18,000 x *g* for 30 min. at 4°C. The supernatant was transferred to a prechilled Eppendorf tube and stored at -80°C. For protein electrophoresis, samples were heated in 1x SDS sample buffer for 5 min. At 95°C, loaded on a stain-free 4-15% SDS gel (Bio-Rad), and migrated at 130 Volts for 90 min. in running buffer (1x Tris-Glycine, 0.1% SDS). The stain-free gel was visualized using a ChemiDoc camera (Bio-Rad). For transfer, a nitrocellulose membrane (VWR) was pre-equilibrated in a transfer buffer (1x Tris-Glycine, 0.025% SDS, 10% methanol). The proteins were transferred for 2 hours at 0.35 A at 4°C. The membrane was blocked in 5% milk in 1x TBS-T at room temperature for 45 minutes and then incubated with the respective antibody (see antibodies below) in 5% milk in 1x TBS-T overnight at 4°C. After extensive washes in TBS-T (3× 10 min), the membrane was incubated for 1h with the appropriate secondary HRP-antibody at room temperature on a shaker. After 3 more washes in TBS-T, the membrane was developed using ECL prime western blotting detection reagent (VWR) and visualized using a chemiluminescence imaging system either ChemiDoc camera (BIO-RAD) or Amersham Imager 680.

### Subcellular fractionation

For Extended Data Fig. 6a, 1×10^7^ MCF10A cells were trypsinized, pelleted and incubated on ice for 20 minutes with 500 µl of BAD buffer (10mM HEPES pH 7.9, 10mM KCl, 10% glycerol, 1.5mM MgCl_2_, 0.34M sucrose, 1mM DTT, 0.1mM PMSF, 1X protease inhibitor cocktail (EDTA-free, Roche)) complemented with 0.1% of Triton X-100. After collecting 20 µl of total cell lysate, the pellets were centrifuged at 1300 x *g* for 5 min at 4°C. The supernatant, containing the cytoplasmic fraction, was transferred to a prechilled tube. Pellets containing the chromatin fraction were then washed twice with 1.5ml BAD buffer and centrifuged at 1300 x*g* for 5 min at 4°C. Finally, the pellets were incubated in 500 µl of lysis buffer (50mM HEPES pH 7.5, 250mM NaCl, 5mM EDTA, 1% NP-40,1mM DTT, 1mM PMSF, 1x protease inhibitor cocktail (EDTA-free, Roche)) for 40 min. Protein samples were subsequently boiled in 1X Laemmli buffer at 95°C for 5 min, separated on 4–15% SDS-PAGE precast protein gels (BIO-RAD), and analysed by western blot.

The subcellular fractionation in Fig. 6b and Extended Data Fig. 6f was performed as follows: MCF10A (WT, +/delT) and CAPAN-1 cells from an 80% confluent 60mm culture dish were pelleted, resuspended in 1 ml of buffer 1 (10mM Tris pH 7.4, 10mM NaCl, 3mM MgCl_2_) and centrifuged at 2500 x *g* for 5 min at 4°C. The cell pellets were incubated with 200μl of buffer 2 (10mM Tris 1M pH 7.4, 10mM NaCl, 3mM MgCl_2_, 10% Glycerol, 0.5% NP-40, 0.5mM DTT, 1X protease inhibitor cocktail (EDTA-free; Roche) for 3 min and centrifuged at 5250 x *g* for 5 min at 4°C. The supernatant was collected in a new tube (cytoplasmic fraction), and the pellet was resuspended in 250μl lysis buffer (50mM HEPES pH 7.5, 1mM DTT, 1mM MgCl_2_, 150mM NaCl, 1% NP-40, 0.1 % Triton X-100, 100U Benzonase, and 1X protease inhibitor cocktail (EDTA-free; Roche)) and lysed for 30 min on ice in a new tube. The nuclear lysates were further sonicated once at 50 amplitudes for 20 sec. using a probe sonicator (Sonics & Materials, Inc.). Protein samples were then prepared as above for western blot.

### Antibodies for western blot and immunofluorescence

Mouse anti-BRCA2 (1:500, OP95, EMD Millipore), rabbit anti-N-ter-BRCA2 (raised for this study using BRCA2 aa 188-563 as antigen; 1:1000, GenScript), rabbit anti-RAD51 (1:2000, Abcam), mouse anti-alpha Tubulin (1:5000, Euromedex), rabbit anti-histone H3 (1:2000, Euromedex), rat anti-Vimentin (1:1000, MAB2105, RyD), mouse anti-histone H4 total (1:1000 Cell Signaling), (rabbit anti-histone H4 (acetyl K5 + K8 + K12 + K16)(abcam,1:5000), mouse anti-Ki67 (1:2000, 8D5, Cell Signaling), mouse anti-NF-kB p65 (1:1000, Cell Signaling Technology), rabbit anti-NF-kB p65 (phospho S536, 1:2000, abcam). Mouse anti-PCAF (1:500, Santa Cruz, sc-13124), mouse anti-GFP (1:1000, Roche, 11814460001), rabbit anti-RFP (1:1000, Enzo, AB233). Secondary antibodies: Horseradish peroxidase (HRP) conjugated secondary antibodies used: goat anti-mouse IgG-HRP (1:10,000, Jackson ImmunoResearch), goat anti-rabbit IgG-HRP (1:10,000, Jackson ImmunoResearch). Alexa Fluor 488 Donkey anti-Mouse IgG (H+L) (1:500, Thermo Fisher A-21202), Alexa Fluor 594 Donkey anti-Rat IgG (1:500, Thermo Fisher A-21209), DAPI (Merck 508741).

### Plasmid transfection and RNAi silencing

siRNA transfections were performed in a serum-containing medium with the transfection reagent JetPrime (Ozyme) following the manufacturer’s instructions. For BRCA2 silencing, we transfected 20 nM of BRCA2 siRNA (Dharmacon D-003462-0, GAAGAAUGCAGGUUUAAUA). The non-targeting siRNA #4 (Dharmacon D-001810-04-20, 20 nM) was used as a control (siCTRL) in the cells. Experiments were performed 48h after transfection.

PCAF-GFP plasmid and GFP control vector (kind gift from Prof. Kyle Miller) were transfected using TurboFect™ (Thermo Fisher Scientific Inc.) using a reverse transfection protocol per the manufacturer’s instructions. Experiments were performed 6 days after a single round of transfection.

### Immunoprecipitation

Endogenous full-length or truncated BRCA2 was immunoprecipitated from MCF10A, +/delT, or CAPAN-1 cells as follows: For MCF10A, cell pellets were resuspended in 1 ml Buffer 1 (10mM Tris pH 7.4, 10mM NaCl, 3mM MgCl_2_) and incubated on ice for 1 min followed by centrifugation at 2,500 *xg* for 5 min. The pellets containing the nuclear fractions were resuspended in 100-200μl of Buffer 2 (10mM Tris 1M pH 7.4, 10mM NaCl, 3mM MgCl_2_, 10% Glycerol, 0.5% NP-40, 0.5mM DTT) and centrifuged at 5,250 *xg* for 5 min. The pellets were then lysed in 1ml lysis buffer (50mM HEPES pH 7.5, 1mM DTT, 1mM MgCl_2_, 150mM NaCl, 1% NP-40, 0.1 % Triton X-100, 250U Benzonase, and 1X protease inhibitor cocktail (Complete, EDTA-free; Roche)) for 30 min at 4^°^C under rotation. For CAPAN-1, total cell lysates were used; cell pellets were lysed in 1ml of lysis buffer for at least 30 min at 4° under rotation. After clearing the lysates by centrifugation at 16,000 *xg* for 15 min, they were divided into two halves. One half was incubated with 5μg of BRCA2 antibody (BRCA2 N-terminal (in-house)) and the other with 5μg rabbit IgG (Thermo Fisher Scientific, 02-6102) overnight under rotation at 4^°^C. BRCA2 complexes were captured using 50μl Dynabeads Protein G (Life Technologies) for 2h under rotation at 4^°^C. The beads were collected using a magnet and washed five times with wash buffer 1 (50mM HEPES pH 7.5, 1mM MgCl_2_, 250mM NaCl, 1% NP-40, and 0.1 % Triton X-100) in the case of the MCF10A, or once with wash buffer 1 and once with wash buffer 2 (50mM HEPES pH 7.5, 1mM MgCl_2_, 500mM NaCl, 1% NP-40, and 0.1 % Triton X-100) in the case of CAPAN-1 cells. Protein complexes were eluted by boiling the beads in 3X Laemmli buffer, separated by 4–15% SDS-PAGE precast protein gels (BIO-RAD), and analysed by western blot.

### GFP-TRAP

A 10-cm plate of HEK293T cells was transiently transfected with Turbofect and the following plasmids: GFP-MBP-NLS, GFP-MBP-BRCA2(WT or the delT) together with MBP-BRCA2-WT-RFP (5μg each), as indicated in the legend, following the manufacturer’s specifications. 36h later, cells were collected and lysed in co-IP lysis buffer (50mM HEPES pH 7.5, 1mM DTT, 1mM MgCl_2_, 150mM NaCl, 1% NP-40, 0.1 % Triton X-100 including 1x protease inhibitor cocktail (EDTA-free) and 250 U Benzonase. Samples were incubated for 30 min at 4°C under rotation. Total protein in the lysates was quantified by Bradford assay. A 50 μg input sample was collected in a separate tube and mixed with 3x Laemmli buffer. The samples were cleared by centrifugation at 20,000 *xg* for 15 min and the cleared lysates were subjected to GFP pull-down with 25 μl of GFP-Trap beads (Chromotek) for 90 min at 4°C under rotation. The beads were then washed trice with wash buffer (50mM HEPES pH 7.5, 1mM MgCl_2_, 250mM NaCl, 1% NP-40, and 0.1 % Triton X-100) and boiled for 5 min at 95°C in 3x Laemmli buffer along with the input samples. Proteins were separated in SDS-PAGE 4-15% precast gels (BIO-RAD) and analyzed by western blot using an Odyssey Infrared Imaging System (Licor; V3.0) and Image Studio Lite (version 5.2) except for PCAF that was detected using ECL prime and Amersham Imager 680.

### PARPi, MMC and HU sensitivity assessed by MTT assay

Cell viability was assessed in MCF10A parental cells or MCF10A BRCA2-mutated clones using MCF10A transfected with either siCTRL or siBRCA2 as controls, as indicated in the legends.

Olaparib (AZD2281, Selleck Chemicals): Cells were seeded in triplicate in 96 well-plates (TPP). The following day they were treated at increasing concentrations of the drug (0.5, 1.0, and 2.5 µM) for 6 days. MMC (Sigma-Aldrich) and HU (Sigma-Aldrich): 3×10^5^ cells were seeded in a 6 well-plate (TPP). For MMC, cells were treated for one hour with the following concentrations of the drug: 0, 1.0, 2.0, and 4.0 µM. For HU, cells were treated for 24h with the following concentrations of the drug: 0, 2.5, and 5 mM. Subsequently, cells were harvested and seeded in triplicate in 96 well plates. On the 6^th^ day, the media was removed and cells were washed with 1x PBS. Cell viability was assessed with 3-[4,5-Dimethylthiazol-2-yl]-2,5-diphenyltetrazolium bromide (MTT, #M5655, Sigma Aldrich). The solution was removed and MTT crystals were dissolved in 100µl 100% DMSO (Sigma-Aldrich). The absorbance was obtained from the Trista 3 microplate reader (Berthold Technologies) at 570 nm. The relative surviving cells were calculated by dividing the absorbance of the treated cells by the absorbance obtained in the untreated condition of the same clone.

In the case of the EGF survival assay (Extended Data Fig. 6c), 2,000 MCF10A and +/delT cells were seeded in 96-well plates in MCF10A normal media (containing 20 ng/ml EGF) or in the same media without EGF. After 72 h incubation at 37°C, cell viability was assessed as above using a FLUOstar OPTIMA microplate reader (BMG Labtech). The relative surviving cells were normalized against the MCF10A parental cell line in conditions of normal media (+EGF).

### Clonogenic survival to assess sensitivity to HU

Cells seeded at 70% of confluence in a 6-well plate were treated with HU (Sigma-Aldrich) at concentrations 0, 2.5, 5, or 10 mM. After 24 h treatment, the cells were serially diluted in complete growth media (Eurobio) and seeded at 500, 1000, 2000, and 4000 cells in triplicates in 60 mm dishes depending on the drug concentration. The media was changed every third day, after 14 days in culture the plates were stained with crystal violet (Sigma-Aldrich), colonies were counted and the surviving fraction was determined for calculated as the ratio of the number of colonies formed to the total number of cells plated.

### Cell-based homologous recombination assay

2.5 × 10^6^ cells for control, 3× 10^6^ for BRCA2-mutated clones, were seeded in 10 cm plates. The following day, cells were detached and counted. 1×10^6^ cells were mixed in a total amount of 400 µl of Optimem (Gibco) with 5 µg of each of these three plasmids: spCas9-GFP, AAVS1-gRNA, and the plasmid coding for the mCherry protein flanked with two regions of homology for the AAVS1 site. For control cells, AAVS1-gRNA (gRNA) plasmid was excluded from the mix. Cells were electroporated in a 0.4 µm cuvette with a Gene Pulser (BioRad) at 350V and 960 µF, and seeded into a P6 well plate. 2 days later, cells were detached and GFP+ cells (expressing spCas9) were sorted using FACS-Aria Fusion Sorting Cytometer. The same number of GFP+ sorted cells were seeded for each condition. 6 days after (8 days from electroporation), the percentage of mCherry+ of cells was quantified using a FACS-Aria Fusion Cytometer. The oligonucleotides with the target sequence used for the CRISPR are indicated in Table S5.

### DNA combing assay with S1 treatment to assess PARPi-induced ssDNA gaps

Cells were seeded in a 10 cm plate and allowed to adhere for 24 h (2 × 10^6^ cells for control, 3.5× 10^6^ for BRCA2-mutated clones). The following day, DNA was labeled for 30 min with 100 µM IdU (Sigma-Aldrich) and washed twice with PBS followed by simultaneous incubation for 2h with 100 µM CIdU (Sigma-Aldrich) and 30 µM Olaparib (AZD2281, Selleck Chemicals). After labeling, cells were washed twice with PBS and permeabilized with 5 ml CSK buffer in a 10 cm dish (10 mM PIPES, pH 6.8, 0.1 M NaCl, 0.3 M sucrose, 3 mM MgCl_2_, EDTA-free Protease Inhibitor Cocktail (Roche)) at room temperature for 10 min. Subsequently, cells were incubated with 1ml of S1 buffer (30 mM sodium acetate pH 4.6, 1 mM zinc sulfate, 50 mM NaCl) was added with or without S1 nuclease (20 U) (Thermofisher # 18001016) for 30 min at 37 °C. Finally, cells were collected by scraping, pelleted, and resuspended in PBS (45 µl per 400,000 cells).

500 µl of cells were transferred to a new tube, briefly heated at 42 °C, and resuspended with 500 µl melted 2% agarose type VII (SIGMA) to make the agarose plugs. Plugs were left to solidify for 20 min at 4 °C and were then digested with Proteinase K (400 mM EDTA pH 8, 10% Proteinase K, 1% Sarcosyl) in a water bath at 42 °C overnight. The next day, the plugs were washed trice with TE 1X buffer. TE solution was removed and a solution containing 50 mM MES pH 5.5,100 mM NaCl was added to the plugs that were heated at 68 °C for 20 min. Agarose plugs were then dissolved by adding 2 µl of β-agarase (NEB) and incubated at 42 °C overnight. The following day, dissolved agarose plugs were transferred to the combing machine (Genomic Vision) where DNA was combed onto silane-coated coverslips (Genomic Vision COV-002-RUO) following the manufacturer’s specifications. Combed coverslips were baked overnight at 60 °C, denatured in denaturation buffer (25 mM NaOH, 200 mM NaCl in H2O) for 15 min, washed trice with 1X PBS, and dehydrated by incubation with increasing concentrations of ethanol 70%, 90%, and 100% each for 5 min. For the IF staining, the coverslips were incubated with BlockAid for 15 min (Life Technologies) at RT followed by the primary anti-IdU and anti-CldU antibodies (1 h, 1:25 anti-mouse Becton Dickinson 347580 for IdU and 1:50 anti-rat Abcam ab6326 for CIdU) and then incubated 1 h with the following secondary antibodies: 1:50 Alexa donkey anti-mouse 488 (Life Technologies ref. 21202), 1:50 Alexa goat anti-rat 555 (Life Technologies ref. A21434) in BlockAid (Thermo Scientific). Slides were air-dried for 5-10 min and were mounted with 7 µl mounting media (80% Glycerol and 20% PBS) and sealed with clear nail polish. Track lengths were measured in Fiji and the ratio was calculated by dividing the IdU (green) by the CldU (red) signal.

### Metaphase spreads

12,500 MCF10A cells were seeded into glass coverslips in P24 wells. The day after, cells were treated with 10 µM of Olaparib for another 24h. Cell division was blocked with 100 ng/ml colcemid (Thermo Fisher) for 3h. Then, a hypotonic shock was performed with 1:7 FBS in distilled water for 40 min. after which the cells were fixed adding 1 volume of 3:1 Methanol-Acetic Acid for 15 min at RT, then exchanged with 3:1 Methanol-acetic acid for 15 min at RT and then for 30 min at 4°C. Coverslips were air-dried and stained with 2% Giemsa solution (Thermo Fisher) diluted in Gurr buffer (Thermo Fisher) for 10 min. The coverslips were rinsed in water and air-dried at RT, then mounted with Eukitt mounting media (Sigma-Aldrich). Images were acquired with a 100X objective with a CMOS camera (LEICA).

### Cell cycle analysis by flow cytometry

1×10^6^ cells were harvested and washed twice with PBS. Cells were then fixed in ice-cold 70% ethanol overnight, before being pelleted, resuspended in ice-cold PBS, and then incubated in the dark in propidium iodide (PI) staining buffer (3.5 mM Tris HCl, pH 7.6, 50 μg/ml PI, 50 μg/ml RNase A, 0.1% NP-40, 10mM NaCl, ddH20) for 30 min at room temperature. Cell cycle distribution was analyzed by flow cytometry BD FACSAria III (BD Bioscience) using the FACSDiva software and data were analyzed with the FlowJo 10.5 software (Tree Star Inc.).

### γH2AX and RAD51 immunofluorescence staining and quantitative image-based cytometry (QIBC)

Cell treatments: MCF10A WT, +/delT, and +/del5 cells were treated with DMSO as control, 10 μM PARPi olaparib (Selleckchem, S1060), or 3 μM mitomycin C (Sigma-Aldrich, M7949) for 24h as indicated.

Immunofluorescence staining: Cells were grown on µ-Plate 96 well plates (ibidi, 89626), fixed in 3% formaldehyde in PBS for 15 minutes at RT, washed twice in PBS, permeabilized for 5 minutes at room temperature in 0.2% Triton X-100 (Sigma-Aldrich) in PBS, and washed twice in PBS. Primary and secondary antibodies were diluted in filtered DMEM containing 10% FBS and 0.02% sodium azide. Incubations in primary and secondary antibodies were performed at room temperature for 2h and 1h, respectively, with three washes in PBS in between. Cells were then washed once with PBS and incubated for 10 minutes at room temperature with PBS containing 4′,6-diamidino-2-phenylindole dihydrochloride (DAPI, 0.5 µg/ml) to stain DNA. Cells were subsequently washed three times in PBS before imaging. The following primary antibodies were used: RAD51 (rabbit, Bioacademia 70-002, 1:1000), H2AX Phospho S139 (mouse, Biolegend 613401, 1:1000). Secondary antibodies were: Alexa Fluor 488 Goat Anti-Rabbit (Life Technologies, A11034, 1:500), Alexa Fluor 568 Goat Anti-Mouse (Life Technologies, A11031, 1:500).

Automated multichannel widefield microscopy for quantitative image-based cytometry (QIBC) was performed as described previously^65,66^ on an Olympus ScanR Screening System (ScanR Image Acquisition 3.01) equipped with an inverted motorized Olympus IX83 microscope, a motorized stage, IR-laser hardware autofocus, a fast emission filter wheel with one set of bandpass filters for multi-wavelength acquisition (DAPI (ex BP 395/25, em BP 435/26), FITC (ex BP 470/24, em BP 511/23), TRITC (ex BP 550/15, em BP 595/40), Cy5 (ex BP 640/30, em BP 705/72)), and a Hamamatsu ORCA-FLASH 4.0 V2 sCMOS camera (2048 x 2048 pixel, pixel size 6.5 x 6.5 µm) with a x20 UPLSAPO (NA 0.75) air objective. For each condition, image information of large cohorts of cells (typically at least 1000 cells for the UPLSAPO 20x objective (NA 0.75)) was acquired under non-saturating conditions, and identical settings were applied to all samples within one experiment. Images were analyzed with the Olympus ScanR Image Analysis Software (version 3.3.0), a dynamic background correction was applied, and nuclei segmentation was performed using an integrated intensity-based object-detection module based on the DAPI signal. Nuclear foci segmentation was performed using an integrated spot-detection module. All downstream analyses were focused on properly detected interphase nuclei containing a 2N-4N DNA content as measured by total and mean DAPI intensities. Color-coded scatter plots of asynchronous cell populations were generated with Spotfire data visualization software (version 10.10.1.7, TIBCO). Within one experiment, similar cell numbers were compared for the different conditions. For visualization of discrete data in scatter plots, mild jittering (random displacement of data points along the discrete data axes) was applied to demerge overlapping data points. Representative scatter plots and quantifications of independent experiments are shown.

### Spheroid invasion assay

Around 60 20 μl drops containing 3000 cells resuspended in the medium were seeded to form spheroids of MCF10A or +/delT cells by hanging drop method on the underside of a 10 cm petri dish lid. The lid was then inverted over the dish (filled with PBS). 24h after incubation at 37°C, drops were carefully collected with a P1000 tip (∼150 μl), and mixed 1:1 with a pH-adjusted solution of Collagen type 1 (Corning) to a final concentration of 1.5 mg/ml. The 300 μl mix was seeded in a well of a 24-well plate. After 1h of incubation at 37°C, 500 μl of complete media was added on top of the collagen gel. The day after the spheroids were imaged using a Leica Flexicam C1 camera, and both, the area of the spheroid, and the area of spheroids+protusions (total spheroid area) were quantified drawing a line around the limit of the spheroid or the protrusions using ImageJ software. We used the ratio of total area/spheroid area to compare the invasion capacity.

### Whole genome sequencing

The internal review boards of each participating institution approved the collection and use of samples of all patients in this study. Informed consent was obtained by the relevant participating institution.

DNA was extracted from both tumor and corresponding normal tissue and samples were subjected to whole-genome sequencing. The resulting BAM files were aligned to the reference human genome (GRCh37) using the Burrows-Wheeler aligner, BWA (v0.5.9). Mutation calling was performed as described previously^44,45^. Briefly, CaVEMan (Cancer Variants Through Expectation Maximization; http://cancerit.github.io/CaVEMan/) was used to call somatic substitutions. Indels in the tumor and normal genomes were called using modified Pindel version 2.0 (http://cancerit.github.io/cgpPindel/) on the NCBI37 genome build. Structural variants were discovered using a bespoke algorithm, BRASS (BReakpoint AnalySiS) (https://github.com/cancerit/BRASS) through discordantly mapping paired-end reads followed by de novo local assembly using Velvet^67^ to determine exact coordinates and features of breakpoint junction sequence.

### Extraction of mutational signatures

Mutational signatures analysis was performed following a three-step process as previously described^44^: (i) hierarchical de novo extraction based on somatic substitutions and their immediate sequence context, (ii) updating the set of consensus signatures using the mutational signatures extracted from breast cancer genomes, and (iii) evaluating the contributions of each of the updated consensus signatures in each of the breast cancer samples.

The complete compendium of consensus mutational signatures that were found in our cohort of breast cancers with the two pathogenic variants includes signatures 1, 2, 3, 5, 8, 13, 17, 18, and 127 (Fig. 3, Extended Data Fig. 3). Mutational signatures that did not contribute large numbers (or proportions) of mutations or that did not significantly improve the correlation between the original mutational pattern of the sample and the one generated by the mutational signatures were excluded from the sample. This procedure reduced the over-fitting of the data and allowed only the essential mutational signatures to be present in each sample.

### HRD score

The HRD score determination was conducted using the HRDetect predictor^44^, considering the following genomic features, listed in their respective order of importance: deletions with microhomology, substitution signature 3, rearrangement signatures 3 and 5, HRD index, and substitution signature 8. For further details, see^44^.

### Allele-specific LOH status

For the BRCA2-del5 Icelandic cohort (published + 1 new)^15,17^ the proportion of the BRCA2 wild-type (wt) alleles was quantitatively measured relative to the c.999del5 BRCA2 mutated alleles in tumor DNA samples by quantitative polymerase chain reaction (qPCR) (7500 Realtime PCR system; Applied Biosystems) with a TaqMan method as previously described^15,17,23^. Briefly, using a single forward primer and two different reverse primers, one designed to amplify the wild-type allele and another for the *BRCA2 999del5* allele, and a FAM-labelled TaqMan probe, the average Ct value was determined by duplicate qPCRs for the two primer pairs in 2 separate reactions. When the *BRCA2* wild-type allele proportion was below 33% the sample was defined as presenting *BRCA2* LOH. Some of these results were further confirmed by array comparative genomic hybridization (aCGH) (Table S1).

For the BRCA2-delT Penn cohort (n =2 + published)^13^, a combination of VarScan2^68^, allele frequency comparisons, and allele-specific copy number calls were used to determine BRCA locus-specific LOH. Estimates of tumor purity (cellularity) were determined using Sequenza and inputted into VarScan2 variant calling. The sample was assigned a locus-specific LOH positive status if the VarScan2 somatic P-value was significant and a locus-specific LOH negative status if the VarScan2 germline P-value was significant. Allele-specific copy number calls of the genomic region containing the BRCA2 mutation were determined by Sequenza. The copy number of the genomic region surrounding the germline BRCA2 mutation (CN) and the number of mutant (m) alleles as per the output of the Sequenza program were used to assign two states of absent locus-specific LOH—heterozygous diploid (CN = 2;m = 1) or amplified with gain of non-mutant (wildtype) allele (CN>2;m = 1)—and three states of locus-specific LOH—loss (CN = 1;m = 1), copy neutral LOH (CN = 2;m = 2), and amplified with LOH (CN>2;m>2). The genomic regions surrounding the germline BRCA2 mutation ranged from less than one to over 100 Mb in length. In cases where the VarScan2 and ASCN calls differed, the difference between cellularity-corrected tumor allele frequency and blood allele frequency (ΔAF) was determined; the sample was assigned a locus-specific LOH positive status if ΔAF>0.2028.

For the I. Curie cohort, raw sequences were generated per sample using the PCR Free Roche KAPA Hyper Prep Library on the Illumina Novaseq 6000 (run D458) at the NGS platform of the Institut Curie, resulting in the creation of 150bp paired-end fastq files. Raw data were then processed using the Institut Curie raw-qc pipeline (10.5281/zenodo.8340106). The overall quality of the reads was checked using FastQC (v0.11.8). Reads were then aligned on the hg19 reference genome using bwa mem (v.0.7.15). Aligned data were processed using Facets (v0.5.1) that take advantage of both the B-allele ratio (BAF) and the read depth to estimate the allele-specific copy number variants and the optimal ploidy and tumor cellularity, after a GC percentage correction. Total copy number (number of A and B in the genotype) and minor copy number (number of B in the genotype) were defined for each segment. LOH was considered for segments with a minor copy number of 0.

### Statistical analysis

Statistical analyses were performed using GraphPad Prism (Version 10.0). Statistical parameters and tests are reported in the figures and corresponding figure legends. The number of samples (n) in figure legends represents independent biological replicates unless stated otherwise. The statistical significance was obtained from the mean of the independent experiments. No statistical methods were used to determine the sample size before starting experiments.

## Data and materials availability

All materials related to this study are available upon request from the corresponding author. Any further inquiries or requests for resources and reagents should be directed to the corresponding author, Raw sequence files listed in Table S7 have been deposited at the European Genome-Phenome Archive with accession numbers EGAS50000000613 and EGAS50000000614.

## Extended Data Figures

**Extended Data Fig. 1.**
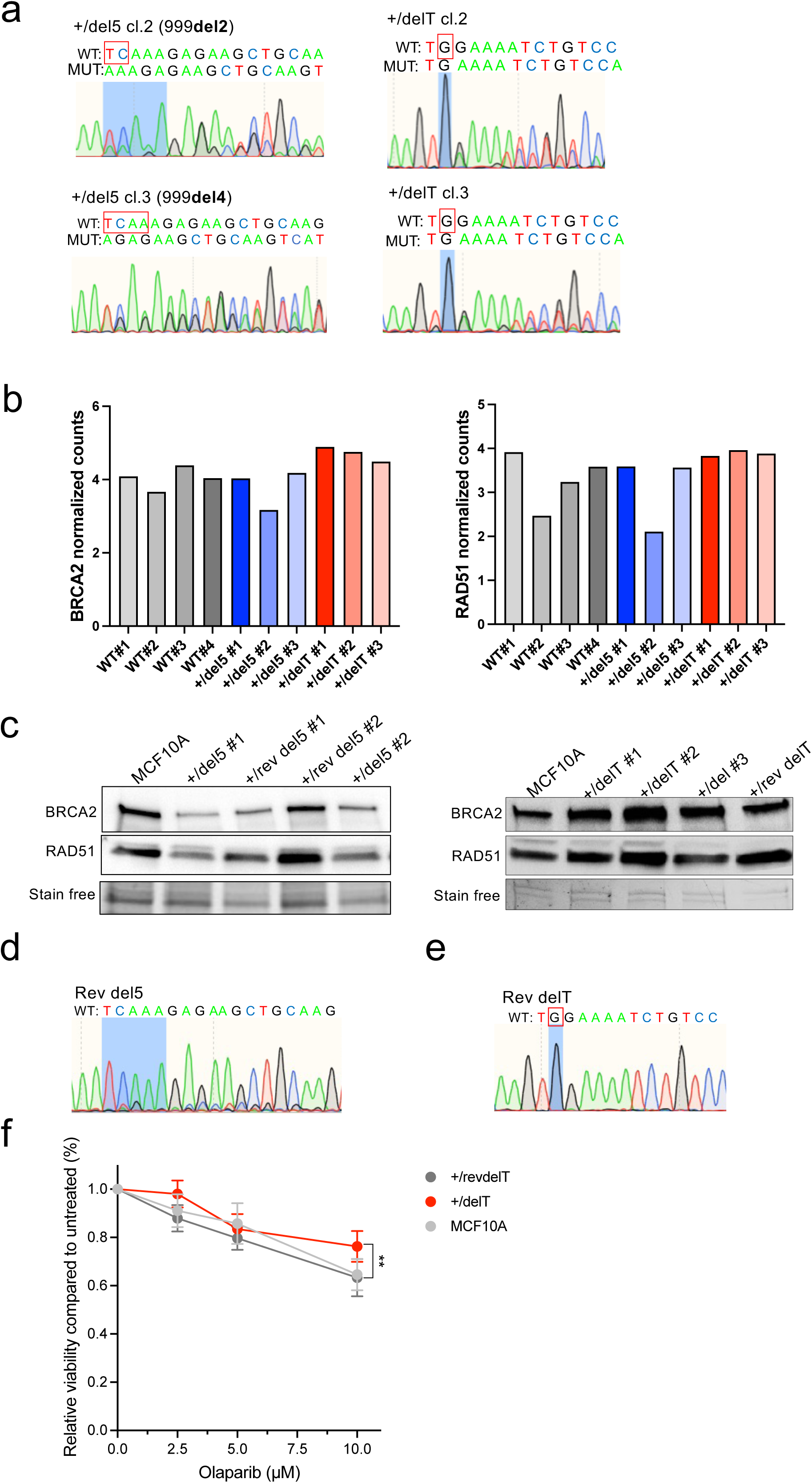
**a,** Sanger sequencing chromatograms confirming the mono-allelic BRCA2 (**left**) del5 and (**right**) delT mutation for the additional biological clones used for the screen. **b,** Normalized RNAseq counts for BRCA2 and RAD51 mRNA levels in the 3 biological replicates of MCF10A (WT#1, 3, 4), +/del5 (#1-3) and +/delT (#1-3). **c,** BRCA2 and RAD51 protein levels for all the clones used in this study. **d,** Sanger Sequencing chromatogram confirming the correction of the BRCA2 del5 mutation to WT. **e,** Sanger sequencing chromatogram showing the correction of the BRCA2 delT(delG) mutation to WT. **f,** Quantification of the relative cell viability of MCF10A, +/delT and +/revdelT monitored by MTT assay upon treatment with increasing doses of the PARP inhibitor Olaparib after 6 days. The data represent the mean ± SD of four independent experiments. Statistical difference was determined by a Kruskal-Wallis test followed by Dunn’s multiple comparison test comparing to the parental MCF10A cells. Only significant values are shown. ** p < 0.01.

**Extended Data Fig. 2.**
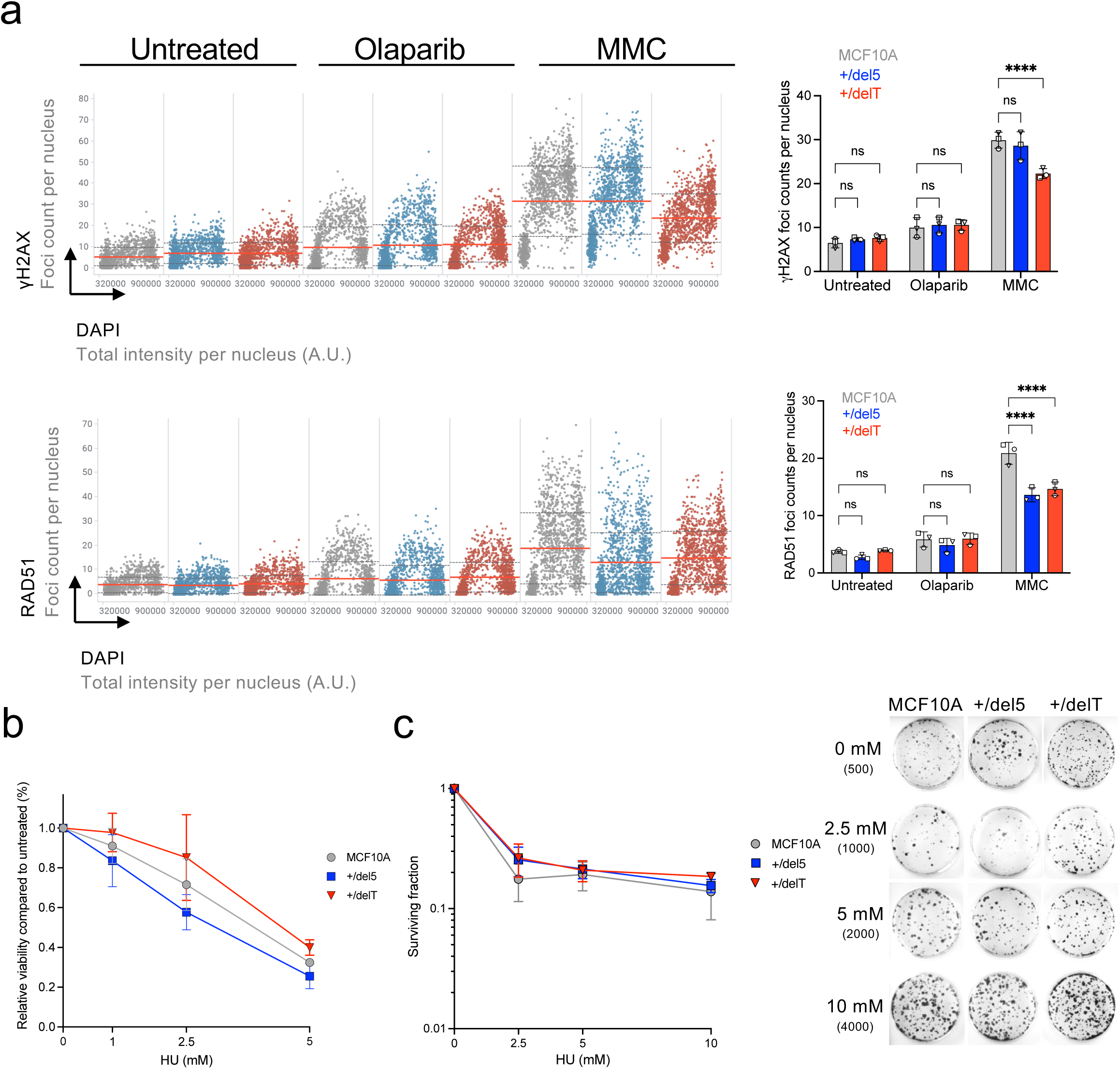
**a,** (**Left**) Representative QIBC experiment to monitor γH2AX (top) and RAD51 foci (bottom) over DAPI intensity in MCF10A clones treated for 24 h with 10 μM of Olaparib or 3 μM of MMC. (**Right**) Quantification of γH2AX (top) and RAD51 foci (bottom). Foci counts per nucleus were analyzed from at least 1000 cells per condition. Data represent counts from three independent experiments and values indicate the mean + SD. Statistical difference was determined by 2-way ANOVA. The p-values show significant differences compared to the MCF10A clone: **** p < 0.0001, ns, not significant. **b,** Quantification of the relative cell viability monitored by MTT assay. Cells were treated 4h with increasing doses of HU as indicated, followed by one day release. After seeding, viability was monitored after 6 days (n=2). **c,** (**Left**) Quantification of the relative cell viability monitored by clonogenic survival assay. Cells were treated 24 h and then seeded. Colonies were stained after 14 days. The data represent the mean ± SD of three independent experiments. Statistical difference in **b** and **c** was determined by a two-way ANOVA test with Dunnett’s multiple comparisons test. Only significant values are shown. **(Right)** Representative images of a clonogenic survival experiment at the indicated seeding densities, shown in brackets, and HU treatment conditions.

**Extended Data Fig. 3.**
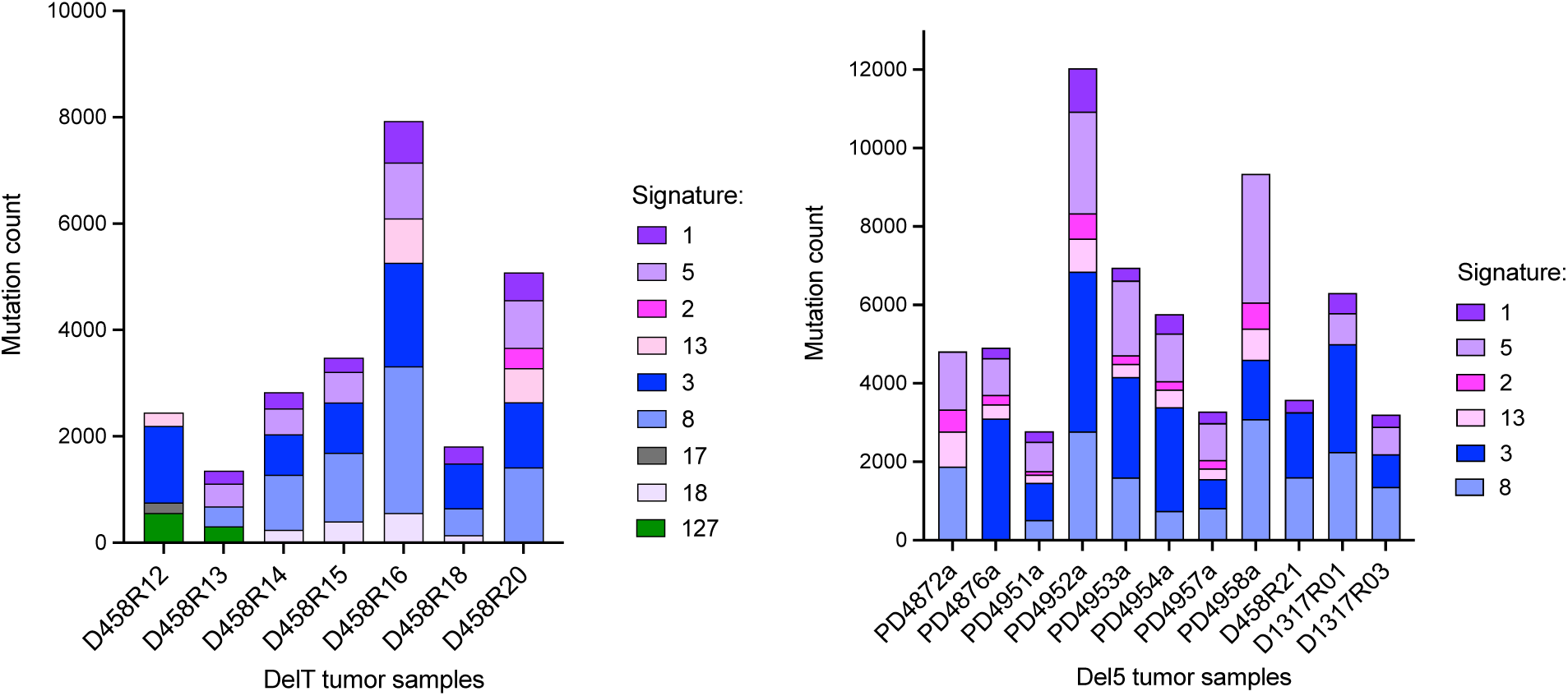
Number of base substitution mutations contributing to each signature (legend on the right) for the 7 individual primary tumors bearing the BRCA2delT variant (**left**) and the 11 expressing the BRCA2 del5 variant (**right**).

**Extended Data Fig. 4.**
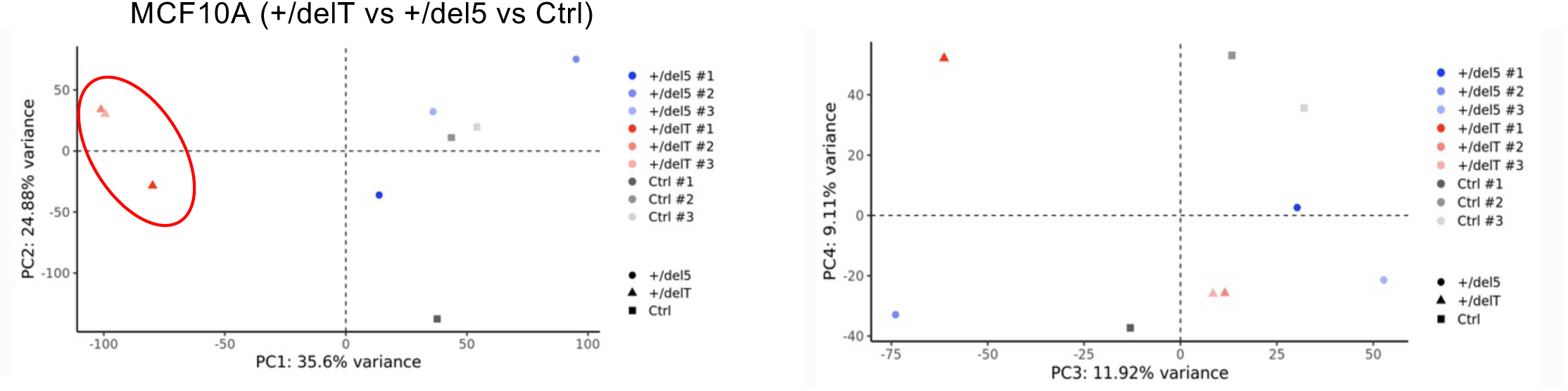
Principal component analysis (PCA) showing the clustering of the gene expression profile of 3 biological replicates (from three different isolated clones) of the MCF10A control (grey) vs +/delT (red) vs +/del5 (blue). DEGs were obtained applying a log2 fold change > 1.5.

**Extended Data Fig. 5.**
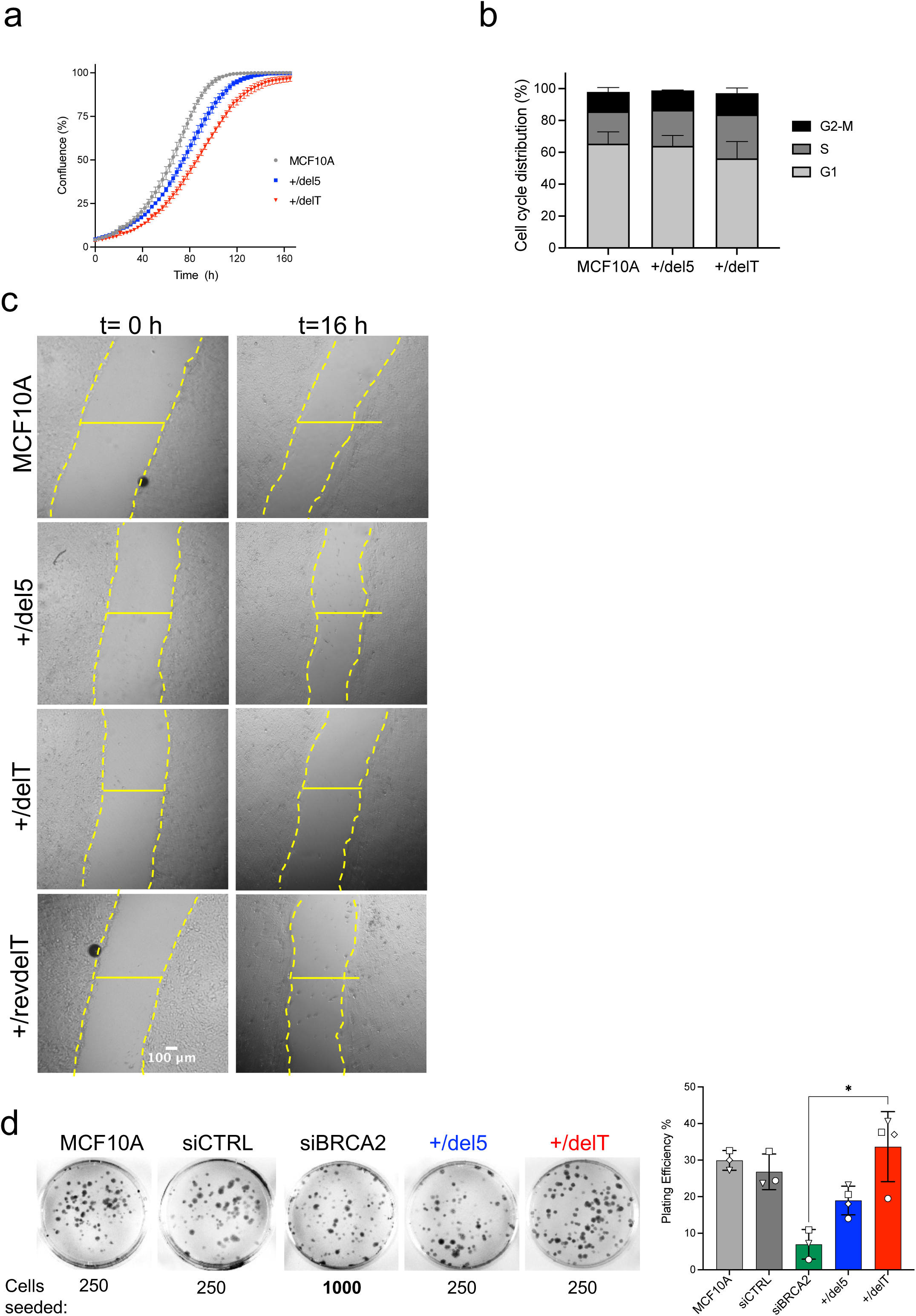
**a,** Representative curves of the mean + SEM of the confluence over time of three technical replicates of the indicated MCF10A cell lines monitored in an Incucyte system. **b,** Histogram showing the percentage of cells in the G1, G1, S and G2/M phase of the cell cycle obtained after FACS analysis from three independent experiments. For each sample 10,000 cells were acquired. Error bars show SD. Statistical difference was determined by a two-way ANOVA test with Dunnett’s multiple comparisons tests. Only significant values are shown. **c,** Representative 4X brightfield images of MCF10A, +/del5, +/delT and +/revdelT wound healing assay at t=0h and t=16h quantified in Fig. 5D. Yellow dashed lines delineate the scratch, and a yellow line of equal size was drawn across the wound to visualize the difference in size at the two time points. **d,** (**Left**) Representative plates showing surviving colonies for the cell lines indicated. Cells were seeded in triplicates and the number of cells seeded is shown below each image. Colonies were stained with crystal violet after 7 days in cell culture. (**Right**) Bar graph showing the quantification of the fraction of surviving colonies from three (MCF10A, siCTRL and siBRCA2) or four (+/del5 and +/delT) independent experiments. The different experiments are represented with different symbol shapes. Statistical difference was determined by a Kruskal-Wallis test with Dunn’s multiple comparisons tests. * p = 0.0287. Only significant p-values are shown.

**Extended Data Fig. 6.**
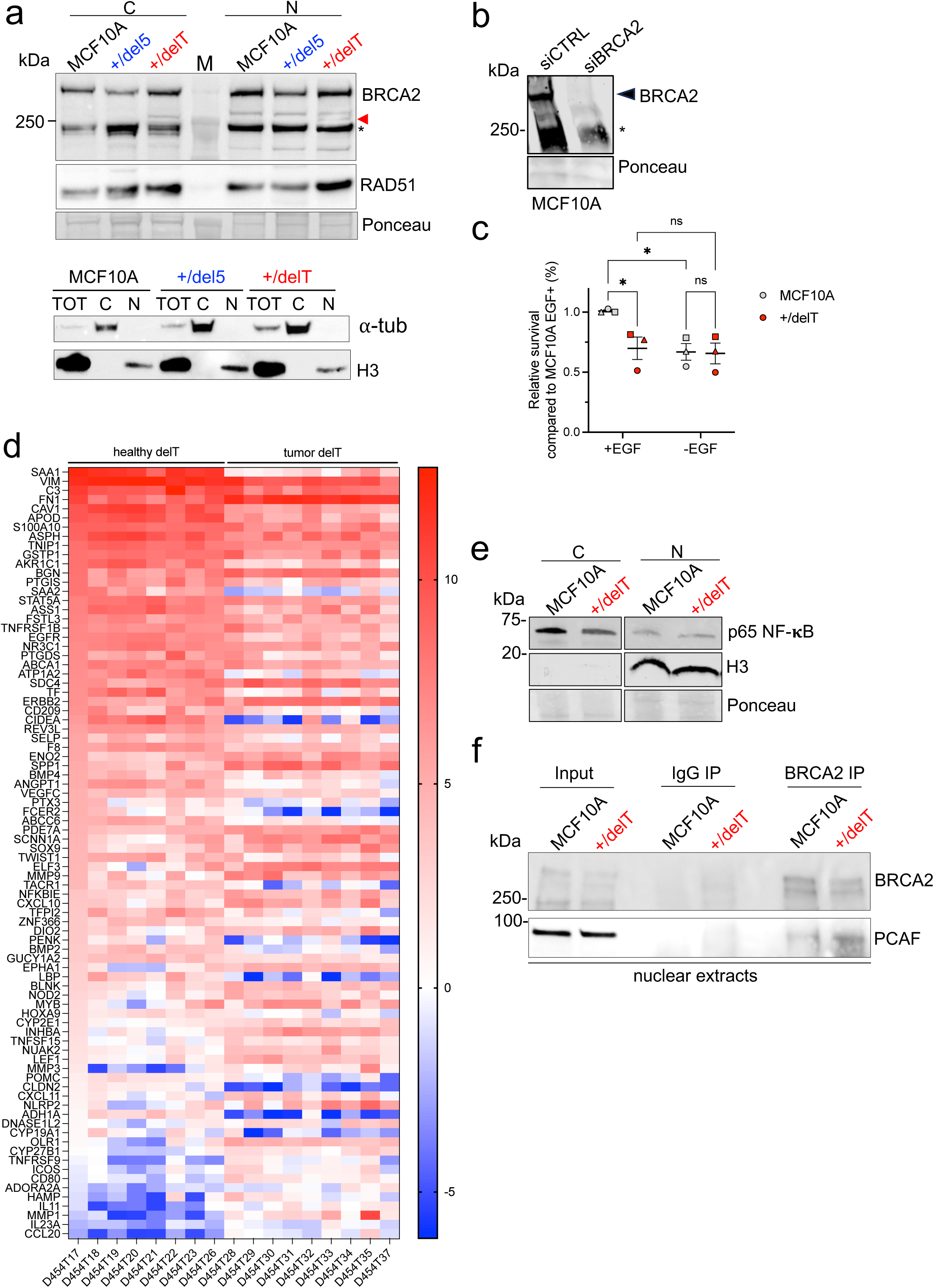
**a, (Top).** Immunoblot showing nuclear/cytoplasmic fractionation of the MCF10A, +/del5 and +/delT cells. The red wedge denotes a band with the size (predicted for the truncated BRCA2-delT product); * indicates a non-specific band. (**Bottom**). Western blot analysis of the samples above using α-tubulin and histone H3 as cytoplasmic/nuclear markers, respectively. TOT, total cell lysate, C, cytoplasm, N, nuclear fraction. (n=2) **b,** Validation of the BRCA2 N-terminal antibody used in cells transfected with siRNA control or BRCA2 siRNA. * Indicates a non-specific band. **c,** Quantification of survival by MTT assay of MCF10A and +/delT cells cultured in normal media (+EGF) or without EGF (-EGF) for 72h. The means of three independent biological experiments are plotted. Two-way-ANOVA followed by Fisher LSD test was performed on the means. ns, not significant, * p= 0.016 (MCF10A EGF+ vs +/delT EGF+), 0.010 (MCF10A EGF+ vs MCF10A EGF-). **d,** Heatmap showing the expression of NF-κB target genes in delT tumors versus their healthy controls, as indicated and assessed by RNAseq. One control sample (D454T24) was discarded because of suspected tumor contamination. **e,** Immunoblots showing the NF-κB (p65 subunit) levels in the cytoplasm (C) and nuclear (N) fractions of MCF10A (WT, +/delT). H3 immunoblot served as control of the nuclear fraction (n=2). **f,** Nuclear co-immunoprecipitation of BRCA2 in complex with PCAF in MCF10A or +/delT cells. BRCA2 and PCAF were detected by western blot using specific antibodies. Rabbit IgG was used as control (n=3).

**Extended Data Fig. 7.**
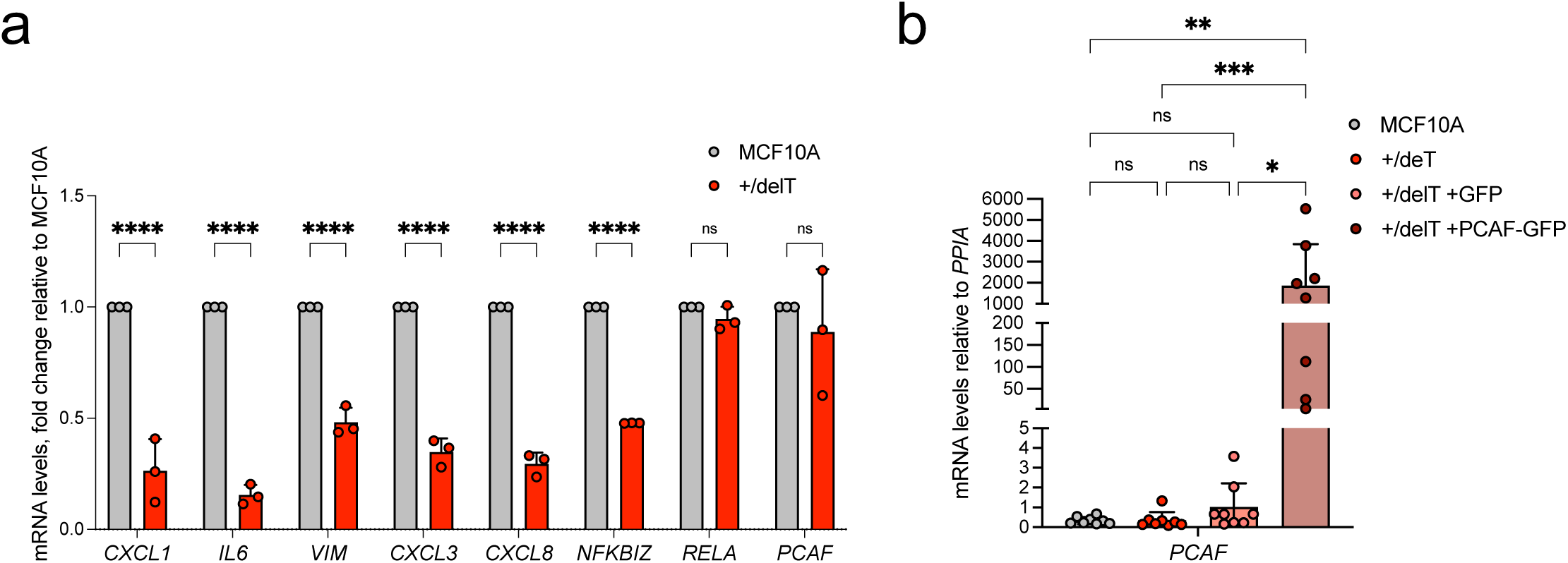
a, RT-qPCR of NF-κB regulated genes (*CXCL1*, *IL6, VIM, CXCL3, CXCL8, NFKBIZ*), *NF-κB* (*RelA*), and *PCAF* in MCF10A parental and +/delT cells. The data was first normalized using *U6snRNA* as a reference gene and then represented as the fold-change expression of the indicated genes relative to MCF10A cells (n=7). Statistical difference was determined by two-way ANOVA with Šídák’s multiple comparisons test. ****p<0.0001 and ns: not significant, p>0.05. **b,** RT-qPCR of *PCAF* in MCF10A, non-transfected +/delT cells, or +/delT transfected with GFP or PCAF-GFP, as indicated. The data was normalized using *PPIA* as a reference gene (n=8). MCF10A and non-transfected +/delT cells data for *PCAF* are the same as in Extended Data Fig. 7a. Statistical difference was determined by the Kruskal-Wallis test (with Dunn’s multiple comparisons test). *p=0.0500, **p=0.0026, ***p=0.0003, and ns: not significant, p>0.05.

## REFERENCES

1. Minello, A. & Carreira, A. BRCA1/2 haploinsufficiency: exploring the impact of losing one allele. J. Mol. Biol. 168277 (2023) doi:10.1016/j.jmb.2023.168277.

2. Sessa, G. et al. BRCA2 promotes R-loop resolution by DDX5 helicase at DNA breaks to facilitate their repair by homologous recombination. Embo J 40, e106018 (2021).

3. Schlacher, K. et al. Double-Strand Break Repair-Independent Role for BRCA2 in Blocking Stalled Replication Fork Degradation by MRE11. Cell 145, 529–542 (2011).

4. Lomonosov, M., Anand, S., Sangrithi, M., Davies, R. & Venkitaraman, A. R. Stabilization of stalled DNA replication forks by the BRCA2 breast cancer susceptibility protein. Gene Dev 17, 3017–3022 (2003).

5. Kolinjivadi, A. M. et al. Smarcal1-Mediated Fork Reversal Triggers Mre11-Dependent Degradation of Nascent DNA in the Absence of Brca2 and Stable Rad51 Nucleofilaments. Mol. Cell 67, 867–881.e7 (2017).

6. Panzarino, N. J. et al. Replication Gaps Underlie BRCA Deficiency and Therapy Response. Cancer Res. 81, 1388–1397 (2021).

7. Quinet, A. et al. PRIMPOL-Mediated Adaptive Response Suppresses Replication Fork Reversal in BRCA-Deficient Cells. Molecular cell 77, 461–474.e9 (2020).

8. Vugic, D. et al. Replication gap suppression depends on the double-strand DNA binding activity of BRCA2. Nat Commun 14, 446 (2023).

9. Daniels, M. J., Wang, Y., Lee, M. & Venkitaraman, A. R. Abnormal cytokinesis in cells deficient in the breast cancer susceptibility protein BRCA2. Science 306, 876–879 (2004).

10. Mondal, G. et al. BRCA2 localization to the midbody by filamin A regulates cep55 signaling and completion of cytokinesis. Dev. Cell 23, 137–152 (2012).

11. Ehlen, A. et al. Proper chromosome alignment depends on BRCA2 phosphorylation by PLK1. Nat. Commun. 11, 1–21 (2020).

12. Fuks, F., Milner, J. & Kouzarides, T. BRCA2 associates with acetyltransferase activity when bound to P/CAF. Oncogene 17, 2531–2534 (1998).

13. Maxwell, K. N. et al. BRCA locus-specific loss of heterozygosity in germline BRCA1 and BRCA2 carriers. Nat Commun 8, 319–11 (2017).

14. King, T. A. et al. Heterogenic Loss of the Wild-Type BRCA Allele in Human Breast Tumorigenesis. Ann. Surg. Oncol. 14, 2510–2518 (2007).

15. Stefansson, O. A. et al. Genomic and phenotypic analysis of BRCA2 mutated breast cancers reveals co-occurring changes linked to progression. Breast Cancer Res. 13, R95 (2011).

16. Heetvelde, M. V. et al. Accurate detection and quantification of epigenetic and genetic second hits in BRCA1 and BRCA2-associated hereditary breast and ovarian cancer reveals multiple co-acting second hits. Cancer Lett. 425, 125–133 (2018).

17. Aradottir, M. et al. Aurora A is a prognostic marker for breast cancer arising in BRCA2 mutation carriers. J. Pathol.: Clin. Res. 1, 33–40 (2015).

18. Karaayvaz-Yildirim, M. et al. Aneuploidy and a deregulated DNA damage response suggest haploinsufficiency in breast tissues of BRCA2 mutation carriers. Sci Adv 6, eaay2611 (2020).

19. Gruber, J. J. et al. Chromatin Remodeling in Response to BRCA2-Crisis. Cell Rep. 28, 2182–2193.e6 (2019).

20. Lim, P. X., Zaman, M., Feng, W. & Jasin, M. BRCA2 promotes genomic integrity and therapy resistance primarily through its role in homology-directed repair. Mol. Cell (2024) doi:10.1016/j.molcel.2023.12.025.

21. Tan, S. L. W. et al. A Class of Environmental and Endogenous Toxins Induces BRCA2 Haploinsufficiency and Genome Instability. Cell 169, 1105–1118.e15 (2017).

22. Arnold, K. et al. Lower level of BRCA2 protein in heterozygous mutation carriers is correlated with an increase in DNA double strand breaks and an impaired DSB repair. Cancer Lett 243, 90–100 (2006).

23. Skoulidis, F. et al. Germline Brca2 heterozygosity promotes Kras(G12D) -driven carcinogenesis in a murine model of familial pancreatic cancer. Cancer Cell 18, 499–509 (2010).

24. Bellacosa, A. et al. Altered gene expression in morphologically normal epithelial cells from heterozygous carriers of BRCA1 or BRCA2 mutations. Cancer Prev Res 3, 48–61 (2010).

25. Tannenbaum, B., Mofunanya, T. & Schoenfeld, A. R. DNA damage repair is unaffected by mimicked heterozygous levels of BRCA2 in HT-29 cells. Int J Biol Sci 3, 402–407 (2007).

26. Warren, M., Lord, C. J., Masabanda, J., Griffin, D. & Ashworth, A. Phenotypic effects of heterozygosity for a BRCA2 mutation. Hum Mol Genet 12, 2645–2656 (2003).

27. Kim, M.-K. et al. Increased rates of spontaneous sister chromatid exchange in lymphocytes of BRCA2+/-carriers of familial breast cancer clusters. Cancer Lett. 210, 85–94 (2004).

28. Kong, L. R. et al. A glycolytic metabolite bypasses “two-hit” tumor suppression by BRCA2. Cell (2024) doi:10.1016/j.cell.2024.03.006.

29. Rebbeck, T. R. et al. Association of type and location of BRCA1 and BRCA2 mutations with risk of breast and ovarian cancer. Jama 313, 1347–1361 (2015).

30. Ware, M. D. et al. Does nonsense-mediated mRNA decay explain the ovarian cancer cluster region of the BRCA2 gene? Oncogene 25, 323–328 (2006).

31. Berman, D. B. et al. A common mutation in BRCA2 that predisposes to a variety of cancers is found in both Jewish Ashkenazi and non-Jewish individuals. Cancer Res. 56, 3409–14 (1996).

32. Neuhausen, S. Recurrent BRCA2 6174delT mutations in Ashkenazi Jewish women affected by breast cancer. Nature Genet. 13, 126–128 (1996).

33. Soule, H. D. et al. Isolation and characterization of a spontaneously immortalized human breast epithelial cell line, MCF-10. Cancer research 50, 6075–6086 (1990).

34. Oddoux, C. et al. The carrier frequency of the BRCA2 6174delT mutation among Ashkenazi Jewish individuals is approximately 1%. Nat. Genet. 14, 188–190 (1996).

35. Spain, B. H., Larson, C. J., Shihabuddin, L. S., Gage, F. H. & Verma, I. M. Truncated BRCA2 is cytoplasmic: implications for cancer-linked mutations. Proc. Natl. Acad. Sci. 96, 13920–13925 (1999).

36. Jensson, B. O. et al. Actionable Genotypes and Their Association with Life Span in Iceland. N. Engl. J. Med. 389, 1741–1752 (2023).

37. Mikaelsdottir, E. K., Valgeirsdottir, S., Eyfjord, J. E. & Rafnar, T. The Icelandic founder mutation BRCA2 999del5: analysis of expression. Breast Cancer Res. 6, R284–R290 (2003).

38. Bryant, H. E. et al. Specific killing of BRCA2-deficient tumours with inhibitors of poly(ADP-ribose) polymerase. Nature 434, 913–917 (2005).

39. Farmer, H. et al. Targeting the DNA repair defect in BRCA mutant cells as a therapeutic strategy. Nature 434, 917–921 (2005).

40. Vugic, D., Ehlen, A. & Carreira, A. Monitoring Homologous Recombination Activity in Human Cells. Methods Mol. Biol. 2153, 115–126 (2021).

41. Cong, K. et al. Replication gaps are a key determinant of PARP inhibitor synthetic lethality with BRCA deficiency. Mol. Cell 81, 3128–3144.e7 (2021).

42. Michalet, X. et al. Dynamic Molecular Combing: Stretching the Whole Human Genome for High-Resolution Studies. Science 277, 1518–1523 (1997).

43. Razavi, P. et al. The Genomic Landscape of Endocrine-Resistant Advanced Breast Cancers. Cancer Cell 34, 427–438.e6 (2018).

44. Davies, H. et al. HRDetect is a predictor of BRCA1 and BRCA2 deficiency based on mutational signatures. Nat. Med. 23, 517–525 (2017).

45. Nik-Zainal, S. et al. Landscape of somatic mutations in 560 breast cancer whole-genome sequences. Nature 534, 47–54 (2016).

46. Mendez, M. G., Kojima, S. & Goldman, R. D. Vimentin induces changes in cell shape, motility, and adhesion during the epithelial to mesenchymal transition. FASEB J. 24, 1838–1851 (2010).

47. Gilles, C. et al. Vimentin contributes to human mammary epithelial cell migration. J. Cell Sci. 112, 4615–4625 (1999).

48. Cifone, M. A. & Fidler, I. J. Correlation of patterns of anchorage-independent growth with in vivo behavior of cells from a murine fibrosarcoma. Proc. Natl. Acad. Sci. 77, 1039–1043 (1980).

49. Sachs, N. et al. A Living Biobank of Breast Cancer Organoids Captures Disease Heterogeneity. Cell 172, 373–386.e10 (2018).

50. Milner, J., Ponder, B., Hughes-Davies, L., Seltmann, M. & Kouzarides, T. Transcriptional activation functions in BRCA2. Nature 386, 772–773 (1997).

51. Edwards, S. L. et al. Resistance to therapy caused by intragenic deletion in BRCA2. Nature 451, 1111–1115 (2008).

52. Sun, L. & Carpenter, G. Epidermal growth factor activation of NF-κB is mediated through IκBα degradation and intracellular free calcium. Oncogene 16, 2095–2102 (1998).

53. Alvaro-Aranda, L. et al. The BRCA2 R2645G variant increases DNA binding and induces hyper-recombination. Nucleic Acids Res. gkad1222 (2023) doi:10.1093/nar/gkad1222.

54. Xia, B. et al. Control of BRCA2 cellular and clinical functions by a nuclear partner, PALB2. Mol. Cell 22, 719–729 (2006).

55. Le, H. P. et al. DSS1 and ssDNA regulate oligomerization of BRCA2. Nucleic Acids Res. 45, 4507–16 (2020).

56. Shahid, T. et al. Structure and mechanism of action of the BRCA2 breast cancer tumor suppressor. Nat. Struct. Mol. Biol. 21, 962–968 (2014).

57. Zhong, H., Voll, R. E. & Ghosh, S. Phosphorylation of NF-κB p65 by PKA Stimulates Transcriptional Activity by Promoting a Novel Bivalent Interaction with the Coactivator CBP/p300. Mol. Cell 1, 661–671 (1998).

58. Kiernan, R. et al. Post-activation Turn-off of NF-κB-dependent Transcription Is Regulated by Acetylation of p65*. J. Biol. Chem. 278, 2758–2766 (2003).

59. Yang, X.-J., Ogryzko, V. V., Nishikawa, J., Howard, B. H. & Nakatani, Y. A p300/CBP-associated factor that competes with the adenoviral oncoprotein E1A. Nature 382, 319–324 (1996).

60. Willis, A., Jung, E. J., Wakefield, T. & Chen, X. Mutant p53 exerts a dominant negative effect by preventing wild-type p53 from binding to the promoter of its target genes. Oncogene 23, 2330–2338 (2004).

61. Kim, J. J. et al. PCAF-Mediated Histone Acetylation Promotes Replication Fork Degradation by MRE11 and EXO1 in BRCA-Deficient Cells. Mol. Cell 80, 327–344.e8 (2020).

62. Shimelis, H. et al. BRCA2 Hypomorphic Missense Variants Confer Moderate Risks of Breast Cancer. Cancer Res. 77, 2789–2799 (2017).

63. Berger, A. H., Knudson, A. G. & Pandolfi, P. P. A continuum model for tumour suppression. Nature 476, 163–169 (2011).

64. Li, C. M.-C. et al. Brca1 haploinsufficiency promotes early tumor onset and epigenetic alterations in a mouse model of hereditary breast cancer. Nat. Genet. 1–13 (2024) doi:10.1038/s41588-024-01958-6.

65. Michelena, J. et al. Analysis of PARP inhibitor toxicity by multidimensional fluorescence microscopy reveals mechanisms of sensitivity and resistance. Nat. Commun. 9, 2678 (2018).

66. Spegg, V. et al. Phase separation properties of RPA combine high-affinity ssDNA binding with dynamic condensate functions at telomeres. Nat. Struct. Mol. Biol. 30, 451–462 (2023).

67. Zerbino, D. R. & Birney, E. Velvet: Algorithms for de novo short read assembly using de Bruijn graphs. Genome Res. 18, 821–829 (2008).

68. Koboldt, D. C. et al. VarScan 2: Somatic mutation and copy number alteration discovery in cancer by exome sequencing. Genome Res. 22, 568–576 (2012).

