## Supplementary material for "A recurrent pathogenic *BRCA2* truncating variant reveals a role for BRCA2-PCAF complex in modulating NF-κB-driven transcription": Table S1

**Table S1**. LOH status and age of diagnosis in a set of 999del5 and c.5946delT bearing breast tumors

999del5 (c.771_775del) breast tumors

| **Tumor ID** | **Cohort** | **Gender** | **Age** | **Subtype** | **LOH**  **status** | **Method of detection** |
| --- | --- | --- | --- | --- | --- | --- |
| 1C3BR15Z39 | PMID: 27499891 | F | 39 | LumB | LOH | Taqman |
| 1C3BV22J99 | PMID: 27499891 | F | 59 | LumA | No | Taqman |
| 1C6DY24V99 | PMID: 27499891 | F | 59 | LumB | No | Taqman |
| 1C6LW39K79 | PMID: 27499891 | F | 48 | LumB | LOH | Taqman |
| 1C6PW09W49 | PMID: 27499891 | F | 44 | LumB | LOH | Taqman |
| 1E6DZ53M99 | PMID: 27499891 | F | 46 | LumB | LOH | Taqman |
| 1G6DY49K99 | PMID: 27499891 | F | 54 | LumA | No | aCGH+Taqman |
| 1G6NY45K59 | PMID: 27499891 | F | 49 | LumA | LOH | Taqman |
| 1G6NY69W59 | PMID: 27499891 | F | 45 | LumA | No | Taqman |
| 1G6TY99J09 | PMID: 27499891 | F | 53 | LumB | LOH | Taqman |
| 1I3BZ82V79 | PMID: 27499891 | F | 40 | LumA | No | aCGH+Taqman |
| 1I6JV54I09 | PMID: 27499891 | F | 54 | NA | LOH | Taqman |
| 1K6FY91X89 | PMID: 27499891 | F | 49 | TNBC | No | aCGH+Taqman |
| 1K6NY52V49 | PMID: 27499891 | F | 61 | LumB | LOH | aCGH+Taqman |
| 1K6TA23K09 | PMID: 27499891 | M | 77 | NA | LOH | Taqman |
| 1O6JR12L59 | PMID: 27499891 | F | 35 | LumA | LOH | aCGH+Taqman |
| 1O6LZ89K99 | PMID: 27499891 | F | 45 | TNBC | No | aCGH+Taqman |
| 1O6RW27Z79 | PMID: 27499891 | F | 46 | LumB | No | Taqman |
| 1Q6HW52K19 | PMID: 27499891 | F | 41 | LumA | No | aCGH+Taqman |
| 1Q6HW79W49 | PMID: 27499891 | F | 42 | LumB | No | aCGH+Taqman |
| 1S3BW99Y89 | PMID: 27499891 | F | 40 | TNBC | LOH | aCGH+Taqman |
| 1S6NY19K09 | PMID: 27499891 | F | 51 | LumA | LOH | Taqman |
| 1U6JY02Z19 | PMID: 27499891 | F | 60 | LumB | LOH | aCGH+Taqman |
| 1U6JZ62V49 | PMID: 27499891 | F | 48 | LumA | LOH | Taqman |
| 5C3BZ02K29 | PMID: 27499891 | F | 41 | LumB | No | aCGH+Taqman |
| 5C3DA42V29 | PMID: 27499891 | F | 77 | TNBC | No | aCGH+Taqman |
| 5C6HA02J39 | PMID: 27499891 | F | 73 | LumA | No | aCGH+Taqman |
| 5G3DA99I99 | PMID: 27499891 | F | 76 | LumB | LOH | Taqman |
| 5G3FY33Z09 | PMID: 27499891 | F | 64 | TNBC | No | aCGH+Taqman |
| 5G6DY82W49 | PMID: 27499891 | F | 57 | TNBC | LOH | aCGH+Taqman |
| 5G6HY53L09 | PMID: 27499891 | F | 56 | TNBC | No | Taqman |
| 5G6LV49L09 | PMID: 27499891 | F | 53 | LumA | No | Taqman |
| 5O3BW44M89 | PMID: 27499891 | F | 44 | LumB | LOH | Taqman |
| 5O3BY94V69 | PMID: 27499891 | F | 57 | LumB | LOH | aCGH+Taqman |
| 5Q3BZ74L39 | PMID: 27499891 | F | 57 | LumB | LOH | aCGH+Taqman |
| 5Q6LY34V99 | PMID: 27499891 | F | 70 | TNBC | No | aCGH+Taqman |
| 5S6LV49K39 | PMID: 27499891 | F | 66 | TNBC | No | Taqman |
| 5S6NY32Y09 | PMID: 27499891 | F | 67 | LumB | LOH | Taqman |
| 7S3BZ84M99 | PMID: 27499891 | F | 40 | LumA | No | aCGH+Taqman |
| 7U3DW54M09 | PMID: 27499891 | F | 51 | LumB | LOH | Taqman |
| 7U6JY57W89 | PMID: 27499891 | F | 50 | TNBC | No | Taqman |
| 7U6LZ49J59 | PMID: 27499891 | F | 43 | LumB | LOH | aCGH+Taqman |
| 8C3BY92V99 | PMID: 27499891 | F | 52 | NA | No | Taqman |
| 8C6DW33J79 | PMID: 27499891 | F | 40 | LumA | No | aCGH+Taqman |
| 8C6LY13K99 | PMID: 27499891 | F | 49 | TNBC | No | aCGH+Taqman |
| 8E6LW24J79 | PMID: 27499891 | F | 41 | LumB | LOH | aCGH+Taqman |
| 8G3DY69J69 | PMID: 27499891 | F | 60 | TNBC | No | aCGH+Taqman |
| 8I6FW99I29 | PMID: 27499891 | F | 49 | LumB | LOH | Taqman |
| 8K3FR12V39 | PMID: 27499891 | F | 30 | LumB | LOH | aCGH+Taqman |
| 8M3BZ29X49 | PMID: 27499891 | F | 46 | LumB | No | aCGH+Taqman |
| 8M3FR72W29 | PMID: 27499891 | F | 32 | LumB | LOH | Taqman |
| 8M6LW47W79 | PMID: 27499891 | F | 37 | LumA | No | Taqman |
| 8Q3FR34M29 | PMID: 27499891 | F | 38 | LumA | LOH | Taqman |
| 8Q6PY44Z69 | PMID: 27499891 | F | 55 | TNBC | No | Taqman |
| 8S3BY77V79 | PMID: 27499891 | F | 50 | LumB | LOH | Taqman |
| 8S6TW84Y89 | PMID: 27499891 | F | 34 | LumB | LOH | aCGH+Taqman |
| 8U3FZ64K89 | PMID: 27499891 | F | 44 | LumB | LOH | aCGH+Taqman |
| 8U6TZ84Z19 | PMID: 27499891 | F | 42 | TNBC | No | aCGH+Taqman |
| 371480H | Curie | F | 42 | NA | No | WGS |

c.5946delT breast tumors

| **Tumor ID** | **Cohort** | **Gender** | **Age** | **Subtype** | **LOH**  **status** | **Method of detection** |
| --- | --- | --- | --- | --- | --- | --- |
| TCGA-AO-A03V | PMID: 28831036 | F | 41 | Lum | No | WGS |
| 4756-Brca2Br1 | PMID: 28831036 | F | 70 | TNBC | No | WGS |
| 6013-Brca2Br49 | PMID: 28831036 | F | 62 | Lum | LOH | WGS |
| 6035-Brca2Br10 | PMID: 28831036 | F | 43 | Lum | LOH | WGS |
| 6537 | Penn Univ. | F | 39 | ER+ | LOH | WGS |
| 7281 | Penn Univ. | F | 39 | TNBC | LOH | WGS |
| 462295H | Curie | F | 47 | Lum | LOH | WGS |
| 549025H | Curie | F | 73 | Lum | No | WGS |
| 619138H | Curie | F | 33 | LumB | LOH | WGS |
| 631699H | Curie | F | 49 | NA | LOH | WGS |
| 638538H | Curie | F | 79 | NA | No | WGS |
| H150105 | Curie | F | 49 | TNBC | LOH | WGS |
| 514207H | Curie | F | 65 | Lum | LOH | WGS |
| 439267H | Curie | F | 46 | Lum | No | pyrosequencing |
| 15267 | N.N. Petrov NMRC of Oncology | F | 50 | Lum | LOH | Allele-specific qPCR |
| 36489 | N.N. Petrov NMRC of Oncology | F | 46 | Lum | No | Allele-specific qPCR |
| 36561 | N.N. Petrov NMRC of Oncology | F | 50 | Lum | LOH | Allele-specific qPCR |
| 37559 | N.N. Petrov NMRC of Oncology | F | 28 | NA | LOH | Allele-specific qPCR |
| 40435 | N.N. Petrov NMRC of Oncology | F | 48 | Lum | No | Allele-specific qPCR |
| 5650 | N.N. Petrov NMRC of Oncology | F | 63 | Lum | LOH | Allele-specific qPCR |
| 6601 | N.N. Petrov NMRC of Oncology | F | 45 | NA | LOH | Allele-specific qPCR |
| 29471 | N.N. Petrov NMRC of Oncology | F | 43 | Lum | No | Allele-specific qPCR |
| 9 | PMID: 27836010 | F | NA | NA | LOH | Micro-satellite analysis |
| 11a | PMID: 27836010 | F | NA | NA | LOH | Micro-satellite analysis |
| 12a | PMID: 27836010 | F | NA | NA | No | Micro-satellite analysis |
| P-0000584 | PMID: 30205045 | F | 29 | NA | LOH | WGS |
| P-0001308 | PMID: 30205045 | F | 44 | NA | No | WGS |

NA= not available; Lum= luminal; TNBC= triple negative breast cancer; ER+= estrogen receptor positive
