## Supplementary material for "A recurrent pathogenic *BRCA2* truncating variant reveals a role for BRCA2-PCAF complex in modulating NF-κB-driven transcription": Table S2

**Table S2**. Tumors characteristics in our study cohort

c.999del5 tumors

| **Sample n°** | **Cohort** | **Gender** | **Age** | **Specimen**  **type** | **Histo-**  **pathological**  **subtype** | **Molecular subtype** | **Tumor**  **grade** | **Mitotic**  **Score**  **(Ki67)** | **ER** | **PR** | **HER2** |
| --- | --- | --- | --- | --- | --- | --- | --- | --- | --- | --- | --- |
| PD4872a | PMID: 27135926 | F | 40 | NA | ductal | LumA | II | high | + | + | - |
| PD4876a | PMID: 27135926 | F | 30 | NA | ductal | LumB | III | high | + | + | - |
| PD4951a | PMID: 27135926 | F | 39 | blood | ductal | LumB | II | high | + | + | - |
| PD4952a | PMID: 27135926 | F | 44 | blood | ductal | LumB | III | high | + | + | - |
| PD4953a | PMID: 27135926 | F | 67 | blood | ductal | LumB | II | low | + | - | - |
| PD4954a | PMID: 27135926 | F | 51 | blood | ductal | LumB | II | high | + | + | - |
| PD4957a | PMID: 27135926 | F | 38 | blood | ductal | LumA | II | low | + | + | - |
| PD4958a | PMID: 27135926 | F | 48 | blood | ductal | LumB | II | high | + | + | - |
| 371480H | I.Curie | F | 42 | tissue | Adeno  carcinoma | NA | III | NA | - | - | NA |
| R01 | S. Sigurdsson | F | 49 | blood | ductal | TNBC | III | high | - | - | - |
| R03 | S. Sigurdsson | F | 32 | blood | ductal | LumB | NA | high | + | + | - |

c.5946delT tumors

| **Sample n°** | **Cohort** | | **Gender** | **Age** | **Specimen**  **type** | **Histo-**  **pathological**  **subtype** | **Molecular subtype** | **Tumor**  **grade** | **Mitotic**  **Score** | **ER** | **PR** | **HER2** |
| --- | --- | --- | --- | --- | --- | --- | --- | --- | --- | --- | --- | --- |
| 514207H | I.Curie | F | | 65 | tissue | Adeno  carcinoma | Lum | III | high | + | - | - |
| 462295H | I.Curie | F | | 47 | tissue | Adeno  carcinoma | NA | III | moderate | + | NA | - |
| 631699H | I.Curie | F | | 49 | tissue | Adeno  carcinoma | NA | II | moderate | NA | + | - |
| 452087H | I.Curie | F | | 65 | tissue | Adeno  carcinoma | NA | NA | NA | NA | NA | NA |
| 619138H | I.Curie | F | | 33 | tissue | Adeno  carcinoma | Lum B | III | high | + | + | - |
| H150105 | I.Curie | F | | 49 | tissue | Adeno  carcinoma | TNBC | III | high | + | - | - |
| 638538H | I.Curie | F | | 79 | tissue | Adeno  carcinoma | NA | NA | NA | NA | NA | NA |
| 549025H | I.Curie | F | | 73 | tissue | Adeno  carcinoma | Lum | NA | NA | + | + | - |
| H091731 | I.Curie | F | | 76 | tissue | Adeno  carcinoma | Lum A | II | NA | + | + | - |
