## Supplementary material for "A recurrent pathogenic *BRCA2* truncating variant reveals a role for BRCA2-PCAF complex in modulating NF-κB-driven transcription": Table S3

| Gene | logFC | adj.P.Val | DEG | Pathway* |
|---|---|---|---|---|
| CADM3 | -10.0769212246182 | 0.00843245301498783 | down | NA |
| CNTN3 | -10.0075320995425 | 0.000651317326288731 | down | NA |
| PTGFR | -9.37069239988164 | 0.000651317326288731 | down | Pathway |
| NLGN4X | -8.75409369509799 | 0.0036004679777663 | down | Pathway |
| ATOH8 | -8.74717426003849 | 0.0116363535575482 | down | Pathway |
| PIK3C2G | -8.6611722206756 | 0.00102396540561436 | down | Pathway |
| TDRD5 | -8.65152316618328 | 0.00258763172862039 | down | NA |
| DARC | -8.47101963256215 | 0.00494483479413562 | down | NA |
| GALNT15 | -8.45825887290362 | 0.00178142130125749 | down | NA |
| ZNF22 | -8.39469796886022 | 0.00178142130125749 | down | NA |
| IL1RL1 | -8.33700758787499 | 0.00113925180685624 | down | NA |
| CYP4X1 | -8.20024741790453 | 0.000768858666668757 | down | NA |
| MAGEE1 | -8.14111396026603 | 0.0000455338772482485 | down | NA |
| ZNF880 | -8.12751741524047 | 0.0000957164820963686 | down | NA |
| GGT5 | -8.07500230431774 | 0.0054148113177371 | down | Pathway |
| ESYT3 | -7.62540478225012 | 0.0000615221659977426 | down | NA |
| APOBEC3F | -7.60488533116241 | 0.0019749530781067 | down | NA |
| ADD2 | -7.39358841085771 | 0.00135340220393029 | down | Pathway |
| SLC6A11 | -7.33558128096658 | 0.00405714344667696 | down | NA |
| KCNQ3 | -7.23798264090939 | 0.00252194956679289 | down | NA |
| MYBPC1 | -7.19909708708476 | 0.0135609845534393 | down | Pathway |
| SRGN | -7.14069728421559 | 0.004556189885561 | down | Pathway |
| FPR1 | -7.12875854689089 | 0.00227280474998497 | down | Pathway |
| DEPDC7 | -7.12874549277296 | 0.005098243527636 | down | NA |
| CDKN1C | -7.11886535171895 | 0.010554643572276 | down | Pathway |
| BST2 | -7.06815725058892 | 0.00150823880964305 | down | Pathway |
| RERG | -7.02274993699795 | 0.00244648580914608 | down | Pathway |
| ADH1B | -7.01971183947436 | 0.00695131164785899 | down | Pathway |
| SFTPB | -7.00458383313603 | 0.00896855015239253 | down | NA |
| CES4A | -6.99162761938316 | 0.010554643572276 | down | NA |
| ZNF883 | -6.97364514431225 | 0.022139039655041 | down | NA |
| SPARC | -6.97125832504797 | 0.013123721662465 | down | Pathway |
| TCN2 | -6.90101534952136 | 0.0071472086286553 | down | NA |
| SLC16A4 | -6.88746115165406 | 0.000298560501862181 | down | NA |
| EXO5 | -6.79574876068121 | 0.0036004679777663 | down | NA |
| CES1 | -6.78053429403356 | 0.015148327326822 | down | Pathway |
| FAM9B | -6.71251398942241 | 0.00178142130125749 | down | NA |
| TCIRG1 | -6.61143563539239 | 0.00118647188968779 | down | Pathway |
| RIBC2 | -6.52194325885767 | 0.00178142130125749 | down | NA |
| ZNF287 | -6.46222752547173 | 0.00521287207365194 | down | NA |
| CYP4B1 | -6.41980171474149 | 0.0154004198295072 | down | Pathway |
| TMC5 | -6.40051142063544 | 0.0211252987656376 | down | NA |
| FAM65B | -6.38391732810947 | 0.0054148113177371 | down | NA |
| PLCB4 | -6.36568574657157 | 0.0086998136871913 | down | Pathway |
| NMNAT3 | -6.29780650179915 | 0.00252194956679289 | down | NA |
| SLCO2A1 | -6.25575770547476 | 0.00265176893220175 | down | NA |
| WIPF1 | -6.21318992002155 | 0.000433169655547905 | down | Pathway |
| NRSN2 | -6.19352479787727 | 0.00517258555667806 | down | NA |
| ABCA6 | -6.13924845981803 | 0.00113925180685624 | down | NA |
| KLRC2 | -6.12684128739877 | 0.00546863024390161 | down | NA |
| ZNF732 | -6.10460071159755 | 0.00252194956679289 | down | NA |
| LAPTM5 | -6.08639737887826 | 0.00308695918387617 | down | NA |

| | | | | |
|---|---|---|---|---|
| C10orf10 | -6.08425994434404 | 0.00227280474998497 | down | NA |
| COL8A1 | -6.01340068147648 | 0.00776131275376969 | down | Pathway |
| ACSL5 | -5.99282027310203 | 0.00240061511093794 | down | Pathway |
| PNLIPRP3 | -5.98011623686289 | 0.00302788431548657 | down | NA |
| BBOX1 | -5.95367842783029 | 0.0201799378678864 | down | NA |
| AKR1C1 | -5.92839201021625 | 0.0294494188378723 | down | Pathway |
| CXCR1 | -5.85138574473949 | 0.0147053724716601 | down | Pathway |
| PSCA | -5.84032605193917 | 0.0122796814813349 | down | Pathway |
| RTN1 | -5.78831199827481 | 0.005098243527636 | down | NA |
| RXFP1 | -5.77805024692885 | 0.0111728595655869 | down | Pathway |
| SULT1E1 | -5.74206001232705 | 0.00426004391502599 | down | Pathway |
| EBF1 | -5.69594750219649 | 0.0267622767466902 | down | NA |
| FMN2 | -5.69196895725043 | 0.0230605480994224 | down | Pathway |
| TMEM100 | -5.68321684367156 | 0.0169765317240416 | down | Pathway |
| AKR1C2 | -5.6606962006623 | 0.0210999421546606 | down | Pathway |
| RNF157 | -5.65093268415879 | 0.00258763172862039 | down | Pathway |
| STAT5A | -5.63360391301583 | 0.0071472086286553 | down | Pathway |
| AKR1B10 | -5.5721820376958 | 0.00925030815276274 | down | Pathway |
| DCN | -5.56340487756776 | 0.0128730119818305 | down | Pathway |
| FAM174B | -5.5136385112208 | 0.00147975674650345 | down | NA |
| ZNF671 | -5.49750636086815 | 0.00144411613966197 | down | NA |
| IL8 | -5.49532829394296 | 0.00182265900362016 | down | NA |
| ABCD2 | -5.469314911834 | 0.000674032279543325 | down | Pathway |
| STAC2 | -5.46486908201147 | 0.015148327326822 | down | Pathway |
| HSD17B2 | -5.45914621479473 | 0.00169541382127618 | down | NA |
| C5orf46 | -5.43914412534879 | 0.0237393482940505 | down | NA |
| VASH1 | -5.42517953943667 | 0.0435818424778398 | down | Pathway |
| ADAM12 | -5.38080234544027 | 0.00142892100222063 | down | Pathway |
| CACNA2D3 | -5.35009626279719 | 0.0086998136871913 | down | NA |
| KCTD8 | -5.2917111365642 | 0.00405714344667696 | down | Pathway |
| IFI44L | -5.23380228513537 | 0.00272789854482884 | down | NA |
| TNFAIP6 | -5.23333534485907 | 0.0245187619921072 | down | Pathway |
| PPAPDC1A | -5.23298446385275 | 0.00896855015239253 | down | NA |
| ASRGL1 | -5.2305275168271 | 0.00106261579796104 | down | NA |
| KLRC3 | -5.18712066174357 | 0.00334081127395787 | down | NA |
| IL22RA2 | -5.16674528640392 | 0.0311750959621759 | down | Pathway |
| TMEM156 | -5.13932091494979 | 0.0138941160958012 | down | NA |
| CARD17 | -5.13474321824394 | 0.000255957655179796 | down | Pathway |
| CACNB2 | -5.10775179990458 | 0.00200626005588499 | down | Pathway |
| ANGPT1 | -5.08882390251926 | 0.00144411613966197 | down | Pathway |
| FAM127C | -5.0820399494582 | 0.0170255390968829 | down | NA |
| TRPV4 | -5.05773722929208 | 0.016330308338876 | down | Pathway |
| IL1R2 | -5.03842039174573 | 0.0400066123929638 | down | Pathway |
| SLC22A18AS | -5.01763793908743 | 0.00337324148306265 | down | NA |
| ARL9 | -5.00105625063694 | 0.0053996054992121 | down | NA |
| RORC | -4.99078642166687 | 0.00911536903416098 | down | Pathway |
| ARMC4 | -4.97462225715944 | 0.00386066187299184 | down | NA |
| S1PR3 | -4.95389352577834 | 0.00593830100427534 | down | Pathway |
| S100A8 | -4.91935364857387 | 0.0267622767466902 | down | Pathway |
| SMOC1 | -4.91000872172288 | 0.0282135400546423 | down | Pathway |
| KCNJ2 | -4.90700934847716 | 0.0138967236371032 | down | Pathway |
| CYP4Z1 | -4.89175792801591 | 0.000433169655547905 | down | Pathway |
| SCNN1B | -4.88574730909804 | 0.0245483411558718 | down | Pathway |

| | | | | |
|---|---|---|---|---|
| SYN3 | -4.87549528860559 | 0.0238682080454715 | down | NA |
| CDIP1 | -4.87272128688214 | 0.00643719647476864 | down | NA |
| AKR1B15 | -4.85915013092891 | 0.0439328688173528 | down | Pathway |
| IL6 | -4.85660764527301 | 0.0179687117421246 | down | Pathway |
| LAMA4 | -4.82939780051801 | 0.0152705952579538 | down | NA |
| TBL1X | -4.77266222817638 | 0.0259503018863846 | down | NA |
| ATP8B4 | -4.72383950697525 | 0.0269963322570122 | down | NA |
| AOX1 | -4.69980974367011 | 0.0174970232259838 | down | NA |
| ST18 | -4.61729274151051 | 0.0062306377236223 | down | Pathway |
| WNT3A | -4.60811036597581 | 0.0124796565780751 | down | Pathway |
| CHI3L2 | -4.58456521778898 | 0.0054148113177371 | down | Pathway |
| LRRN1 | -4.56279161946741 | 0.0193505295868115 | down | Pathway |
| AKR1C3 | -4.55231597159733 | 0.0128702987042865 | down | Pathway |
| CXCL1 | -4.52557001629118 | 0.00144411613966197 | down | Pathway |
| SAMSN1 | -4.5016777573781 | 0.0321381722961886 | down | Pathway |
| METTL7B | -4.49672446090401 | 0.0380576218383549 | down | NA |
| SLC9A2 | -4.47575338172601 | 0.0194925393212716 | down | Pathway |
| SLC25A18 | -4.46569017651924 | 0.00515269958448667 | down | NA |
| SERPINA5 | -4.45591533301858 | 0.00381572485124622 | down | NA |
| ABCA9 | -4.44473460555452 | 0.015148327326822 | down | NA |
| KLHL4 | -4.43873330584292 | 0.00298509759394963 | down | NA |
| GSDMC | -4.42953756157726 | 0.018188252251489 | down | NA |
| ABCC2 | -4.42285227719579 | 0.00147975674650345 | down | Pathway |
| CACNA1D | -4.41060631145483 | 0.0420973154335111 | down | Pathway |
| CXCR4 | -4.41049048048176 | 0.00381572485124622 | down | Pathway |
| PNPLA3 | -4.39286367056188 | 0.0054148113177371 | down | Pathway |
| C12orf39 | -4.38280863434205 | 0.0304605322782198 | down | NA |
| CFH | -4.37913480515151 | 0.00728949742823313 | down | NA |
| PAOX | -4.3774088485098 | 0.0352596894579683 | down | NA |
| PLAC8 | -4.35170853777466 | 0.0176519651049434 | down | Pathway |
| ASAP3 | -4.33843863024477 | 0.000651317326288731 | down | Pathway |
| TEK | -4.33680599990967 | 0.0054148113177371 | down | Pathway |
| ANKRD35 | -4.3293994605873 | 0.00824205122503509 | down | NA |
| VNN3 | -4.2917540483228 | 0.016330308338876 | down | NA |
| NEXN | -4.28724795190354 | 0.00178142130125749 | down | Pathway |
| HSD11B1 | -4.27052345424236 | 0.0148641268014274 | down | NA |
| SOWAHB | -4.26605770745868 | 0.0111219906172995 | down | NA |
| PDE7B | -4.25068089415059 | 0.0035229097927994 | down | Pathway |
| GPR141 | -4.22579111216431 | 0.00182265900362016 | down | NA |
| ABLIM3 | -4.1802031782391 | 0.00102396540561436 | down | NA |
| C2CD4A | -4.15855680294418 | 0.0431558003872985 | down | Pathway |
| OLAH | -4.14403652760964 | 0.0287819607414773 | down | Pathway |
| CTSE | -4.12178233130885 | 0.0196813478636481 | down | NA |
| SULT1B1 | -4.11889955014662 | 0.00728651967850821 | down | Pathway |
| GYS2 | -4.10080660386731 | 0.00178142130125749 | down | NA |
| RSPO2 | -4.09525565611273 | 0.0400720386567861 | down | Pathway |
| LAT2 | -4.08765455954097 | 0.0058845229487207 | down | Pathway |
| AHRR | -4.07036767266042 | 0.0374113323533681 | down | NA |
| ZG16B | -4.06162036860832 | 0.00178142130125749 | down | Pathway |
| FIBIN | -4.05117585896073 | 0.00591289489354877 | down | NA |
| STARD8 | -4.04709467016028 | 0.00357363537533407 | down | NA |
| CCDC28B | -4.02446033080069 | 0.0251424344181079 | down | NA |
| TENC1 | -4.01354926454108 | 0.00770585223804868 | down | NA |

| | | | | |
|---|---|---|---|---|
| VNN2 | -4.00455165609079 | 0.0130391966139351 | down | NA |
| PTPN22 | -4.00320638365487 | 0.0111824216484844 | down | Pathway |
| ZNF32 | -3.96931574158628 | 0.00496606953909166 | down | NA |
| COL24A1 | -3.96738947464767 | 0.0180795351180147 | down | Pathway |
| MCTP1 | -3.96122805867111 | 0.00721317781275581 | down | Pathway |
| NPR3 | -3.9414128998595 | 0.00147975674650345 | down | Pathway |
| DHRS9 | -3.91304489361247 | 0.00517258555667806 | down | Pathway |
| TSPAN1 | -3.89622576341669 | 0.00246457991558625 | down | NA |
| KLRC1 | -3.86321633509054 | 0.0111728595655869 | down | NA |
| ETNK2 | -3.8580811396102 | 0.0054148113177371 | down | NA |
| SRSF12 | -3.83358170406834 | 0.00749532301861601 | down | NA |
| SERPINA1 | -3.83118685625967 | 0.0147308841871448 | down | Pathway |
| LMCD1 | -3.82222410425676 | 0.00178142130125749 | down | Pathway |
| ITGB2 | -3.77348855480112 | 0.00591289489354877 | down | Pathway |
| EVI2A | -3.72071964303897 | 0.0149974875776845 | down | NA |
| SNED1 | -3.71141920170722 | 0.00777074350823438 | down | Pathway |
| ITGA10 | -3.70494853600732 | 0.0400066123929638 | down | Pathway |
| GLDC | -3.70404763540145 | 0.0428564414308668 | down | Pathway |
| XKRX | -3.67942065580646 | 0.00178142130125749 | down | NA |
| ST8SIA1 | -3.64689346590561 | 0.0134270017038353 | down | NA |
| GAS1 | -3.64120282641341 | 0.0104492197841631 | down | Pathway |
| C1S | -3.61344619179263 | 0.0106259042981105 | down | NA |
| S100A9 | -3.60618805887194 | 0.0483357666963776 | down | Pathway |
| FETUB | -3.60159737359391 | 0.0132160745645083 | down | NA |
| LYPD3 | -3.59161197929005 | 0.00404689710333156 | down | Pathway |
| SELENBP1 | -3.58227099715642 | 0.0106259042981105 | down | NA |
| AGR2 | -3.57597744931816 | 0.0144140090034824 | down | Pathway |
| SUSD1 | -3.53618294678528 | 0.00178142130125749 | down | NA |
| SEPP1 | -3.53315372279056 | 0.0031672425850628 | down | NA |
| ARID5B | -3.52212691600468 | 0.00189349418112886 | down | Pathway |
| MILR1 | -3.49537296079981 | 0.0121433833434938 | down | NA |
| ACPP | -3.49371757488574 | 0.00977426819056698 | down | NA |
| CXCL3 | -3.477904071283 | 0.00507823891194564 | down | Pathway |
| KCNJ15 | -3.47709969568016 | 0.00676515248648476 | down | NA |
| SEMA5A | -3.46374666549465 | 0.00250674240741419 | down | Pathway |
| KCNK2 | -3.458090381502 | 0.00923052977989724 | down | Pathway |
| PLEKHA6 | -3.45530884537139 | 0.00777074350823438 | down | NA |
| POU5F1 | -3.44169431706877 | 0.0356786735999177 | down | Pathway |
| SIPA1L2 | -3.42961493139144 | 0.0262422747603064 | down | NA |
| IL1R1 | -3.42844055020407 | 0.00531139971143219 | down | Pathway |
| TACC2 | -3.4257443499772 | 0.00227280474998497 | down | NA |
| CXCL2 | -3.41735756642211 | 0.0130498140241582 | down | Pathway |
| CLEC2B | -3.41725323566111 | 0.0107090702638709 | down | NA |
| PANX2 | -3.41274995769234 | 0.0186597863997083 | down | NA |
| HAPLN3 | -3.40339858601798 | 0.0065222493677266 | down | NA |
| ICAM5 | -3.40151325634085 | 0.0229056489929945 | down | NA |
| C10orf90 | -3.39877917871842 | 0.00113925180685624 | down | NA |
| PRR16 | -3.38558512157684 | 0.0176519651049434 | down | Pathway |
| EYA1 | -3.35574755703977 | 0.00405714344667696 | down | Pathway |
| PAK3 | -3.35339773410099 | 0.0143949538686486 | down | Pathway |
| BCL6 | -3.33887601028235 | 0.00178142130125749 | down | Pathway |
| FADS2 | -3.33542002406174 | 0.00828039871964914 | down | Pathway |
| PAPPA | -3.33125072936095 | 0.0201810661313687 | down | NA |

| | | | | |
|---|---|---|---|---|
| TCEA3 | -3.31214157734595 | 0.00478424392890516 | down | NA |
| P2RX6 | -3.29683611565857 | 0.0138941160958012 | down | Pathway |
| ANKS1B | -3.29225669445252 | 0.0213570845460382 | down | NA |
| PADI4 | -3.28608328534351 | 0.0268024230152425 | down | NA |
| ALDH3A1 | -3.28105197025773 | 0.00144411613966197 | down | NA |
| PLCB1 | -3.27964223854134 | 0.0267325809773597 | down | Pathway |
| IFI44 | -3.23694062436466 | 0.007147208628655З | down | NA |
| SCNN1G | -3.23681564030817 | 0.00702066841229523 | down | Pathway |
| CYBA | -3.2280001961561 | 0.0204810479500326 | down | Pathway |
| PCDH18 | -3.22736828121204 | 0.0102106481651848 | down | NA |
| POLN | -3.2009408126027 | 0.0102295968756798 | down | NA |
| C4orf19 | -3.1963354426263 | 0.0222504819698704 | down | NA |
| RBPMS2 | -3.18041356919321 | 0.0133137063525308 | down | Pathway |
| ACOT2 | -3.17010742969204 | 0.0054706052525586 | down | Pathway |
| PCDH7 | -3.15763460986353 | 0.0247862496022762 | down | NA |
| PRR15L | -3.14773735839437 | 0.0262422747603064 | down | NA |
| MPZ | -3.14453742793331 | 0.0463496604912139 | down | NA |
| IKZF2 | -3.1429011432161 | 0.0053996054992121 | down | NA |
| TM4SF1 | -3.14276826026722 | 0.0124408531882066 | down | NA |
| HIST1H2BN | -3.14223532653703 | 0.00824205122503509 | down | NA |
| DDC | -3.13874492736003 | 0.00176517598072332 | down | NA |
| ANXA6 | -3.13825860488312 | 0.00606913744496404 | down | Pathway |
| SP140 | -3.12819539372151 | 0.00979169933051757 | down | NA |
| NFATC4 | -3.11746326941695 | 0.0374596033802236 | down | Pathway |
| HPGD | -3.11117851327996 | 0.00484852546514122 | down | Pathway |
| KLF15 | -3.10927760204747 | 0.0418967487771853 | down | Pathway |
| ADRA1B | -3.07363922890274 | 0.0105896693547357 | down | Pathway |
| NBEA | -3.068376909426 | 0.00184215937779317 | down | NA |
| DYSF | -3.04968111803876 | 0.0473275296212984 | down | NA |
| WNT5A | -3.04129265651873 | 0.0226494393819051 | down | Pathway |
| MAP7D2 | -3.02560000712188 | 0.0315493990815156 | down | NA |
| KLHL38 | -3.01961509639703 | 0.033088935669882 | down | NA |
| GLRX | -3.01955642801035 | 0.005098243527636 | down | NA |
| LYPD5 | -3.0113886338427 | 0.016275865818765 | down | NA |
| GAB3 | -3.00917746058268 | 0.00235306830255325 | down | NA |
| EPHA10 | -2.99152801500484 | 0.0392467988097998 | down | Pathway |
| DSEL | -2.95760713521936 | 0.0400066123929638 | down | Pathway |
| TNS1 | -2.95015552985914 | 0.00484852546514122 | down | NA |
| KRT80 | -2.94122286676619 | 0.0124408531882066 | down | NA |
| CPM | -2.93261495524312 | 0.00426004391502599 | down | NA |
| PHACTR3 | -2.9292024965973 | 0.0352596894579683 | down | NA |
| RARRES3 | -2.92002351306651 | 0.0191725614207769 | down | NA |
| EYS | -2.91717949335216 | 0.00956392481487896 | down | NA |
| GSDMA | -2.90808185699999 | 0.0175135800028153 | down | NA |
| FIBCD1 | -2.88144416277495 | 0.00782765597435922 | down | NA |
| TAS1R3 | -2.87800749361472 | 0.0321456879645613 | down | NA |
| XKR3 | -2.87486904945327 | 0.0222504819698704 | down | NA |
| TNFSF14 | -2.87282750904331 | 0.0201799378678864 | down | Pathway |
| FGFBP1 | -2.86171931511438 | 0.010554643572276 | down | Pathway |
| HERC5 | -2.85920851864778 | 0.0107090702638709 | down | NA |
| PDE6A | -2.83970224239313 | 0.00189349418112886 | down | Pathway |
| PADI1 | -2.83693395367278 | 0.010846865278975 | down | NA |
| GPR64 | -2.83478895965305 | 0.0106235251353713 | down | NA |

| | | | | |
|---|---|---|---|---|
| SERPINE2 | -2.81670895300662 | 0.0182216278626592 | down | Pathway |
| CHST6 | -2.81111803878256 | 0.00591289489354877 | down | Pathway |
| MAB21L3 | -2.7895545327976 | 0.010554643572276 | down | NA |
| TMEM92 | -2.78898605783004 | 0.0132160745645083 | down | NA |
| GBP2 | -2.77857864349472 | 0.00113925180685624 | down | NA |
| UBE2L6 | -2.77296064831407 | 0.00500099367069865 | down | NA |
| ACSF2 | -2.77108978146178 | 0.00178142130125749 | down | Pathway |
| FAT4 | -2.76428089853214 | 0.0216151761529874 | down | NA |
| LINC00087 | -2.76176894943265 | 0.00113925180685624 | down | NA |
| ADRB2 | -2.7481891198717 | 0.0053996054992121 | down | Pathway |
| AK7 | -2.74818801173301 | 0.00807750674119537 | down | NA |
| OVCH1 | -2.74361295752132 | 0.0122640526852166 | down | NA |
| THBS1 | -2.71183420247972 | 0.00165808241694114 | down | Pathway |
| CAPN8 | -2.69455791161197 | 0.0216603737210304 | down | NA |
| BTBD16 | -2.69420775788434 | 0.0483525873183295 | down | NA |
| ZEB2 | -2.68707162136257 | 0.00654637280781179 | down | Pathway |
| RNF165 | -2.68140340372184 | 0.0214058597012822 | down | Pathway |
| PDK4 | -2.67199964991104 | 0.0109209797785481 | down | Pathway |
| ATP6V0A4 | -2.67147179185494 | 0.045389458846911 | down | Pathway |
| NAV3 | -2.66341581750932 | 0.0156376081715616 | down | NA |
| GPM6B | -2.66215733948199 | 0.00200253059330313 | down | Pathway |
| ATP1B1 | -2.65730562643278 | 0.00196838376763342 | down | Pathway |
| MGAT5B | -2.64985907541065 | 0.00751072730065212 | down | NA |
| TNIP3 | -2.64825017088986 | 0.0253573488752213 | down | Pathway |
| EVA1A | -2.64551013174725 | 0.00843499908287437 | down | NA |
| SHROOM2 | -2.64487391068103 | 0.0148641268014274 | down | Pathway |
| APOBR | -2.6411456197207 | 0.0288722112538373 | down | Pathway |
| EHF | -2.6211363590428 | 0.044450112293341 | down | Pathway |
| ABLIM2 | -2.6150940154245 | 0.0362774417492569 | down | NA |
| IRAK2 | -2.60613429158804 | 0.0130391966139351 | down | Pathway |
| CHADL | -2.606053522805 | 0.0499850218635955 | down | Pathway |
| COL16A1 | -2.59784882678571 | 0.0258824603631041 | down | Pathway |
| PHEX | -2.57290465264704 | 0.0258682839732607 | down | NA |
| ARHGAP29 | -2.56728052500938 | 0.0120131837342731 | down | NA |
| SULT1A2 | -2.56648950684192 | 0.00381572485124622 | down | Pathway |
| C3orf18 | -2.5607752535623 | 0.0211252987656376 | down | NA |
| SLC45A1 | -2.55935385010352 | 0.0242532429594263 | down | NA |
| CYP39A1 | -2.54898740303625 | 0.0130498140241582 | down | Pathway |
| C5AR1 | -2.54261712586456 | 0.0253740510953186 | down | Pathway |
| GOLT1A | -2.54213754959566 | 0.0189719132259805 | down | NA |
| SLC7A2 | -2.5350093684124 | 0.00843245301498783 | down | Pathway |
| NFKBIZ | -2.52754149739335 | 0.0467726086929653 | down | NA |
| MAP3K8 | -2.52646092424234 | 0.0333731272492139 | down | NA |
| MLPH | -2.5242682472131 | 0.0144526470266525 | down | NA |
| TNFAIP2 | -2.50939984180351 | 0.0239516473700319 | down | NA |
| HPD | -2.50850949368972 | 0.0148205349578936 | down | NA |
| PROM2 | -2.50210262071385 | 0.0199465653162187 | down | NA |
| LPIN1 | -2.49402693791024 | 0.00102396540561436 | down | Pathway |
| PPAP2B | -2.4893377761031 | 0.0331790410698035 | down | NA |
| ST3GAL5 | -2.48848758249687 | 0.0040765885132371 | down | NA |
| OSGIN1 | -2.48829476829703 | 0.0104492197841631 | down | Pathway |
| COL13A1 | -2.48803665841247 | 0.0365507866500216 | down | Pathway |
| CDH2 | -2.47391088430983 | 0.0141322061675459 | down | Pathway |

| | | | | |
|---|---|---|---|---|
| SOD2 | -2.4733853599522 | 0.0277647033898298 | down | Pathway |
| VWA5A | -2.46856796756716 | 0.00934617699878462 | down | NA |
| CPPED1 | -2.46423931302883 | 0.00910729534474426 | down | NA |
| FGF5 | -2.43601263036684 | 0.0387244156283981 | down | Pathway |
| C3 | -2.42718690387025 | 0.0213570845460382 | down | Pathway |
| FCHO1 | -2.42311241910122 | 0.0112334062705262 | down | NA |
| GCNT2 | -2.40928635784101 | 0.0488543966375156 | down | Pathway |
| MMP19 | -2.40613997917646 | 0.025389981851985 | down | Pathway |
| BEAN1 | -2.39751029212169 | 0.0420973154335111 | down | NA |
| SNCG | -2.39595627074911 | 0.0078380861499063 | down | Pathway |
| PTX3 | -2.3922046438315 | 0.00500099367069865 | down | Pathway |
| ALDH1L1 | -2.38966712894157 | 0.0430014060922334 | down | NA |
| HYAL1 | -2.38586072616896 | 0.019520871963596 | down | Pathway |
| ERRFI1 | -2.38442455606618 | 0.0096274444680014 | down | Pathway |
| PLSCR4 | -2.38354902713834 | 0.00702066841229523 | down | Pathway |
| C15orf62 | -2.38354052986661 | 0.00546863024390161 | down | Pathway |
| PIK3R3 | -2.37493518614131 | 0.0102106481651848 | down | Pathway |
| SPSB1 | -2.37402510698613 | 0.00778937934344466 | down | NA |
| IQCD | -2.37174978217299 | 0.0333455930308587 | down | NA |
| CEACAM19 | -2.36764110252446 | 0.0392467988097998 | down | NA |
| PTGER4 | -2.36405099246591 | 0.00749532301861601 | down | Pathway |
| GLIS2 | -2.36319138587381 | 0.0424202879380085 | down | NA |
| SLC26A2 | -2.35006977463642 | 0.00308695918387617 | down | Pathway |
| TGFBR3 | -2.34737523821035 | 0.00606913744496404 | down | Pathway |
| SYT12 | -2.34708137067009 | 0.0445792339514482 | down | NA |
| DPYD | -2.32644981909286 | 0.00598193964559136 | down | NA |
| PDGFRL | -2.32132403993271 | 0.00178142130125749 | down | NA |
| GMPR | -2.31638262406424 | 0.0230948034985204 | down | NA |
| ABCC3 | -2.31583471089037 | 0.004556189885561 | down | Pathway |
| ACSL1 | -2.29929139793535 | 0.00843499908287437 | down | Pathway |
| SPOCK1 | -2.29066040729576 | 0.0234847752774467 | down | Pathway |
| TNFSF13 | -2.2888715649924 | 0.0272971111673572 | down | NA |
| ZNF285 | -2.26574021209546 | 0.0122716776209058 | down | NA |
| DNAJB4 | -2.26559925461207 | 0.00484852546514122 | down | NA |
| PLLP | -2.26407778173342 | 0.033927613893117 | down | NA |
| LURAP1L | -2.24428654011536 | 0.0275994272677768 | down | NA |
| GPRC5C | -2.23759785560294 | 0.0435818424778398 | down | NA |
| ENTPD2 | -2.23731426785559 | 0.0367990281061773 | down | Pathway |
| GBP1 | -2.2329945049602 | 0.0106546715302698 | down | Pathway |
| SLC26A7 | -2.23225562948774 | 0.0459841737280008 | down | NA |
| GCLC | -2.22841640094057 | 0.0482482911547309 | down | Pathway |
| NCR3LG1 | -2.22604055632574 | 0.0138967236371032 | down | NA |
| PIK3IP1 | -2.21829714383733 | 0.0315175207440936 | down | Pathway |
| RAB30 | -2.21444226283941 | 0.0054706052525586 | down | NA |
| CTF1 | -2.21085904558607 | 0.0367187431476375 | down | Pathway |
| ARRB1 | -2.20291384758641 | 0.0111728595655869 | down | Pathway |
| NABP1 | -2.20280448999705 | 0.00178142130125749 | down | NA |
| SLC35E4 | -2.20073990483268 | 0.00734849165405664 | down | NA |
| C9orf84 | -2.19911039634548 | 0.0147308841871448 | down | NA |
| MYEOV | -2.18755542115678 | 0.0323190626882029 | down | NA |
| SFMBT2 | -2.18597837930002 | 0.0212024004701557 | down | NA |
| COL18A1 | -2.18513762160105 | 0.0344113680692851 | down | Pathway |
| GHR | -2.15944166731983 | 0.0472721070917943 | down | Pathway |

| | | | | |
|---|---|---|---|---|
| ACOX2 | -2.15875629122349 | 0.0113258262475813 | down | Pathway |
| GPLD1 | -2.15195571592288 | 0.00189349418112886 | down | Pathway |
| TNFAIP8L3 | -2.12980703568241 | 0.0111646381833105 | down | Pathway |
| PDCD1LG2 | -2.1287470756759 | 0.0345026198173697 | down | Pathway |
| ZNF608 | -2.12499544090566 | 0.00734849165405664 | down | NA |
| KIAA1683 | -2.10434937602039 | 0.045389458846911 | down | NA |
| PDGFB | -2.09445760163709 | 0.00910729534474426 | down | Pathway |
| FLVCR2 | -2.09367424557246 | 0.0347137603169297 | down | NA |
| LINC00961 | -2.08468812442111 | 0.0277523049848282 | down | NA |
| ADAMTS1 | -2.07457220692292 | 0.0201750048585462 | down | Pathway |
| GLUL | -2.06910885707242 | 0.0053996054992121 | down | Pathway |
| TRPV3 | -2.06833370509147 | 0.033088935669882 | down | NA |
| PPARGC1A | -2.06324208870936 | 0.0104492197841631 | down | Pathway |
| KYNU | -2.0574625941896 | 0.00200626005588499 | down | Pathway |
| GVQW1 | -2.04964262900557 | 0.00645724017769797 | down | NA |
| S100A4 | -2.04627088234322 | 0.00447116000598364 | down | Pathway |
| NWD1 | -2.04143074130351 | 0.0143949538686486 | down | NA |
| LRIG1 | -2.04113194559935 | 0.0054148113177371 | down | Pathway |
| ZNF140 | -2.04001825107405 | 0.00273170545121021 | down | NA |
| CFI | -2.03664748142653 | 0.016130762944312 | down | NA |
| BEST2 | -2.03430776172745 | 0.0320257136017401 | down | NA |
| METTL7A | -2.03337034616659 | 0.00252194956679289 | down | NA |
| RGCC | -2.03172762333376 | 0.0146228523478505 | down | Pathway |
| G6PD | -2.02740755834384 | 0.0130498140241582 | down | Pathway |
| LEPR | -2.01490189016982 | 0.0307656843123698 | down | Pathway |
| TDRD6 | -2.00614363291672 | 0.033927613893117 | down | NA |
| TOX2 | -1.99866810784007 | 0.0238682080454715 | down | NA |
| TMEM2 | -1.99689162081945 | 0.00144411613966197 | down | NA |
| CHN2 | -1.98347096563066 | 0.00702066841229523 | down | NA |
| EGF | -1.98173969182649 | 0.0130498140241582 | down | Pathway |
| CD99L2 | -1.98064256360459 | 0.0137416567035548 | down | Pathway |
| TSC22D3 | -1.97614888249614 | 0.0115451559763567 | down | NA |
| FAXDC2 | -1.97494486978331 | 0.0138941160958012 | down | NA |
| DUSP5 | -1.97228054152274 | 0.00154021253051307 | down | Pathway |
| CTGF | -1.96922560176924 | 0.0307086016026121 | down | NA |
| EVI2B | -1.9686410201414 | 0.00687763451878185 | down | NA |
| KDR | -1.96566640084641 | 0.0222504819698704 | down | Pathway |
| S1PR1 | -1.95941053373089 | 0.0403142608678876 | down | Pathway |
| TBXAS1 | -1.95053408622542 | 0.00246457991558625 | down | Pathway |
| PSD4 | -1.94306732356667 | 0.0124408531882066 | down | NA |
| IL6R | -1.94086633772377 | 0.00843245301498783 | down | Pathway |
| NUP210 | -1.93060179732407 | 0.00624317365825117 | down | NA |
| SQRDL | -1.92403471513223 | 0.00178142130125749 | down | NA |
| FBLN1 | -1.91713140121349 | 0.00778937934344466 | down | Pathway |
| NRCAM | -1.91480635132426 | 0.0311651363263454 | down | Pathway |
| ELFN2 | -1.91062112173957 | 0.00484852546514122 | down | NA |
| ZEB1 | -1.90694388982459 | 0.0253740510953186 | down | Pathway |
| CBLN3 | -1.90258178087191 | 0.0108434289526 | down | Pathway |
| TMTC1 | -1.89998430136756 | 0.0157457400676371 | down | NA |
| CARD16 | -1.89633909098668 | 0.0268805994466039 | down | Pathway |
| NALCN | -1.89052373282431 | 0.0355292297896454 | down | NA |
| DUSP10 | -1.89002682727801 | 0.0352596894579683 | down | Pathway |
| CORIN | -1.88958792347061 | 0.0355292297896454 | down | Pathway |

| | | | | |
|---|---|---|---|---|
| NCOA7 | -1.88197039269448 | 0.0333455930308587 | down | NA |
| ARHGAP24 | -1.88120340272534 | 0.005098243527636 | down | Pathway |
| ANXA9 | -1.87970579353244 | 0.00950911307403478 | down | NA |
| BTG1 | -1.8790745330628 | 0.00337324148306265 | down | Pathway |
| IFNGR1 | -1.87782077244719 | 0.005098243527636 | down | Pathway |
| PLXNC1 | -1.87714370490105 | 0.0362981021104693 | down | Pathway |
| NRP1 | -1.87463161096705 | 0.0405759606948465 | down | Pathway |
| ARHGDIB | -1.86518167109416 | 0.0231195994191715 | down | Pathway |
| KLF9 | -1.86426441234149 | 0.0246029301731497 | down | Pathway |
| NAMPT | -1.86351051487368 | 0.0282646904660126 | down | NA |
| MFSD2A | -1.84577291287313 | 0.0118528227912586 | down | Pathway |
| FGFR1 | -1.83725465742417 | 0.020321185017871 | down | Pathway |
| ACPL2 | -1.83583931448804 | 0.0463496604912139 | down | NA |
| CLDN1 | -1.82025055318413 | 0.00252194956679289 | down | Pathway |
| SDPR | -1.81785102289455 | 0.00934617699878462 | down | NA |
| IL18R1 | -1.81569465237602 | 0.0122071261074245 | down | NA |
| PBLD | -1.80647291224383 | 0.00734849165405664 | down | Pathway |
| KIAA0319 | -1.80367052416425 | 0.0230934876660662 | down | Pathway |
| LPAR1 | -1.80094481498405 | 0.0428564414308668 | down | Pathway |
| DENND3 | -1.78969754303401 | 0.0173389359340561 | down | NA |
| PPARG | -1.77290161223229 | 0.0111555943002799 | down | Pathway |
| LPXN | -1.77002784982689 | 0.035902780740593 | down | Pathway |
| CCDC149 | -1.76922113088419 | 0.00178142130125749 | down | NA |
| CSF1R | -1.76056128757959 | 0.0291693300805134 | down | Pathway |
| MYOM1 | -1.7595275225616 | 0.0367187431476375 | down | Pathway |
| SGK1 | -1.75937806816457 | 0.00824205122503509 | down | Pathway |
| GABRA2 | -1.75274457527744 | 0.0416514378514478 | down | Pathway |
| ERC2 | -1.74928545450506 | 0.0152636611117527 | down | Pathway |
| EPS8 | -1.74839546733414 | 0.0096274444680014 | down | Pathway |
| KRT8 | -1.74602921232848 | 0.0147053724716601 | down | NA |
| ACOT1 | -1.7425334403336 | 0.0294613574555184 | down | Pathway |
| C1RL | -1.74146046111781 | 0.0116655741495269 | down | NA |
| PNRC1 | -1.73818321052469 | 0.0121135876833331 | down | NA |
| PRKD1 | -1.72373086335577 | 0.0108082132007718 | down | Pathway |
| KIAA0513 | -1.72154903675032 | 0.0296230124526444 | down | NA |
| IFIT3 | -1.71945698834821 | 0.0176519651049434 | down | NA |
| TFPI | -1.71413057353098 | 0.0147657330463027 | down | Pathway |
| GRAMD2 | -1.71039441980636 | 0.0174970232259838 | down | NA |
| BTN3A3 | -1.70156039141806 | 0.00933842120625558 | down | NA |
| PIGZ | -1.69964199387693 | 0.0203221355260243 | down | NA |
| SLC16A5 | -1.69612152590791 | 0.0262422747603064 | down | NA |
| BASP1 | -1.69544756044847 | 0.0201810661313687 | down | Pathway |
| ERICH2 | -1.6879993991793 | 0.0479428131353168 | down | NA |
| CCBE1 | -1.6811610035329 | 0.0217206551740143 | down | Pathway |
| CUBN | -1.6647516507586 | 0.020321185017871 | down | Pathway |
| OPN3 | -1.66219857599509 | 0.0122640526852166 | down | NA |
| BLNK | -1.66205992990849 | 0.0086998136871913 | down | NA |
| PRDM8 | -1.65942898305794 | 0.0223802245837849 | down | Pathway |
| NFIL3 | -1.65528478017036 | 0.0411587988695176 | down | NA |
| SPTSSA | -1.64927885555657 | 0.0133552152128459 | down | Pathway |
| MEGF9 | -1.64520163860567 | 0.00531139971143219 | down | Pathway |
| F2R | -1.64439211816467 | 0.0147053724716601 | down | Pathway |
| AIF1L | -1.63977065741369 | 0.0081017485417304 | down | Pathway |

| | | | | |
|---|---|---|---|---|
| GDPD3 | -1.63847089791954 | 0.0395186687480931 | down | Pathway |
| EPGN | -1.63616812900671 | 0.0170233078156277 | down | Pathway |
| C1orf21 | -1.6179607158412 | 0.00749532301861601 | down | NA |
| TMEM139 | -1.61406899088515 | 0.0400066123929638 | down | NA |
| MFGE8 | -1.6097888632034 | 0.0458038100332125 | down | NA |
| C3orf55 | -1.60807417984038 | 0.0413751768823673 | down | NA |
| EMP3 | -1.60503722255139 | 0.0495460136912076 | down | NA |
| ARMC12 | -1.60126894504496 | 0.0304059231674357 | down | Pathway |
| ZBED2 | -1.60007467935654 | 0.0213557792211214 | down | Pathway |
| MMD | -1.59815042401767 | 0.00343450619497383 | down | Pathway |
| ELAVL2 | -1.59646044399569 | 0.0428564414308668 | down | NA |
| VNN1 | -1.58910602163286 | 0.0296404297305883 | down | NA |
| ABHD5 | -1.5878293757438 | 0.0106546715302698 | down | Pathway |
| GPR37 | -1.58104579462665 | 0.0418967487771853 | down | NA |
| TMEM171 | -1.57942700771782 | 0.0483525873183295 | down | NA |
| GNE | -1.57731219752451 | 0.0115451559763567 | down | NA |
| COL12A1 | -1.57500497998156 | 0.0407325428536912 | down | Pathway |
| SERPINF2 | -1.56882387486446 | 0.0474776341741952 | down | Pathway |
| ARRDC3 | -1.56752247634379 | 0.0305049395736605 | down | Pathway |
| FOXF2 | -1.56338501636205 | 0.0156832778778444 | down | Pathway |
| C12orf55 | -1.55859541882309 | 0.0354553594021197 | down | NA |
| GK5 | -1.55271502107824 | 0.00979169933051757 | down | Pathway |
| MTHFD2L | -1.55093414651196 | 0.0185302032146686 | down | Pathway |
| SQSTM1 | -1.54865573372781 | 0.0176519651049434 | down | NA |
| RHCE | -1.54522484961405 | 0.0152871043929697 | down | Pathway |
| TNIK | -1.54286715774992 | 0.00283367842044329 | down | Pathway |
| PDLIM2 | -1.54043451914192 | 0.0461192758143927 | down | NA |
| CXXC5 | -1.53843475667468 | 0.0362981021104693 | down | NA |
| MXD4 | -1.53716146254266 | 0.0201750048585462 | down | NA |
| RASL11B | -1.53596429378793 | 0.0307412992125011 | down | Pathway |
| SVEP1 | -1.53311616389069 | 0.0417854019048567 | down | NA |
| HYPK | -1.52947244130942 | 0.0333455930308587 | down | NA |
| GRHL3 | -1.52475184114589 | 0.00702066841229523 | down | Pathway |
| PKP2 | -1.52215180937134 | 0.00575191630450542 | down | Pathway |
| SLC22A23 | -1.51719897038609 | 0.0203221355260243 | down | NA |
| ADM | -1.50726568316134 | 0.0344113680692851 | down | Pathway |
| TFAP2C | -1.50045088044781 | 0.0176519651049434 | down | NA |
| PNKP | 1.50410979243787 | 0.018188252251489 | up | NA |
| CORO2A | 1.50728807342785 | 0.0215564972815284 | up | NA |
| FBXO24 | 1.51204499444072 | 0.0361207566281088 | up | NA |
| MARC1 | 1.51687530016793 | 0.0387244156283981 | up | NA |
| HOXA9 | 1.51954436221579 | 0.00445986866069448 | up | Pathway |
| CDKN2D | 1.54384268137695 | 0.0133552152128459 | up | Pathway |
| CHD7 | 1.54504988145161 | 0.00591289489354877 | up | Pathway |
| DPF1 | 1.54707531393634 | 0.00777074350823438 | up | NA |
| S100P | 1.54874375510821 | 0.00778937934344466 | up | Pathway |
| ASIC1 | 1.55319287714883 | 0.0326778073702989 | up | NA |
| DIP2C | 1.55343852872653 | 0.0253573488752213 | up | NA |
| ACVR1C | 1.55930293783424 | 0.0226915804630063 | up | Pathway |
| DLX1 | 1.55953323846319 | 0.0147657330463027 | up | Pathway |
| SLITRK5 | 1.56191808488437 | 0.0215096637229127 | up | Pathway |
| UBE2S | 1.56266135597323 | 0.0457186412007819 | up | NA |
| RTN2 | 1.56974007162677 | 0.0449693831571452 | up | NA |

| | | | | |
|---|---|---|---|---|
| CAPN5 | 1.57368570878507 | 0.0358770664997914 | up | NA |
| HOXC5 | 1.57658877666766 | 0.0456952115225833 | up | Pathway |
| MFAP5 | 1.57920527803465 | 0.0453305002744757 | up | Pathway |
| HECW1 | 1.58619392950615 | 0.00789561487621582 | up | Pathway |
| EGR2 | 1.58653861492125 | 0.0251424344181079 | up | Pathway |
| KAZN | 1.5910863801269 | 0.0237923779921136 | up | NA |
| ZNF382 | 1.60250101490696 | 0.0139180203012815 | up | NA |
| KDELC2 | 1.60634304721551 | 0.0108434289526 | up | NA |
| CD3EAP | 1.60920845703608 | 0.0260852286893381 | up | NA |
| ONECUT2 | 1.61585999399463 | 0.042575469587858 | up | Pathway |
| LINGO2 | 1.62713509673622 | 0.0497002842683898 | up | Pathway |
| AP1M2 | 1.63683619888376 | 0.0174970232259838 | up | NA |
| SOX9 | 1.64596668204803 | 0.0434350860308345 | up | Pathway |
| TMEM150C | 1.6515194692314 | 0.0176728714069161 | up | NA |
| RND2 | 1.65400500214745 | 0.0138967236371032 | up | Pathway |
| TUBB2B | 1.65765568416471 | 0.0383727100135226 | up | Pathway |
| STAP2 | 1.66607490110829 | 0.0191725614207769 | up | NA |
| FAM43A | 1.67037625736452 | 0.00827020212835214 | up | NA |
| ZNF726 | 1.68680849352484 | 0.0162053198162007 | up | NA |
| AMN1 | 1.6927256561861 | 0.031000309837461 | up | NA |
| DUOX1 | 1.70003102600758 | 0.0162389203409003 | up | Pathway |
| PLA2G4C | 1.70438939543828 | 0.0182216278626592 | up | Pathway |
| MAN1A1 | 1.70924048591687 | 0.0062306377236223 | up | NA |
| RARRES1 | 1.74088205579501 | 0.0231445034793347 | up | NA |
| B4GALNT3 | 1.74171083996537 | 0.0180795351180147 | up | NA |
| PRDM5 | 1.74339890675768 | 0.038877877511596 | up | Pathway |
| RIMS3 | 1.7542339957991 | 0.0262422747603064 | up | NA |
| HOXA4 | 1.76311599233231 | 0.0367187431476375 | up | Pathway |
| DGKG | 1.78938180958211 | 0.019520871963596 | up | Pathway |
| TMEM169 | 1.81201775330409 | 0.0207380072336541 | up | NA |
| PMP22 | 1.8161688167926 | 0.013123721662465 | up | Pathway |
| ZNF570 | 1.83126352517518 | 0.0182775772887824 | up | NA |
| ZNF736 | 1.84605997523405 | 0.00484852546514122 | up | NA |
| PKN3 | 1.84669349560363 | 0.0431340318024439 | up | Pathway |
| MAN1C1 | 1.84729076979827 | 0.0231886035049557 | up | NA |
| C14orf37 | 1.85246366941902 | 0.0258575048667508 | up | NA |
| HOXB2 | 1.85869385157628 | 0.015148327326822 | up | Pathway |
| ZFP69 | 1.88016700337596 | 0.00546863024390161 | up | NA |
| LAD1 | 1.88263848575406 | 0.0443500696802338 | up | NA |
| IRF5 | 1.89818977323317 | 0.0210188309491636 | up | Pathway |
| CNFN | 1.91280607821432 | 0.0221097626779986 | up | NA |
| CRAT | 1.91611489819772 | 0.0440114424695586 | up | Pathway |
| POF1B | 1.92554569444853 | 0.0198658427915667 | up | Pathway |
| PITX2 | 1.93639174113406 | 0.010554643572276 | up | Pathway |
| KLC3 | 1.93859929834466 | 0.00498453710201918 | up | NA |
| ZNF85 | 1.94238255252242 | 0.018188252251489 | up | NA |
| CHRNB2 | 1.95368320534814 | 0.0230605480994224 | up | Pathway |
| C2CD4C | 1.96147975472164 | 0.030603444527413 | up | NA |
| ALDH1A3 | 1.96279474149 | 0.0250861591912763 | up | Pathway |
| PPM1J | 1.96581974762857 | 0.0245274541092311 | up | NA |
| GUCY2D | 1.96645726047907 | 0.0122716776209058 | up | Pathway |
| ILDR1 | 1.98112918802895 | 0.0105896693547357 | up | Pathway |
| SCN8A | 1.98615563013751 | 0.0231886035049557 | up | NA |

| | | | | |
|---|---|---|---|---|
| ZNF738 | 1.99729931731016 | 0.00527703435214645 | up | NA |
| CSF3 | 2.00224071679486 | 0.0147053724716601 | up | Pathway |
| HNF4G | 2.00891534718149 | 0.00200626005588499 | up | NA |
| NAPSA | 2.00992512967087 | 0.0136296663580451 | up | Pathway |
| LEF1 | 2.01107564524578 | 0.00165808241694114 | up | Pathway |
| MCF2 | 2.01982400664569 | 0.0366936404727024 | up | Pathway |
| PNMAL1 | 2.02020078469744 | 0.0197443936746592 | up | NA |
| STOX2 | 2.02839415189789 | 0.0228523296583246 | up | NA |
| FGF12 | 2.03723777773883 | 0.00888234694118875 | up | Pathway |
| GNG4 | 2.04118727083542 | 0.0201799378678864 | up | Pathway |
| CA2 | 2.04856185118612 | 0.0402452542329434 | up | Pathway |
| ACKR3 | 2.04902973248934 | 0.00517258555667806 | up | Pathway |
| SEPT3 | 2.07301596537564 | 0.00405714344667696 | up | NA |
| S1PR2 | 2.08906983555846 | 0.0221603263711135 | up | Pathway |
| SBSN | 2.12458553044432 | 0.00782765597435922 | up | NA |
| MT1F | 2.12813431976606 | 0.0297909624468199 | up | Pathway |
| DMC1 | 2.13401638169462 | 0.00734849165405664 | up | NA |
| C7orf31 | 2.13432300199142 | 0.0188202094513492 | up | NA |
| LY6D | 2.13766933845014 | 0.0223784739433146 | up | NA |
| ARID3A | 2.17786609517234 | 0.0152871043929697 | up | NA |
| TMOD2 | 2.19739599012135 | 0.00910729534474426 | up | Pathway |
| TMEM121 | 2.20758571928776 | 0.0311617985964184 | up | NA |
| RASIP1 | 2.21080616935792 | 0.00734849165405664 | up | NA |
| HOXA3 | 2.22624054288561 | 0.0254006500509984 | up | Pathway |
| MPO | 2.23068031833919 | 0.0254006500509984 | up | Pathway |
| KIF5C | 2.27063656742454 | 0.00896855015239253 | up | Pathway |
| ZNF790 | 2.27837745665157 | 0.0362981021104693 | up | NA |
| HOXB6 | 2.27848872211882 | 0.00240061511093794 | up | Pathway |
| CACNB4 | 2.30118095348005 | 0.0112429120342136 | up | Pathway |
| SCAMP5 | 2.30188997932893 | 0.00135340220393029 | up | NA |
| IL11 | 2.32439257056052 | 0.0197443936746592 | up | Pathway |
| WNT9A | 2.33109319343344 | 0.00546863024390161 | up | Pathway |
| CERCAM | 2.33231034263902 | 0.020321185017871 | up | NA |
| KIF26B | 2.35183503044323 | 0.00888234694118875 | up | NA |
| SMARCA1 | 2.36873365948593 | 0.0362511271953595 | up | NA |
| FUT1 | 2.374236427726 | 0.0277523049848282 | up | Pathway |
| FAM49A | 2.38494109055686 | 0.0271497464282466 | up | NA |
| TCF7 | 2.39336983203155 | 0.0139993108938463 | up | NA |
| TMEM74 | 2.39701095895352 | 0.0173294107278013 | up | NA |
| PMEPA1 | 2.40745620657191 | 0.00104973959334669 | up | Pathway |
| ATP6V1B1 | 2.42016803942517 | 0.0253740510953186 | up | Pathway |
| NTNG1 | 2.42323867933844 | 0.0273428427535571 | up | Pathway |
| FAM20C | 2.4259979057931 | 0.0139776732228738 | up | NA |
| GAL3ST4 | 2.48059386044945 | 0.0167337516259899 | up | Pathway |
| CCDC74B | 2.48704670123546 | 0.0106259042981105 | up | NA |
| REEP2 | 2.48895084228683 | 0.0472685879694989 | up | NA |
| TMEM145 | 2.49074414981529 | 0.0162389203409003 | up | NA |
| BCL11B | 2.49935834746551 | 0.0103245757363459 | up | Pathway |
| DENND1C | 2.52023357727887 | 0.0203221355260243 | up | NA |
| ISYNA1 | 2.52755837461445 | 0.0483357666963776 | up | Pathway |
| FN3K | 2.53753476710101 | 0.0227171426497814 | up | NA |
| RUNDC3A | 2.54436832648417 | 0.0201457704981271 | up | Pathway |
| IGFBP6 | 2.54497480834999 | 0.0125442401271443 | up | NA |

| | | | | |
|---|---|---|---|---|
| SEMA5B | 2.60300063807701 | 0.0397000040828547 | up | Pathway |
| DDN | 2.62813550096078 | 0.0114511616058178 | up | NA |
| ADAP2 | 2.64601624069951 | 0.0305049395736605 | up | Pathway |
| CCDC74A | 2.6921475562264 | 0.00500099367069865 | up | NA |
| MACC1 | 2.69586234758879 | 0.0379518318008679 | up | NA |
| WNT10A | 2.71162080313761 | 0.00910729534474426 | up | Pathway |
| TMEM54 | 2.71818719739216 | 0.0128082261076923 | up | NA |
| MAPK4 | 2.73069766438104 | 0.0337855529672982 | up | NA |
| NKX6-1 | 2.73491670319605 | 0.0291693300805134 | up | Pathway |
| ZBTB46 | 2.73811292207235 | 0.0118528227912586 | up | NA |
| HOXB3 | 2.75981680963186 | 0.0430808655473894 | up | Pathway |
| SYT11 | 2.76320713416538 | 0.015148327326822 | up | NA |
| MAPT | 2.7704825088656 | 0.00860208157094257 | up | Pathway |
| IGDCC4 | 2.79317022835875 | 0.00224714602281571 | up | NA |
| CYP4F2 | 2.79769319136304 | 0.0454652501932256 | up | Pathway |
| MEIS3 | 2.8225686267687 | 0.00676515248648476 | up | Pathway |
| FAM124A | 2.83447760573776 | 0.0053996054992121 | up | NA |
| UNC13D | 2.87965410184515 | 0.00191992729452257 | up | Pathway |
| ANKRD6 | 2.9000666591975 | 0.0444233195990525 | up | NA |
| LYPD6B | 2.90341667215587 | 0.0201799378678864 | up | NA |
| MYL9 | 2.90501203659934 | 0.0139993108938463 | up | Pathway |
| BMP6 | 2.91171718937032 | 0.019520871963596 | up | Pathway |
| GOLGA7B | 2.91669983223161 | 0.0204810479500326 | up | NA |
| APBB1 | 2.93502545969685 | 0.00381572485124622 | up | Pathway |
| BARX2 | 2.94358957650831 | 0.00871732607004357 | up | Pathway |
| GATA2 | 2.95225153798948 | 0.0122640526852166 | up | Pathway |
| B3GNT8 | 2.96018383092207 | 0.0420973154335111 | up | Pathway |
| C2orf70 | 2.98240063020513 | 0.00405714344667696 | up | NA |
| ILDR2 | 3.00314904008401 | 0.0400066123929638 | up | Pathway |
| COL5A3 | 3.00715992039817 | 0.005098243527636 | up | Pathway |
| GNAZ | 3.02869241504876 | 0.0042211714376257 | up | NA |
| SUSD4 | 3.03888826453616 | 0.00979169933051757 | up | NA |
| NOSTRIN | 3.04512456755733 | 0.00624317365825117 | up | NA |
| CTD-2303H24.2 | 3.04678837439348 | 0.0062760062540132 | up | NA |
| PAPPA2 | 3.09649591006573 | 0.0376869821208327 | up | Pathway |
| COX6B2 | 3.12975300629497 | 0.0212024004701557 | up | NA |
| DYNC1I1 | 3.13030596561342 | 0.0106259042981105 | up | NA |
| TMEM74B | 3.1710320938747 | 0.0416187654679112 | up | NA |
| BSN | 3.17725441398464 | 0.0096274444680014 | up | Pathway |
| LMX1B | 3.2404575597649 | 0.00428685517651297 | up | NA |
| DNAH3 | 3.25394045860248 | 0.00517258555667806 | up | NA |
| CCDC170 | 3.27528149517691 | 0.0102295968756798 | up | NA |
| B3GAT2 | 3.28491931761414 | 0.040755288029091 | up | NA |
| C15orf48 | 3.28948169891221 | 0.0300282042013176 | up | NA |
| MAK | 3.29738441471838 | 0.0053996054992121 | up | Pathway |
| SOX18 | 3.30195625329366 | 0.0425872288408441 | up | Pathway |
| CAMK4 | 3.32634540061397 | 0.0489481001743869 | up | NA |
| LRRC16B | 3.32712181161162 | 0.0297909624468199 | up | NA |
| SLC44A2 | 3.36850196182657 | 0.0054706052525586 | up | NA |
| MYO1G | 3.40897607286603 | 0.00189349418112886 | up | Pathway |
| ISL1 | 3.43800505747927 | 0.0457186412007819 | up | Pathway |
| RAB39B | 3.52285282846834 | 0.0210188309491636 | up | Pathway |
| STK32A | 3.53126687085302 | 0.000566246025402591 | up | NA |

| | | | | |
|---|---|---|---|---|
| ARTN | 3.57916258387679 | 0.0152705952579538 | up | Pathway |
| STMN3 | 3.60955201502082 | 0.0134270017038353 | up | NA |
| PTPRB | 3.63319374803193 | 0.00113925180685624 | up | NA |
| ARHGAP9 | 3.65689966355647 | 0.0202559703011228 | up | NA |
| KRTCAP3 | 3.71538040437006 | 0.00503074927760451 | up | NA |
| COPZ2 | 3.7473983250838 | 0.00200626005588499 | up | NA |
| SLC16A9 | 3.75151369682845 | 0.0435818424778398 | up | NA |
| SMIM17 | 3.77861052804065 | 0.0089536451107292 | up | NA |
| KCNMB3 | 3.79552829486219 | 0.025389981851985 | up | NA |
| ATP2C2 | 3.7978154098254 | 0.0152871043929697 | up | NA |
| SLC52A3 | 3.81027715746352 | 0.0106259042981105 | up | NA |
| VILL | 3.86980970898178 | 0.0131470809572953 | up | Pathway |
| GIPR | 3.88303822436617 | 0.0267622767466902 | up | Pathway |
| A2ML1 | 3.88768072071325 | 0.013626546440128 | up | NA |
| ADRA2C | 3.92964026309317 | 0.0143949538686486 | up | Pathway |
| CRYBA2 | 3.95157106616437 | 0.00210684384260289 | up | Pathway |
| CDKL2 | 4.07605795705377 | 0.00144411613966197 | up | NA |
| CCDC106 | 4.07657308154538 | 0.021111240347148 | up | NA |
| RGL3 | 4.10730843198005 | 0.0139776732228738 | up | NA |
| NES | 4.10956726759073 | 0.00595640055506055 | up | Pathway |
| KLK6 | 4.13573873911957 | 0.00431734994257802 | up | Pathway |
| TMEM176B | 4.16640457989916 | 0.0405759606948465 | up | NA |
| HRASLS | 4.17276982064808 | 0.0291693300805134 | up | NA |
| ELMOD1 | 4.3212031886277 | 0.0152636611117527 | up | NA |
| SLC2A6 | 4.32225750610165 | 0.0492499585613165 | up | NA |
| RNF128 | 4.36804657004704 | 0.0258824603631041 | up | NA |
| GPR63 | 4.37527387824218 | 0.0165514063897871 | up | NA |
| LZTS1 | 4.37983067420727 | 0.00265176893220175 | up | Pathway |
| SULT2B1 | 4.39937353954524 | 0.0147308841871448 | up | Pathway |
| EBF4 | 4.56950432649132 | 0.0119744930114424 | up | NA |
| PRKG2 | 4.65762229827064 | 0.00896855015239253 | up | Pathway |
| ACP5 | 4.74331668174934 | 0.00910729534474426 | up | Pathway |
| RHCG | 4.79047000054141 | 0.0453297049187041 | up | Pathway |
| LY75 | 4.94496721558681 | 0.00273170545121021 | up | NA |
| OVOL1 | 5.02132692186817 | 0.0170233078156277 | up | NA |
| ATAD3C | 5.04424578041965 | 0.0400066123929638 | up | NA |
| CDH8 | 5.11613145969445 | 0.0250861591912763 | up | Pathway |
| NUDT11 | 5.11781093793545 | 0.0352596894579683 | up | Pathway |
| FABP6 | 5.12873212750697 | 0.0277523049848282 | up | NA |
| TINCR | 5.14817045464318 | 0.0179687117421246 | up | NA |
| SATB1 | 5.16021534335003 | 0.0362981021104693 | up | NA |
| BRSK2 | 5.1882327309153 | 0.00749532301861601 | up | Pathway |
| DTNA | 5.2449164109858 | 0.0173294107278013 | up | Pathway |
| ZNF501 | 5.36482863873611 | 0.0022065719328289 | up | NA |
| ARL11 | 5.39536121590824 | 0.0405415370308813 | up | NA |
| SPRR1B | 5.42545388024377 | 0.0230605480994224 | up | NA |
| ADAM22 | 5.52214711004794 | 0.0210999421546606 | up | NA |
| KLK11 | 5.53324899328983 | 0.00106875874786922 | up | NA |
| TBX20 | 5.55510567672913 | 0.00012760111491141 | up | Pathway |
| TMEM216 | 5.69059514494095 | 0.0053996054992121 | up | NA |
| PARVB | 5.76552021059124 | 0.00144411613966197 | up | Pathway |
| CNTN1 | 5.76939255575019 | 0.000259361393150236 | up | Pathway |
| FAM129A | 5.89112561454683 | 0.0211252987656376 | up | NA |

| | | | | |
|---|---|---|---|---|
| TLR2 | 5.90790145086905 | 0.0453864161294708 | up | Pathway |
| NAP1L5 | 6.21700529708365 | 0.0213570845460382 | up | NA |
| AOC1 | 6.25129675651158 | 0.00469637367881301 | up | NA |
| KLK8 | 6.30833645387338 | 0.001597350253811 | up | Pathway |
| SEMA3E | 6.31131166411311 | 0.0350822106932278 | up | Pathway |
| KLK5 | 6.34715091015079 | 0.00517258555667806 | up | Pathway |
| DHRS2 | 6.38505916622061 | 0.0267622767466902 | up | Pathway |
| RIPPLY2 | 6.45088131404535 | 0.00142892100222063 | up | Pathway |
| KLK7 | 6.4655497068858 | 0.0245483411558718 | up | Pathway |
| RUNDC3B | 6.66234293758365 | 0.000651317326288731 | up | NA |
| HEY2 | 6.77905678428081 | 0.000137900118880927 | up | Pathway |
| SBSPON | 6.84177013749781 | 0.00102396540561436 | up | NA |
| DLX6 | 6.8794482937516 | 0.0000615221659977426 | up | Pathway |
| KRT13 | 7.00160685512286 | 0.000827483118042297 | up | NA |
| NME5 | 7.08877943114356 | 0.000433169655547905 | up | NA |
| SLCO1B3 | 7.21939253292425 | 0.00360046797777663 | up | NA |
| AKR1E2 | 7.2231052762594 | 0.00178142130125749 | up | NA |
| CD302 | 7.24904823675022 | 0.00776131275376969 | up | NA |
| TENM1 | 7.34142169266763 | 0.00189349418112886 | up | Pathway |
| NPM2 | 7.36582567425246 | 0.00702066841229523 | up | NA |
| SLC47A1 | 7.57842070339323 | 0.00405714344667696 | up | NA |
| CSRNP3 | 7.61153033123016 | 0.00360996867394443 | up | NA |
| GALNT12 | 7.61482313938681 | 0.00734849165405664 | up | NA |
| ERBB4 | 7.69009424371806 | 0.00113925180685624 | up | Pathway |
| BAIAP2L2 | 7.79310293801417 | 0.0054148113177371 | up | Pathway |
| PDE3B | 7.81217782542374 | 0.00254626552489978 | up | Pathway |
| ZNF256 | 7.89607916469048 | 0.00734849165405664 | up | NA |
| ZNF681 | 8.14051191375033 | 0.0000455338772482485 | up | NA |
| LRRC3 | 8.30221502517773 | 0.00147975674650345 | up | NA |
| ZNF506 | 8.40673203175828 | 0.00254626552489978 | up | NA |
| PCLO | 8.4690865689218 | 0.0231640718401788 | up | Pathway |
| WFDC2 | 8.56359831702183 | 0.0204810479500326 | up | NA |
| CDC42EP5 | 8.58450033049152 | 0.0259150270319065 | up | Pathway |
| ENPP4 | 8.81644981081593 | 0.0103069519864546 | up | Pathway |
| KANK4 | 8.9222651974298 | 0.00178142130125749 | up | Pathway |
| ZNF711 | 9.6603101649764 | 0.0120736455545121 | up | NA |
| MYH14 | 9.87232565011336 | 0.00240061511093794 | up | Pathway |
| KLK10 | 9.98500202500899 | 0.00210185778327598 | up | NA |
| SLAIN1 | 10.5517171918516 | 0.0000455338772482485 | up | NA |
| NELL2 | 10.7911671366455 | 0.00113925180685624 | up | Pathway |

*Genes used in the GO enrichment analysis in Fig 4 are indicated as "Pathway" if not they are listed as NA, not applicable
