## Supplementary material for "A recurrent pathogenic *BRCA2* truncating variant reveals a role for BRCA2-PCAF complex in modulating NF-κB-driven transcription": Table S4

| Gene | logFC | adj.P.Val | DEG | Pathway* |
|---|---|---|---|---|
| ADIPOQ | -11.2799984945786 | 0.000237620653586152 | down | Pathway |
| LEP | -11.04162666931 | 0.0000116557440237734 | down | Pathway |
| CIDEC | -10.9237113376168 | 0.000047501407051521 | down | Pathway |
| TUSC5 | -10.892530578022 | 0.0000407286167700333 | down | NA |
| C14orf180 | -10.3951166446475 | 0.0000207215494866922 | down | NA |
| ADH1B | -10.0348847231457 | 0.0000727757884061152 | down | Pathway |
| CIDEA | -9.81501533065702 | 0.0000643485145994096 | down | Pathway |
| MRAP | -9.48287652297978 | 4.09983852314949E-06 | down | Pathway |
| FABP4 | -9.4629221134978 | 0.0000343880181021525 | down | Pathway |
| GLYAT | -9.3383563752576 | 4.93078487897174E-06 | down | Pathway |
| GPD1 | -9.16483851105949 | 9.35003131715643E-06 | down | Pathway |
| AQP7 | -9.04760103210204 | 0.0000338376108967206 | down | Pathway |
| PLIN4 | -8.68200463307713 | 9.54545132091244E-06 | down | NA |
| MYOC | -8.59979985474771 | 0.000026490529190438 | down | Pathway |
| SAA1 | -8.58181891294256 | 2.81821035777139E-06 | down | Pathway |
| RBP4 | -8.49687879812433 | 0.0000292644696970919 | down | Pathway |
| CA4 | -8.49176095763971 | 0.000014088441156621 | down | Pathway |
| PLIN1 | -8.41598880305268 | 0.0000148509983906441 | down | Pathway |
| CD300LG | -8.3915670816539 | 0.0000867662331740805 | down | NA |
| PCK1 | -8.361689120799 | 0.000336144586198737 | down | Pathway |
| PPP1R1A | -8.34613891837922 | 0.0000256185428275925 | down | Pathway |
| HEPACAM | -8.12409196617378 | 7.89799979427237E-06 | down | NA |
| TRDN | -7.91719798164115 | 0.000193689291524232 | down | Pathway |
| LGALS12 | -7.88060073973149 | 0.00005852044257214 | down | Pathway |
| TIMP4 | -7.85491625848605 | 7.81149829611418E-06 | down | Pathway |
| APOB | -7.78831691146445 | 0.000035799033508745 | down | Pathway |

| | | | | |
|---|---|---|---|---|
| TMEM132C | -7.75689081133308 | 0.0000539430429661821 | down | NA |
| SLC7A10 | -7.71233467254346 | 0.0000276807225929861 | down | Pathway |
| HSPB7 | -7.70166949287452 | 2.81821035777139E-06 | down | Pathway |
| ADRA1A | -7.68893525290817 | 4.93078487897174E-06 | down | Pathway |
| SLC19A3 | -7.67870954509574 | 0.0000317201556733218 | down | Pathway |
| ALDH1L1 | -7.65644003152199 | 0.0000136429042467373 | down | Pathway |
| CHRDL1 | -7.63492251896158 | 0.0000111856259866863 | down | Pathway |
| LIPE | -7.6021268005524 | 9.06819054236689E-06 | down | Pathway |
| SCARA5 | -7.55797087194618 | 0.000053782526435052 | down | NA |
| PLA2G2A | -7.52870009022432 | 0.000260191438164158 | down | Pathway |
| PCOLCE2 | -7.48802416424739 | 4.93078487897174E-06 | down | NA |
| C12orf39 | -7.41335210397584 | 0.000969962540091329 | down | NA |
| HBA2 | -7.35652021202049 | 0.0000138892049247438 | down | Pathway |
| DEFB132 | -7.33465563420126 | 0.000286324374916553 | down | NA |
| C19orf80 | -7.229904995647 | 6.64655070757393E-06 | down | NA |
| TNMD | -7.21905152294525 | 0.0000154285478081755 | down | Pathway |
| KLB | -7.17262497335337 | 0.000467178037277391 | down | Pathway |
| HBB | -7.13031291421792 | 0.0000136527038725634 | down | Pathway |
| LHCGR | -7.11106383434347 | 5.57848613541443E-06 | down | Pathway |
| FRMD1 | -7.05576665608617 | 0.0000483036379967917 | down | NA |
| G0S2 | -6.98343738057125 | 4.87087154292548E-06 | down | Pathway |
| CD36 | -6.96248418130756 | 0.0000253346585062879 | down | Pathway |
| AADAC | -6.95341075588744 | 0.000644903205574493 | down | Pathway |
| HBA1 | -6.93541623219882 | 0.0000231355064402411 | down | Pathway |

| | | | | |
|---|---|---|---|---|
| SGCG | -6.89600023604883 | 0.0000276380671060869 | down | Pathway |
| GLP2R | -6.85677035912559 | 0.0000035134961308845 | down | Pathway |
| HSPB6 | -6.81741042521839 | 0.0000108446700708311 | down | Pathway |
| SAA2 | -6.74908718204407 | 0.0000496954027098307 | down | Pathway |
| TRHDE | -6.73574957145066 | 0.0000785780320716064 | down | Pathway |
| AKR1C2 | -6.69409644957238 | 6.97446154261324E-06 | down | Pathway |
| LPL | -6.68534489773007 | 0.0000130933903747851 | down | Pathway |
| CSN1S1 | -6.61339162163147 | 0.0000035100225008365 | down | Pathway |
| ACVR1C | -6.57482246037446 | 0.0000486314964949464 | down | Pathway |
| ACADL | -6.56964579966812 | 0.000861267488020354 | down | Pathway |
| RP1-302G2.5 | -6.5694022325165 | 2.81821035777139E-06 | down | NA |
| ADH1A | -6.56022667449111 | 0.0000231355064402411 | down | Pathway |
| THRSP | -6.54891149059753 | 0.00274707467455701 | down | Pathway |
| CPA1 | -6.52777781376075 | 7.68521024514109E-06 | down | Pathway |
| ADH1C | -6.504056065898 | 0.000035273576192804 | down | Pathway |
| LBP | -6.43700257141555 | 0.0000606073407913894 | down | Pathway |
| CFD | -6.40257266997557 | 0.000107098315490048 | down | NA |
| PI16 | -6.36924449234117 | 0.000166776688431061 | down | Pathway |
| AQPEP | -6.31070459343645 | 0.0000333215231415449 | down | NA |
| KCNB1 | -6.29510590119429 | 8.41402063974025E-06 | down | Pathway |
| FHL1 | -6.28826939180485 | 3.43545566418516E-06 | down | Pathway |
| ATP1A2 | -6.24817336209922 | 0.0000281834185229659 | down | Pathway |
| EPB42 | -6.23084003974254 | 3.46935980267288E-06 | down | Pathway |
| ABCA8 | -6.22752696528936 | 9.40865569656448E-06 | down | Pathway |

| | | | | |
|---|---|---|---|---|
| FXYD1 | -6.22283948732946 | 0.0000804534278010506 | down | Pathway |
| NNAT | -6.21130609655285 | 0.00362554039333159 | down | Pathway |
| GYG2 | -6.21042313518195 | 0.0000539241510291624 | down | Pathway |
| CES1 | -6.20019151235811 | 0.0000439434001789324 | down | Pathway |
| PCDH9 | -6.14018855505827 | 4.19222087788146E-06 | down | Pathway |
| S100B | -6.13031499886417 | 0.000574338703708593 | down | Pathway |
| IRX6 | -6.08699641304794 | 0.0000546581306106511 | down | NA |
| DGAT2 | -6.08525752949882 | 0.0000138892049247438 | down | Pathway |
| C6 | -6.04795347045673 | 0.000790526835161806 | down | Pathway |
| HP | -6.03239459981034 | 0.00316977218857638 | down | Pathway |
| FGFBP2 | -6.01516274980477 | 0.0000216825353897115 | down | NA |
| CDH20 | -6.00120843351918 | 0.000050846170747786 | down | Pathway |
| PRRT4 | -5.99543861891853 | 0.0000950283437031099 | down | NA |
| CDO1 | -5.99162353208315 | 0.0000208276182863052 | down | Pathway |
| KIAA1239 | -5.98645509596242 | 0.0000838494407218363 | down | NA |
| PLXNA4 | -5.98625381324915 | 2.81821035777139E-06 | down | Pathway |
| NECAB1 | -5.96674789312811 | 0.000403474561536482 | down | Pathway |
| CLEC3B | -5.93955152451728 | 0.0000043740272661602 | down | Pathway |
| ITIH5 | -5.93017942364147 | 3.46935980267288E-06 | down | NA |
| TMEM100 | -5.89863018393158 | 0.0000256658752137784 | down | Pathway |
| SLC2A4 | -5.88620463454955 | 0.0000683419627159911 | down | Pathway |
| CLEC4G | -5.86368013399247 | 0.000123180519229478 | down | Pathway |
| MLXIPL | -5.83615725473236 | 0.000153027504055082 | down | Pathway |

| | | | | |
|---|---|---|---|---|
| LYVE1 | -5.79629678092306 | 0.0000183329851889496 | down | Pathway |
| CNTFR | -5.79214714357932 | 0.0000710861640679176 | down | Pathway |
| TMC2 | -5.77919501124227 | 6.94705115187937E-06 | down | Pathway |
| AOC3 | -5.73333098190958 | 4.87087154292548E-06 | down | Pathway |
| PRG4 | -5.71557200922911 | 0.000931600112341234 | down | NA |
| SDPR | -5.66891802046878 | 4.09983852314949E-06 | down | NA |
| ANO3 | -5.65413954502693 | 0.000104447813318692 | down | Pathway |
| NPY5R | -5.65138499639375 | 0.00142304010132895 | down | Pathway |
| NLGN1 | -5.64663402009522 | 0.0000143427643652586 | down | Pathway |
| VIT | -5.63529327980966 | 0.0000138892049247438 | down | Pathway |
| SLC25A18 | -5.62251045634725 | 0.000589908254731518 | down | Pathway |
| CALB2 | -5.6004799572688 | 0.000231776048398995 | down | Pathway |
| PTPRQ | -5.55792365440674 | 0.000350006354206234 | down | Pathway |
| DPT | -5.5505834094039 | 0.0000657083374773792 | down | Pathway |
| RERGL | -5.54812509569477 | 0.000181968093700162 | down | NA |
| SLC16A7 | -5.5243368209878 | 0.0000838494407218363 | down | Pathway |
| GPAM | -5.4970091966537 | 7.81149829611418E-06 | down | Pathway |
| HSD17B13 | -5.47561697751682 | 0.0000174267445510039 | down | Pathway |
| ANGPTL5 | -5.42104162415191 | 0.0000300655359075657 | down | NA |
| BTNL9 | -5.41269247874485 | 0.0000415862988437535 | down | NA |
| NAT8L | -5.38602529007275 | 0.00118459136238005 | down | Pathway |
| MAOA | -5.37333476692732 | 9.54160563038411E-06 | down | Pathway |
| ABCD2 | -5.36928399810394 | 0.000082096864667323 | down | Pathway |
| C1QTNF9 | -5.36432131152871 | 6.64655070757393E-06 | down | NA |
| ANGPTL1 | -5.36141589861936 | 0.0000216013010133142 | down | NA |

| | | | | |
|---|---|---|---|---|
| MASP1 | -5.35848273482595 | 0.000776285145144357 | down | Pathway |
| TF | -5.3570756995815 | 0.00015963265949586З | down | Pathway |
| AKR1C1 | -5.35022234436963 | 0.0000148509983906441 | down | Pathway |
| PFKFB1 | -5.34656824572607 | 0.0000127212743298798 | down | Pathway |
| SHISA3 | -5.3233117720389 | 0.0000105415976727814 | down | Pathway |
| EYS | -5.31544341521218 | 0.000287680835523724 | down | Pathway |
| AGXT | -5.3023745900857 | 2.81821035777139E-06 | down | Pathway |
| CMA1 | -5.29446730293529 | 0.000615223146382208 | down | Pathway |
| CRYAB | -5.28535890423489 | 0.0000274499364663672 | down | Pathway |
| CASQ2 | -5.28364250830396 | 0.0000852629833450245 | down | Pathway |
| PLAC9 | -5.27548817796761 | 0.0000540795957791983 | down | NA |
| GRIN2B | -5.27180374698126 | 0.000348079392491984 | down | Pathway |
| TNNT3 | -5.26911612491664 | 0.000544691369367457 | down | Pathway |
| ANGPTL4 | -5.21267393681027 | 6.28823464173383E-06 | down | Pathway |
| KCNIP2 | -5.19920706047672 | 0.0000135579150977941 | down | Pathway |
| GPC3 | -5.19087047785838 | 0.0000596138417871747 | down | Pathway |
| PDE3B | -5.18714712908329 | 0.000345803265669088 | down | Pathway |
| SORBS1 | -5.16681019433352 | 0.0000035100225008365 | down | Pathway |
| NMUR1 | -5.16291090954816 | 6.28823464173383E-06 | down | Pathway |
| LRRN4CL | -5.0960477306019 | 0.0000197702668193731 | down | NA |
| KCNA4 | -5.08634667382832 | 0.000118806257223279 | down | Pathway |
| CLDN2 | -5.06143832093782 | 0.0000197273938950305 | down | Pathway |
| GPX3 | -5.06139966425636 | 0.0000363555584077947 | down | Pathway |
| C7 | -5.05808616192662 | 0.0012221591731031 | down | NA |
| SLC4A4 | -5.05230508714062 | 0.0000300655359075657 | down | Pathway |

| | | | | |
|---|---|---|---|---|
| RXRG | -5.04595553331751 | 0.000145928809403184 | down | Pathway |
| SLC29A4 | -5.03621671966199 | 6.86538373931544E-06 | down | Pathway |
| SCN4A | -5.03334332874219 | 0.000546802942568236 | down | Pathway |
| BMP5 | -5.03068356456753 | 0.000132942602781429 | down | Pathway |
| FCER2 | -5.00225589922485 | 0.00170874121676437 | down | Pathway |
| DMRT2 | -4.98040493530196 | 0.00106915086935269 | down | Pathway |
| RBP7 | -4.9751811610409 | 0.0000143427643652586 | down | Pathway |
| NPR3 | -4.9709691783695 | 0.000251905287744471 | down | Pathway |
| ADRB1 | -4.96780638738935 | 0.0000356687838731038 | down | Pathway |
| ASPA | -4.95588510267213 | 0.0000151140255854889 | down | Pathway |
| AC131097.4 | -4.95419498887348 | 0.000591030766927739 | down | NA |
| ANGPT1 | -4.95190272751288 | 6.64655070757393E-06 | down | Pathway |
| PPARG | -4.93081938788796 | 0.000115299872397173 | down | Pathway |
| NAALAD2 | -4.92958608966029 | 0.00110812514122035 | down | NA |
| GHR | -4.90852944072827 | 0.000568722138742832 | down | Pathway |
| SYN2 | -4.90782149531593 | 0.000162681118293746 | down | Pathway |
| SLC14A2 | -4.89521964251252 | 0.0000302660001749592 | down | NA |
| ANGPTL7 | -4.87337384447628 | 0.000753962027795544 | down | Pathway |
| SCGN | -4.86656312248055 | 0.0000256185428275925 | down | Pathway |
| CCDC3 | -4.85656284089692 | 0.0000035100225008365 | down | Pathway |
| SCEL | -4.832319642538 | 0.00801423985618286 | down | Pathway |
| HPSE2 | -4.82087851483714 | 0.000145021548110666 | down | Pathway |
| AOX1 | -4.80799101739722 | 0.0000216825353897115 | down | Pathway |
| ANGPT4 | -4.80278724577424 | 0.0000559870939467668 | down | Pathway |
| BCHE | -4.8026786730482 | 0.000286164054547944 | down | Pathway |

| | | | | |
|---|---|---|---|---|
| SOD3 | -4.79692529271791 | 9.35003131715643E-06 | down | Pathway |
| NTRK2 | -4.79587682368983 | 0.000174685514057309 | down | Pathway |
| CKMT2 | -4.79258170577608 | 0.0000599824438302514 | down | Pathway |
| CLDN5 | -4.78319476742042 | 0.0000035100225008365 | down | Pathway |
| CAV1 | -4.78128094441621 | 4.09983852314949E-06 | down | Pathway |
| SFRP1 | -4.76831666191754 | 0.0000540795957791983 | down | Pathway |
| DNASE1L3 | -4.76548699624983 | 0.00090945726120555 | down | Pathway |
| CRHBP | -4.7508475115544 | 0.000471013104407419 | down | Pathway |
| TSLP | -4.74926907813875 | 0.0000300655359075657 | down | Pathway |
| ZBTB16 | -4.74703213518503 | 0.00392715490930315 | down | Pathway |
| MARC1 | -4.74654075460485 | 0.0000903371821431127 | down | NA |
| MAP1LC3C | -4.73119620545657 | 0.0000231355064402411 | down | Pathway |
| PLCXD3 | -4.72720798648349 | 0.000719977459913792 | down | Pathway |
| TMEM37 | -4.72577514878088 | 0.0000922735054215946 | down | Pathway |
| HRASLS5 | -4.72395839821317 | 0.000140674677925602 | down | NA |
| TNXB | -4.72158564039523 | 0.0000408366581746066 | down | Pathway |
| HRCT1 | -4.71923307981687 | 0.0000852629833450245 | down | NA |
| MEOX2 | -4.7105923847279 | 9.40865569656448E-06 | down | Pathway |
| HHIP | -4.69893025503394 | 0.000442845218508403 | down | Pathway |
| TSPAN8 | -4.69050830972325 | 0.00218930866629397 | down | Pathway |
| ITGA7 | -4.68486411242365 | 9.40570684581767E-06 | down | Pathway |
| PRKAR2B | -4.67609544419828 | 0.0000705182148924997 | down | Pathway |
| NPR1 | -4.67484613808943 | 0.0000746099172986083 | down | Pathway |
| ELANE | -4.67479756796384 | 0.0000167341009768103 | down | Pathway |
| FIGF | -4.6642775561754 | 4.47749536812041E-06 | down | NA |

| | | | | |
|---|---|---|---|---|
| GYS2 | -4.64336046316919 | 0.00115836595329297 | down | Pathway |
| GRIA1 | -4.63352735184862 | 0.00221847159308067 | down | Pathway |
| TACR1 | -4.63259389477629 | 0.000336027226324442 | down | Pathway |
| SLC22A3 | -4.63071484546116 | 0.0000256779395398172 | down | Pathway |
| SLC1A7 | -4.61773838281641 | 0.0000727757884061152 | down | Pathway |
| HIF3A | -4.60931540706125 | 0.00177737088492446 | down | Pathway |
| CAV2 | -4.60740408793919 | 0.0000035100225008365 | down | Pathway |
| IL33 | -4.6057480995729 | 0.0000216812107614539 | down | Pathway |
| PPP1R1B | -4.59413807866562 | 0.00177497231862233 | down | Pathway |
| FAM107A | -4.5727325905125 | 0.000214593620021976 | down | Pathway |
| FCN2 | -4.57272468160267 | 0.00626296663035763 | down | Pathway |
| FMO2 | -4.56276252430863 | 0.000503557897516188 | down | Pathway |
| APOD | -4.55434387938808 | 0.000053782526435052 | down | Pathway |
| RDH5 | -4.54850251643279 | 4.93078487897174E-06 | down | Pathway |
| HCAR2 | -4.54182863025332 | 0.0000581124054029902 | down | Pathway |
| MYOM1 | -4.53728004985574 | 0.0000118863962808334 | down | Pathway |
| BHMT2 | -4.53172530501595 | 4.42708939954965E-06 | down | Pathway |
| GABRA2 | -4.53086216620513 | 0.00131486752518726 | down | Pathway |
| KLF15 | -4.52402504300866 | 0.000493412706639896 | down | Pathway |
| SEMA3G | -4.52077053856664 | 4.93078487897174E-06 | down | Pathway |
| GALNT15 | -4.51725638379422 | 0.0000573557135571416 | down | NA |
| ATOH8 | -4.50894194598401 | 0.0000363429564403213 | down | Pathway |
| MT1M | -4.50874798985914 | 0.0000375375960516261 | down | Pathway |
| HSD11B1 | -4.49988278916708 | 0.00023087769011815 | down | Pathway |

| | | | | |
|---|---|---|---|---|
| C1QTNF7 | -4.49426143052251 | 0.000612958213159945 | down | NA |
| DARC | -4.47576987106342 | 0.00117951915899427 | down | NA |
| CCDC69 | -4.47422239497144 | 0.0000276279299739945 | down | Pathway |
| PDK4 | -4.46735164696395 | 0.0000736794698513773 | down | Pathway |
| FAM180B | -4.46326897506829 | 0.0000805725609347576 | down | NA |
| ACSM5 | -4.45073839223754 | 0.000420534806909944 | down | Pathway |
| DES | -4.4398129649725 | 0.0176217438468198 | down | Pathway |
| SVEP1 | -4.43585438224386 | 4.09983852314949E-06 | down | Pathway |
| MME | -4.43138021968257 | 0.000660059208830781 | down | Pathway |
| C2orf40 | -4.43031117521089 | 0.000574338703708593 | down | NA |
| KIF25 | -4.40985128148907 | 0.0000228719746794813 | down | Pathway |
| SNTG2 | -4.39767937742971 | 0.000331787661833553 | down | NA |
| PKNOX2 | -4.38051533475157 | 0.0000936258389767269 | down | NA |
| AVPR2 | -4.34876898435487 | 0.000502108817876077 | down | Pathway |
| OGN | -4.33104978755481 | 0.00328464595664734 | down | Pathway |
| C8orf34 | -4.32775631377141 | 0.000166533427059547 | down | NA |
| FREM1 | -4.32018173432822 | 0.0000231355064402411 | down | Pathway |
| TWIST2 | -4.31760014726583 | 0.000974221204416011 | down | Pathway |
| CHL1 | -4.30534423264893 | 0.0000136429042467373 | down | Pathway |
| C3orf55 | -4.30532039552845 | 0.000207183989810153 | down | NA |
| F10 | -4.29627600536128 | 0.0000785780320716064 | down | Pathway |
| HPR | -4.29336046457851 | 0.00248304237219427 | down | Pathway |
| CPED1 | -4.27893041447912 | 0.0000220455135734751 | down | NA |
| BMX | -4.27328620831966 | 7.57103304085244E-06 | down | Pathway |

| | | | | |
|---|---|---|---|---|
| DTX1 | -4.27271976955953 | 0.0000297807856142578 | down | Pathway |
| CCDC85A | -4.27014420384844 | 0.000467071501866575 | down | NA |
| PDE2A | -4.25891016733054 | 7.03617665926645E-06 | down | Pathway |
| TPO | -4.25229980850458 | 0.000805427054623683 | down | Pathway |
| CEBPA | -4.24977280918949 | 0.00103328181299301 | down | Pathway |
| CCDC178 | -4.2492206782888 | 0.0000315490420068815 | down | NA |
| XPNPEP2 | -4.24594118468079 | 0.000841961641436113 | down | NA |
| FGF2 | -4.23469720337943 | 0.000208755177709903 | down | Pathway |
| FZD4 | -4.23114092495479 | 4.93078487897174E-06 | down | Pathway |
| GFAP | -4.22586921794054 | 0.00810123338750151 | down | Pathway |
| CNTNAP3B | -4.22034205754551 | 0.000248543237413034 | down | NA |
| ENPP2 | -4.19577885483851 | 0.0000138892049247438 | down | Pathway |
| ACACB | -4.19577088402362 | 0.0000725483837037814 | down | Pathway |
| ALDH1A1 | -4.19375229923244 | 0.000888268179667604 | down | Pathway |
| PDZD2 | -4.19164325573677 | 0.0000281969501506849 | down | NA |
| GPR146 | -4.18747792560292 | 6.28823464173383E-06 | down | NA |
| INMT | -4.18265804870926 | 0.0000702001460353275 | down | Pathway |
| LRRN3 | -4.17875052773583 | 0.0000035100225008365 | down | Pathway |
| LEFTY2 | -4.1653917739465 | 0.0008890430012595 | down | Pathway |
| NRN1 | -4.16389771669909 | 0.000337787104676769 | down | Pathway |
| PRR26 | -4.14863250265683 | 0.0000317201556733218 | down | NA |
| PDE11A | -4.14538313861459 | 0.000888253953419409 | down | Pathway |
| SLC17A7 | -4.14514316769474 | 0.000035799033508745 | down | Pathway |
| IGFBP6 | -4.14207751689887 | 0.000133214633794614 | down | Pathway |

| | | | | |
|---|---|---|---|---|
| PALMD | -4.14181839373815 | 9.40865569656448E-06 | down | Pathway |
| AKAP12 | -4.13623408059883 | 4.19222087788146E-06 | down | Pathway |
| CDKN1C | -4.13164517991594 | 0.000043451807242662 | down | Pathway |
| FAM89A | -4.13025876217922 | 0.0000242339676738567 | down | NA |
| CLCA2 | -4.12502805412794 | 0.000334143519914029 | down | Pathway |
| PPP1R14A | -4.1132251342614 | 0.000243299091832372 | down | NA |
| PKD1L2 | -4.11299553932838 | 0.00407139800958276 | down | Pathway |
| P2RX6 | -4.10864488786676 | 0.000169316799157137 | down | Pathway |
| PRELP | -4.09619244852288 | 0.0000334916128673879 | down | NA |
| WNT11 | -4.08370330131689 | 0.00199327714734499 | down | Pathway |
| CPA2 | -4.08020423765729 | 0.000623555058149339 | down | NA |
| MYO16 | -4.06772952328102 | 0.0000231600052310895 | down | Pathway |
| MAB21L1 | -4.06673623502035 | 0.000190284078984827 | down | Pathway |
| AKR1C3 | -4.066023259862 | 4.19222087788146E-06 | down | Pathway |
| DMRTA1 | -4.06311010577887 | 0.00050374057871714 | down | Pathway |
| DMGDH | -4.06052880393303 | 0.0000216825353897115 | down | Pathway |
| C10orf10 | -4.04568582036774 | 0.0000054276263364332 | down | NA |
| PYGM | -4.04389037521481 | 0.0000281834185229659 | down | Pathway |
| IGF1 | -4.03854532661337 | 0.000295212942885313 | down | Pathway |
| KLHL30 | -4.03733179136833 | 0.000203752307486219 | down | NA |
| KLHL33 | -4.03130993770455 | 0.000295212942885313 | down | NA |
| LRP1B | -4.02112284390598 | 0.00947119322050621 | down | NA |
| RELN | -4.01931740579915 | 0.000815822151727438 | down | Pathway |
| ABCC6 | -4.01168093319001 | 0.0000229075862355889 | down | Pathway |

| | | | | |
|---|---|---|---|---|
| NMNAT2 | -4.00946638294773 | 0.000303085671835874 | down | NA |
| RASD1 | -4.0061211531694 | 0.000108792788264765 | down | Pathway |
| BMPER | -4.00443602973403 | 0.000142409141081999 | down | Pathway |
| PENK | -4.00409539087765 | 0.00895166323314354 | down | Pathway |
| AVPR1A | -3.99962115511668 | 0.000101669495917888 | down | Pathway |
| CBLN1 | -3.99956504416236 | 0.000111254350827603 | down | Pathway |
| TNS1 | -3.99846098407643 | 0.0000164035762947684 | down | Pathway |
| CYP2A6 | -3.99701701651163 | 0.0420665200077647 | down | Pathway |
| HLF | -3.99564647148942 | 0.0000182494380782862 | down | Pathway |
| TMOD1 | -3.98665347468514 | 0.000248543237413034 | down | Pathway |
| BMP2 | -3.96701673621564 | 0.000083142834261072 | down | Pathway |
| ASPG | -3.95981348713451 | 0.0004784401782741 | down | Pathway |
| HOXA10 | -3.95905956401771 | 0.0000822637069617463 | down | Pathway |
| HOGA1 | -3.95382152802359 | 0.000237620653586152 | down | Pathway |
| AGMO | -3.94886454977826 | 0.000334143519914029 | down | Pathway |
| MLIP | -3.94288259186442 | 0.00106915086935269 | down | Pathway |
| C1QTNF9B | -3.92297627028357 | 8.13603719666152E-06 | down | NA |
| TCF15 | -3.91650227230598 | 0.000270155397232282 | down | Pathway |
| MEOX1 | -3.91636060626135 | 0.000597980642242288 | down | Pathway |
| CLMP | -3.90267562843546 | 0.000077538075857269 | down | Pathway |
| BOK | -3.90135032435998 | 0.000076246527908556 | down | Pathway |
| TGFBR3 | -3.89428690790126 | 0.000337787104676769 | down | Pathway |
| BMP3 | -3.87521056652805 | 0.0206499838504648 | down | Pathway |
| EBF1 | -3.8683134600187 | 9.35003131715643E-06 | down | NA |

| | | | | |
|---|---|---|---|---|
| NMB | -3.86818207470585 | 0.0000281834185229659 | down | Pathway |
| CTSG | -3.86621616687013 | 0.00367689322002081 | down | Pathway |
| ALDH1A2 | -3.86608312629764 | 0.0000776285145144357 | down | Pathway |
| VLDLR | -3.86469605167026 | 0.0000300655359075657 | down | Pathway |
| NKAPL | -3.86049444743007 | 0.000014088441156621 | down | NA |
| SNCG | -3.85958757577897 | 0.00212912302558871 | down | Pathway |
| NTRK3 | -3.85938608594833 | 0.00206532857401165 | down | Pathway |
| EBF3 | -3.85873267926423 | 0.0000297807856142578 | down | NA |
| KCNAB1 | -3.84618753012724 | 0.0000302660001749592 | down | Pathway |
| ADIRF | -3.83969064579344 | 0.000131656086833542 | down | Pathway |
| SCN4B | -3.8285080852239 | 0.000137832569291202 | down | Pathway |
| LUZP2 | -3.82541438557985 | 0.0000985411462195632 | down | NA |
| ECM2 | -3.81816568389154 | 0.0000208276182863052 | down | Pathway |
| ARHGDIG | -3.81622222812331 | 0.0143867933717587 | down | Pathway |
| DDR2 | -3.81391766650675 | 9.54545132091244E-06 | down | Pathway |
| RGS7BP | -3.80880822504656 | 0.00130689265584673 | down | Pathway |
| AGPAT2 | -3.80465158454141 | 3.53589780530754E-06 | down | Pathway |
| PTH1R | -3.80347625210138 | 0.000129439610476009 | down | Pathway |
| KCNA2 | -3.80335927722175 | 0.00121028633221942 | down | Pathway |
| PGM5 | -3.80053375824283 | 0.0000135579150977941 | down | Pathway |
| PTGER3 | -3.79537805752215 | 0.00222750025134191 | down | Pathway |
| HOXA6 | -3.79112497240483 | 0.000406484763669773 | down | Pathway |
| FOXP2 | -3.78896095542037 | 0.0000852629833450245 | down | Pathway |
| C16orf89 | -3.78508628838183 | 0.00467081786442399 | down | NA |

| | | | | |
|---|---|---|---|---|
| SRPX | -3.7766560194451 | 0.000101569544108791 | down | NA |
| S100A1 | -3.77330752478411 | 0.0100714300020009 | down | Pathway |
| ABCA9 | -3.77117167231045 | 0.000018291536526029 | down | Pathway |
| ANKRD53 | -3.76894493510949 | 0.000788212727933024 | down | Pathway |
| GNAL | -3.76782091755097 | 0.0000518100577819199 | down | Pathway |
| WISP2 | -3.76471112788533 | 0.00389757481976342 | down | NA |
| GPR133 | -3.75513821242355 | 0.00042051996284283 | down | NA |
| FAM47E-STBD1 | -3.75505521734822 | 0.0000293106134078413 | down | NA |
| SLCO1C1 | -3.75152528974678 | 0.000618712960997213 | down | Pathway |
| TFPI2 | -3.75069535949488 | 0.00271174209364322 | down | Pathway |
| SRL | -3.74380952947644 | 0.000419733193335058 | down | NA |
| TRPM3 | -3.74170591506731 | 0.0010098666038566 | down | Pathway |
| GNAI1 | -3.74170310034176 | 0.0000160561685248542 | down | Pathway |
| ADAMTS18 | -3.73965130826001 | 0.00218930866629397 | down | Pathway |
| STXBP6 | -3.73053837507406 | 0.000239011377535071 | down | Pathway |
| SPTB | -3.72748827479201 | 0.00035511239248199 | down | Pathway |
| ADCY5 | -3.72157590114608 | 0.00236376952926664 | down | Pathway |
| COX7A1 | -3.71494575961583 | 0.000244249546016284 | down | NA |
| HOXA7 | -3.71487827616766 | 0.000334044156735741 | down | Pathway |
| GSTM5 | -3.71466076186456 | 0.000179838232954031 | down | Pathway |
| DMD | -3.70382411581314 | 0.0000281834185229659 | down | Pathway |
| MMD | -3.70240448935793 | 8.82277213345037E-06 | down | Pathway |
| LINC00961 | -3.70209100576518 | 0.0000165697766706899 | down | NA |
| TYRO3 | -3.7018155097038 | 6.28823464173383E-06 | down | Pathway |

| | | | | |
|---|---|---|---|---|
| KRT79 | -3.69924349421895 | 0.0013012448199426 | down | Pathway |
| FHL5 | -3.67506393100664 | 0.0000165697766706899 | down | NA |
| ALDH2 | -3.67119530260337 | 0.0000669530180311546 | down | Pathway |
| CADM2 | -3.66367535393974 | 0.00752424512668045 | down | NA |
| ZNF676 | -3.66198885790012 | 0.0000835210703882731 | down | NA |
| DOC2B | -3.65926374698171 | 0.00150353420711914 | down | Pathway |
| ARHGAP20 | -3.65862152471 | 0.000145928809403184 | down | Pathway |
| FGF10 | -3.65821207212732 | 0.0275520600374499 | down | Pathway |
| FCGR3B | -3.65420280362623 | 0.000507568307577507 | down | NA |
| ABCC9 | -3.64513567649009 | 0.0000486314964949464 | down | Pathway |
| FAM228A | -3.64033611005617 | 0.000158378878195677 | down | NA |
| CCL24 | -3.63215705182308 | 0.00220286488602987 | down | Pathway |
| GPBAR1 | -3.62954114793159 | 7.02934969351015E-06 | down | Pathway |
| FFAR3 | -3.62658971112777 | 0.000350006354206234 | down | Pathway |
| DNAH9 | -3.62476158852461 | 0.00119702906199665 | down | Pathway |
| MATN2 | -3.62448284915844 | 0.000290682434526154 | down | NA |
| GSN | -3.62374538605296 | 6.23165598484117E-06 | down | Pathway |
| COLEC11 | -3.62237647806452 | 0.0000529984249830006 | down | Pathway |
| TENM1 | -3.62195844355212 | 0.00214689919819285 | down | Pathway |
| SIM1 | -3.615285884196 | 0.0282893565224409 | down | Pathway |
| SYNPO | -3.59949724112553 | 4.60415811305413E-06 | down | Pathway |
| MMRN1 | -3.59874918511679 | 0.00147697324715866 | down | Pathway |
| GABRE | -3.59839153167981 | 0.00481201807973802 | down | Pathway |
| TLCD2 | -3.59086761666984 | 0.00035108323967454 | down | Pathway |

| | | | | |
|---|---|---|---|---|
| GOLGA8M | -3.58845394971181 | 0.00188975830332752 | down | NA |
| GNG11 | -3.58207051070969 | 2.81821035777139E-06 | down | NA |
| CNR1 | -3.57258439497634 | 0.00233570228654904 | down | Pathway |
| SYNE3 | -3.5692208770669 | 0.0000383001339113691 | down | Pathway |
| SLIT3 | -3.56448467245205 | 0.000119309514671337 | down | Pathway |
| DENND2A | -3.56370142870472 | 3.46935980267288E-06 | down | NA |
| PID1 | -3.56053552742711 | 0.000205838892437834 | down | Pathway |
| ABCA6 | -3.55667688669961 | 0.000011339465151072 | down | Pathway |
| ABCB5 | -3.55433235640515 | 0.0188435994940584 | down | Pathway |
| GLDN | -3.55422856825764 | 0.000608080952459838 | down | Pathway |
| COL25A1 | -3.55305947937497 | 0.00644007706582593 | down | NA |
| SPTBN1 | -3.54968221482542 | 4.19222087788146E-06 | down | Pathway |
| BMP6 | -3.54493581011831 | 0.000395784895508871 | down | Pathway |
| RAB3C | -3.54253182503072 | 0.00696313971949208 | down | Pathway |
| KANK4 | -3.5409385285132 | 0.00145553954574202 | down | Pathway |
| ADAMTS5 | -3.53666296454508 | 0.0000231600052310895 | down | Pathway |
| MEDAG | -3.52065293024942 | 0.0001158000930272 | down | Pathway |
| EGFR | -3.51603321986057 | 0.0000776285145144357 | down | Pathway |
| SLITRK2 | -3.51431167350137 | 0.000613752329335866 | down | Pathway |
| MRAS | -3.51043297436587 | 0.000204039481344666 | down | Pathway |
| CLIC5 | -3.50650277180751 | 0.000911933852378139 | down | Pathway |
| THSD7B | -3.50644150804196 | 0.000248543237413034 | down | Pathway |
| RORB | -3.49831192125561 | 0.00680934672390898 | down | Pathway |
| THBS4 | -3.49610826999243 | 0.00458278986785771 | down | Pathway |

| | | | | |
|---|---|---|---|---|
| PPARGC1A | -3.49604806001425 | 0.00182534060977757 | down | Pathway |
| EHD2 | -3.49442231432448 | 0.0000160561685248542 | down | Pathway |
| KANK3 | -3.49307724904695 | 0.0000276380671060869 | down | Pathway |
| C1QTNF1 | -3.48954802734576 | 0.0000138892049247438 | down | Pathway |
| DCT | -3.48877176609222 | 0.000116452235987473 | down | Pathway |
| C7orf41 | -3.46002711728378 | 0.0000246359191821229 | down | NA |
| SV2B | -3.45982092338506 | 0.00135253883046239 | down | Pathway |
| RBPMS2 | -3.45956498724779 | 0.0000127212743298798 | down | Pathway |
| HOXA5 | -3.4582836667281 | 0.000467124973211234 | down | Pathway |
| TMEM88 | -3.45636731239783 | 7.75610919931282E-06 | down | Pathway |
| ITM2A | -3.45004719791818 | 0.0000276807225929861 | down | NA |
| MGLL | -3.44761532324938 | 0.000486714959662508 | down | Pathway |
| WASF3 | -3.43333945711301 | 0.000467178037277391 | down | Pathway |
| PKDCC | -3.42992369754716 | 0.000141818334379598 | down | Pathway |
| LEPR | -3.4295854795498 | 6.86538373931544E-06 | down | Pathway |
| NDNF | -3.42799904143851 | 0.00138620547389207 | down | Pathway |
| PTGFR | -3.42098100677971 | 0.000868075912579947 | down | Pathway |
| ELOVL3 | -3.41574758232844 | 0.00352129257719098 | down | Pathway |
| ADCYAP1R1 | -3.41289170387051 | 0.00887174517079537 | down | Pathway |
| CD209 | -3.40804605619145 | 0.00688492820022576 | down | Pathway |
| BPIFB2 | -3.40677274698719 | 0.00963174662568759 | down | NA |
| GYPC | -3.40139450447447 | 0.000250656412124445 | down | NA |
| MAOB | -3.40110246209627 | 0.000141818334379598 | down | Pathway |
| MCAM | -3.40037108980182 | 6.23165598484117E-06 | down | Pathway |

| | | | | |
|---|---|---|---|---|
| GPER1 | -3.39631730077417 | 0.000113657142831213 | down | Pathway |
| MYRIP | -3.39241961899401 | 0.00280636574446582 | down | Pathway |
| SSTR1 | -3.38636902546674 | 0.00281418192569626 | down | Pathway |
| LGI4 | -3.38606701718793 | 0.0000987598037500726 | down | Pathway |
| KIAA1045 | -3.38574565620428 | 0.000485274796211091 | down | NA |
| LPHN3 | -3.384021402635 | 0.000226269035045052 | down | NA |
| ASS1 | -3.37785573469757 | 0.00244230817713593 | down | Pathway |
| PREX2 | -3.37595713583181 | 0.0000501761030412632 | down | Pathway |
| CCL13 | -3.37071825644541 | 0.0178855844568774 | down | Pathway |
| SCN9A | -3.37040215792798 | 0.000374117349314525 | down | Pathway |
| C10orf90 | -3.36801542581684 | 0.00332097079768363 | down | Pathway |
| TTPA | -3.36774146458754 | 0.00866293633700327 | down | Pathway |
| MT1X | -3.35756631792791 | 0.000165237414605702 | down | Pathway |
| CRTAC1 | -3.35520666387272 | 0.0472725533412473 | down | Pathway |
| ESR2 | -3.35256171475675 | 0.000441084063429536 | down | Pathway |
| FMOD | -3.35095883316811 | 0.000223227171581174 | down | Pathway |
| LRRC2 | -3.34735599647031 | 0.000113657142831213 | down | NA |
| FAM150B | -3.34429481477358 | 0.0209400512349099 | down | NA |
| MFAP4 | -3.34228344466251 | 0.00115661684218583 | down | Pathway |
| LIFR | -3.33925284963003 | 0.0000233944443705919 | down | NA |
| AFF2 | -3.33911829215836 | 0.00338219740239093 | down | Pathway |
| EDNRB | -3.33003493632662 | 0.0000142621697148002 | down | Pathway |
| STX11 | -3.32809780468118 | 0.000151222201296785 | down | Pathway |
| SCN3A | -3.32362316446504 | 0.00100321475980003 | down | Pathway |

| | | | | |
|---|---|---|---|---|
| IGSF10 | -3.32297982853027 | 0.000345501284740521 | down | Pathway |
| FAM149A | -3.32084053134004 | 0.000131292341600427 | down | NA |
| FAM162B | -3.32070639137785 | 0.000248543237413034 | down | NA |
| DIO3 | -3.31823132905632 | 0.0113296757803969 | down | Pathway |
| ADM | -3.31328389720215 | 0.000118517326107118 | down | Pathway |
| PRPH | -3.3094818784071 | 0.00181969213656071 | down | NA |
| TGFBR2 | -3.30267924595493 | 0.0000114475091836265 | down | Pathway |
| GDF5 | -3.30176958731517 | 0.0014394023965887 | down | Pathway |
| MEST | -3.29709032503795 | 0.0000573557135571416 | down | Pathway |
| MGST1 | -3.29325392755476 | 0.00209453764137192 | down | Pathway |
| RGS6 | -3.29003281800119 | 0.00471885773183127 | down | Pathway |
| PNPLA2 | -3.28907813553096 | 6.86538373931544E-06 | down | Pathway |
| ABLIM3 | -3.28800542314957 | 0.000589275416381324 | down | Pathway |
| EBF2 | -3.28146582577916 | 0.0000705182148924997 | down | Pathway |
| PHYHIP | -3.27970407635738 | 0.0000165697766706899 | down | NA |
| CSMD1 | -3.27919997651096 | 0.0355721711712099 | down | Pathway |
| CADM3 | -3.27835797177992 | 0.000292548645182149 | down | Pathway |
| ACSL1 | -3.27682817541346 | 0.0000983664357695885 | down | Pathway |
| ANK2 | -3.27682268655122 | 0.000241403265626349 | down | Pathway |
| GPM6A | -3.27647402547181 | 0.0235103775207698 | down | Pathway |
| FAXDC2 | -3.27554805905379 | 0.000325264961088344 | down | Pathway |
| FAM110D | -3.27303209724073 | 0.0000302660001749592 | down | NA |
| TRIM61 | -3.27055685152505 | 0.000915919013007425 | down | NA |
| WSCD1 | -3.26519625581866 | 0.000236331886537201 | down | NA |

| | | | | |
|---|---|---|---|---|
| CCDC80 | -3.26271475919778 | 0.000248598917320596 | down | Pathway |
| ACO1 | -3.25868312763603 | 0.0000216013010133142 | down | Pathway |
| CYP26B1 | -3.25682777968583 | 0.00056566482513222 | down | Pathway |
| CXCR1 | -3.25589960724005 | 0.0032737444367043 | down | Pathway |
| CLVS1 | -3.25318026186152 | 0.000736715997381874 | down | NA |
| TBX15 | -3.24916438416385 | 0.000929013260278517 | down | Pathway |
| FGF7 | -3.24488159461145 | 0.000102523997037046 | down | Pathway |
| HOXB8 | -3.24342132985722 | 0.000900034428774405 | down | Pathway |
| PLP1 | -3.24229118994353 | 0.0389058721932742 | down | Pathway |
| RNF157 | -3.23775445426018 | 0.000079070782724783 | down | Pathway |
| ART4 | -3.23523764281573 | 0.00767670442226362 | down | Pathway |
| ALDOC | -3.2343597503791 | 0.00013536565337438 | down | Pathway |
| RNF150 | -3.22240820614812 | 0.0126888140897027 | down | NA |
| BAI3 | -3.22164570523051 | 0.00385388988673581 | down | NA |
| NIPSNAP3B | -3.22132478548041 | 0.000214581904479832 | down | NA |
| CYP3A5 | -3.220853939586 | 0.00280867451928625 | down | Pathway |
| PM20D1 | -3.22010823349126 | 0.0167291356033649 | down | Pathway |
| CCL21 | -3.21873801302757 | 0.0202295448469118 | down | Pathway |
| MAMDC2 | -3.21588629680419 | 0.00048557048018908 | down | NA |
| EFEMP1 | -3.21302285404775 | 0.000922833019169213 | down | Pathway |
| EPAS1 | -3.21068203304844 | 0.0000820462790595965 | down | Pathway |
| NOVA1 | -3.2064302600185 | 0.00752613011903416 | down | Pathway |
| AQP1 | -3.20168793657922 | 0.0000674325767586959 | down | Pathway |
| FNDC4 | -3.19839471371748 | 0.0000907487497202508 | down | Pathway |
| ACE2 | -3.19812143276693 | 0.00549872263106959 | down | Pathway |

| | | | | |
|---|---|---|---|---|
| PALM | -3.19755955545334 | 0.0000271212961336546 | down | Pathway |
| DOCK11 | -3.19702208367297 | 0.0000541162231374976 | down | Pathway |
| CACNA2D1 | -3.19698253253938 | 0.0000983664357695885 | down | Pathway |
| CRHR2 | -3.19050292552157 | 0.00846125009380684 | down | Pathway |
| SMOC1 | -3.18522117690626 | 0.00544629483212838 | down | Pathway |
| GIPC2 | -3.17600417156214 | 0.000119265125967958 | down | NA |
| GNG2 | -3.17558358397625 | 0.0000298697963818535 | down | Pathway |
| PLA2G16 | -3.17535928544061 | 0.000146406827745791 | down | NA |
| GALNT13 | -3.17341956275921 | 0.000700595594406309 | down | Pathway |
| ALX1 | -3.17211707451355 | 0.00902106186430693 | down | Pathway |
| LHFP | -3.16301924839933 | 0.0000110582399127156 | down | NA |
| LMOD1 | -3.16037908030158 | 0.0000695787681704741 | down | Pathway |
| C1orf115 | -3.15979025904395 | 6.55608097662842E-06 | down | NA |
| SHE | -3.15780698370336 | 0.0000126485088705199 | down | NA |
| PTRF | -3.15449542457307 | 0.0000165697766706899 | down | NA |
| DPP6 | -3.14904952110205 | 0.0216834725309011 | down | Pathway |
| RARRES2 | -3.1484071041035 | 0.000813858716747745 | down | Pathway |
| C2CD4B | -3.14752997297815 | 0.018819221318244 | down | Pathway |
| P2RY12 | -3.14486322301323 | 0.00017903656295909 | down | Pathway |
| CAT | -3.1441189146174 | 0.0000279906111157143 | down | Pathway |
| VGLL3 | -3.1427578294669 | 0.0000271212961336546 | down | NA |
| S1PR1 | -3.14049986331424 | 0.0000174267445510039 | down | Pathway |
| AIF1L | -3.13832523451778 | 0.000942497585998621 | down | Pathway |
| PTGIS | -3.13584278391706 | 0.00302199038600061 | down | Pathway |
| CA3 | -3.13309328206816 | 0.032682009523293 | down | Pathway |

| | | | | |
|---|---|---|---|---|
| PLIN5 | -3.13273994632366 | 0.0188435994940584 | down | Pathway |
| TEK | -3.13255535949579 | 0.0000222356336549405 | down | Pathway |
| GRB14 | -3.13209680954944 | 0.022660886497866 | down | Pathway |
| RGN | -3.1229125305068 | 0.0095518537332813 | down | Pathway |
| SOX6 | -3.12254074838663 | 0.0120190094198583 | down | Pathway |
| MAPK10 | -3.12063968967659 | 0.00051428028943494 | down | Pathway |
| KLHL31 | -3.11971654627897 | 0.00128877260305796 | down | Pathway |
| FAM13A | -3.11887778023604 | 0.000162681118293746 | down | Pathway |
| ZCCHC24 | -3.1183839927436 | 0.0000231600052310895 | down | NA |
| SYNM | -3.11802734581242 | 0.00161805377113719 | down | Pathway |
| ABCA10 | -3.11631522845126 | 0.0000525970188393347 | down | Pathway |
| FBLN2 | -3.10952818441719 | 0.00119633534347166 | down | Pathway |
| RYR3 | -3.10786226317265 | 0.0202714868055221 | down | Pathway |
| BANK1 | -3.10574695247517 | 0.000117854090787491 | down | Pathway |
| CYP19A1 | -3.09888831648864 | 0.0251428803736838 | down | Pathway |
| SIK2 | -3.09085697631705 | 0.0000319476602110509 | down | Pathway |
| PPAP2A | -3.09062394922178 | 7.81149829611418E-06 | down | NA |
| TSPAN7 | -3.08993648380612 | 0.00230802063145906 | down | NA |
| CXorf36 | -3.08425795625045 | 0.0000108596544569222 | down | NA |
| MYL3 | -3.07953033708248 | 0.0058038808891796 | down | Pathway |
| CABP1 | -3.05945850835388 | 0.000216635632680683 | down | Pathway |
| DEFB1 | -3.05573198223699 | 0.0231177358048044 | down | Pathway |
| NRG2 | -3.0533931724567 | 0.000473536753678606 | down | NA |
| CORO2B | -3.05246013638984 | 0.000269175251478561 | down | Pathway |
| FAT2 | -3.05115150709159 | 0.0222470908853414 | down | Pathway |

| | | | | |
|---|---|---|---|---|
| RETSAT | -3.05017527464941 | 0.0000182694821537976 | down | Pathway |
| STEAP4 | -3.04911552136889 | 0.00286476027223067 | down | Pathway |
| TACC1 | -3.04602836514345 | 0.0000309776681751308 | down | Pathway |
| CCDC36 | -3.0439638961412 | 0.000163991887276702 | down | NA |
| CDKN2C | -3.0372701571782 | 0.0000776285145144357 | down | Pathway |
| ECSCR | -3.03410194972266 | 0.0000116557440237734 | down | Pathway |
| KCNK3 | -3.02949375734383 | 0.0415187246244953 | down | Pathway |
| ABI3BP | -3.0225932865783 | 0.00205809357022652 | down | Pathway |
| SLC16A12 | -3.02109016488271 | 0.00233203390978014 | down | Pathway |
| RBMS3 | -3.02094824872171 | 0.0000532787255821952 | down | Pathway |
| KCNJ8 | -3.02066702402038 | 0.0000608423462801813 | down | Pathway |
| CETP | -3.01977125620677 | 0.00484530835517679 | down | Pathway |
| MGAT3 | -3.01915840412785 | 0.00345665780095536 | down | Pathway |
| MYOCD | -3.01652652210509 | 0.00348108678866668 | down | Pathway |
| ZNF728 | -3.01153253696136 | 0.0100641863927781 | down | NA |
| MAP7D3 | -3.00937911927696 | 0.000262144502419119 | down | Pathway |
| PNMA6C | -3.00864597333944 | 0.02141119530514 | down | NA |
| GPLD1 | -3.00709992881942 | 0.00418628908115978 | down | Pathway |
| TMEM246 | -3.00568705760756 | 0.000431332646885772 | down | NA |
| LDB2 | -3.00165239414993 | 0.0000111856259866863 | down | Pathway |
| ALS2CR11 | -2.99404147134348 | 0.011627420777486 | down | NA |
| TLN2 | -2.98807871352204 | 0.0000105415976727814 | down | Pathway |
| NEGR1 | -2.97906364419617 | 0.000508647596353905 | down | Pathway |
| CCM2L | -2.97679589041738 | 0.0000271212961336546 | down | Pathway |
| STS | -2.97638804775021 | 0.0000146139227501972 | down | Pathway |

| | | | | |
|---|---|---|---|---|
| KL | -2.97361109266607 | 0.00320652401274349 | down | Pathway |
| LOXL4 | -2.96853558489894 | 0.000861267488020354 | down | Pathway |
| CNKSR2 | -2.96806099999378 | 0.000940606639141062 | down | Pathway |
| MT1A | -2.96574301841181 | 0.0299020955501491 | down | Pathway |
| PNMAL2 | -2.96290736189915 | 0.000877977769201698 | down | NA |
| TMTC1 | -2.96216186861126 | 0.000512606771232984 | down | NA |
| ANXA1 | -2.96147830241599 | 0.000508647596353905 | down | Pathway |
| LHX6 | -2.95966190829793 | 0.000190284078984827 | down | Pathway |
| USP44 | -2.95820176324187 | 0.00321714455979539 | down | Pathway |
| SYT15 | -2.9567327287024 | 0.012311606703728 | down | Pathway |
| EMCN | -2.95467130479812 | 0.0000279906111157143 | down | NA |
| ROBO4 | -2.95438490260642 | 0.0000279906111157143 | down | Pathway |
| WSCD2 | -2.95294866509061 | 0.0000706988760858014 | down | NA |
| SH3BGRL2 | -2.95199645455275 | 0.0000231600052310895 | down | NA |
| COX4I2 | -2.9447757911505 | 0.00042386930184934 | down | NA |
| GSC | -2.94339198228953 | 0.000251899046000088 | down | Pathway |
| GFRA2 | -2.94147641209315 | 0.0159750748527446 | down | NA |
| RNASE1 | -2.94068896780737 | 0.00265938938578874 | down | NA |
| HCAR3 | -2.93887786240111 | 0.00117672943473209 | down | NA |
| EGFL7 | -2.93864955029254 | 0.0000159440884555377 | down | Pathway |
| MYEOV | -2.93659854350643 | 0.0121694911558815 | down | NA |
| C2CD4C | -2.93508305935116 | 0.000431332646885772 | down | NA |
| IGF2 | -2.93380638618972 | 0.00246376790917567 | down | Pathway |
| PCBP3 | -2.93064271768449 | 0.00508167804998552 | down | NA |

| | | | | |
|---|---|---|---|---|
| JAM2 | -2.92950876708732 | 0.0000673604236703592 | down | Pathway |
| PC | -2.92861507630627 | 0.000215840636591643 | down | Pathway |
| CNGA1 | -2.92643083117656 | 0.00922407955483978 | down | NA |
| SBK3 | -2.92462741051851 | 0.00056566482513222 | down | NA |
| PDE8B | -2.92139309675684 | 0.0180524208154652 | down | Pathway |
| PCSK5 | -2.92130823430796 | 0.000015696620047865 | down | Pathway |
| PTPRB | -2.91949672347883 | 0.0000501761030412632 | down | Pathway |
| KCNE1L | -2.91889142896902 | 0.00490926927885822 | down | NA |
| CD34 | -2.91663601983918 | 0.0000127084518999542 | down | Pathway |
| SPATA22 | -2.91306157317499 | 0.0351453590456273 | down | Pathway |
| SOX5 | -2.9121153469593 | 0.0188259132023721 | down | Pathway |
| OLFM2 | -2.91086893679128 | 0.00840275837207068 | down | Pathway |
| PPP2R1B | -2.90813136994828 | 4.09983852314949E-06 | down | NA |
| SCN7A | -2.90533089788951 | 0.00520369582389385 | down | Pathway |
| CDH5 | -2.9043119193405 | 0.0000276807225929861 | down | Pathway |
| RSPO3 | -2.89724963531218 | 0.00909669683764579 | down | Pathway |
| CLSTN2 | -2.8964698110971 | 0.0348331726387353 | down | Pathway |
| F13A1 | -2.8895883997282 | 0.00270351074517001 | down | Pathway |
| SEMA3D | -2.88333470452205 | 0.00480292293458476 | down | Pathway |
| GIMAP7 | -2.88287854789225 | 0.000644903205574493 | down | NA |
| FAT4 | -2.8806864326546 | 0.0000518100577819199 | down | Pathway |
| FLRT2 | -2.88062089988889 | 0.00139185330339306 | down | Pathway |
| FGF14 | -2.88051153527911 | 0.00229984948079451 | down | Pathway |
| KCTD16 | -2.87586186937316 | 0.00454687995958166 | down | Pathway |
| CTNNA3 | -2.8732816265294 | 0.000350006354206234 | down | Pathway |
| KANK1 | -2.87072176871455 | 0.000212119686593945 | down | Pathway |

| | | | | |
|---|---|---|---|---|
| APCDD1 | -2.86587939077875 | 0.00169512196123347 | down | Pathway |
| PCDH18 | -2.86374269990646 | 0.000153027504055082 | down | Pathway |
| SPARCL1 | -2.86073747822022 | 0.0000412403604997038 | down | Pathway |
| SLC35G2 | -2.86048319738658 | 0.000197745261188871 | down | NA |
| IQSEC3 | -2.85971199089039 | 0.00905557304949794 | down | Pathway |
| AGTR1 | -2.85925819123501 | 0.0446401695534864 | down | Pathway |
| FOLR2 | -2.85702266177438 | 0.0207989872550089 | down | Pathway |
| ICAM2 | -2.85292399944758 | 0.0000776285145144357 | down | NA |
| SOX7 | -2.85279566486268 | 0.000116452235987473 | down | Pathway |
| ARHGEF15 | -2.85124240274834 | 0.0000970166031874953 | down | Pathway |
| VWF | -2.84981612172864 | 0.000188487593620814 | down | Pathway |
| TMEM220 | -2.84674215663736 | 0.000649370895645139 | down | NA |
| DCN | -2.84271563645101 | 0.0011272928520189 | down | Pathway |
| HOXA3 | -2.84258464832799 | 0.000663244467285107 | down | Pathway |
| ADRB2 | -2.84236433856635 | 0.001549696519053 | down | Pathway |
| LDHD | -2.84156274820099 | 0.00144348008265578 | down | Pathway |
| HYAL1 | -2.83994889795468 | 7.89799979427237E-06 | down | Pathway |
| ANGPTL2 | -2.83863474870753 | 0.0000608423462801813 | down | NA |
| USHBP1 | -2.83599201745087 | 0.0000256658752137784 | down | NA |
| GIMAP6 | -2.83394526666309 | 0.000139807852494195 | down | NA |
| FAM178B | -2.82930845962009 | 0.000869120605228221 | down | NA |
| SERPINF1 | -2.82806675146491 | 0.000738068453006212 | down | Pathway |
| HSPA12B | -2.82452270709192 | 0.000175174974499993 | down | NA |

| | | | | |
|---|---|---|---|---|
| AFAP1L1 | -2.82138205512249 | 0.000076246527908556 | down | NA |
| RASIP1 | -2.81588967061164 | 0.0000279906111157143 | down | Pathway |
| RASL11A | -2.81134613233941 | 0.0000119501501165282 | down | NA |
| ANKRD29 | -2.81058859522513 | 0.000915276692190789 | down | NA |
| MYCT1 | -2.80842201673919 | 7.81149829611418E-06 | down | NA |
| ADRA2C | -2.80507526640384 | 0.0490175716730538 | down | Pathway |
| LILRB5 | -2.80429600230025 | 0.0280582289809411 | down | NA |
| IGFN1 | -2.8005141473809 | 0.0205280709928409 | down | Pathway |
| CLEC14A | -2.79975060927304 | 0.000014035296417122 | down | Pathway |
| OSR1 | -2.79899881041283 | 0.0188240857257247 | down | Pathway |
| KIF26A | -2.79693553357693 | 0.0000279906111157143 | down | Pathway |
| FAM13C | -2.79650539543417 | 0.000431772835303863 | down | NA |
| GPR116 | -2.79259380102607 | 0.0000197273938950305 | down | NA |
| EGFLAM | -2.79182521897109 | 0.000158964635373281 | down | Pathway |
| FERMT2 | -2.78479368516653 | 8.31905963221189E-06 | down | Pathway |
| MRGPRF | -2.78376174663366 | 0.00249446761649592 | down | NA |
| CAPN11 | -2.7827415082666 | 0.00241236489864774 | down | NA |
| ASXL3 | -2.78213404053242 | 0.0106162702848304 | down | Pathway |
| MMRN2 | -2.7804565164217 | 0.0000804534278010506 | down | Pathway |
| PPP1R16B | -2.77884167767368 | 0.000971678499600897 | down | Pathway |
| ARC | -2.77609853797032 | 0.0226674284559493 | down | Pathway |
| HSPA12A | -2.77239174655165 | 0.0000317201556733218 | down | NA |
| NDN | -2.76815363001688 | 0.000759925664784329 | down | Pathway |
| KLHL4 | -2.76805918342618 | 0.00535468215420713 | down | NA |
| PARK2 | -2.76695661257463 | 0.0000121719061601948 | down | NA |

| | | | | |
|---|---|---|---|---|
| TDRD10 | -2.76570189791637 | 0.00326614339243186 | down | Pathway |
| LAMA4 | -2.76168288609567 | 0.0000135579150977941 | down | Pathway |
| PFKFB3 | -2.75912851645476 | 0.00445620677212947 | down | Pathway |
| SPATA9 | -2.75859133613808 | 0.00452864868731021 | down | NA |
| PPAPDC3 | -2.75592713248859 | 0.000601594805013111 | down | NA |
| TAL1 | -2.75163448334989 | 0.0000154779552885175 | down | Pathway |
| CD248 | -2.74990154804813 | 0.00145915024249602 | down | NA |
| DLC1 | -2.73842536294813 | 0.0000078671363756136 | down | Pathway |
| HOXA4 | -2.73530309216713 | 0.000562760464026854 | down | Pathway |
| ACKR3 | -2.73436898586493 | 0.000704610227076125 | down | Pathway |
| COL6A6 | -2.73362172895113 | 0.0244636475164922 | down | Pathway |
| ALPK3 | -2.73163640246981 | 0.00145915024249602 | down | Pathway |
| DDX43 | -2.72730639328562 | 0.00832876345793972 | down | NA |
| MET | -2.72331879103082 | 0.00750187247990144 | down | Pathway |
| FGF16 | -2.72223024078183 | 0.0253422132360867 | down | Pathway |
| MMP27 | -2.72040305806932 | 0.0190860427790771 | down | Pathway |
| CKB | -2.71610030490195 | 0.0152623935077707 | down | Pathway |
| FBXO27 | -2.71496027234879 | 0.0108365691865681 | down | NA |
| FCN3 | -2.71351494056913 | 0.000644903205574493 | down | Pathway |
| CNTNAP3 | -2.71139486532355 | 0.000844752341507983 | down | NA |
| ADRA2A | -2.70575002161044 | 0.00482686394488731 | down | Pathway |
| NGFR | -2.69674677324854 | 0.00307631715715913 | down | Pathway |
| HOXA2 | -2.69333834899702 | 0.00325856338812637 | down | Pathway |

| | | | | |
|---|---|---|---|---|
| PLSCR4 | -2.69214363422623 | 8.13603719666152E-06 | down | Pathway |
| PTGDS | -2.69073722228498 | 0.0164672022868194 | down | Pathway |
| TRPM6 | -2.68444828840117 | 0.00275966451272313 | down | Pathway |
| ST6GALNAC3 | -2.68434004773936 | 0.0000494797234835793 | down | Pathway |
| TLL1 | -2.68404840495517 | 0.00145102726031926 | down | Pathway |
| TNNC2 | -2.67821458840019 | 9.40865569656448E-06 | down | Pathway |
| CDC42EP2 | -2.67150529289564 | 2.81821035777139E-06 | down | Pathway |
| FFAR4 | -2.66880068173883 | 0.00163768504147142 | down | Pathway |
| MSRB3 | -2.66851798580099 | 0.0000300655359075657 | down | NA |
| KCNA1 | -2.66843292725997 | 0.0487983501383243 | down | Pathway |
| CNRIP1 | -2.66745937300511 | 0.000292548645182149 | down | NA |
| C4orf19 | -2.66484161463065 | 0.0330371322498722 | down | NA |
| MYOM2 | -2.66280593962407 | 0.00133901541050801 | down | Pathway |
| VSTM4 | -2.65933793212794 | 0.000228503668915116 | down | Pathway |
| SSPN | -2.65855337681769 | 0.00027255436347788 | down | Pathway |
| EMP1 | -2.65814733216708 | 0.0000272524090656503 | down | Pathway |
| POMC | -2.65759653459094 | 0.00769842241759571 | down | Pathway |
| EPDR1 | -2.65569162182974 | 0.0030941928897933 | down | Pathway |
| ECHDC3 | -2.6530440055515 | 0.0000800308091501004 | down | Pathway |
| FZD9 | -2.65110135447346 | 0.004124635248681 | down | Pathway |
| ACADS | -2.65060963323293 | 6.94705115187937E-06 | down | Pathway |
| CYP4F12 | -2.65001218368114 | 0.0108793314762571 | down | Pathway |
| FBLN5 | -2.64955653126288 | 0.00451789014201225 | down | Pathway |
| RILP | -2.64758207191416 | 0.000339081478581284 | down | Pathway |

| | | | | |
|---|---|---|---|---|
| AIFM2 | -2.64726768738435 | 0.0000182694821537976 | down | Pathway |
| PODN | -2.64691304020582 | 0.00219769730433524 | down | Pathway |
| TSHZ2 | -2.64653179405992 | 0.000083142834261072 | down | NA |
| ADAM33 | -2.64288539518565 | 0.0225282775518651 | down | NA |
| MLTK | -2.64184667164346 | 0.000128228570138333 | down | NA |
| ANTXR2 | -2.64106622210983 | 0.000035273576192804 | down | NA |
| NYAP1 | -2.64012770599837 | 0.00334856571634844 | down | Pathway |
| CCDC141 | -2.63869092580596 | 0.000906295089778017 | down | Pathway |
| PGM1 | -2.63760342746625 | 0.000120581223029634 | down | Pathway |
| SGCA | -2.63607459824016 | 0.000529387418480805 | down | Pathway |
| CPM | -2.63549478216561 | 0.00794135678819648 | down | NA |
| DMTN | -2.63485034249489 | 0.000405010877871049 | down | Pathway |
| FAM212B | -2.63113106379777 | 0.0000608423462801813 | down | NA |
| SPIN2A | -2.63096440645751 | 0.0000523496120669737 | down | NA |
| CSPG4 | -2.63085342084907 | 0.000303936242259401 | down | Pathway |
| EPHB6 | -2.62598513778448 | 0.00226918892700237 | down | Pathway |
| MPZ | -2.62440696351078 | 0.00247461331178703 | down | Pathway |
| KCNJ12 | -2.62370971414237 | 0.00307890462092542 | down | Pathway |
| KCNA6 | -2.62291837784999 | 0.00757408497251189 | down | Pathway |
| NFIB | -2.62041845173125 | 0.000510167641940358 | down | Pathway |
| ZBED6 | -2.61657566021846 | 0.0232111884782677 | down | Pathway |
| ZNF385D | -2.61338523859428 | 0.000472155211243409 | down | NA |
| GPT | -2.61195663821609 | 0.027525311654449 | down | Pathway |

| | | | | |
|---|---|---|---|---|
| WDR86 | -2.60891267991497 | 0.00376035975460225 | down | NA |
| TNFRSF8 | -2.60882200247278 | 0.00929335649705296 | down | Pathway |
| CRIM1 | -2.60867839082788 | 0.000507335957373077 | down | Pathway |
| GBE1 | -2.60540159876039 | 0.000162681118293746 | down | Pathway |
| MESP1 | -2.60216497660424 | 0.0151300399591247 | down | Pathway |
| P2RY14 | -2.60017370411147 | 0.000540113325507902 | down | NA |
| LIMS2 | -2.59898239411857 | 0.00023087769011815 | down | Pathway |
| ARHGEF4 | -2.59764202096039 | 0.00387307674648601 | down | Pathway |
| LOXHD1 | -2.59660556750998 | 0.00113798502039897 | down | Pathway |
| IL17D | -2.585692736402 | 0.00187345393893482 | down | NA |
| ERG | -2.58538014693109 | 6.64655070757393E-06 | down | NA |
| ADAMTSL3 | -2.58279339053288 | 0.000499903596536925 | down | Pathway |
| ACOT2 | -2.58217515789505 | 0.000667550086311639 | down | Pathway |
| ELTD1 | -2.58110609287168 | 0.0000116557440237734 | down | NA |
| ZFPM2 | -2.58059920399693 | 0.000273291651608255 | down | Pathway |
| RHCG | -2.58034084717333 | 0.0443365910316329 | down | Pathway |
| TINAGL1 | -2.57668869913375 | 0.0000603309823850517 | down | NA |
| CXCR2 | -2.57640166854751 | 0.000349273418373698 | down | Pathway |
| SCN3B | -2.57421825077591 | 0.00281418192569626 | down | Pathway |
| MARK1 | -2.56673833137386 | 0.00258052237930075 | down | Pathway |
| PHKG1 | -2.56596077574796 | 0.000408291543525218 | down | Pathway |
| ITSN1 | -2.56551025551206 | 4.19179441432916E-06 | down | Pathway |
| ME1 | -2.5626691791733 | 0.00185530742960061 | down | NA |
| SELP | -2.56243923932324 | 0.0103768441045141 | down | Pathway |

| | | | | |
|---|---|---|---|---|
| RHOXF1 | -2.56037134492675 | 0.000609602651536263 | down | Pathway |
| CDH23 | -2.56035259622217 | 0.013147532255673 | down | Pathway |
| LRRTM2 | -2.55578528921217 | 0.000083142834261072 | down | Pathway |
| LARP6 | -2.54975142181683 | 0.000485274796211091 | down | Pathway |
| AMOTL2 | -2.54593383231826 | 0.0000216812107614539 | down | Pathway |
| OR51E1 | -2.54492840927036 | 0.00325856338812637 | down | NA |
| RGMA | -2.54367361456321 | 0.00852337071720228 | down | Pathway |
| RRAS | -2.54276105366424 | 0.0000504491271049433 | down | Pathway |
| ZNF208 | -2.5396428012135 | 0.0100092446603312 | down | NA |
| C2CD2 | -2.5372990602052 | 0.000125494722169181 | down | NA |
| ST6GALNAC1 | -2.53614507538278 | 0.00538642899109318 | down | Pathway |
| RHOBTB3 | -2.534159713907 | 0.0021078631042849 | down | Pathway |
| RAMP3 | -2.52862810288995 | 0.000308899455123123 | down | Pathway |
| RIMBP2 | -2.52847946279442 | 0.0403359145812765 | down | NA |
| SH3D19 | -2.52746933352268 | 0.0000249486515527268 | down | Pathway |
| RAMP2 | -2.52659064793258 | 0.000153733090164043 | down | Pathway |
| GJA4 | -2.52479527896466 | 0.0000351034889861726 | down | Pathway |
| PVRL3 | -2.52414994429935 | 0.0021441037747987 | down | NA |
| PEBP4 | -2.5228878250044 | 0.00331076048162542 | down | NA |
| PEAR1 | -2.52209561522268 | 0.0000601544313076721 | down | Pathway |
| SPRY2 | -2.52077918716469 | 9.40865569656448E-06 | down | Pathway |
| MOCS1 | -2.52030229148641 | 0.00121028633221942 | down | NA |
| COCH | -2.51888344105173 | 0.011322249627765 | down | Pathway |
| CYYR1 | -2.51689945141495 | 0.0000373578845664019 | down | NA |
| SEPP1 | -2.51620804773035 | 0.00507661167408378 | down | NA |
| ZNF366 | -2.5140068793793 | 0.000292548645182149 | down | Pathway |

| | | | | |
|---|---|---|---|---|
| SYNPO2 | -2.51342333019811 | 0.00339097090104914 | down | Pathway |
| COPZ2 | -2.51272345826851 | 0.00100549513202968 | down | NA |
| RUNX1T1 | -2.50845970123274 | 0.000825138483672569 | down | Pathway |
| MYL9 | -2.499711370785 | 0.000433136280562955 | down | Pathway |
| PTX3 | -2.49751371590556 | 0.0366901158454262 | down | Pathway |
| LRP1 | -2.49658237577421 | 0.000249448202162025 | down | Pathway |
| KCNT2 | -2.48894750845244 | 0.00694006308467102 | down | Pathway |
| LCN10 | -2.48891715310964 | 0.00562991818020628 | down | NA |
| CCDC39 | -2.48353628265944 | 0.0109120167142285 | down | NA |
| C1QTNF2 | -2.47469316506232 | 0.00133693059646152 | down | Pathway |
| C11orf53 | -2.4725294127 | 0.0221775227136671 | down | NA |
| NR2F1 | -2.46750930611749 | 0.000441084063429536 | down | Pathway |
| LAMA2 | -2.46724627118946 | 0.00091172111452844 | down | Pathway |
| METTL7A | -2.46582652065669 | 0.000121388009059353 | down | NA |
| CTTNBP2 | -2.46376819325165 | 0.00626770841408716 | down | Pathway |
| PROS1 | -2.45891290075711 | 0.000214009802442619 | down | Pathway |
| SCN11A | -2.45493010303588 | 0.00170874121676437 | down | Pathway |
| TMIE | -2.45478470494593 | 0.00181673268362257 | down | Pathway |
| PPAP2B | -2.45294372958539 | 0.000254485834340982 | down | NA |
| MYH11 | -2.45228808320747 | 0.0199033238851549 | down | Pathway |
| CELF2 | -2.45148256081187 | 0.000688734676719381 | down | Pathway |
| ACSS3 | -2.44916875094374 | 0.00775281933385788 | down | NA |
| GPR124 | -2.44906368648212 | 0.000172264305958254 | down | NA |
| GYPE | -2.44768146049906 | 0.00770119757534949 | down | NA |

| | | | | |
|---|---|---|---|---|
| ANO4 | -2.44676338010463 | 0.00294454909661334 | down | Pathway |
| TEF | -2.44615356076068 | 0.0000297807856142578 | down | Pathway |
| HRC | -2.44594050580498 | 0.00177366881919331 | down | Pathway |
| KLHL29 | -2.44294633149558 | 0.00238262710984893 | down | NA |
| CRLF1 | -2.44268780116646 | 0.0121956352609844 | down | Pathway |
| ADAMTS9 | -2.4380417743832 | 0.0000525970188393347 | down | Pathway |
| RASSF9 | -2.43707407514703 | 0.00438413574157416 | down | NA |
| SASH1 | -2.43697768753088 | 0.000787830615815324 | down | Pathway |
| CDKN2B | -2.43404448046895 | 0.0000279906111157143 | down | Pathway |
| TMEM140 | -2.43266702532725 | 0.000468772268881032 | down | NA |
| PRRG3 | -2.43041251099148 | 0.0374107279167477 | down | NA |
| SYPL2 | -2.43001902660383 | 0.000176117704657276 | down | Pathway |
| FMN2 | -2.42992372426147 | 0.0212930972438783 | down | Pathway |
| GIMAP1 | -2.42966700883336 | 0.00201503545441972 | down | NA |
| PLEKHG6 | -2.42677724593942 | 0.000690022905459024 | down | Pathway |
| REEP2 | -2.42647583739228 | 0.0011492299653367 | down | Pathway |
| ZP1 | -2.4263447230585 | 0.0046103242323531 | down | Pathway |
| CLEC2L | -2.42318078207612 | 0.00604092034650895 | down | NA |
| DHRS3 | -2.42234159704129 | 0.000239200260565704 | down | Pathway |
| NOTCH4 | -2.41811964833808 | 0.000248543237413034 | down | Pathway |
| GIMAP8 | -2.41777986297669 | 0.000911933852378139 | down | NA |
| OXT | -2.41701431006424 | 0.0108972102649181 | down | Pathway |
| PLN | -2.41554782680986 | 0.00108929186953253 | down | Pathway |
| PECR | -2.41532166171923 | 0.000667550086311639 | down | Pathway |

| | | | | |
|---|---|---|---|---|
| MFAP5 | -2.41339894401977 | 0.00326602496705623 | down | Pathway |
| TM4SF18 | -2.41174164539375 | 0.0000136429042467373 | down | NA |
| ESAM | -2.4106558070303 | 0.0000348721026430314 | down | Pathway |
| MSRA | -2.41011108616822 | 0.000145928809403184 | down | Pathway |
| SGCE | -2.40877107632688 | 0.000700503270072223 | down | Pathway |
| BIN1 | -2.40858131567007 | 0.00019746681291275 | down | Pathway |
| SOBP | -2.40593509572562 | 0.0043034161229907 | down | Pathway |
| KLF4 | -2.40463494171611 | 0.00693139176967699 | down | Pathway |
| FAM124B | -2.40299467675363 | 0.000993835869103217 | down | NA |
| SYN3 | -2.40127938698094 | 0.000259201308557463 | down | Pathway |
| C10orf128 | -2.40073646014368 | 0.00196351350704928 | down | NA |
| NDRG2 | -2.39962535270586 | 0.000644903205574493 | down | Pathway |
| TESC | -2.39620847693588 | 0.00312808545879524 | down | Pathway |
| TMEM47 | -2.39558472242743 | 0.000516062519049087 | down | NA |
| ELN | -2.39418084859561 | 0.00788924216501734 | down | Pathway |
| TTN | -2.39212301641035 | 0.00605886133650656 | down | Pathway |
| HOXB7 | -2.39048539932819 | 0.000230684280569815 | down | Pathway |
| FABP5 | -2.38750298787268 | 0.0052221558926021 | down | Pathway |
| TENM3 | -2.38650357112582 | 0.00621178910355922 | down | Pathway |
| KIAA1614 | -2.38142209502988 | 0.000092702025597482 | down | Pathway |
| SMAD9 | -2.38104977438201 | 0.0000182694821537976 | down | Pathway |
| ANKRD35 | -2.38039921539123 | 0.00267947386790323 | down | NA |
| PDE7B | -2.37919066672054 | 0.00190000587193525 | down | Pathway |
| FADS3 | -2.37428622754384 | 0.0000136527038725634 | down | Pathway |

| | | | | |
|---|---|---|---|---|
| FBXO17 | -2.37220728651492 | 0.0220343578088744 | down | NA |
| PLIN2 | -2.37004971058131 | 0.00138550130500484 | down | Pathway |
| RASGRF2 | -2.37003044399895 | 0.000310725913459433 | down | Pathway |
| SEMA3A | -2.36758328909398 | 0.011723248920309 | down | Pathway |
| PLA2G4A | -2.3631351204978 | 0.00209366569254892 | down | Pathway |
| SLC9A9 | -2.36095049781371 | 0.00117672943473209 | down | Pathway |
| TENC1 | -2.35837741716803 | 0.000164493266543036 | down | NA |
| C10orf54 | -2.35699721235557 | 0.0013246982820412 | down | NA |
| FGD5 | -2.35563781241528 | 0.0000244970606182201 | down | Pathway |
| FEZ1 | -2.35358280786176 | 0.00105890810473494 | down | Pathway |
| CRYBG3 | -2.35335217578319 | 0.000215607438062973 | down | NA |
| EPS8 | -2.35264132318553 | 0.0000506791357951246 | down | Pathway |
| ALDH4A1 | -2.35199752979498 | 0.000215842509418116 | down | Pathway |
| TRIM50 | -2.35084384529796 | 0.0177486959427138 | down | NA |
| SH2D1B | -2.34778498050455 | 0.020345024415386 | down | NA |
| PLA2G5 | -2.34624407639458 | 0.0021048761859491 | down | Pathway |
| PHGDH | -2.34329041340047 | 0.00887711457661303 | down | Pathway |
| CFH | -2.342947057539 | 0.00159571332335374 | down | NA |
| CYGB | -2.33472200433035 | 0.000736715997381874 | down | Pathway |
| SGK2 | -2.33348587392094 | 0.00372566789800948 | down | Pathway |
| ZEB2 | -2.3325594630487 | 0.000628853343204391 | down | Pathway |
| HSDL2 | -2.33225760334512 | 0.000207433504676273 | down | NA |
| MYO1H | -2.33089369051936 | 0.0142591766331992 | down | Pathway |
| COBLL1 | -2.33058561401423 | 0.0000292644696970919 | down | NA |
| ADRA1B | -2.32898571788729 | 0.00862350011647218 | down | Pathway |

| | | | | |
|---|---|---|---|---|
| ITGB1BP1 | -2.32647798796142 | 9.35003131715643E-06 | down | Pathway |
| TUBB6 | -2.32519235682511 | 0.0000845369655144294 | down | NA |
| CTNNAL1 | -2.3181357682328 | 0.00023393419048509 | down | Pathway |
| CYB5R3 | -2.31527927466101 | 9.02194580668829E-06 | down | Pathway |
| CXCL12 | -2.31466939814107 | 0.00211137401172175 | down | Pathway |
| TFPI | -2.30344140572727 | 0.0000494797234835793 | down | Pathway |
| PAMR1 | -2.30233901929891 | 0.0228097906880952 | down | NA |
| PRDM16 | -2.29850619560963 | 0.0051534027900591 | down | Pathway |
| ACSS2 | -2.29754619285559 | 0.0000216825353897115 | down | Pathway |
| CEP112 | -2.29581624005838 | 0.000810110470696398 | down | Pathway |
| PAPPA2 | -2.2951129492516 | 0.0372795416383165 | down | Pathway |
| LDHB | -2.29438024825291 | 0.000545535222688912 | down | NA |
| DHH | -2.29154074858214 | 0.0188259132023721 | down | Pathway |
| KCNA5 | -2.28942752562313 | 0.00786067140452324 | down | Pathway |
| CPS1 | -2.28925318235319 | 0.00520369582389385 | down | Pathway |
| HAND2 | -2.28510049266033 | 0.0090083680338739 | down | Pathway |
| FOXN3 | -2.28509159645605 | 0.0000302660001749592 | down | Pathway |
| GIMAP5 | -2.28448548524175 | 0.00491352659349936 | down | NA |
| SHANK3 | -2.27747390167699 | 0.000205446686047126 | down | Pathway |
| AOC2 | -2.27649073559242 | 0.000170060550641866 | down | Pathway |
| RUNDC3B | -2.27491874605935 | 0.0041798401796802 | down | NA |
| ANO6 | -2.27410479368568 | 0.000220589758364881 | down | Pathway |
| RAB6B | -2.27260216536147 | 0.00108922007814007 | down | NA |
| ZNF423 | -2.27171494233054 | 0.00558235305166484 | down | Pathway |

| | | | | |
|---|---|---|---|---|
| KANK2 | -2.26789295541132 | 9.29322477234728E-06 | down | Pathway |
| PDGFRL | -2.26368155167737 | 0.00773311364441068 | down | NA |
| VIM | -2.26278103315448 | 0.00048207700127577 | down | Pathway |
| TWIST1 | -2.26018914511073 | 0.0252744632550311 | down | Pathway |
| ANXA3 | -2.25968353623889 | 0.030288943491289 | down | Pathway |
| FGF1 | -2.25564043498968 | 0.000450824470498984 | down | Pathway |
| ANKRD33B | -2.25532437512456 | 0.00269583076825462 | down | NA |
| PEMT | -2.25172297816225 | 0.0000246359191821229 | down | Pathway |
| ASPH | -2.24954160588975 | 0.0024500709589003 | down | Pathway |
| THSD7A | -2.24705433141967 | 0.000391187746166089 | down | Pathway |
| DLGAP2 | -2.24657367726438 | 0.0318840051621111 | down | Pathway |
| ID1 | -2.2420939418669 | 0.00255359974113374 | down | Pathway |
| CTIF | -2.23867011804749 | 0.0000983664357695885 | down | NA |
| TUBB2B | -2.23809226838988 | 0.0129742483772123 | down | Pathway |
| CLDND2 | -2.2379076738119 | 0.0044236087945341 | down | NA |
| VEGFB | -2.23370081501659 | 0.0000776285145144357 | down | Pathway |
| AC011242.6 | -2.22916325904979 | 0.0123699031226192 | down | NA |
| FMO3 | -2.22822131749059 | 0.00307684048047913 | down | NA |
| SPTA1 | -2.2276675898561 | 0.0334356491927514 | down | Pathway |
| EPHA3 | -2.22299238462671 | 0.0118689936170433 | down | Pathway |
| HFM1 | -2.22131140602015 | 0.0110054393686342 | down | Pathway |
| ABHD6 | -2.22069706594807 | 0.00153501584143883 | down | Pathway |
| KLF9 | -2.21782198739451 | 0.000168232320553012 | down | Pathway |
| C2orf73 | -2.21691843414404 | 0.0199439040369627 | down | NA |
| SLC24A3 | -2.21573584504441 | 0.00951653821392023 | down | Pathway |

| | | | | |
|---|---|---|---|---|
| DAAM2 | -2.21569243326377 | 0.000488192635042567 | down | Pathway |
| F8 | -2.21184555841888 | 0.000088167205817432 | down | Pathway |
| ADAMTSL4 | -2.21182235873485 | 0.00592804137738116 | down | Pathway |
| TMEM132B | -2.21153312541306 | 0.0480684797169861 | down | NA |
| SH3RF2 | -2.21139840047814 | 0.00780913914480076 | down | Pathway |
| CAMK1 | -2.20969124882013 | 0.0000650555176656801 | down | Pathway |
| PRCD | -2.20736475542816 | 0.0110559690241855 | down | NA |
| TIE1 | -2.20730720946347 | 0.0000355328007087224 | down | Pathway |
| MICU3 | -2.20489096631708 | 0.0000834195459999636 | down | Pathway |
| CLCN4 | -2.20466179450951 | 0.0169946930157186 | down | Pathway |
| TCF7L1 | -2.20016274866925 | 0.0109961846083061 | down | Pathway |
| NGF | -2.19994092881192 | 0.00257832651968228 | down | Pathway |
| PDE1B | -2.19710635556537 | 0.00276579133623911 | down | Pathway |
| CDC14B | -2.19704425917901 | 0.0000800308091501004 | down | Pathway |
| PTPRS | -2.19494779603142 | 0.00182534060977757 | down | Pathway |
| PZP | -2.19218334449558 | 0.00141093543080837 | down | Pathway |
| GPR34 | -2.19014704956364 | 0.00251349528441885 | down | NA |
| IL1RL1 | -2.18979877315635 | 0.044776884678522 | down | Pathway |
| PAPSS2 | -2.1879698751783 | 0.000416222788834241 | down | Pathway |
| LY6K | -2.18660242189042 | 0.0444413524124325 | down | NA |
| SLC14A1 | -2.179513384637 | 0.00577702406883004 | down | NA |
| LIMCH1 | -2.17638392433019 | 0.00683453116517172 | down | Pathway |
| PLEKHG5 | -2.17614844578997 | 0.0000820462790595965 | down | Pathway |
| GPR97 | -2.17459565170562 | 0.0238393234010804 | down | NA |

| | | | | |
|---|---|---|---|---|
| SNCAIP | -2.17439684516547 | 0.0131697712379213 | down | Pathway |
| ELMOD3 | -2.16888939512476 | 0.000123180519229478 | down | NA |
| EPB41L4B | -2.1678172480109 | 0.0208693549167121 | down | Pathway |
| TSKS | -2.16723691837424 | 0.0218953657214285 | down | NA |
| STK32A | -2.16628705609853 | 0.0200840340969399 | down | Pathway |
| SOCS2 | -2.1662309558028 | 0.00312929793923999 | down | Pathway |
| STXBP1 | -2.16289231713125 | 0.000162681118293746 | down | Pathway |
| KLF2 | -2.16033796277745 | 0.00188516693694283 | down | Pathway |
| ACVRL1 | -2.15952337078852 | 0.000112551861422631 | down | Pathway |
| FSTL1 | -2.15482458673052 | 0.000711819222909794 | down | Pathway |
| APLNR | -2.15001351852252 | 0.000740009341689421 | down | Pathway |
| NOSTRIN | -2.14837579835896 | 0.00301812310270163 | down | Pathway |
| SLC35F1 | -2.1466788534757 | 0.00704396390732661 | down | NA |
| UST | -2.14639960879839 | 0.000217045050979065 | down | Pathway |
| TPPP3 | -2.14589718531972 | 0.0137536074881435 | down | Pathway |
| VAMP5 | -2.14427410769805 | 0.00136971994879107 | down | Pathway |
| SLC26A7 | -2.14329457543163 | 0.0288288826055289 | down | Pathway |
| ITPK1 | -2.14239623303148 | 0.0000276807225929861 | down | Pathway |
| GABARAPL1 | -2.14212437409292 | 0.0000109146579501064 | down | Pathway |
| BMP4 | -2.14074446212685 | 0.013997789543765 | down | Pathway |
| PDE1A | -2.13099498216153 | 0.000431779887936525 | down | NA |
| PROCR | -2.13047027901179 | 0.00335368941960841 | down | Pathway |
| SNX21 | -2.13040135103782 | 0.0000164035762947684 | down | NA |
| HMCN2 | -2.12614470197237 | 0.0127745751752361 | down | NA |
| CD302 | -2.12577883021267 | 0.000700503270072223 | down | NA |

| | | | | |
|---|---|---|---|---|
| TRIM55 | -2.12562173674477 | 0.0345671917537695 | down | Pathway |
| MYO1C | -2.12120085318963 | 0.0000165697766706899 | down | Pathway |
| CYP11A1 | -2.12061372580603 | 0.016749076455753 | down | Pathway |
| ST6GALNAC6 | -2.1205915423265 | 0.0000138892049247438 | down | Pathway |
| FAM124A | -2.11931241319313 | 0.0118988882291875 | down | NA |
| ASB9 | -2.11921830248658 | 0.000291821939599855 | down | NA |
| KDR | -2.11608972409529 | 0.00017500910237081 | down | Pathway |
| FAM166B | -2.11495282210235 | 0.00810573283533351 | down | NA |
| PLTP | -2.11254408389732 | 0.0327481301295951 | down | Pathway |
| GLI1 | -2.11235530123409 | 0.0312884340823514 | down | Pathway |
| UGP2 | -2.11029135114629 | 0.0000138892049247438 | down | Pathway |
| PYGL | -2.10720068511866 | 0.00235263051078067 | down | Pathway |
| EHBP1 | -2.10697728477521 | 0.0000109146579501064 | down | NA |
| PHLDB2 | -2.10682810162755 | 0.0021078631042849 | down | Pathway |
| GULP1 | -2.10586777803114 | 0.000254485834340982 | down | Pathway |
| CFL2 | -2.1036981088485 | 0.0000174267445510039 | down | Pathway |
| NMT2 | -2.10356565775371 | 0.000116515980185141 | down | Pathway |
| ARHGAP24 | -2.10332047610127 | 0.000239941225756581 | down | Pathway |
| EFNB1 | -2.1018137772875 | 0.00107479335798494 | down | Pathway |
| RGCC | -2.0989362700451 | 0.000448856211033562 | down | Pathway |
| TMEM221 | -2.09718358364698 | 0.0308917236024481 | down | NA |
| CACHD1 | -2.09664538895948 | 0.00870120133366826 | down | Pathway |
| RP11-122A3.2 | -2.09529444148548 | 0.00119633534347166 | down | NA |
| ALDH6A1 | -2.09375143647149 | 0.00193278694790573 | down | Pathway |
| NDUFA4L2 | -2.08962105527002 | 0.000532787255821952 | down | NA |

| | | | | |
|---|---|---|---|---|
| DCLK1 | -2.08897674029832 | 0.019551694635488 | down | Pathway |
| RAPGEF3 | -2.08825772555247 | 0.00181673268362257 | down | Pathway |
| FAM213A | -2.08824360128562 | 0.000356273730229586 | down | NA |
| STON1 | -2.0872455416721 | 0.00499531273823489 | down | NA |
| NUDT7 | -2.08615644202864 | 0.000405010877871049 | down | Pathway |
| CIB2 | -2.08281768843334 | 0.0183883108996659 | down | Pathway |
| MXRA7 | -2.08201550662261 | 0.000283631640767584 | down | NA |
| MSX1 | -2.08169873783649 | 0.00182453898514806 | down | Pathway |
| ECHDC1 | -2.0814798018535 | 0.00247529312572664 | down | Pathway |
| CDH13 | -2.07837882517048 | 0.000471013104407419 | down | Pathway |
| PLCL2 | -2.07580429767431 | 0.0020816284783849 | down | Pathway |
| ROBO3 | -2.07383160099551 | 0.000763802742473218 | down | Pathway |
| CYP39A1 | -2.07261242822938 | 0.0237126773179452 | down | Pathway |
| EEPD1 | -2.06975271416584 | 0.00471405621268965 | down | Pathway |
| CD93 | -2.06883889882302 | 0.0000872680034807056 | down | NA |
| LAMC1 | -2.0682437226186 | 0.000626417394262855 | down | Pathway |
| PLCL1 | -2.06724497102471 | 0.000476126243266669 | down | Pathway |
| PKD1L1 | -2.06676312653583 | 0.00659550276027539 | down | Pathway |
| MYO7B | -2.06673693693965 | 0.00437358676856239 | down | Pathway |
| SPNS2 | -2.06663451485803 | 0.00164634102680297 | down | Pathway |
| TPPP | -2.06554573400674 | 0.0126888140897027 | down | Pathway |
| KIF17 | -2.0644749543305 | 0.00109190146331252 | down | Pathway |
| PLEKHH2 | -2.06357795606748 | 0.00990019919880412 | down | Pathway |
| PTPRM | -2.06251263107762 | 0.000166187451735925 | down | Pathway |
| PCDH12 | -2.06168596479316 | 0.0000323174020300252 | down | Pathway |

| | | | | |
|---|---|---|---|---|
| FGFR1 | -2.06059645160512 | 0.0011272928520189 | down | Pathway |
| MALL | -2.05812820193356 | 0.0074157114338491 | down | Pathway |
| RGL1 | -2.05673307789524 | 0.010403606957345 | down | Pathway |
| SMAD6 | -2.05440778175498 | 0.00630377850534343 | down | Pathway |
| CLEC1A | -2.05295106535494 | 0.000067262681463929 | down | NA |
| AMOTL1 | -2.05237941448808 | 0.000264261417358478 | down | Pathway |
| RECK | -2.05021218354473 | 0.000268320741088932 | down | Pathway |
| RFTN2 | -2.04787905725103 | 0.000253346530197055 | down | NA |
| ITPRIPL1 | -2.04532419141149 | 0.00395298694804414 | down | NA |
| EHHADH | -2.04471350570649 | 0.00074141469115704 | down | Pathway |
| C3 | -2.04441588074182 | 0.00996872930780224 | down | Pathway |
| NLGN4X | -2.04367488517406 | 0.016132695267151 | down | Pathway |
| ADAM22 | -2.04254644404003 | 0.00915072993447028 | down | Pathway |
| C19orf12 | -2.04042816829058 | 0.0000148509983906441 | down | Pathway |
| RASA3 | -2.04029551511782 | 0.000363661472296406 | down | Pathway |
| ADSSL1 | -2.03969129392695 | 0.000248430078437311 | down | NA |
| ACR | -2.0377641039692 | 0.00124590520515647 | down | Pathway |
| NOVA2 | -2.03736996615211 | 0.000420214028467988 | down | Pathway |
| RASGRP2 | -2.0330723146407 | 0.00568630668358794 | down | Pathway |
| PDP2 | -2.03073983128917 | 0.0012117962851961 | down | NA |
| TMC1 | -2.03056737340036 | 0.0300726247451444 | down | Pathway |
| COL21A1 | -2.03047505568641 | 0.010847416030757 | down | NA |
| NRP1 | -2.02789096407208 | 0.0000506791357951246 | down | Pathway |
| TRABD2A | -2.02598439649012 | 0.016232843764579 | down | Pathway |

| | | | | |
|---|---|---|---|---|
| CLDN11 | -2.02596889061728 | 0.019358519395915 | down | Pathway |
| RFTN1 | -2.02567457720216 | 0.000143752115808916 | down | Pathway |
| FFAR2 | -2.02535661703133 | 0.0175476881499325 | down | Pathway |
| PPP1R3G | -2.02148290505592 | 0.00491485921465165 | down | Pathway |
| TSKU | -2.02112901396823 | 0.000704610227076125 | down | Pathway |
| ATP1B2 | -2.02087669741172 | 0.0222863569022569 | down | Pathway |
| PHLDB1 | -2.02024655134705 | 0.000113090619824214 | down | Pathway |
| IRAK3 | -2.01793680434328 | 0.00638740141816541 | down | Pathway |
| TMEM56 | -2.01586424459836 | 0.0472473341140652 | down | NA |
| HOXA9 | -2.01435005111031 | 0.00350510071357994 | down | Pathway |
| ACAT1 | -2.01146720768784 | 0.0000908288860926587 | down | Pathway |
| LYNX1 | -2.01131630709493 | 0.0000947346685538899 | down | Pathway |
| APBB1IP | -2.01102122781169 | 0.0032357903437502 | down | NA |
| SEC14L5 | -2.0051835218949 | 0.0109231314627126 | down | NA |
| PDZD4 | -2.00220032517122 | 0.000308665816614267 | down | NA |
| ARHGAP28 | -1.99931118624393 | 0.0102620111971817 | down | Pathway |
| SPRY1 | -1.99911139672865 | 0.000628853343204391 | down | Pathway |
| PDE3A | -1.99513999273183 | 0.00422499544835807 | down | Pathway |
| FAM212A | -1.99467366892557 | 0.00243192549206861 | down | NA |
| AC006547.14 | -1.99421278406586 | 0.0107110602174137 | down | NA |
| TMEM35 | -1.99229203462872 | 0.0202714868055221 | down | NA |
| NNMT | -1.98865737789243 | 0.00339275830195315 | down | Pathway |
| BOC | -1.98828853255036 | 0.011115775963202 | down | Pathway |
| UTRN | -1.98802569437571 | 0.0000653191222370623 | down | Pathway |

| | | | | |
|---|---|---|---|---|
| BTBD6 | -1.98713703089729 | 0.0000594803468915483 | down | NA |
| PCCA | -1.98629843847428 | 0.0000820462790595965 | down | Pathway |
| TEAD1 | -1.98331812444994 | 0.00254356709610707 | down | Pathway |
| HADH | -1.98179811448032 | 0.000519472046918103 | down | Pathway |
| CRTAP | -1.98140958576944 | 0.0000712512997588031 | down | Pathway |
| KIAA1462 | -1.97903384052347 | 0.00092693459266144 | down | NA |
| LIMA1 | -1.97768464541383 | 0.000357020895717162 | down | Pathway |
| REM1 | -1.97670526578294 | 0.000971678499600897 | down | Pathway |
| XG | -1.97667324554329 | 0.042185682884104 | down | Pathway |
| TCEAL7 | -1.97595858078002 | 0.0055751785436753 | down | NA |
| ACOT1 | -1.97544185090081 | 0.00998545280970259 | down | Pathway |
| POLR3GL | -1.9745655075771 | 0.000448856211033562 | down | NA |
| ADAMTS3 | -1.97347107403116 | 0.00547574673144751 | down | Pathway |
| HIGD1B | -1.96580349230075 | 0.00462133890211985 | down | NA |
| ZDBF2 | -1.96312057106938 | 0.00706190750915601 | down | NA |
| CCBE1 | -1.96195709187234 | 0.0448362270108187 | down | Pathway |
| MCTP1 | -1.96176775442993 | 0.00174267483578896 | down | Pathway |
| STAT5A | -1.9590575768852 | 0.00117951915899427 | down | Pathway |
| KCTD12 | -1.95687921450341 | 0.00138046603939258 | down | NA |
| RNF180 | -1.95634211701375 | 0.014136352287295 | down | Pathway |
| CCDC107 | -1.95616407233923 | 0.000205446686047126 | down | NA |
| TMEM178B | -1.95613675382748 | 0.0102268892774019 | down | NA |
| ORMDL3 | -1.95488627906012 | 0.000763802742473218 | down | Pathway |
| KIF1C | -1.95049310128619 | 0.0000337437858946321 | down | Pathway |

| | | | | |
|---|---|---|---|---|
| SMIM10 | -1.95024235279794 | 0.00220454450982573 | down | NA |
| VKORC1L1 | -1.94908943868579 | 0.000131712152793352 | down | Pathway |
| ZNF106 | -1.94907434386524 | 0.000101349074328654 | down | Pathway |
| SNTA1 | -1.94594570647451 | 0.0000985411462195632 | down | Pathway |
| TNIP1 | -1.94593233430578 | 0.0000293106134078413 | down | Pathway |
| RP11-3N2.13 | -1.94545884137913 | 0.0110635314110508 | down | NA |
| CLU | -1.94448361126248 | 0.00770064857934628 | down | Pathway |
| THRB | -1.94342117920551 | 0.00141093543080837 | down | Pathway |
| VIP | -1.94311589582721 | 0.00132503330749421 | down | Pathway |
| PLS3 | -1.94083106398271 | 0.0000236395522203272 | down | Pathway |
| OXCT1 | -1.94026341193186 | 0.000876859143059688 | down | Pathway |
| TCN2 | -1.93969643597869 | 0.0142716554505583 | down | NA |
| VSIG2 | -1.93947820640905 | 0.0184838246216 | down | NA |
| PLEKHA4 | -1.93824126495257 | 0.00704505451770364 | down | Pathway |
| AVPI1 | -1.93749927921681 | 0.00342730582790048 | down | NA |
| TMEM200B | -1.93697200760582 | 0.0000765099910680501 | down | NA |
| NOV | -1.93637854522149 | 0.00711884141653348 | down | NA |
| THSD1 | -1.93463094643869 | 0.000116145582440322 | down | Pathway |
| SRCRB4D | -1.93302878094115 | 0.00315255128750998 | down | NA |
| RHOQ | -1.93220880271496 | 0.000268320741088932 | down | Pathway |
| ZFHX4 | -1.93197507180845 | 0.00777856227005529 | down | NA |
| C11orf96 | -1.93011057040093 | 0.00757408497251189 | down | NA |
| PDZRN3 | -1.92999927821291 | 0.0142591766331992 | down | Pathway |

| | | | | |
|---|---|---|---|---|
| OSR2 | -1.92839699207954 | 0.0411739247249634 | down | Pathway |
| ZEB1 | -1.92485036819349 | 0.00042051996284283 | down | Pathway |
| CALCRL | -1.92279684601088 | 0.00323032043386279 | down | Pathway |
| SERPING1 | -1.92165756706452 | 0.0082474348441763 | down | Pathway |
| ANO2 | -1.91861158674834 | 0.0026354738940612 | down | Pathway |
| BST1 | -1.91785774448464 | 0.000720894017791845 | down | Pathway |
| OLFML1 | -1.91740443915049 | 0.000783542272291761 | down | NA |
| C10orf11 | -1.9167314595645 | 0.00188855403965796 | down | NA |
| ACAA2 | -1.91668130921037 | 0.000397545133749527 | down | Pathway |
| SUCLA2 | -1.91346922156416 | 0.00033598535174166 | down | Pathway |
| TM7SF2 | -1.90950222905951 | 0.00624888804618131 | down | Pathway |
| ADHFE1 | -1.90822854906813 | 0.0011161451102904 | down | Pathway |
| BCL6B | -1.90661558035348 | 0.0000983664357695885 | down | Pathway |
| ZNF219 | -1.90627682619212 | 0.000083142834261072 | down | Pathway |
| EPB41L2 | -1.90583468570227 | 0.00181969213656071 | down | Pathway |
| CD99L2 | -1.9044465312172 | 0.0000421738197942172 | down | Pathway |
| EDA | -1.90414555245124 | 0.000601178163986356 | down | Pathway |
| THRA | -1.89863431518214 | 0.0000836233586411006 | down | Pathway |
| LGALS1 | -1.89619846036072 | 0.00277543029340354 | down | Pathway |
| PLAGL1 | -1.89568138543154 | 0.00972254968752351 | down | NA |
| ARHGAP31 | -1.89299954834767 | 0.00049018992375733 | down | Pathway |
| TUBB2A | -1.89126934835612 | 0.00206752002085825 | down | Pathway |
| DPP4 | -1.89077843372564 | 0.0407394921189933 | down | Pathway |
| HOXA11 | -1.89066844024791 | 0.0450820104403733 | down | Pathway |

| | | | | |
|---|---|---|---|---|
| AASS | -1.89063502016426 | 0.00127967686689168 | down | Pathway |
| SOX18 | -1.89026515279437 | 0.00511126324033011 | down | Pathway |
| FLI1 | -1.89019972202004 | 0.00334209626234771 | down | Pathway |
| MID1 | -1.88983752179496 | 0.0307426485825176 | down | Pathway |
| AK4 | -1.88838444699877 | 0.00272748625291439 | down | Pathway |
| PDGFD | -1.88786372973981 | 0.00265750967133912 | down | Pathway |
| DAPK2 | -1.88763093786424 | 0.0048512817694514 | down | Pathway |
| MECOM | -1.88746001534679 | 0.0435419820319113 | down | Pathway |
| AAMDC | -1.8863230254208 | 0.0000135579150977941 | down | Pathway |
| ARHGEF6 | -1.88578807362397 | 0.00628844279303909 | down | Pathway |
| ARHGEF40 | -1.88399900610801 | 0.00147697324715866 | down | Pathway |
| PPP1R36 | -1.88128287658951 | 0.0169804738628558 | down | NA |
| ESYT1 | -1.87913254793198 | 0.0000903371821431127 | down | Pathway |
| LRRC34 | -1.87773192344921 | 0.0364561878540886 | down | NA |
| FILIP1 | -1.87581461683338 | 0.00246376790917567 | down | NA |
| LAMB3 | -1.87578466663576 | 0.0212899946146452 | down | Pathway |
| NAV3 | -1.87517195901378 | 0.000205446686047126 | down | Pathway |
| GPATCH11 | -1.87493655289372 | 0.000241403265626349 | down | NA |
| GPX4 | -1.87474023851823 | 0.000176792164193682 | down | Pathway |
| MEIS2 | -1.87413533790853 | 0.00151879405288415 | down | Pathway |
| PTPN21 | -1.87408524640604 | 0.000583409684636326 | down | NA |
| HOXB6 | -1.87120591838017 | 0.000840027060867341 | down | Pathway |
| NPR2 | -1.86974379757694 | 0.000476928494523595 | down | Pathway |
| GRK5 | -1.86897354987849 | 0.000206395232206536 | down | Pathway |

| | | | | |
|---|---|---|---|---|
| SLC10A6 | -1.86624524192032 | 0.0051149138931311 | down | Pathway |
| FAM126A | -1.86576682795893 | 0.000121388009059353 | down | Pathway |
| FLT4 | -1.86062385747162 | 0.000395784895508871 | down | Pathway |
| CSGALNACT1 | -1.85996898329176 | 0.000841961641436113 | down | Pathway |
| ETFB | -1.85886023321821 | 0.00147744477533225 | down | Pathway |
| SULT1C4 | -1.85726366536314 | 0.00613114311322311 | down | Pathway |
| FAM43A | -1.85710502380287 | 0.000459962776335258 | down | NA |
| SNN | -1.85588723511854 | 0.00052570559702797 | down | Pathway |
| TMBIM1 | -1.85530742047031 | 0.00022370204671549 | down | Pathway |
| BTBD8 | -1.85444889058909 | 0.00666528593643077 | down | NA |
| SH3TC2 | -1.85371184236446 | 0.00213856146020748 | down | Pathway |
| TSPAN18 | -1.84965530495259 | 0.00136480741696431 | down | NA |
| PRKCDBP | -1.84920438456763 | 0.00408481621367174 | down | NA |
| LRFN5 | -1.84544680614328 | 0.0389587849772827 | down | Pathway |
| PARD3B | -1.84542850672993 | 0.00064772397790125 | down | Pathway |
| CPE | -1.84432248694664 | 0.00469727338401908 | down | Pathway |
| PRKD1 | -1.84431933443334 | 0.000644903205574493 | down | Pathway |
| ADCY4 | -1.84345427164264 | 0.000203752307486219 | down | Pathway |
| FAM49A | -1.84235220451722 | 0.00651970178464188 | down | NA |
| MARC2 | -1.84198779956807 | 0.00742082169996448 | down | NA |
| CFI | -1.84165102859095 | 0.0147515441940397 | down | NA |
| PXDC1 | -1.84101836686137 | 0.00422050513357063 | down | NA |
| DTX4 | -1.84062895557691 | 0.00180818451145274 | down | NA |

| | | | | |
|---|---|---|---|---|
| HOXC8 | -1.83983040501424 | 0.0138294782983929 | down | Pathway |
| ANKDD1A | -1.83864220789836 | 0.00294876592879868 | down | NA |
| ABLIM1 | -1.83803059510462 | 0.00270351074517001 | down | NA |
| DIXDC1 | -1.83659442710389 | 0.000311104455999785 | down | Pathway |
| ADRBK2 | -1.8353452309721 | 0.00121028633221942 | down | NA |
| ABHD1 | -1.83437150511383 | 0.0142274347695393 | down | Pathway |
| KCNH4 | -1.83436951499374 | 0.0229048097093173 | down | Pathway |
| GPRC5B | -1.83422672947511 | 0.0246070864225715 | down | Pathway |
| HDAC9 | -1.83288011480943 | 0.00194512444061659 | down | Pathway |
| MPDZ | -1.83238721556675 | 0.0020816284783849 | down | Pathway |
| SH2D3C | -1.83230960800321 | 0.00148633732729716 | down | Pathway |
| CHST7 | -1.83193557612623 | 0.00139305578450717 | down | Pathway |
| PCYOX1 | -1.83030749309447 | 0.00148633732729716 | down | Pathway |
| TLE1 | -1.82883361205247 | 0.0239952983476666 | down | Pathway |
| GPR153 | -1.82836841285753 | 0.00146990157599343 | down | NA |
| FYN | -1.82549327392426 | 0.00258052237930075 | down | Pathway |
| TSPAN12 | -1.8254496105367 | 0.0159462793265593 | down | Pathway |
| TK2 | -1.82477126652181 | 0.00011597175870838 | down | NA |
| EPHX1 | -1.82402918382308 | 0.00495217487937342 | down | Pathway |
| LRRC70 | -1.82383448669945 | 0.000779630130756003 | down | NA |
| SIX2 | -1.82081363439266 | 0.0162995678512346 | down | Pathway |
| FOXI2 | -1.82070645603191 | 0.0273331740055999 | down | NA |
| CCDC152 | -1.81951005906336 | 0.00197379925851253 | down | NA |
| ITGA1 | -1.81809356551362 | 0.000649370895645139 | down | Pathway |

| | | | | |
|---|---|---|---|---|
| FAH | -1.81630995734016 | 0.00318854345568665 | down | Pathway |
| ACSL4 | -1.81587639576728 | 0.00111333107244355 | down | Pathway |
| SLITRK4 | -1.81394541987993 | 0.0296223385977939 | down | Pathway |
| ADAMTS1 | -1.8133533588032 | 0.00795240081953702 | down | Pathway |
| SCN2A | -1.81069708549687 | 0.00289259669398799 | down | Pathway |
| PER1 | -1.81033494372688 | 0.00165724320113718 | down | Pathway |
| LRRC4B | -1.80967015015648 | 0.0088003251403496 | down | Pathway |
| HOXB4 | -1.80775345166223 | 0.0027881039353636 | down | Pathway |
| SHROOM4 | -1.80721118446773 | 0.00142596384212134 | down | Pathway |
| ENPEP | -1.80412281335589 | 0.00121028633221942 | down | Pathway |
| FSTL3 | -1.80288179003171 | 0.00622453575354714 | down | Pathway |
| ACKR4 | -1.80237980572757 | 0.0142946430845491 | down | Pathway |
| ALAD | -1.8023217105843 | 0.000583409684636326 | down | Pathway |
| WTIP | -1.80223824419703 | 0.00570085537493398 | down | Pathway |
| APOLD1 | -1.80124034686695 | 0.00562991818020628 | down | Pathway |
| GAS1 | -1.79904092754323 | 0.0233177760667566 | down | Pathway |
| LRMP | -1.79887521133868 | 0.0203724909852069 | down | NA |
| FLNC | -1.79871067942222 | 0.0345731093197016 | down | Pathway |
| HIC1 | -1.79834696866204 | 0.00623399091702636 | down | Pathway |
| SH3KBP1 | -1.79430066904843 | 0.00295295674377838 | down | Pathway |
| FAM69B | -1.79341823401303 | 0.00643343171522469 | down | NA |
| CARD6 | -1.79291787646834 | 0.00281418192569626 | down | NA |
| FAM69A | -1.79172024903371 | 0.000670752647235661 | down | NA |
| LTBP4 | -1.79055586771675 | 0.000644903205574493 | down | Pathway |

| | | | | |
|---|---|---|---|---|
| GPC6 | -1.78974785579891 | 0.00721812611063696 | down | Pathway |
| CPXM2 | -1.78972662715658 | 0.011900316418752 | down | NA |
| NEXN | -1.78799572580657 | 0.00101779294392152 | down | Pathway |
| RNF125 | -1.7872886327798 | 0.006366149643079 | down | NA |
| CPNE2 | -1.78618330928243 | 0.00120369971264521 | down | Pathway |
| KLHL5 | -1.78493541002345 | 0.0000922735054215946 | down | NA |
| FOXO1 | -1.78480763385791 | 0.000194405063018176 | down | Pathway |
| SCARF1 | -1.78413184257021 | 0.00170181458497422 | down | Pathway |
| FLT1 | -1.78362092051412 | 0.000601594805013111 | down | Pathway |
| KLRF1 | -1.78336014789347 | 0.0386204255088704 | down | NA |
| BVES | -1.78202708887565 | 0.000479061444774117 | down | Pathway |
| ANXA6 | -1.78090370111457 | 0.000268320741088932 | down | Pathway |
| MRO | -1.78024399551469 | 0.00281418192569626 | down | NA |
| HABP4 | -1.78005934311956 | 0.0000878102623376788 | down | Pathway |
| NFIX | -1.77894442679598 | 0.00233203390978014 | down | Pathway |
| EPB41L3 | -1.77838491030593 | 0.0084179667371984 | down | Pathway |
| PPARA | -1.7773459370723 | 0.00175975367170712 | down | Pathway |
| SNED1 | -1.77724162036237 | 0.0000762267464760712 | down | Pathway |
| SCIN | -1.77634304788663 | 0.0127684265683857 | down | Pathway |
| RHOJ | -1.7755072328378 | 0.000496500461346152 | down | Pathway |
| GUCY1A2 | -1.7753023272684 | 0.000490830955935515 | down | Pathway |
| CYBRD1 | -1.76811819037851 | 0.0203260089093394 | down | Pathway |
| KALRN | -1.76791268880003 | 0.00937897026901019 | down | Pathway |
| NRIP2 | -1.76735061658381 | 0.0272070316154518 | down | NA |
| LRP3 | -1.76729447626239 | 0.00246696522349474 | down | Pathway |
| ULBP3 | -1.767072959906 | 0.0380460293762841 | down | NA |

| | | | | |
|---|---|---|---|---|
| LPAR1 | -1.76649869949642 | 0.0011621952937401 | down | Pathway |
| C1S | -1.7660056710748 | 0.014605198914439 | down | NA |
| ITGA6 | -1.7657461236148 | 0.000141818334379598 | down | Pathway |
| DUSP6 | -1.76276161344848 | 0.00042386930184934 | down | Pathway |
| CCR10 | -1.76206342660198 | 0.00553942800669223 | down | Pathway |
| MAP1B | -1.7619726927547 | 0.00627514402292473 | down | Pathway |
| CRB1 | -1.75981256377255 | 0.0272780190299227 | down | Pathway |
| NDRG4 | -1.75944830175746 | 0.0149566307950456 | down | Pathway |
| HOXD8 | -1.75889401152841 | 0.0068387118260065 | down | Pathway |
| PPM1L | -1.75752585786238 | 0.0071136519041166 | down | Pathway |
| ABHD5 | -1.75592165561191 | 0.00123650385139297 | down | Pathway |
| BHLHE22 | -1.75484218916872 | 0.0356488232470164 | down | NA |
| PRICKLE2 | -1.75437372138211 | 0.00112531383076672 | down | Pathway |
| CLIC2 | -1.75316559783142 | 0.00144858652271961 | down | Pathway |
| CHCHD10 | -1.75295951626748 | 0.00213262441477563 | down | Pathway |
| ANG | -1.75231269281417 | 0.000364692473193508 | down | Pathway |
| APOL6 | -1.7522576968062 | 0.00233570228654904 | down | Pathway |
| PTGER4 | -1.74788529340768 | 0.00764391344791033 | down | Pathway |
| SH3BP5 | -1.74753938705479 | 0.000433136280562955 | down | Pathway |
| DDIT4L | -1.74484343963263 | 0.0151589477491166 | down | NA |
| TIMP3 | -1.74164298191291 | 0.00347728048226657 | down | Pathway |
| PHLDA3 | -1.74042263275101 | 0.00546883555736987 | down | Pathway |
| S100A10 | -1.73907735758194 | 0.00680967701791538 | down | Pathway |
| FKBP5 | -1.73459858965337 | 0.0188553799757236 | down | NA |
| SYDE1 | -1.73210390612729 | 0.000068298800819865 9 | down | Pathway |

| | | | | |
|---|---|---|---|---|
| SGCB | -1.73202628488218 | 0.00155839873950891 | down | Pathway |
| CYS1 | -1.73048912203414 | 0.0225904851453788 | down | NA |
| CYSTM1 | -1.72932449105665 | 0.00042051996284283 | down | NA |
| PPL | -1.72712524867329 | 0.0140506660780196 | down | Pathway |
| DIAPH2 | -1.72701285518083 | 0.000297754513483632 | down | Pathway |
| NUAK1 | -1.72660852078328 | 0.00158924176152983 | down | Pathway |
| ME3 | -1.72588549702581 | 0.000322109665703003 | down | NA |
| ATP2B4 | -1.72575511076373 | 0.00119780135899441 | down | Pathway |
| OAF | -1.72467935619694 | 0.00915748616039275 | down | NA |
| PDGFRA | -1.72407446527269 | 0.00652367365079394 | down | Pathway |
| MFNG | -1.72327984531446 | 0.00276951318756057 | down | Pathway |
| NUDT8 | -1.72153814730048 | 0.00744512343304905 | down | Pathway |
| STARD9 | -1.72130890868074 | 0.000601594805013111 | down | Pathway |
| GNA14 | -1.71932566602442 | 0.00126193249823639 | down | Pathway |
| GATA2 | -1.71908068524889 | 0.0143302437627075 | down | Pathway |
| ABHD15 | -1.71800841627494 | 0.00145102726031926 | down | NA |
| SLC16A2 | -1.71749084200869 | 0.00626770841408716 | down | Pathway |
| ZNF677 | -1.71736552542127 | 0.0263727942984936 | down | NA |
| QKI | -1.7164838114097 | 0.00120073501325808 | down | Pathway |
| TTC7B | -1.71642964287521 | 0.000830139190852364 | down | Pathway |
| ARHGAP10 | -1.71574863650361 | 0.00145471215127804 | down | Pathway |
| TLR4 | -1.71530617068676 | 0.0116510406628552 | down | Pathway |
| ITIH3 | -1.71318011589327 | 0.0369657423521988 | down | NA |
| NR5A2 | -1.71276141817879 | 0.0062649241155265 | down | Pathway |

| | | | | |
|---|---|---|---|---|
| GIPC3 | -1.70975499190058 | 0.00115836595329297 | down | NA |
| DST | -1.70708716102188 | 0.00162111992478466 | down | Pathway |
| CYB5A | -1.70564890076797 | 0.0320395898745852 | down | NA |
| RGS17 | -1.70292275522234 | 0.0270660984484938 | down | NA |
| TKT | -1.69772721054566 | 0.00199327714734499 | down | NA |
| STEAP1 | -1.69661020201886 | 0.00270351074517001 | down | NA |
| RBPMS | -1.69598078791368 | 0.015622161643148 | down | Pathway |
| NEK7 | -1.69539147784247 | 0.00115836595329297 | down | Pathway |
| PRPH2 | -1.69449444398152 | 0.0260578968111256 | down | Pathway |
| REV3L | -1.69398153625525 | 0.0000981323114659227 | down | Pathway |
| SHMT1 | -1.69358865837991 | 0.0007283132901898 | down | Pathway |
| LONRF3 | -1.69282344300797 | 0.00168542494454958 | down | NA |
| SLC9A3R2 | -1.69233688642646 | 0.000467178037277391 | down | Pathway |
| RRAS2 | -1.69042621404476 | 0.0100479114758072 | down | Pathway |
| SYNE1 | -1.68805682491592 | 0.000235721954118244 | down | Pathway |
| ITGB1 | -1.68801292118626 | 0.00185192627706817 | down | Pathway |
| ZNF354C | -1.68783098970681 | 0.0149034398144706 | down | NA |
| TNFSF12 | -1.68570104225566 | 0.00209453764137192 | down | Pathway |
| PMEPA1 | -1.68447480503974 | 0.000841961641436113 | down | Pathway |
| GDPD5 | -1.68432643490217 | 0.0061798066195251 | down | Pathway |
| MDH1 | -1.67888029409353 | 0.0000674325767586959 | down | NA |
| FAM26E | -1.67617364581364 | 0.000720894017791845 | down | NA |
| PJA1 | -1.6760580713873 | 0.000303280961205611 | down | NA |
| CSRP2 | -1.67410357331232 | 0.0210829232423331 | down | Pathway |

| | | | | |
|---|---|---|---|---|
| SLIT2 | -1.67321587063017 | 0.00283072509777522 | down | Pathway |
| OMD | -1.67242642303182 | 0.0351441363414662 | down | Pathway |
| KIAA1324L | -1.67172291821668 | 0.0108793314762571 | down | NA |
| GFOD1 | -1.67114674339424 | 0.000614423176741171 | down | NA |
| DYNC1I1 | -1.67066971688931 | 0.00154132424937158 | down | Pathway |
| ATP8B4 | -1.67042777788973 | 0.00340418843214864 | down | Pathway |
| MDFIC | -1.67023282617172 | 0.000165578754766909 | down | Pathway |
| PLCE1 | -1.67016655232556 | 0.049680191317224 | down | Pathway |
| C16orf86 | -1.67014265147683 | 0.00106915086935269 | down | NA |
| PARVA | -1.66964291302857 | 0.0000867662331740805 | down | Pathway |
| TSPAN3 | -1.66899010787331 | 0.000644903205574493 | down | NA |
| CCDC68 | -1.6682326698738 | 0.0260770851504337 | down | NA |
| HSD17B1 | -1.66800387980656 | 0.000766378302764361 | down | Pathway |
| PDE9A | -1.66518749551518 | 0.0216005607968023 | down | Pathway |
| WLS | -1.66405071831858 | 0.00385178115848343 | down | Pathway |
| HAAO | -1.66354609957392 | 0.0230570427441117 | down | Pathway |
| LMO2 | -1.66339591739823 | 0.00187418520764299 | down | NA |
| GIMAP4 | -1.66303023368713 | 0.00928600159201033 | down | NA |
| RGS5 | -1.65691765052083 | 0.0344590193497764 | down | Pathway |
| HSD17B11 | -1.65663954721329 | 0.0025953531753237 | down | Pathway |
| KIRREL | -1.65652282305255 | 0.00156894722727738 | down | NA |
| FAM184A | -1.65453742376615 | 0.00717850825503638 | down | NA |
| CPNE8 | -1.65312662773577 | 0.00220454450982573 | down | Pathway |
| ADCK3 | -1.65168300951413 | 0.00106481929211232 | down | NA |

| | | | | |
|---|---|---|---|---|
| DPF3 | -1.65156022767964 | 0.0186894721379191 | down | NA |
| RPS6KA2 | -1.65135978908635 | 0.00158498331759283 | down | Pathway |
| C1R | -1.65118506270233 | 0.0400061635272468 | down | NA |
| PRDX6 | -1.6510122837661 | 0.000339081478581284 | down | Pathway |
| SCRN2 | -1.65014470373559 | 0.00106481398796966 | down | NA |
| MBNL2 | -1.64826189399098 | 0.000146406827745791 | down | NA |
| CHST3 | -1.64758000962539 | 0.000710467195448009 | down | Pathway |
| COL15A1 | -1.64632089944675 | 0.000924520437151173 | down | Pathway |
| SULT1B1 | -1.6448135790078 | 0.0211530176292111 | down | Pathway |
| SHISA6 | -1.64425761560434 | 0.0330775009997538 | down | Pathway |
| COL4A2 | -1.64363628770959 | 0.000750777485406687 | down | Pathway |
| SELENBP1 | -1.64340480231398 | 0.0200399959949144 | down | NA |
| CYSLTR1 | -1.64313146023425 | 0.0127030362887505 | down | Pathway |
| ABHD14B | -1.64236206066742 | 0.000141818334379598 | down | Pathway |
| ENPP3 | -1.64202483151264 | 0.00358870986093913 | down | Pathway |
| LRIG3 | -1.63766839878245 | 0.0314509143231254 | down | Pathway |
| MT1E | -1.6364640709574 | 0.0102071421219476 | down | Pathway |
| MMP15 | -1.63559269976496 | 0.0169021173048264 | down | Pathway |
| SESN3 | -1.63312406076941 | 0.00558149148714527 | down | Pathway |
| ZBTB20 | -1.63267577319873 | 0.00276849610528115 | down | Pathway |
| MAPK11 | -1.63262038288554 | 0.00524068276653727 | down | Pathway |
| INPP5K | -1.6300235450127 | 0.0000698332812978121 | down | Pathway |
| MFSD4 | -1.62941353615014 | 0.0102771029268131 | down | NA |
| ARL4D | -1.62921180279832 | 0.0321454471072378 | down | Pathway |
| LMCD1 | -1.62887328802373 | 0.00105875870252309 | down | Pathway |

| | | | | |
|---|---|---|---|---|
| GOLGA8H | -1.62866606192409 | 0.0192046020571693 | down | NA |
| STARD8 | -1.62766771547948 | 0.000170060550641866 | down | Pathway |
| NBEAL1 | -1.62586527469024 | 0.0081659346742982 | down | NA |
| IRS2 | -1.62445979809151 | 0.0359249486499276 | down | Pathway |
| TNFRSF21 | -1.62080692254596 | 0.0188411356067593 | down | Pathway |
| ECH1 | -1.6207895439662 | 0.000110023001841881 | down | Pathway |
| PDK2 | -1.6194692910933 | 0.00440701294263352 | down | Pathway |
| TCF4 | -1.61816644188004 | 0.00145553954574202 | down | Pathway |
| DLG2 | -1.61807951060545 | 0.00518239962225961 | down | Pathway |
| GPAT2 | -1.61772449277428 | 0.0269958170015769 | down | Pathway |
| NR1D1 | -1.6162018138138 | 0.000253726815949543 | down | Pathway |
| A4GALT | -1.61367703335569 | 0.00632939494484344 | down | Pathway |
| LRRC8C | -1.61178983779845 | 0.000698758446656829 | down | Pathway |
| CREB5 | -1.61158583795391 | 0.00887711457661303 | down | NA |
| RPH3AL | -1.61000230503452 | 0.000700595594406309 | down | Pathway |
| FBN1 | -1.60961838773746 | 0.0110559690241855 | down | Pathway |
| ZNF541 | -1.60924511279575 | 0.0228270099570833 | down | NA |
| BTD | -1.6073852715901 | 0.000942091324717271 | down | Pathway |
| SBF2 | -1.60680236470459 | 0.00178890584146715 | down | Pathway |
| SEPT4 | -1.60605156286947 | 0.000420397875164547 | down | NA |
| GMFG | -1.60594932457125 | 0.0400293334856324 | down | Pathway |
| PHYHD1 | -1.6059424153342 | 0.0356592623789359 | down | NA |
| VEGFC | -1.60579248067103 | 0.00185553461592785 | down | Pathway |
| FGL2 | -1.60441151753782 | 0.0187723830610946 | down | Pathway |
| CRY2 | -1.59977444132678 | 0.0000683419627159911 | down | Pathway |
| HOXB5 | -1.59927489365955 | 0.00845646038238792 | down | Pathway |

| | | | | |
|---|---|---|---|---|
| ZBTB47 | -1.59899330440535 | 0.0000704163887660646 | down | NA |
| ABHD14A | -1.59797080181479 | 0.0000922735054215946 | down | NA |
| MARCH2 | -1.59652019071146 | 0.000535614279950669 | down | NA |
| DHDDS | -1.59623172004825 | 0.00106869234681663 | down | Pathway |
| PLVAP | -1.59587340673552 | 0.00115836595329297 | down | Pathway |
| CPEB1 | -1.59381495805623 | 0.00480015029323151 | down | Pathway |
| NFIL3 | -1.59368835246355 | 0.0293803406992471 | down | Pathway |
| KAT2B | -1.59128685886812 | 0.000638241665109871 | down | Pathway |
| STEAP1B | -1.58943546961055 | 0.00848953306601889 | down | NA |
| FILIP1L | -1.58488517356793 | 0.0020816284783849 | down | NA |
| RGS20 | -1.58163077714468 | 0.0484134164678539 | down | Pathway |
| VTI1B | -1.58067567316065 | 0.00027467719353159 | down | Pathway |
| WDPCP | -1.57795765433337 | 0.00108922007814007 | down | Pathway |
| TMEM64 | -1.57784410771412 | 0.0247460601671352 | down | Pathway |
| HMCN1 | -1.57611583009785 | 0.0405130478051652 | down | Pathway |
| SLC12A4 | -1.57611534506235 | 0.000364692473193508 | down | Pathway |
| HIBADH | -1.57481489735826 | 0.000253346530197055 | down | Pathway |
| COX14 | -1.57457259397878 | 0.00160027080252149 | down | NA |
| GRASP | -1.57349155087996 | 0.00292100081529363 | down | NA |
| TMEM204 | -1.57322757690567 | 0.000589908254731518 | down | Pathway |
| HNMT | -1.571014792615 | 0.00134483697054065 | down | Pathway |
| ECHS1 | -1.57039831685081 | 0.00184622010573845 | down | Pathway |
| ARHGAP42 | -1.56970577146935 | 0.00132033926647642 | down | Pathway |
| LINC00087 | -1.5679916421295 | 0.00124880008254324 | down | NA |
| MYCBP2 | -1.5675851539617 | 0.000188885063372902 | down | Pathway |

| | | | | |
|---|---|---|---|---|
| NES | -1.56682252912057 | 0.00295087976804644 | down | Pathway |
| FXYD6 | -1.56651183625225 | 0.0357679087705267 | down | Pathway |
| SKAP2 | -1.56440550313912 | 0.00542673173830875 | down | NA |
| AHNAK | -1.56345790625852 | 0.000574338703708593 | down | Pathway |
| PDE5A | -1.56142246759745 | 0.00827144400648057 | down | Pathway |
| GSTP1 | -1.56140254815608 | 0.0348718054713023 | down | Pathway |
| COL4A1 | -1.56014652742222 | 0.00207822151532915 | down | Pathway |
| COLEC12 | -1.5586537354391 | 0.0273178528554498 | down | NA |
| CYP2U1 | -1.55859446729157 | 0.000235214577539393 | down | Pathway |
| WWC2 | -1.55479747343833 | 0.0119163659385013 | down | Pathway |
| IL1R1 | -1.55417251038411 | 0.0136332819593479 | down | Pathway |
| PINK1 | -1.55350952404931 | 0.0000859453541881024 | down | Pathway |
| PIK3R1 | -1.55347749105943 | 0.00176602131458252 | down | Pathway |
| FAM101B | -1.55337584645669 | 0.0001672376444689 | down | NA |
| ID3 | -1.55260091231612 | 0.000845640412420472 | down | Pathway |
| BNIP3L | -1.55241024033735 | 0.00145915024249602 | down | Pathway |
| PEX19 | -1.5522938087804 | 0.000549682823155422 | down | NA |
| GCKR | -1.55190746950297 | 0.0145065711448319 | down | Pathway |
| SDC2 | -1.55114879032921 | 0.0198793867220357 | down | Pathway |
| TMEM163 | -1.54948097113376 | 0.00153501584143883 | down | NA |
| A2M | -1.54759866929622 | 0.0109810184792256 | down | Pathway |
| DAB2 | -1.54706604274031 | 0.0237601310887327 | down | Pathway |
| CCDC82 | -1.54655382092615 | 0.00149459685630432 | down | NA |
| PRUNE2 | -1.54462865071697 | 0.00157914855010636 | down | NA |

| | | | | |
|---|---|---|---|---|
| NR3C1 | -1.54318547960041 | 0.00203055769170349 | down | Pathway |
| TSTD3 | -1.5422732535006 | 0.0000818111593372942 | down | NA |
| ST3GAL3 | -1.54009395416019 | 0.00152495166073959 | down | Pathway |
| DOCK6 | -1.53974351847176 | 0.00127941014151396 | down | Pathway |
| ADH5 | -1.53961949540766 | 0.000118744457927505 | down | Pathway |
| ZSCAN23 | -1.53853359458118 | 0.00345806640655465 | down | NA |
| BEND5 | -1.53551574026704 | 0.00371588644748172 | down | NA |
| TRIM7 | -1.53075066300483 | 0.0251479160968384 | down | NA |
| KLHDC8B | -1.52907186252115 | 0.00773646547756223 | down | Pathway |
| MARCH3 | -1.52814720765958 | 0.0241480842247122 | down | NA |
| CDC42EP5 | -1.52655161831466 | 0.0462131225007804 | down | Pathway |
| THBD | -1.52552540406613 | 0.0175026181017843 | down | Pathway |
| XRCC6BP1 | -1.5250863249887 | 0.000111398699557408 | down | NA |
| LPHN2 | -1.52490011297328 | 0.0379828512949598 | down | NA |
| STAT5B | -1.52318348790952 | 0.00298161373057095 | down | Pathway |
| TNFRSF1B | -1.52309171417316 | 0.0325420765599535 | down | Pathway |
| ROCK2 | -1.52234656156349 | 0.00202831071785562 | down | Pathway |
| ABCA1 | -1.52231136840371 | 0.00917817391011895 | down | Pathway |
| EPHA4 | -1.52115480612799 | 0.00608635739126197 | down | Pathway |
| SRPX2 | -1.52095654896299 | 0.029004708672121 | down | Pathway |
| SLC25A20 | -1.5208833060191 | 0.00166274493222347 | down | Pathway |
| HLA-E | -1.52066245738095 | 0.00880779721169707 | down | Pathway |
| PIR | -1.52052165416424 | 0.0000674325767586959 | down | Pathway |
| PMP22 | -1.51975252594395 | 0.00136971994879107 | down | Pathway |
| HOXD3 | -1.51838959345506 | 0.0306738867016833 | down | Pathway |

| | | | | |
|---|---|---|---|---|
| ZNF132 | -1.51500413912894 | 0.00385388988673581 | down | NA |
| EIF4E3 | -1.51456082005811 | 0.0053894257113624 | down | NA |
| C11orf63 | -1.50972544421516 | 0.00659550276027539 | down | NA |
| PTGDR2 | -1.50561771412522 | 0.0262782614007196 | down | Pathway |
| CYTH3 | -1.50524872286748 | 0.00288565652822036 | down | Pathway |
| SAMD4A | -1.50445590982619 | 0.0295010587002061 | down | NA |
| PAM | -1.50229384523732 | 0.0317104190407771 | down | Pathway |
| TST | -1.50179022359915 | 0.0247830034629426 | down | Pathway |
| SUCLG1 | -1.50093768656181 | 0.000142682493544042 | down | NA |
| PRR5 | -1.50040315435138 | 0.000476723974429816 | down | Pathway |
| TMEM206 | 1.50206300104083 | 0.00357032289951928 | up | NA |
| TPBG | 1.50229584961093 | 0.0282569515546027 | up | Pathway |
| TSEN54 | 1.5024374048994 | 0.000142828599519749 | up | NA |
| SLC26A11 | 1.50249394964651 | 0.00182534060977757 | up | Pathway |
| DENND1B | 1.50384594642119 | 0.00417108439348636 | up | Pathway |
| RFC4 | 1.50797964312409 | 0.00110916714653022 | up | Pathway |
| PACSIN1 | 1.50855361716523 | 0.0341748238107089 | up | Pathway |
| SLC39A11 | 1.50960290021139 | 0.00840837982487324 | up | NA |
| ASB13 | 1.51000745685354 | 0.00085848026156221 | up | NA |
| LETM2 | 1.51057873914715 | 0.026261623539383 | up | NA |
| STMN1 | 1.51240027339684 | 0.014328989318616 | up | Pathway |
| DOPEY2 | 1.51326242017417 | 0.000856772455144523 | up | NA |
| WDR34 | 1.51465431122731 | 0.00161970755890169 | up | NA |
| SOX12 | 1.51629243090994 | 0.00138046603939258 | up | NA |
| CENPP | 1.51664710221258 | 0.0271581850169887 | up | Pathway |
| SLC7A1 | 1.51673650584593 | 0.00366644821557256 | up | Pathway |
| ADAMTS6 | 1.51776550039531 | 0.0126162752683544 | up | Pathway |
| MTHFD2 | 1.51919235044522 | 0.000237946724054582 | up | Pathway |
| RELT | 1.51986167034583 | 0.0182138016689834 | up | Pathway |
| RTN4RL2 | 1.52006305015226 | 0.00142230717359239 | up | Pathway |
| FAM208B | 1.52436324698379 | 0.0130403891592346 | up | NA |
| BAI2 | 1.52495173116242 | 0.00553942800669223 | up | NA |
| IL23A | 1.52630300766491 | 0.015726010656474 | up | Pathway |
| RCC2 | 1.52832996948861 | 0.00122526659088574 | up | Pathway |
| CCSAP | 1.53343372453694 | 0.000565246610366704 | up | Pathway |
| MXRA5 | 1.53368500369042 | 0.0438787835422273 | up | Pathway |
| SIGLEC10 | 1.53816659844681 | 0.0246070864225715 | up | Pathway |

| | | | | |
|---|---|---|---|---|
| IRF9 | 1.53876821781306 | 0.0012160510220832 | up | NA |
| CLHC1 | 1.54054927192331 | 0.00555180461055724 | up | NA |
| PRR22 | 1.54164656110791 | 0.0178565327011306 | up | NA |
| WEE1 | 1.54225888663569 | 0.0340534062987513 | up | Pathway |
| ZNF26 | 1.54239421099693 | 0.000895982352114403 | up | NA |
| RBM47 | 1.542763109021 | 0.00294898602444929 | up | NA |
| ZNF710 | 1.54330818194659 | 0.00117093193320375 | up | NA |
| TCP10L | 1.54491181660693 | 0.0105109351990559 | up | NA |
| YJEFN3 | 1.54839080010154 | 0.00949004004081481 | up | Pathway |
| DOCK5 | 1.54973479981792 | 0.00638797079785207 | up | Pathway |
| SLC39A4 | 1.55168833115641 | 0.0022895754405791 | up | Pathway |
| EEF1E1 | 1.55198005610615 | 0.00112358637543383 | up | NA |
| NDUFV2 | 1.55204203609894 | 0.00102489591778388 | up | Pathway |
| DNMT3B | 1.55230749329307 | 0.00145915024249602 | up | NA |
| COL13A1 | 1.55244583824536 | 0.0455392936512178 | up | Pathway |
| INADL | 1.55258629530641 | 0.0235128775463151 | up | NA |
| FKBP10 | 1.55639330155245 | 0.00528967351329939 | up | Pathway |
| TIGD1 | 1.55785411643925 | 0.000840027060867341 | up | NA |
| TMEM97 | 1.55871948552029 | 0.00375394041768101 | up | Pathway |
| ZDHHC13 | 1.55948073405172 | 0.00130976520489504 | up | NA |
| LYG1 | 1.56045611107563 | 0.00390784209576858 | up | NA |
| AGRN | 1.56081999819592 | 0.00517956726880787 | up | Pathway |
| DDR1 | 1.56164383722412 | 0.03366291933718 | up | Pathway |
| AHRR | 1.56180980447807 | 0.00535468215420713 | up | Pathway |
| RCC1 | 1.56347980155084 | 0.00052570559702797 | up | Pathway |
| LCA5L | 1.56535774643928 | 0.0168518161798243 | up | Pathway |
| PYCRL | 1.56541381906321 | 0.000479061444774117 | up | NA |
| SLC23A3 | 1.56714486097106 | 0.00656201067176639 | up | NA |
| MSMO1 | 1.56717141541727 | 0.00735482662378602 | up | Pathway |
| GPR3 | 1.56743609470815 | 0.0475313872246112 | up | Pathway |
| UNC5CL | 1.56872278606834 | 0.0259938630067968 | up | Pathway |
| DCAF4L1 | 1.56910827716803 | 0.00139305578450717 | up | NA |
| BGN | 1.56943185114513 | 0.022772865904403 | up | Pathway |
| RCAN3 | 1.57280535324492 | 0.000841961641436113 | up | Pathway |
| ZNF682 | 1.57306297288513 | 0.00274639569119427 | up | NA |
| JPH1 | 1.57328302702301 | 0.0359633964652047 | up | Pathway |
| SLC37A1 | 1.57333814113294 | 0.00100650911501318 | up | Pathway |
| CDK18 | 1.57379334379924 | 0.0241480842247122 | up | Pathway |
| PLXNA3 | 1.57539000137844 | 0.000376032581302456 | up | Pathway |
| LEPREL4 | 1.57650205603703 | 0.000276020147346874 | up | NA |
| KRTCAP2 | 1.57712908720979 | 0.000717488454114974 | up | NA |
| CARD9 | 1.58026509260192 | 0.0449735123596271 | up | Pathway |
| NDUFAF6 | 1.58268849382377 | 0.000431772835303863 | up | NA |
| GPATCH2 | 1.58396476474232 | 0.00422296522054893 | up | NA |
| ZFP69B | 1.5850641341049 | 0.000350006354206234 | up | NA |
| ARHGEF19 | 1.58591487156012 | 0.0160438324150357 | up | Pathway |
| MBOAT1 | 1.5859664062831 | 0.00323520335464546 | up | Pathway |

| | | | | |
|---|---|---|---|---|
| MSI2 | 1.58815838000723 | 0.0168370907274003 | up | Pathway |
| ADC | 1.59159254444061 | 0.000391187746166089 | up | NA |
| KIAA1524 | 1.59273197180937 | 0.00271177006687139 | up | NA |
| BBS1 | 1.59442819508788 | 0.00131695493965 | up | Pathway |
| RPS6KA1 | 1.5969476116621 | 0.0100758667659766 | up | Pathway |
| RHOH | 1.59709494530642 | 0.0339725270594287 | up | Pathway |
| STK36 | 1.60076956301881 | 0.000292548645182149 | up | Pathway |
| PLEKHF2 | 1.60151306084402 | 0.00624288728893613 | up | NA |
| C19orf55 | 1.60396087215197 | 0.0000504491271049433 | up | NA |
| SEMA7A | 1.60453606503841 | 0.00470658635764137 | up | Pathway |
| CEP85 | 1.60477740849697 | 0.000111254350827603 | up | Pathway |
| MURC | 1.60507649327326 | 0.0142328962616396 | up | NA |
| RACGAP1 | 1.60700632335893 | 0.000035799033508745 | up | Pathway |
| EPT1 | 1.60765538758763 | 0.000206281533083547 | up | NA |
| GPC2 | 1.60884138256646 | 0.00453457564841719 | up | Pathway |
| MSANTD1 | 1.61393615629104 | 0.00790087356097827 | up | NA |
| RMI1 | 1.61537395234195 | 0.00142596384212134 | up | Pathway |
| C1orf233 | 1.61558465083718 | 0.0345290535500279 | up | NA |
| ZNF860 | 1.61584938566485 | 0.0139784975205989 | up | NA |
| TRIM11 | 1.62155963093558 | 0.000119175187363853 | up | Pathway |
| TBC1D30 | 1.62169359832503 | 0.0120390461542805 | up | Pathway |
| SLFNL1 | 1.62214199900018 | 0.00709856689335476 | up | NA |
| PPAT | 1.62526083175885 | 0.000135932393250195 | up | Pathway |
| ARRDC1 | 1.62606275487232 | 0.0000983664357695885 | up | Pathway |
| ZSWIM4 | 1.62799203471121 | 0.00420702857911762 | up | NA |
| C1QTNF3 | 1.62800844863545 | 0.00709624883671271 | up | Pathway |
| CAPS | 1.6290198686653 | 0.000302676112610582 | up | NA |
| PHLDA1 | 1.63108281687154 | 0.029406115022519 | up | NA |
| GPSM2 | 1.63574136923103 | 0.00591591608877752 | up | Pathway |
| ZNF107 | 1.63607786589428 | 0.00138848522194885 | up | NA |
| KCNS3 | 1.63653474080368 | 0.0204096430793536 | up | Pathway |
| SFXN2 | 1.64282241748174 | 0.00455346619506465 | up | Pathway |
| UBE2S | 1.64321463682342 | 0.00097481393680394 | up | Pathway |
| WFIKKN1 | 1.64345417625807 | 0.0114153028650823 | up | Pathway |
| CCDC40 | 1.64700438097154 | 0.0123659286727766 | up | Pathway |
| CACNB1 | 1.64709148557311 | 0.000762652416362351 | up | Pathway |
| FKBP4 | 1.64709970048809 | 0.000635415412915421 | up | Pathway |
| DEF6 | 1.64723134475746 | 0.0142774440656343 | up | Pathway |
| TRIM14 | 1.64897950436321 | 0.0149535239839532 | up | NA |
| DUSP5 | 1.64957921764173 | 0.0257179312757213 | up | Pathway |
| MTCP1 | 1.65256513291551 | 0.00151909420811683 | up | Pathway |
| ZNF474 | 1.65568428665533 | 0.00182453898514806 | up | NA |
| CA13 | 1.65577218574636 | 0.0196205807507955 | up | NA |
| OASL | 1.65911771430353 | 0.0308174165789104 | up | NA |
| MTG1 | 1.65922804795595 | 0.00152929163816738 | up | Pathway |
| LRRC37A3 | 1.66001557591926 | 0.00400762796617777 | up | NA |
| CYP2C8 | 1.66084681607716 | 0.00972901529699187 | up | Pathway |

| | | | | |
|---|---|---|---|---|
| VSIG1 | 1.66124338472104 | 0.00399434834763171 | up | Pathway |
| TNK1 | 1.66270515447493 | 0.00625942784465069 | up | Pathway |
| HSF2BP | 1.66349746082565 | 0.00392394620749101 | up | Pathway |
| CYP2D6 | 1.66646206153411 | 0.016520170236806 | up | Pathway |
| CHEK2 | 1.66684595075556 | 0.00372685603120057 | up | Pathway |
| FZD3 | 1.66956636141288 | 0.0196684058833414 | up | Pathway |
| CTPS1 | 1.66989560690413 | 0.000664676837799438 | up | Pathway |
| RAD54B | 1.67221694949107 | 0.00327517765179247 | up | Pathway |
| ZNF681 | 1.67435713536117 | 0.000831732132767331 | up | NA |
| EZR | 1.6747635648032 | 0.00393984991773987 | up | Pathway |
| CRYBB3 | 1.67591804094302 | 0.0395445275486255 | up | Pathway |
| SLC6A9 | 1.6763416392367 | 0.00659162986452108 | up | Pathway |
| BHLHE40 | 1.67673383387216 | 0.0189414705647988 | up | Pathway |
| AKAP5 | 1.67736617216759 | 0.00128174034231144 | up | Pathway |
| SYCE2 | 1.67750106497689 | 0.00537292428569105 | up | Pathway |
| KIAA1598 | 1.67887103515952 | 0.000291476034775124 | up | NA |
| SYNJ2 | 1.68130660105356 | 0.00245298894765407 | up | Pathway |
| STX1A | 1.68166516680496 | 0.00556965729927086 | up | Pathway |
| RNFT2 | 1.68274967466312 | 0.00220382262363427 | up | NA |
| CCDC154 | 1.68511348641169 | 0.00141113946069284 | up | Pathway |
| KNTC1 | 1.68848450731226 | 0.0000907487497202508 | up | Pathway |
| TARBP1 | 1.69130510247299 | 0.000859990336724413 | up | NA |
| PRSS27 | 1.6946126421249 | 0.0303636392380581 | up | NA |
| TRAF5 | 1.69836691946884 | 0.00589782611744088 | up | Pathway |
| PIP5KL1 | 1.69887300275775 | 0.00774965323616244 | up | Pathway |
| GRIN2D | 1.70123067829244 | 0.0370839698448013 | up | Pathway |
| MAATS1 | 1.70296744905278 | 0.0352387339369508 | up | NA |
| P2RY2 | 1.70318557609015 | 0.0312326090184183 | up | Pathway |
| SEC61A2 | 1.70507230947944 | 0.0000498615716999355 | up | NA |
| MAP7 | 1.70662228121255 | 0.00587517942299147 | up | Pathway |
| BAIAP2L1 | 1.71091260394476 | 0.000970047607259142 | up | Pathway |
| CTHRC1 | 1.71123030481394 | 0.0447490543568287 | up | Pathway |
| C3orf35 | 1.71231769090466 | 0.00130881877702352 | up | NA |
| TMEM51 | 1.71402854175122 | 0.0151313744375043 | up | NA |
| TTLL4 | 1.71485480111769 | 0.000549682823155422 | up | Pathway |
| OSBP2 | 1.71714837282394 | 0.00552420139638654 | up | Pathway |
| NFKBIE | 1.71831714185 | 0.0163955359713008 | up | Pathway |
| PASK | 1.72457916558553 | 0.0000276807225929861 | up | Pathway |
| CCDC11 | 1.72529554079856 | 0.00599418024721437 | up | NA |
| CYP2S1 | 1.72856015393124 | 0.0483848664517235 | up | Pathway |
| KIAA1244 | 1.73283045060703 | 0.0496919423366155 | up | NA |
| EXPH5 | 1.73500701169007 | 0.00324326249555947 | up | Pathway |
| ST20 | 1.73826683499567 | 0.00469072761669965 | up | Pathway |
| ZNF239 | 1.73880063088585 | 0.00469342995142372 | up | NA |
| SDC4 | 1.74095173997828 | 0.0131794853683773 | up | Pathway |
| TMEM79 | 1.74337771005287 | 0.00268498881833785 | up | Pathway |
| FAP | 1.74385835601803 | 0.00857867564802506 | up | Pathway |

| | | | | |
|---|---|---|---|---|
| CYB561 | 1.7452163536641 | 0.000587143854522317 | up | NA |
| ECT2 | 1.74555228400968 | 0.00855565502546065 | up | Pathway |
| RAB3IP | 1.74557687664274 | 0.00151454343948195 | up | Pathway |
| GALE | 1.74557785001671 | 0.00065086546894402 | up | Pathway |
| UNC5B | 1.74852831969865 | 0.00251873286739583 | up | Pathway |
| TPRN | 1.75183895275097 | 0.000841961641436113 | up | Pathway |
| MST1R | 1.75859943158356 | 0.0161599332334302 | up | Pathway |
| STRIP2 | 1.76002952803855 | 0.00276169247196343 | up | Pathway |
| TEX14 | 1.76154745766429 | 0.0381907572894224 | up | Pathway |
| TNFSF15 | 1.76278936214184 | 0.0121566532629491 | up | Pathway |
| ZNF367 | 1.76341592001691 | 0.00247576653171771 | up | NA |
| PDE6B | 1.76397959019458 | 0.04591131837312 | up | Pathway |
| SLC20A1 | 1.76461761468269 | 0.000170757522420596 | up | Pathway |
| PLAUR | 1.76475898257391 | 0.0121583559556684 | up | Pathway |
| RPS6KA6 | 1.76635373111518 | 0.0202824634713753 | up | Pathway |
| SEMA4D | 1.76850066509891 | 0.00247529312572664 | up | Pathway |
| ATAD5 | 1.76998696574414 | 0.000329307780244191 | up | Pathway |
| C19orf26 | 1.77009084386549 | 0.0486961408605182 | up | NA |
| KPNA2 | 1.77086340456787 | 0.000468452937123437 | up | NA |
| PLAC8L1 | 1.7714695071229 | 0.0115706337358168 | up | NA |
| DFNB31 | 1.77179470819072 | 0.00114539280751961 | up | NA |
| CYP2E1 | 1.7761519756944 | 0.00170637096365094 | up | Pathway |
| CCDC157 | 1.77729562392004 | 0.000313795619829168 | up | NA |
| RUNX1 | 1.77841411228472 | 0.0171675049022399 | up | Pathway |
| C2ORF15 | 1.77901798456221 | 0.00961806054540523 | up | NA |
| MFAP2 | 1.77908470786454 | 0.0481911870672368 | up | Pathway |
| CCDC57 | 1.77912349870609 | 0.000724733363954482 | up | Pathway |
| NME1-NME2 | 1.78006121621948 | 0.00245522107807217 | up | NA |
| LRRC48 | 1.78228830850883 | 0.0303140983516673 | up | NA |
| KIAA0907 | 1.78314680309554 | 0.000138870378028143 | up | NA |
| MTBP | 1.7837790483157 | 0.000519472046918103 | up | Pathway |
| SMC4 | 1.7877375837084 | 0.000056381582449123 | up | Pathway |
| KCNQ3 | 1.79183945673057 | 0.0264791426351144 | up | Pathway |
| SLC4A3 | 1.79288469072593 | 0.0079601083748213 | up | Pathway |
| NR6A1 | 1.79345420384346 | 0.0120589690265858 | up | NA |
| LAT | 1.79423453629297 | 0.0013012448199426 | up | Pathway |
| GPRIN1 | 1.79579612897357 | 0.00265369164960224 | up | NA |
| FAM60A | 1.79626696914553 | 0.00760429837564804 | up | NA |
| ARMC9 | 1.79636244447 | 0.0000800308091501004 | up | NA |
| AP1G2 | 1.79753296299595 | 0.000901124896509146 | up | NA |
| C9orf169 | 1.79907994121397 | 0.0130724014767002 | up | NA |
| SAMD10 | 1.8002743579483 | 0.000273291651608255 | up | NA |
| SHROOM2 | 1.8019108300946 | 0.0283452070632807 | up | Pathway |
| TET3 | 1.80252743310879 | 0.0000408366581746066 | up | NA |
| CHRNA5 | 1.80663073774293 | 0.0233385373569767 | up | Pathway |
| KIF24 | 1.80926940057137 | 0.00328464595664734 | up | Pathway |
| SSX2IP | 1.80929082152247 | 0.00162111992478466 | up | Pathway |

| | | | | |
|---|---|---|---|---|
| HIST1H2BJ | 1.8094895129272 | 0.00724666334109843 | up | NA |
| HID1 | 1.81035795413169 | 0.00455346619506465 | up | NA |
| BMP8B | 1.81085606962392 | 0.0295220277242881 | up | Pathway |
| TREML1 | 1.81514929961941 | 0.0473995801739087 | up | Pathway |
| NOD2 | 1.81651625465167 | 0.0460243861722245 | up | Pathway |
| CCDC19 | 1.81657786807188 | 0.0415187246244953 | up | NA |
| C3orf14 | 1.81703322785888 | 0.00526272322711181 | up | NA |
| SIX4 | 1.81824215296974 | 0.00513013504633908 | up | Pathway |
| BARD1 | 1.81889029977561 | 0.000212119686593945 | up | Pathway |
| IL18 | 1.82031494726246 | 0.0401056338960979 | up | Pathway |
| APOBR | 1.82173582640823 | 0.0142233426430729 | up | Pathway |
| AP1S3 | 1.82434403265144 | 0.0139253746321529 | up | NA |
| WDR52 | 1.82434737416377 | 0.00336688723201336 | up | NA |
| PARPBP | 1.82451045911272 | 0.00204486228999566 | up | NA |
| BAIAP2 | 1.82546779189156 | 0.000337262245187832 | up | Pathway |
| RPL36A | 1.82691921541605 | 0.000929013260278517 | up | NA |
| EPHX4 | 1.82749980052079 | 0.0400107234455228 | up | NA |
| WDR90 | 1.82777347211467 | 0.000810110470696398 | up | NA |
| FAM110A | 1.82793726254355 | 0.00142596384212134 | up | NA |
| SLC52A3 | 1.83179534507823 | 0.00880390044578562 | up | Pathway |
| ZNF469 | 1.83205861525468 | 0.0449735123596271 | up | Pathway |
| FAM102A | 1.83270774104383 | 0.00197572406218136 | up | NA |
| C1QTNF6 | 1.83315805474228 | 0.0000820462790595965 | up | NA |
| MPZL3 | 1.83569354517953 | 0.0172229037697034 | up | Pathway |
| CENPI | 1.83975916025388 | 0.00390443188665039 | up | Pathway |
| FCHO1 | 1.84041326059938 | 0.040428282763154 | up | NA |
| ABHD11 | 1.84142225869088 | 0.00078152247691911 | up | NA |
| ST6GALNAC2 | 1.8427025308541 | 0.0302871779978074 | up | Pathway |
| PLK4 | 1.84274395977602 | 0.00160438663234401 | up | Pathway |
| ZNF692 | 1.84351601062697 | 0.000554683115890849 | up | Pathway |
| RSPH9 | 1.84392537488736 | 0.0382716197100738 | up | Pathway |
| C11orf80 | 1.84628226505607 | 0.00213856146020748 | up | Pathway |
| LRGUK | 1.84665171422977 | 0.0392962172989077 | up | Pathway |
| FJX1 | 1.84757683951405 | 0.014245696298618 | up | Pathway |
| HIST1H2BN | 1.84858695435824 | 0.000732720734417925 | up | NA |
| CREB3L4 | 1.85110681511233 | 0.0110206353212982 | up | NA |
| SPOCD1 | 1.85180769096815 | 0.012510148231495 | up | NA |
| STAP2 | 1.85400441565752 | 0.0171132458612732 | up | NA |
| MUC20 | 1.85437701662394 | 0.0286484005193526 | up | NA |
| TMED3 | 1.85449124395025 | 0.00228257346191973 | up | NA |
| NRIP3 | 1.85461375004861 | 0.0489658709529967 | up | NA |
| ZNF93 | 1.85524361644699 | 0.00125534704783508 | up | NA |
| ATP1B1 | 1.8562865299406 | 0.0104599466767885 | up | Pathway |
| CCDC120 | 1.85635131367281 | 0.000178098843747599 | up | NA |
| CD9 | 1.8568445332454 | 0.00110360381131248 | up | Pathway |
| SFI1 | 1.85720954537198 | 0.000705608245898618 | up | NA |
| CDRT4 | 1.86002495688867 | 0.00372976604558838 | up | NA |

| | | | | |
|---|---|---|---|---|
| VMP1 | 1.86156231615018 | 0.000175174974499993 | up | Pathway |
| HMGA1 | 1.86157300358246 | 0.000217793877768999 | up | Pathway |
| ADORA2A | 1.86174771029096 | 0.0157281824916152 | up | Pathway |
| VEPH1 | 1.86303074677734 | 0.0498575277424041 | up | Pathway |
| OSBPL3 | 1.86331660305469 | 0.00565263646504274 | up | Pathway |
| PRR19 | 1.8638871356642 | 0.00191754479124839 | up | NA |
| ECE2 | 1.86450625781846 | 0.000949528071534232 | up | Pathway |
| SRGAP3 | 1.86534054835595 | 0.000320626940631743 | up | Pathway |
| SLC2A1 | 1.86632297497778 | 0.00177292501128082 | up | Pathway |
| MICAL2 | 1.86896073917152 | 0.000158964635373281 | up | Pathway |
| MCM4 | 1.86934899238048 | 0.000355843204294783 | up | Pathway |
| TNFRSF18 | 1.86954972410187 | 0.0238769799332038 | up | Pathway |
| TIGD4 | 1.86989171085738 | 0.00222443579069482 | up | NA |
| CASZ1 | 1.87652394198697 | 0.00988847123234526 | up | Pathway |
| LSR | 1.87735682324264 | 0.000345275370612215 | up | Pathway |
| GGT1 | 1.87855962239138 | 0.0242356839126835 | up | Pathway |
| ZDHHC23 | 1.87925705682134 | 0.00136971994879107 | up | Pathway |
| PRR11 | 1.87955376125878 | 0.00115445867089498 | up | NA |
| FAM174B | 1.88087268549348 | 0.00328966424894828 | up | NA |
| LAG3 | 1.88757504818882 | 0.0217328589941478 | up | Pathway |
| FAM83F | 1.88867399956913 | 0.0486044815845015 | up | NA |
| SIPA1L3 | 1.88958194557047 | 0.000662754921180765 | up | Pathway |
| YDJC | 1.89718686930348 | 0.000803271289386904 | up | NA |
| WNK3 | 1.89765262878956 | 0.0182929091605755 | up | Pathway |
| KCNK13 | 1.89773784110942 | 0.0165493220651772 | up | Pathway |
| AKR7L | 1.8988690293041 | 0.0257179312757213 | up | NA |
| MLLT11 | 1.90034712895784 | 0.00389757481976342 | up | Pathway |
| IPO4 | 1.90186186626524 | 0.0142192978643513 | up | NA |
| CPT1B | 1.90251687519514 | 0.00663943471616634 | up | Pathway |
| JAKMIP2 | 1.90778489086119 | 0.00213262441477563 | up | NA |
| SLC9A3R1 | 1.90789481412149 | 0.0162091537426571 | up | Pathway |
| LIG1 | 1.90866121347148 | 0.000158964635373281 | up | Pathway |
| JMJD7 | 1.90956936358976 | 0.00112332921428468 | up | NA |
| WNT5A | 1.91337897484017 | 0.0365428119178917 | up | Pathway |
| SPAG1 | 1.91441838334382 | 0.00352129257719098 | up | Pathway |
| SLC7A9 | 1.91544721839241 | 0.00390502642158636 | up | Pathway |
| CDC25A | 1.91728511991343 | 0.0115370744888728 | up | Pathway |
| HMGB3 | 1.92001124885234 | 0.0012238966854728 | up | Pathway |
| SGIP1 | 1.92111194822398 | 0.00460728611558834 | up | Pathway |
| KHDC1 | 1.92175061565146 | 0.0202824634713753 | up | NA |
| CCR4 | 1.92472591830702 | 0.0341914764137012 | up | Pathway |
| BLNK | 1.92538668657015 | 0.0057620729902112 | up | NA |
| CYFIP2 | 1.93026774088401 | 0.012081853020791 | up | Pathway |
| CELSR3 | 1.93058858353974 | 0.00233570228654904 | up | Pathway |
| GDPD1 | 1.93063998957767 | 0.0122380663877698 | up | Pathway |
| GPR141 | 1.93146928901371 | 0.00157236669490859 | up | NA |
| SPTBN5 | 1.9344401135729 | 0.00945737576368963 | up | Pathway |

| | | | | |
|---|---|---|---|---|
| BEST4 | 1.93471232883252 | 0.0248395598250645 | up | Pathway |
| TMEM238 | 1.93574749529154 | 0.00633149712061773 | up | NA |
| MYBL1 | 1.93795349709182 | 0.0044252766818311 | up | Pathway |
| SORD | 1.93823373524049 | 0.0130184411777078 | up | Pathway |
| GEN1 | 1.93986403219953 | 0.00025212751751663 | up | Pathway |
| P4HA3 | 1.94222963578284 | 0.0120387050254416 | up | NA |
| CLEC18B | 1.94896405702212 | 0.0132012206740265 | up | NA |
| LINC00176 | 1.94965716312986 | 0.011329964921088 | up | NA |
| SPIRE2 | 1.95055303249334 | 0.00121028633221942 | up | Pathway |
| CHAF1B | 1.9514366142783 | 0.000963778501952083 | up | Pathway |
| RASGRP1 | 1.95415655283487 | 0.0311552233583491 | up | Pathway |
| FAM222A | 1.95494501881519 | 0.00571236145696468 | up | NA |
| ATAD2 | 1.95598488813116 | 0.00788924216501734 | up | NA |
| CBS | 1.95626620775725 | 0.0330653920827545 | up | Pathway |
| F2RL1 | 1.95819721403649 | 0.0453778089154887 | up | Pathway |
| PSD | 1.95824952643197 | 0.014601409805193 | up | Pathway |
| GAPT | 1.95909849876975 | 0.0366901158454262 | up | NA |
| SKAP1 | 1.95972021062097 | 0.00115836595329297 | up | Pathway |
| ICOS | 1.96143581415875 | 0.0348684404319181 | up | NA |
| SEMA4A | 1.96143726462157 | 0.014245696298618 | up | Pathway |
| FANK1 | 1.96196361205294 | 0.0117086629742809 | up | NA |
| GET4 | 1.96359865682973 | 0.00105938301189131 | up | Pathway |
| PLS1 | 1.96429630720241 | 0.0230008926696326 | up | Pathway |
| CCDC24 | 1.9644982619757 | 0.00121028633221942 | up | Pathway |
| PHLDA2 | 1.96664859833097 | 0.0210871693102494 | up | Pathway |
| TLL2 | 1.96689684499544 | 0.0054739891480849 | up | Pathway |
| TRIB3 | 1.96722865536691 | 0.000349273418373698 | up | Pathway |
| CDKN2A | 1.96927363120504 | 0.0477539858737158 | up | Pathway |
| EME2 | 1.9699712095378 | 0.0131577181271704 | up | Pathway |
| TMEM30B | 1.97373468974793 | 0.0219683117028708 | up | Pathway |
| LRRC46 | 1.97390244025386 | 0.0031856931561156 | up | NA |
| SLC1A2 | 1.97874687666682 | 0.045352296765963 | up | Pathway |
| NEBL | 1.98006373642082 | 0.0411399224762018 | up | Pathway |
| RAB15 | 1.982718466858 | 0.0074157114338491 | up | Pathway |
| PHLDB3 | 1.98601820728202 | 0.00359636766911974 | up | NA |
| MAPK13 | 1.98629109153321 | 0.00194002085474097 | up | Pathway |
| CHTF18 | 1.9876670045417 | 0.000664676837799438 | up | Pathway |
| POLE2 | 1.99080666551803 | 0.000358181427641145 | up | Pathway |
| FAM81A | 1.99081877109647 | 0.00915882627269218 | up | NA |
| FTCD | 1.99315666088371 | 0.0326294115086425 | up | Pathway |
| SERPINE1 | 1.99330450122001 | 0.0312902307609046 | up | Pathway |
| DRP2 | 1.99666084852252 | 0.00537292428569105 | up | Pathway |
| KIAA1522 | 2.00034991605364 | 0.0020816284783849 | up | NA |
| ERBB2 | 2.00045147658858 | 0.00799086665294254 | up | Pathway |
| ENO2 | 2.00138238192368 | 0.00560887118460066 | up | Pathway |
| KCNN4 | 2.00161151505859 | 0.0449735123596271 | up | Pathway |
| SLC10A5 | 2.00179651785127 | 0.00236917230691088 | up | Pathway |

| | | | | |
|---|---|---|---|---|
| ACTL10 | 2.00330045605841 | 0.00565270531637202 | up | NA |
| SLC16A3 | 2.01021033862367 | 0.00122603999975415 | up | Pathway |
| MARCKSL1 | 2.01709120557987 | 0.00112938350159127 | up | Pathway |
| DNA2 | 2.01720698131317 | 0.000398274781386981 | up | Pathway |
| IFI30 | 2.01745566914113 | 0.0354635862272344 | up | Pathway |
| CHDH | 2.01909451449362 | 0.0168199353774218 | up | Pathway |
| DAPP1 | 2.01950656333019 | 0.0242032100760598 | up | NA |
| GINS4 | 2.0206987842829 | 0.0000594803468915483 | up | Pathway |
| ADAM8 | 2.02301148276919 | 0.000570638544941905 | up | Pathway |
| ZNF726 | 2.02558749654901 | 0.000600930863194888 | up | NA |
| C3orf80 | 2.02755702995697 | 0.00385452088387352 | up | NA |
| PAQR4 | 2.02867277830812 | 0.00182534060977757 | up | NA |
| SAMD12 | 2.02947217587159 | 0.0131337575959185 | up | NA |
| MICALL2 | 2.03090005502542 | 0.00167969947195216 | up | Pathway |
| MACC1 | 2.03168216742573 | 0.0380637370125983 | up | Pathway |
| RTKN | 2.03291403624554 | 0.000162681118293746 | up | Pathway |
| FLVCR1 | 2.03733309118873 | 0.0000532576945494599 | up | Pathway |
| NAT14 | 2.04013652141053 | 0.00229309730356969 | up | NA |
| CIT | 2.04179220686261 | 0.00237132491024521 | up | Pathway |
| PDE7A | 2.04338251976899 | 0.00025212751751663 | up | Pathway |
| OSCAR | 2.04584960318003 | 0.0370358228254132 | up | NA |
| FARP1 | 2.04711836178318 | 0.000094244323740427 | up | Pathway |
| C11orf35 | 2.05034815693214 | 0.00397360345476787 | up | NA |
| PFKFB4 | 2.05367654617467 | 0.000589285919388508 | up | Pathway |
| MCM2 | 2.05720389748042 | 0.000100688489044776 | up | Pathway |
| ABCC3 | 2.05727799037014 | 0.0215039520146331 | up | Pathway |
| LPAR2 | 2.05891018651618 | 0.00241236489864774 | up | Pathway |
| ZMYND10 | 2.06136256506857 | 0.00663943471616634 | up | Pathway |
| CHEK1 | 2.06209426365006 | 0.000828858884724498 | up | Pathway |
| ZNF587 | 2.06454475120993 | 0.00382033136612923 | up | NA |
| KIF15 | 2.06458111989272 | 0.0014712269319643 | up | Pathway |
| HS3ST3A1 | 2.06748252545463 | 0.0189614464643186 | up | NA |
| CTNND2 | 2.06868631345028 | 0.0377304304252964 | up | Pathway |
| CCNB1 | 2.07041276540608 | 0.00097390877809808 | up | Pathway |
| FOXO6 | 2.07081974260465 | 0.00228208135608419 | up | Pathway |
| MROH6 | 2.07093891150315 | 0.000111398699557408 | up | NA |
| CABLES2 | 2.07614231584558 | 0.0000983664357695885 | up | NA |
| GRID2IP | 2.07702561949466 | 0.000829905375575771 | up | Pathway |
| GSG2 | 2.08040679034609 | 0.00121028633221942 | up | NA |
| EGLN3 | 2.08108145195475 | 0.00430768835890432 | up | Pathway |
| CCDC148 | 2.08119913877075 | 0.0232383566339407 | up | NA |
| SLC22A18AS | 2.08250896342528 | 0.0360316518070838 | up | NA |
| CNIH2 | 2.08518724520868 | 0.000879322624593572 | up | Pathway |
| IL21R | 2.08613064064116 | 0.0282893565224409 | up | NA |
| HIST1H2AG | 2.08955072418456 | 0.000924332041181922 | up | NA |
| RAB39B | 2.09095445638808 | 0.0331292231104064 | up | Pathway |
| MAL2 | 2.09412450100466 | 0.0334752784866362 | up | Pathway |

| | | | | |
|---|---|---|---|---|
| LMO7 | 2.09524581563262 | 0.000677663935937173 | up | NA |
| FOXH1 | 2.09610636076858 | 0.0210715582621193 | up | Pathway |
| PAQR6 | 2.09695587884413 | 0.000232628787921521 | up | NA |
| XBP1 | 2.0982906711908 | 0.0352387339369508 | up | Pathway |
| SPC24 | 2.10144234581041 | 0.00215045672405595 | up | Pathway |
| MANEAL | 2.10582056395922 | 0.00131660031069336 | up | NA |
| TNFRSF12A | 2.10961827034957 | 0.0279405107787787 | up | Pathway |
| POSTN | 2.11032252078921 | 0.00200769638373584 | up | Pathway |
| GABRD | 2.11085764776192 | 0.00277830814423671 | up | Pathway |
| PANX2 | 2.11348325884533 | 0.0196892398352766 | up | NA |
| CNNM4 | 2.11456894245633 | 0.000408463821676327 | up | Pathway |
| HN1 | 2.11487003068832 | 0.000378487820262663 | up | NA |
| TLCD1 | 2.11558824152168 | 0.000574338703708593 | up | Pathway |
| FAM46C | 2.11750292353305 | 0.0201521533689393 | up | NA |
| CILP2 | 2.13257719810452 | 0.0140732295635356 | up | NA |
| RP5-1180C10.2 | 2.13561652110693 | 0.0000506385213250383 | up | NA |
| IGSF3 | 2.13817839166893 | 0.00892776600221047 | up | Pathway |
| ARHGAP11A | 2.14453734129862 | 0.000279298057998729 | up | Pathway |
| GRHL1 | 2.1487923819976 | 0.0180245130770724 | up | Pathway |
| GPR19 | 2.15126628125187 | 0.00130531727742229 | up | NA |
| SPATA17 | 2.15427875465114 | 0.0183883108996659 | up | NA |
| ABCC5 | 2.15428453475241 | 0.000235214577539393 | up | Pathway |
| HIST4H4 | 2.15709615825255 | 0.00934580710566081 | up | NA |
| TIGD3 | 2.16120957131676 | 0.00382033136612923 | up | NA |
| PHEX | 2.1614855193307 | 0.0123139997378851 | up | Pathway |
| SPEF1 | 2.16154484749408 | 0.0275377785953495 | up | Pathway |
| C15orf27 | 2.16180719191911 | 0.0493317391087599 | up | NA |
| KIAA1875 | 2.16365053650506 | 0.000736853069612146 | up | NA |
| KIAA1024 | 2.16729087373567 | 0.00170721093631515 | up | NA |
| ZNF90 | 2.16966353955302 | 0.00102838916679661 | up | NA |
| NSUN7 | 2.17186150938782 | 0.0138234450676705 | up | NA |
| A1BG | 2.17676393530839 | 0.00304131564925523 | up | NA |
| SLC23A1 | 2.17873771360133 | 0.0362026060557744 | up | Pathway |
| ZNF165 | 2.17907180674099 | 0.00379169275051211 | up | NA |
| CENPE | 2.18038579928963 | 0.00150727664972473 | up | Pathway |
| HOMER2 | 2.1806894367172 | 0.00493799547182434 | up | Pathway |
| KLHDC7B | 2.1807196170985 | 0.03433532476578 | up | NA |
| PMAIP1 | 2.18273339710478 | 0.0332165437967119 | up | Pathway |
| WWC1 | 2.18358958685347 | 0.0145239361597585 | up | Pathway |
| RASGRF1 | 2.19552098948723 | 0.00956606303894765 | up | Pathway |
| SOGA2 | 2.19560080914491 | 0.0170882038343085 | up | NA |
| FA2H | 2.19658377597048 | 0.0209399330720711 | up | Pathway |
| LRIT3 | 2.19980964482206 | 0.0244864170677308 | up | Pathway |
| NAGS | 2.19992688195777 | 0.00634286587794532 | up | Pathway |
| PODNL1 | 2.20151815611567 | 0.00779426574577066 | up | NA |
| CELSR2 | 2.20254354103578 | 0.0105711871070646 | up | Pathway |

| | | | | |
|---|---|---|---|---|
| C21orf58 | 2.20315050070318 | 0.000645663856617067 | up | NA |
| KIAA1467 | 2.20424094717622 | 0.0311491827122035 | up | NA |
| CSMD2 | 2.21679457581323 | 0.00387933783746444 | up | NA |
| FANCI | 2.21858068212235 | 0.00020066982285047 | up | NA |
| MCOLN2 | 2.22023909707506 | 0.00474603524970248 | up | Pathway |
| ASPHD2 | 2.22178567528248 | 0.0141538541072011 | up | NA |
| LRRC56 | 2.22253905621289 | 0.0085184269846312 | up | NA |
| DLG3 | 2.23108513192215 | 0.000592339872086392 | up | Pathway |
| BLM | 2.23479555266424 | 0.000197550176171525 | up | Pathway |
| PRRT2 | 2.24887018761541 | 0.00511581920671087 | up | Pathway |
| PLK1 | 2.25578217261358 | 0.000248730069891533 | up | Pathway |
| MAD2L1 | 2.25642351105252 | 0.000589908254731518 | up | Pathway |
| CDC7 | 2.25711456116415 | 0.000700595594406309 | up | Pathway |
| RASSF7 | 2.25779981616525 | 0.000191471622938169 | up | Pathway |
| TMEM61 | 2.25782456494324 | 0.0355721711712099 | up | NA |
| ARSE | 2.25851272551206 | 0.0394864668202511 | up | NA |
| SOX9 | 2.2603938930621 | 0.0184838246216 | up | Pathway |
| C2CD4D | 2.26433515383903 | 0.0128537793291694 | up | NA |
| AGMAT | 2.26646091562159 | 0.000334143519914029 | up | Pathway |
| ELMO3 | 2.26668789498689 | 0.000706264663722071 | up | Pathway |
| RELL2 | 2.26902998728133 | 0.00174230901888893 | up | Pathway |
| E2F1 | 2.27019200165327 | 0.00187635791697833 | up | Pathway |
| CCDC74B | 2.27240670753972 | 0.0303139848964262 | up | NA |
| EDARADD | 2.27614152117276 | 0.0497958395177395 | up | NA |
| CATSPERB | 2.27841905759205 | 0.0137140266551214 | up | NA |
| ABHD17C | 2.27914810657698 | 0.00648851095978662 | up | Pathway |
| RNF183 | 2.28043777555485 | 0.0347962131080547 | up | NA |
| CACNA1F | 2.2813273944392 | 0.00392715490930315 | up | Pathway |
| FSCN2 | 2.28528514722746 | 0.0216672179894037 | up | Pathway |
| LRP8 | 2.28609527081911 | 0.000519472046918103 | up | Pathway |
| ITGAX | 2.28681707119471 | 0.00744150879819119 | up | Pathway |
| MFSD2A | 2.28755117590655 | 0.0329264258250978 | up | Pathway |
| CAPN12 | 2.28811300911374 | 0.00188632887174866 | up | NA |
| PALM3 | 2.28985258758381 | 0.0236282185075584 | up | NA |
| LYPD1 | 2.29036853798518 | 0.0153365514896887 | up | Pathway |
| PRR7 | 2.29168754948722 | 0.00304614305296458 | up | NA |
| TMEM132A | 2.2930568549816 | 0.0017270286051826 | up | NA |
| MED12L | 2.29916654494698 | 0.00110509301084367 | up | NA |
| SPAG5 | 2.30072209196753 | 0.0000595770953396175 | up | Pathway |
| STIL | 2.30088047308066 | 0.000295209929728523 | up | Pathway |
| CORO2A | 2.30570068993054 | 0.00157709677579446 | up | NA |
| TRIP13 | 2.30898376291953 | 0.000369158002758366 | up | Pathway |
| DIO2 | 2.31096046472317 | 0.0247241021814546 | up | Pathway |
| GALNT7 | 2.31223109971324 | 0.0069801442531268 | up | NA |
| RAD51AP1 | 2.31224890187173 | 0.0000858818717921015 | up | Pathway |
| CTAGE8 | 2.31342513662882 | 0.0104252659821406 | up | NA |
| FAM169A | 2.31566160750784 | 0.0255384889473093 | up | NA |

| | | | | |
|---|---|---|---|---|
| KIFC2 | 2.31582405346442 | 0.000861267488020354 | up | Pathway |
| TDRD5 | 2.31605679391115 | 0.0322513805624948 | up | Pathway |
| KMO | 2.32096883545577 | 0.0380637634784904 | up | Pathway |
| MORN3 | 2.3254712573663 | 0.000404412180985721 | up | NA |
| CLEC7A | 2.32576252270712 | 0.00990834920104383 | up | Pathway |
| MCM10 | 2.32609524508807 | 0.00281392518247231 | up | Pathway |
| PIF1 | 2.32760608465273 | 0.00076654389496254 | up | Pathway |
| STARD10 | 2.32841272740542 | 0.00654297865903201 | up | Pathway |
| CAMK2A | 2.33455248045524 | 0.0171571843940662 | up | Pathway |
| RBM11 | 2.34136958867761 | 0.0126974212948193 | up | NA |
| IFNLR1 | 2.34205275114261 | 0.0411399224762018 | up | NA |
| SYTL1 | 2.34225353185344 | 0.00148633732729716 | up | NA |
| TFR2 | 2.34831480014765 | 0.0329264258250978 | up | Pathway |
| CCDC87 | 2.35594595559719 | 0.0096739140380748 | up | Pathway |
| SLC6A12 | 2.35872057751111 | 0.0131719812342726 | up | Pathway |
| DNASE1L2 | 2.36207165737849 | 0.0218618669133301 | up | Pathway |
| PSD4 | 2.36251230583047 | 0.00136523329018138 | up | Pathway |
| GUCY2D | 2.36812396377955 | 0.00989624148676905 | up | Pathway |
| NUAK2 | 2.37477236607306 | 0.00194219014779668 | up | Pathway |
| FANCD2 | 2.37649963822651 | 0.0000136429042467373 | up | Pathway |
| C6orf132 | 2.38134570944634 | 0.00942486447520965 | up | NA |
| PRRG2 | 2.38303030900865 | 0.00381070411012472 | up | NA |
| ATAD3C | 2.38558260413373 | 0.00422296522054893 | up | NA |
| CABYR | 2.38689549298517 | 0.00689884602230553 | up | Pathway |
| SYNGR3 | 2.39328998798477 | 0.0188549137608228 | up | Pathway |
| FRMD5 | 2.39355124382191 | 0.00760429837564804 | up | Pathway |
| CCDC150 | 2.39965750952209 | 0.00148676846448783 | up | NA |
| KIAA1549L | 2.40305670792245 | 0.000888253953419409 | up | NA |
| MLF1IP | 2.40460769257855 | 0.000417674733615524 | up | NA |
| NDC80 | 2.40546693136275 | 0.00118116352687324 | up | Pathway |
| PKIB | 2.40576798684473 | 0.0188365641459818 | up | Pathway |
| TYMS | 2.4099024464997 | 0.00388301788607342 | up | Pathway |
| WDR96 | 2.41900898769751 | 0.0371167201276631 | up | NA |
| RASGEF1A | 2.42702614003973 | 0.00120161497101071 | up | Pathway |
| FAM155B | 2.43116568810355 | 0.0158963601296545 | up | NA |
| RMI2 | 2.43200363772794 | 0.000844752341507983 | up | Pathway |
| PAFAH1B3 | 2.43521199169801 | 0.000104275122471357 | up | Pathway |
| MAP3K9 | 2.4355612849158 | 0.00189000128758172 | up | Pathway |
| PRRG4 | 2.437488614549 | 0.00329542057938119 | up | NA |
| AGBL2 | 2.44171628417771 | 0.00170538473396595 | up | NA |
| FOXP3 | 2.44258017577835 | 0.00165819933787852 | up | Pathway |
| FHDC1 | 2.44561701396806 | 0.0010675210015105 | up | Pathway |
| ZNF497 | 2.44762146369836 | 0.00100113982575308 | up | NA |
| PADI2 | 2.45282635888805 | 0.0191073887562299 | up | Pathway |
| CTSV | 2.45478955734052 | 0.00148635421887215 | up | Pathway |
| MBOAT2 | 2.45826908175214 | 0.000596665455601958 | up | Pathway |
| TRAF4 | 2.45844942668894 | 0.00153501584143883 | up | Pathway |

| | | | | |
|---|---|---|---|---|
| FAAH2 | 2.46024688893089 | 0.00507034402206909 | up | Pathway |
| CLDN9 | 2.4618422063815 | 0.0138967916718629 | up | Pathway |
| VWA7 | 2.46396745809571 | 0.0298674517852519 | up | NA |
| TRIM46 | 2.46458828224824 | 0.000757598198797455 | up | Pathway |
| TNFRSF9 | 2.46823043454399 | 0.0209053368295463 | up | NA |
| ZNF730 | 2.47029522597212 | 0.00659550276027539 | up | NA |
| NOXO1 | 2.47067385670745 | 0.0126162752683544 | up | Pathway |
| ZNF296 | 2.47297606863114 | 0.00145553954574202 | up | NA |
| ORC1 | 2.47350667986994 | 0.000263936114358879 | up | Pathway |
| STRC | 2.47380401165399 | 0.012782163899609 | up | Pathway |
| ILDR2 | 2.47395575442676 | 0.0468587190881898 | up | Pathway |
| MTFR2 | 2.47545789641534 | 0.000391187746166089 | up | NA |
| C11orf82 | 2.47718405972168 | 0.000736715997381874 | up | NA |
| ALCAM | 2.47997026183023 | 0.00713006730167117 | up | Pathway |
| PRC1 | 2.48062392120945 | 0.000123180519229478 | up | Pathway |
| E2F5 | 2.48375219114796 | 0.000516368145033116 | up | NA |
| SIGLEC8 | 2.48378082774085 | 0.0250663268582413 | up | NA |
| CTLA4 | 2.48809214164754 | 0.00987216328055093 | up | Pathway |
| KCNK1 | 2.49255929859111 | 0.0470695703615074 | up | Pathway |
| CDKN3 | 2.49365167929146 | 0.00491317584754662 | up | Pathway |
| CXCL11 | 2.49430245504768 | 0.0142926677598967 | up | Pathway |
| FAM72D | 2.49458436278261 | 0.00179694623517401 | up | NA |
| SLC22A15 | 2.49683109521124 | 0.0000838494407218363 | up | NA |
| MYO5B | 2.49888815642849 | 0.00965359416639844 | up | Pathway |
| CENPK | 2.51041709796336 | 0.000420208419873881 | up | Pathway |
| PPM1J | 2.51061634948531 | 0.0043034161229907 | up | NA |
| BRIP1 | 2.51320202579864 | 0.00106481929211232 | up | Pathway |
| FBN2 | 2.51369209579831 | 0.0033441245341242 | up | Pathway |
| SLC9A7 | 2.52103153511927 | 0.000377306285542243 | up | Pathway |
| SYCP2 | 2.52200814018741 | 0.0340264145065728 | up | Pathway |
| CCDC160 | 2.52226163260424 | 0.0413941573038584 | up | NA |
| ACTA1 | 2.52536535659551 | 0.0140709049290675 | up | Pathway |
| C5orf49 | 2.52798671057402 | 0.0346917710932833 | up | NA |
| C16orf59 | 2.53055965953179 | 0.000753195264254039 | up | NA |
| CCNA2 | 2.53123754320533 | 0.000268320741088932 | up | Pathway |
| AURKA | 2.53470940819506 | 0.000519472046918103 | up | Pathway |
| VDR | 2.53616919173651 | 0.00566837447176071 | up | Pathway |
| CLDN7 | 2.53727480999132 | 0.00165254468618578 | up | Pathway |
| LCN12 | 2.54100370929875 | 0.02601879802371 | up | NA |
| DOC2A | 2.54656391914207 | 0.00196351350704928 | up | Pathway |
| XK | 2.54690956873837 | 0.0234589142307055 | up | Pathway |
| PSRC1 | 2.54958429932507 | 0.0000351034889861726 | up | Pathway |
| CCDC74A | 2.5497781178798 | 0.0142988254831046 | up | NA |
| ENTPD2 | 2.5515592314666 | 0.00478310802381797 | up | Pathway |
| CDC6 | 2.55818198337214 | 0.000509006221677753 | up | Pathway |
| COL9A2 | 2.56124698138419 | 0.0169946930157186 | up | Pathway |
| CD80 | 2.56284127663467 | 0.000660059208830781 | up | Pathway |

| | | | | |
|---|---|---|---|---|
| HIST2H2BE | 2.56307771177952 | 0.00273471678689332 | up | NA |
| STXBP2 | 2.5676932658001 | 0.00158213708083203 | up | Pathway |
| TMEM92 | 2.57078680772653 | 0.0245649944482044 | up | NA |
| CHMP4C | 2.57992344576007 | 0.00051428028943494 | up | Pathway |
| KIAA0319 | 2.58185619404056 | 0.014832060538293 | up | Pathway |
| HIST1H4H | 2.58204635982678 | 0.00213856146020748 | up | NA |
| XRCC2 | 2.58230576983352 | 0.000501010287690176 | up | Pathway |
| ITPKA | 2.58320262333549 | 0.00100650911501318 | up | Pathway |
| ENTPD8 | 2.58859782850866 | 0.0212611060697359 | up | NA |
| C16orf93 | 2.59473981160759 | 0.000667898335859802 | up | NA |
| OCLN | 2.59474475193916 | 0.00689884602230553 | up | Pathway |
| FAM135B | 2.59966133675486 | 0.0317418480867959 | up | NA |
| CKS2 | 2.59987786495862 | 0.0000683419627159911 | up | Pathway |
| MROH7 | 2.60287736689686 | 0.0118873829995837 | up | NA |
| HIST1H2BD | 2.6029080781236 | 0.00428965940298131 | up | NA |
| CDH2 | 2.60509612222196 | 0.0198171702368437 | up | Pathway |
| PPAP2C | 2.60620066968865 | 0.0103681124456128 | up | NA |
| SP6 | 2.60802945637212 | 0.00542556753044505 | up | Pathway |
| SLC17A9 | 2.61109259786683 | 0.00810350097701753 | up | Pathway |
| PTK7 | 2.61303522787182 | 0.011329964921088 | up | Pathway |
| NETO2 | 2.61577453891867 | 0.00315255128750998 | up | Pathway |
| ZNF552 | 2.63132661245883 | 0.00796429223262063 | up | NA |
| ARHGAP39 | 2.64244090466502 | 0.000273291651608255 | up | Pathway |
| RAD51 | 2.64987979549768 | 0.000284014161466381 | up | Pathway |
| FCGR1A | 2.65089109711693 | 0.0408767389241535 | up | Pathway |
| SCG5 | 2.66172029999168 | 0.03433532476578 | up | Pathway |
| TFCP2L1 | 2.66309725871786 | 0.038557167127307 | up | Pathway |
| PAQR5 | 2.66655037144985 | 0.00502192460041498 | up | NA |
| EPHB3 | 2.67864522247714 | 0.00104676061895256 | up | Pathway |
| DLEU7 | 2.68039299619485 | 0.0028601567717847 | up | NA |
| PAK6 | 2.68725242011282 | 0.00422113960257483 | up | Pathway |
| GINS1 | 2.69054225239314 | 0.000421604515831116 | up | Pathway |
| CDK1 | 2.69148349978716 | 0.000805427054623683 | up | Pathway |
| RAB11FIP4 | 2.69213024485355 | 0.0027333224875454 | up | Pathway |
| TCTEX1D2 | 2.69561276322163 | 0.00142304010132895 | up | NA |
| SQLE | 2.70870992982269 | 0.00155839873950891 | up | Pathway |
| THSD4 | 2.71474932693327 | 0.049878905310642 | up | Pathway |
| MICALCL | 2.71536877208103 | 0.00261779260408553 | up | NA |
| CYP27B1 | 2.71543594279712 | 0.000698758446656829 | up | Pathway |
| SH3D21 | 2.726711532435 | 0.000719977459913792 | up | Pathway |
| EME1 | 2.72691943101819 | 0.000164752278344025 | up | Pathway |
| LMNB1 | 2.73340706641435 | 0.000127345443716461 | up | NA |
| CTXN1 | 2.73646047942656 | 0.0189660633835277 | up | NA |
| PRKCZ | 2.74022870238551 | 0.00270525293884244 | up | Pathway |
| FN1 | 2.74214056642138 | 0.000216635632680683 | up | Pathway |
| CCNI2 | 2.75752426597714 | 0.00810123338750151 | up | Pathway |
| CLSPN | 2.77726393889825 | 0.00177018203518982 | up | Pathway |

| | | | | |
|---|---|---|---|---|
| CCDC114 | 2.78315566490019 | 0.000378487820262663 | up | NA |
| DCDC1 | 2.78457813499501 | 0.0236187649805771 | up | Pathway |
| CXCR4 | 2.78515813817608 | 0.00177796070266155 | up | Pathway |
| STMND1 | 2.78699850324552 | 0.0433861818136822 | up | Pathway |
| UBE2T | 2.788000342649 | 0.000221168752651162 | up | NA |
| TRPS1 | 2.79472265464135 | 0.0035710718737098 | up | Pathway |
| ZWINT | 2.80146262374958 | 0.000615223146382208 | up | Pathway |
| HAGHL | 2.80364082312427 | 0.00180018263739458 | up | NA |
| CXCL10 | 2.8038001292575 | 0.00694679944106855 | up | Pathway |
| RASL11B | 2.81305886632066 | 0.00325856338812637 | up | Pathway |
| FAM84A | 2.81722175951065 | 0.0182929091605755 | up | NA |
| ESCO2 | 2.81731828505936 | 0.000927755789375418 | up | Pathway |
| IL17RB | 2.81831566972187 | 0.0257725501443704 | up | Pathway |
| ZSCAN1 | 2.82449301473976 | 0.0399378067214851 | up | NA |
| SLC38A1 | 2.82594293868553 | 0.00982997895865081 | up | Pathway |
| CDCA8 | 2.82697182207155 | 0.000162681118293746 | up | Pathway |
| EZH2 | 2.83285267878316 | 0.0000763833155876388 | up | Pathway |
| RHPN1 | 2.83870723140828 | 0.00152232638180539 | up | Pathway |
| PTTG1 | 2.8397587121969 | 0.000398274781386981 | up | Pathway |
| RUNX2 | 2.84255268365644 | 0.000408291543525218 | up | Pathway |
| SALL4 | 2.84685073118852 | 0.000607184694399605 | up | Pathway |
| POLQ | 2.86014546357065 | 0.000942497585998621 | up | Pathway |
| DHDH | 2.86257856881118 | 0.00283676451622642 | up | Pathway |
| BHLHA15 | 2.86270135581658 | 0.025151441560057 | up | Pathway |
| F7 | 2.86457474140936 | 0.0304047498136272 | up | Pathway |
| DNAH14 | 2.86626488520998 | 0.00187182897619392 | up | Pathway |
| GLYATL1 | 2.86802713945782 | 0.0111908227923717 | up | Pathway |
| EPS8L2 | 2.86870465417311 | 0.00112035390923959 | up | Pathway |
| GPR84 | 2.8738132506443 | 0.00138415355447896 | up | NA |
| HAMP | 2.87465195539205 | 0.0411399224762018 | up | Pathway |
| CDCA2 | 2.87928232953819 | 0.000260191438164158 | up | Pathway |
| SH2D3A | 2.87935187068646 | 0.0011802314378548 | up | Pathway |
| OIP5 | 2.88528515345055 | 0.000645364554820782 | up | Pathway |
| KCNC3 | 2.89697711933345 | 0.0188192131037244 | up | Pathway |
| MLK4 | 2.89898663224134 | 0.0160929878763038 | up | NA |
| TONSL | 2.90313350717423 | 0.0000680525117142832 | up | Pathway |
| KIAA1211 | 2.90617092695015 | 0.0106685439259569 | up | NA |
| SLCO5A1 | 2.90659646699214 | 0.0092925061324682 | up | Pathway |
| SLC12A8 | 2.90675913504687 | 0.000317717294706102 | up | Pathway |
| FBXO41 | 2.9071341563397 | 0.0000859453541881024 | up | NA |
| PABPC1L | 2.90782607125705 | 0.00083195246942083 | up | NA |
| KRT86 | 2.91206370152457 | 0.0111273098206496 | up | Pathway |
| TTC23L | 2.91526789837396 | 0.00639544851342644 | up | NA |
| SERINC2 | 2.91709592125491 | 0.00165254468618578 | up | Pathway |
| LRRC8E | 2.91849283416323 | 0.00160959441921452 | up | Pathway |
| SHCBP1 | 2.9264093555418 | 0.000191471622938169 | up | Pathway |
| ST14 | 2.93241207101729 | 0.00578133544523018 | up | Pathway |

| | | | | |
|---|---|---|---|---|
| GALNT5 | 2.93429929504314 | 0.00859723006064256 | up | NA |
| HPDL | 2.93437796088137 | 0.00799395219515198 | up | NA |
| FAM227A | 2.94671424470215 | 0.000793176632112059 | up | NA |
| LLGL2 | 2.9499687596398 | 0.000264003411359058 | up | Pathway |
| HSH2D | 2.95615260262081 | 0.000830139190852364 | up | Pathway |
| MARVELD2 | 2.96014094586494 | 0.00219255414040764 | up | Pathway |
| ROR2 | 2.96253306201531 | 0.0178314304358834 | up | Pathway |
| HTRA4 | 2.96277927353652 | 0.00761480424673054 | up | Pathway |
| IL4I1 | 2.96590477915471 | 0.00455346619506465 | up | Pathway |
| KLHDC8A | 2.96802920658119 | 0.0126290954502962 | up | NA |
| KIF11 | 2.96813647583934 | 0.000235725359934501 | up | Pathway |
| ERMN | 2.97139983903798 | 0.000287242442401643 | up | Pathway |
| TTBK1 | 2.97584659820847 | 0.00624626167325063 | up | Pathway |
| HIST3H2A | 2.98801758206312 | 0.000915276692190789 | up | NA |
| BRSK1 | 2.99424285613442 | 0.00415363598686419 | up | Pathway |
| GPRC5A | 2.99482112809304 | 0.00652618662248596 | up | Pathway |
| KRTCAP3 | 2.99942870206529 | 0.000709098607885016 | up | NA |
| FAM183A | 3.0087410389475 | 0.00895166323314354 | up | NA |
| KIF5C | 3.01086678116142 | 0.00936281328230613 | up | Pathway |
| INPP5J | 3.01451039465407 | 0.0183432371170624 | up | Pathway |
| CRABP2 | 3.02005592438813 | 0.00526819892395613 | up | Pathway |
| LYPD3 | 3.02084554240366 | 0.0125569495365769 | up | Pathway |
| GSTO2 | 3.02569658890095 | 0.0118980584902469 | up | Pathway |
| CCDC64 | 3.0293297812533 | 0.0162659280165425 | up | NA |
| TREM2 | 3.0395442019228 | 0.0190860427790771 | up | Pathway |
| RP11-366M4.3 | 3.0427526848575 | 0.0344821602330359 | up | NA |
| IRF6 | 3.04362050741323 | 0.00297707597284382 | up | Pathway |
| NCCRP1 | 3.04427010253701 | 0.0343460382512441 | up | NA |
| HNF4G | 3.04623368783228 | 0.0216005607968023 | up | NA |
| GINS2 | 3.04672116519102 | 0.000472155211243409 | up | Pathway |
| LEF1 | 3.04677334246805 | 0.000519990796027708 | up | Pathway |
| GRB7 | 3.06323170471271 | 0.00654297865903201 | up | Pathway |
| PITX1 | 3.06532636489792 | 0.017292770855018 | up | Pathway |
| FBXO43 | 3.07620713262274 | 0.000166776688431061 | up | Pathway |
| HIST1H1E | 3.07852481150222 | 0.000319134550957803 | up | NA |
| ARNT2 | 3.08284633796126 | 0.0101726972918696 | up | Pathway |
| ARHGEF39 | 3.08381091449985 | 0.000164917390145855 | up | NA |
| WDR62 | 3.08970402935171 | 0.000121388009059353 | up | Pathway |
| BARX2 | 3.09236897767064 | 0.0396562999793072 | up | Pathway |
| CDC42BPG | 3.09357405576571 | 0.000676026891912816 | up | Pathway |
| FANCA | 3.09488832123915 | 0.00011974337283636 | up | Pathway |
| BCL2L14 | 3.09683697456186 | 0.00864837331015004 | up | NA |
| WNT10A | 3.10040687601188 | 0.0295475886274442 | up | Pathway |
| COL7A1 | 3.10440342985429 | 0.00291972984985498 | up | Pathway |
| SFN | 3.10900978404063 | 0.0269450875724438 | up | Pathway |
| FBXO16 | 3.11028520435468 | 0.00651302524025588 | up | NA |
| LAMP3 | 3.1107001023753 | 0.0105220563316939 | up | NA |

| | | | | |
|---|---|---|---|---|
| TUBB3 | 3.11613773092279 | 0.0149034398144706 | up | NA |
| RECQL4 | 3.11675315389405 | 0.000689337876851608 | up | Pathway |
| SOWAHB | 3.11880757224355 | 0.0328182105721982 | up | NA |
| GOLT1A | 3.12598894961611 | 0.0113583676412547 | up | NA |
| ZNF365 | 3.12701846828141 | 0.000470998749679388 | up | Pathway |
| NCAPH | 3.13152879600656 | 0.0000447686575281947 | up | Pathway |
| CCNE2 | 3.1466895671318 | 0.000587515290771729 | up | Pathway |
| KIF5A | 3.15001980490232 | 0.01673747327592 | up | Pathway |
| NEURL3 | 3.15318477182362 | 0.00340359852612911 | up | NA |
| CASC5 | 3.15879452747965 | 0.000157468476137043 | up | NA |
| FSTL4 | 3.16210673100883 | 0.0179492886797449 | up | Pathway |
| BNIPL | 3.17033826322089 | 0.0395053252125013 | up | NA |
| PYCR1 | 3.17194977794952 | 0.000141818334379598 | up | Pathway |
| MND1 | 3.17317505078218 | 0.000761477801047362 | up | Pathway |
| ORC6 | 3.17616317606445 | 0.0000845369655144294 | up | Pathway |
| CDT1 | 3.17653812276869 | 0.0000735297797622699 | up | Pathway |
| ZYG11A | 3.17751027729174 | 0.00436192495129313 | up | NA |
| KIF18A | 3.17752560590404 | 0.0000884529995345068 | up | Pathway |
| MELK | 3.18221421817838 | 0.00018153478515827 | up | Pathway |
| SKA1 | 3.18346027027458 | 0.00233676442224468 | up | Pathway |
| MMP9 | 3.18524332374093 | 0.011329964921088 | up | Pathway |
| SULT2B1 | 3.18776631231172 | 0.0211117305656323 | up | Pathway |
| RIPK4 | 3.19389911911508 | 0.0353739221455307 | up | NA |
| GPR143 | 3.19435437858625 | 0.0283892247568998 | up | Pathway |
| PLEKHD1 | 3.19449260951295 | 0.0144108400502652 | up | NA |
| CDCA5 | 3.20033048794123 | 0.000146406827745791 | up | Pathway |
| FAM160A1 | 3.20167411932727 | 0.00348731366816943 | up | NA |
| RTKN2 | 3.20185744150486 | 0.0000146445125206015 | up | NA |
| MTL5 | 3.20881258281239 | 0.00462953827534328 | up | NA |
| MYBL2 | 3.21168938478698 | 0.000712904123714457 | up | Pathway |
| ANKRD22 | 3.21787879524466 | 0.0073949253539407 | up | NA |
| NKAIN1 | 3.22258824169313 | 0.048710658195052 | up | Pathway |
| CDCP1 | 3.22736945599455 | 0.0090021306941423 | up | NA |
| ADM2 | 3.23656981948364 | 0.00111536330428464 | up | Pathway |
| GALNT6 | 3.2368808381892 | 0.00900685147647643 | up | NA |
| TACSTD2 | 3.23860591723531 | 0.0383105053069992 | up | Pathway |
| RUFY4 | 3.2421812302422 | 0.00392339755665027 | up | NA |
| DNAH5 | 3.24222771045047 | 0.0227271145178156 | up | Pathway |
| DNAAF1 | 3.26018485974854 | 0.0090083680338739 | up | Pathway |
| TTC22 | 3.26194361152205 | 0.00739850157452355 | up | NA |
| HMMR | 3.26256579242262 | 0.000687630657276183 | up | NA |
| ASF1B | 3.26624226770606 | 0.0000836233586411006 | up | Pathway |
| TK1 | 3.26808488153447 | 0.000744473918723129 | up | NA |
| RHPN2 | 3.26877212228062 | 0.00196351350704928 | up | Pathway |
| FAM110C | 3.26901130196188 | 0.0204757896035019 | up | Pathway |
| SLC4A8 | 3.27791512643376 | 0.00360908416623356 | up | Pathway |
| ARHGAP8 | 3.27795900561669 | 0.00233676442224468 | up | Pathway |

| | | | | |
|---|---|---|---|---|
| PLEK2 | 3.28225582720762 | 0.000406484763669773 | up | Pathway |
| NUPR1L | 3.28523923047316 | 0.0329145813752973 | up | NA |
| ITGB6 | 3.28703567691495 | 0.0264052107779987 | up | Pathway |
| TTK | 3.28918713148792 | 0.000505918606544073 | up | Pathway |
| CACNA1D | 3.29396037926826 | 0.016749076455753 | up | Pathway |
| C3orf83 | 3.29601377907333 | 0.017562390602011 | up | NA |
| MEX3A | 3.29751441109944 | 0.00166626800176831 | up | NA |
| FAM83H | 3.29814661531851 | 0.000158964635373281 | up | Pathway |
| KIF23 | 3.30393504998237 | 0.000066995981626043 | up | Pathway |
| WNK2 | 3.30463568931765 | 0.0119995487134115 | up | Pathway |
| NUP210 | 3.31438919765552 | 0.0000439374577569317 | up | NA |
| CDS1 | 3.32384280686887 | 0.00148633732729716 | up | Pathway |
| TP73 | 3.32854796702327 | 0.019601973959696 | up | Pathway |
| AURKB | 3.33780801439234 | 0.00019746681291275 | up | Pathway |
| CDCA3 | 3.34808738247404 | 0.000205838892437834 | up | NA |
| CADPS | 3.35032103268006 | 0.00194003331594711 | up | Pathway |
| GSDMC | 3.35042979382447 | 0.00621178910355922 | up | NA |
| PLEKHN1 | 3.36040863334945 | 0.00994244746253262 | up | Pathway |
| TPX2 | 3.36085349409231 | 0.0000983664357695885 | up | Pathway |
| SBK1 | 3.36115244921663 | 0.00047340108910888 | up | Pathway |
| BCAS4 | 3.36288652249769 | 0.00192525122037535 | up | NA |
| NCAPG | 3.36470085820474 | 0.000408330584021694 | up | Pathway |
| C19orf45 | 3.36781242494206 | 0.00342730582790048 | up | NA |
| FAXC | 3.37373306723134 | 0.0047812945006144 | up | NA |
| ANXA9 | 3.37992037794648 | 0.0145609647351334 | up | Pathway |
| TPD52 | 3.38211274060569 | 0.0012733433616297 | up | NA |
| ZG16B | 3.38603430968805 | 0.0480230696749663 | up | Pathway |
| RNF223 | 3.39750573151363 | 0.0129136410098457 | up | NA |
| RAB17 | 3.39966206784184 | 0.00763884093437723 | up | Pathway |
| SCNN1A | 3.403639151982 | 0.00775383288365438 | up | Pathway |
| ERBB3 | 3.40394461614036 | 0.0126587341342774 | up | Pathway |
| SMKR1 | 3.40401618260149 | 0.000644903205574493 | up | NA |
| ARTN | 3.40880584416723 | 0.00205664522328738 | up | Pathway |
| CNKSR1 | 3.4097680433331 | 0.00122253999389884 | up | Pathway |
| KRT8 | 3.41733794124054 | 0.0044252766818311 | up | Pathway |
| ATG9B | 3.42176383473723 | 0.0031025774574955 | up | Pathway |
| HPN | 3.42377546215446 | 0.000882233408713756 | up | Pathway |
| EHF | 3.43673938663593 | 0.0270875601970393 | up | Pathway |
| RIBC2 | 3.44568344584613 | 0.001549696519053 | up | NA |
| PARD6B | 3.44691190036949 | 0.00872320757293315 | up | Pathway |
| RAB3B | 3.45099919038184 | 0.0120274146099159 | up | Pathway |
| E2F2 | 3.45307058589413 | 0.0013048365196367 | up | Pathway |
| CRB3 | 3.45318148062228 | 0.00312472742044704 | up | Pathway |
| CASKIN1 | 3.45679848941781 | 0.00245460802665968 | up | NA |
| CARD14 | 3.46432187983423 | 0.015622161643148 | up | Pathway |
| USP43 | 3.46944199117851 | 0.00104676061895256 | up | NA |
| LMTK3 | 3.47052880879421 | 0.00362660736073139 | up | NA |

| | | | | |
|---|---|---|---|---|
| FAM64A | 3.47651154883203 | 0.000743757915804901 | up | NA |
| GTSE1 | 3.47741352793175 | 0.000129439610476009 | up | Pathway |
| ST6GAL2 | 3.48171007674405 | 0.0135592768876727 | up | Pathway |
| CENPA | 3.48881287661793 | 0.00159967308514979 | up | Pathway |
| TOX3 | 3.4890538329658 | 0.0446369924330441 | up | NA |
| SPINT2 | 3.48972324391476 | 0.000519472046918103 | up | Pathway |
| HOOK1 | 3.50414000157181 | 0.00735443587241716 | up | NA |
| TRPA1 | 3.50545583058889 | 0.00755914818601825 | up | Pathway |
| DTL | 3.50766607678946 | 0.000117823399615808 | up | Pathway |
| HELLS | 3.51231244452813 | 0.0000834322071783268 | up | Pathway |
| KIF2C | 3.51381736620548 | 0.0000835210703882731 | up | Pathway |
| DSP | 3.51531644548929 | 0.00892806785177948 | up | Pathway |
| CENPM | 3.52228396494041 | 0.00223086015864832 | up | NA |
| APOBEC3B | 3.52456799169835 | 0.000940687174566504 | up | Pathway |
| SPP1 | 3.52929594515106 | 0.00042386930184934 | up | Pathway |
| ANLN | 3.53212482603676 | 0.000616152927867891 | up | Pathway |
| ESPL1 | 3.5331433036538 | 0.0000744532949887053 | up | Pathway |
| POU2F3 | 3.53391366387145 | 0.00680934672390898 | up | Pathway |
| KRT18 | 3.54576128887416 | 0.0048512817694514 | up | Pathway |
| PBK | 3.54899562388985 | 0.0014734502545674 | up | Pathway |
| PCP2 | 3.54943641147825 | 0.00972254968752351 | up | NA |
| CCDC78 | 3.55583820322518 | 0.00183181145755457 | up | Pathway |
| BUB1 | 3.55602306472424 | 0.000107255895851857 | up | Pathway |
| CELSR1 | 3.56028445254866 | 0.00292422104868455 | up | Pathway |
| KIFC1 | 3.56214075461254 | 0.00039228933178638 | up | Pathway |
| RP11-503N18.1 | 3.56429145136492 | 0.00131144497005329 | up | NA |
| DERL3 | 3.56821021110152 | 0.00664859283510803 | up | Pathway |
| KRT7 | 3.57102035577866 | 0.0253412692379924 | up | Pathway |
| FUT3 | 3.57232275767961 | 0.0111452924043911 | up | Pathway |
| C15orf48 | 3.57508322848792 | 0.00281418192569626 | up | NA |
| TFAP2A | 3.5803651384462 | 0.0446369924330441 | up | Pathway |
| CDC20 | 3.58233156534829 | 0.000538763688521035 | up | Pathway |
| PPFIA4 | 3.58315527266475 | 0.00090214920106453 | up | Pathway |
| CD164L2 | 3.58400149466253 | 0.0115579685895833 | up | NA |
| FAM57B | 3.58681664626887 | 0.00194156039182602 | up | NA |
| CEP55 | 3.58905628483753 | 0.00151850243845122 | up | Pathway |
| SLC7A5 | 3.59107757568973 | 0.00110281469064072 | up | Pathway |
| CCL11 | 3.59140244405754 | 0.0129473973639108 | up | Pathway |
| NUSAP1 | 3.59270429204966 | 0.000190284078984827 | up | Pathway |
| SLC16A6 | 3.59503590255757 | 0.0138967916718629 | up | Pathway |
| MYB | 3.59814280778534 | 0.0270322999128714 | up | NA |
| LAMP5 | 3.59851196429427 | 0.00299033480640625 | up | NA |
| SYNE4 | 3.60104094312325 | 0.00067186667339516 | up | Pathway |
| CDC45 | 3.60172890881638 | 0.000395483609521161 | up | Pathway |
| TRPV6 | 3.6031790297251 | 0.0328259654347378 | up | Pathway |
| B4GALNT3 | 3.61526578683562 | 0.00978410092385436 | up | NA |

| | | | | |
|---|---|---|---|---|
| EPHA10 | 3.61658228363675 | 0.022241106607553 | up | Pathway |
| KIAA0101 | 3.61740386826171 | 0.00022370204671549 | up | NA |
| CLPSL1 | 3.61859111094221 | 0.0304201532307021 | up | Pathway |
| TMC4 | 3.61880863617499 | 0.00241672990645292 | up | NA |
| KRT19 | 3.62094683264772 | 0.00517925832697476 | up | Pathway |
| B3GNT3 | 3.62480277690589 | 0.0161218932518052 | up | Pathway |
| DEPDC1B | 3.63028160414954 | 0.000363486883128481 | up | Pathway |
| FHAD1 | 3.6304510101069 | 0.00160614389893204 | up | NA |
| ABCA12 | 3.63710042848309 | 0.0341851002603364 | up | Pathway |
| LYPD6B | 3.64026024500636 | 0.0188259132023721 | up | NA |
| SLC44A4 | 3.64565320604247 | 0.0125363972198085 | up | Pathway |
| PPEF1 | 3.64689530642425 | 0.000299618041369827 | up | NA |
| TMPRSS6 | 3.64955452115246 | 0.00855582897386266 | up | Pathway |
| SEPT3 | 3.65089645994458 | 0.00145553954574202 | up | NA |
| RSPH1 | 3.65154552717319 | 0.00304480057341788 | up | Pathway |
| C9orf117 | 3.65445477125754 | 0.00455346619506465 | up | NA |
| BSPRY | 3.65539493297029 | 0.00595423768116199 | up | Pathway |
| RASAL1 | 3.65990479093659 | 0.0231049230020836 | up | Pathway |
| LPAR3 | 3.66229700705356 | 0.0259573302366102 | up | Pathway |
| ARHGEF38 | 3.66264208030312 | 0.0460931862875276 | up | NA |
| RAB26 | 3.66540385946758 | 0.00190145726290614 | up | Pathway |
| SCNN1G | 3.66636663272952 | 0.0225282775518651 | up | Pathway |
| AARD | 3.68662812062512 | 0.0296182361036083 | up | NA |
| DQX1 | 3.7065519727708 | 0.00328721285754064 | up | Pathway |
| FAM83D | 3.73398846940676 | 0.000275632924351592 | up | Pathway |
| CENPF | 3.73919035335862 | 0.000248543237413034 | up | Pathway |
| SHROOM3 | 3.73927930863891 | 0.00387307674648601 | up | Pathway |
| PKP3 | 3.7416722649716 | 0.000544691369367457 | up | Pathway |
| RHOV | 3.74641714687588 | 0.0162502018874337 | up | Pathway |
| LIPH | 3.74911342447966 | 0.0279432409809875 | up | Pathway |
| TMEM63C | 3.75094596025703 | 0.0116851286075757 | up | Pathway |
| TNS4 | 3.75268050754228 | 0.0489594945001134 | up | NA |
| PROM2 | 3.7551015449291 | 0.00678825962044422 | up | Pathway |
| SMPDL3B | 3.75643652902281 | 0.0219776248638008 | up | Pathway |
| RAP1GAP | 3.75813746074463 | 0.00392031743915764 | up | Pathway |
| SAPCD2 | 3.76708076619691 | 0.00314355662610002 | up | Pathway |
| RNF43 | 3.76734409086887 | 0.0120898151991117 | up | Pathway |
| EPS8L1 | 3.77436312669953 | 0.00440701294263352 | up | Pathway |
| TFAP2C | 3.78780124049936 | 0.00744474871211261 | up | Pathway |
| ESPN | 3.78850939035961 | 0.0152297030171976 | up | Pathway |
| ASPM | 3.78869721371203 | 0.00137451446761062 | up | Pathway |
| CGN | 3.79212635383485 | 0.0177739620225969 | up | NA |
| ESRP2 | 3.79345894080691 | 0.000830139190852364 | up | Pathway |
| SPC25 | 3.80186201517328 | 0.000228503668915116 | up | Pathway |
| SKA3 | 3.8034971206409 | 0.000368446268809778 | up | Pathway |
| KIF20A | 3.81091768840926 | 0.000676202287658234 | up | Pathway |
| FAM196A | 3.81387447281287 | 0.0315546195946172 | up | NA |

| | | | | |
|---|---|---|---|---|
| MUC1 | 3.81447877569435 | 0.00599418024721437 | up | Pathway |
| EPHA1 | 3.81872201620797 | 0.00126686845119704 | up | Pathway |
| PTK6 | 3.82079004418819 | 0.00982997895865081 | up | Pathway |
| C1orf106 | 3.82082388838142 | 0.0209823713098639 | up | NA |
| IGSF9 | 3.82246797170305 | 0.0116586765166049 | up | Pathway |
| BIK | 3.82271338164602 | 0.00944783160822367 | up | Pathway |
| KIF12 | 3.82448811844532 | 0.0293803406992471 | up | Pathway |
| OBP2A | 3.82840421160693 | 0.00198047408728795 | up | NA |
| ETV4 | 3.83252795634397 | 0.00212489036267517 | up | Pathway |
| NUF2 | 3.84295445825949 | 0.000281854030114776 | up | Pathway |
| ERVMER34-1 | 3.85331210361321 | 0.00495515236033379 | up | NA |
| CDC25C | 3.85643795214361 | 0.000122263566346754 | up | Pathway |
| PRLR | 3.86991068169011 | 0.00918450044101456 | up | Pathway |
| FOXM1 | 3.87075907180676 | 0.000483531691305894 | up | Pathway |
| PLEKHG4B | 3.87141530574673 | 0.0434662423487181 | up | Pathway |
| HIST1H3D | 3.87293628803239 | 0.00604092034650895 | up | NA |
| TMEM125 | 3.87906571249242 | 0.00271375848231484 | up | NA |
| NEURL | 3.88169433953446 | 0.00222750025134191 | up | NA |
| SRMS | 3.88669558431172 | 0.020762210500034 | up | Pathway |
| ACAN | 3.88937197827282 | 0.000924332041181922 | up | NA |
| SDS | 3.89618759365382 | 0.000669252547627322 | up | Pathway |
| ANO9 | 3.89687216983859 | 0.000519472046918103 | up | Pathway |
| TJP3 | 3.90351129042545 | 0.00324541237160142 | up | Pathway |
| FAM111B | 3.9058693597857 | 0.000549682823155422 | up | Pathway |
| BUB1B | 3.92568730715987 | 0.000262144502419119 | up | Pathway |
| CLUL1 | 3.93136452253606 | 0.00746380444011929 | up | NA |
| CKAP2L | 3.93243815394737 | 0.000146406827745791 | up | NA |
| GGT6 | 3.9333774167559 | 0.0211399492374203 | up | Pathway |
| CTD-3193O13.9 | 3.93820262953079 | 0.00301873399135759 | up | NA |
| GJB2 | 3.94310281643778 | 0.00155839873950891 | up | Pathway |
| IL11 | 3.9573838011835 | 0.000476999976161584 | up | Pathway |
| MFSD6L | 3.96356072091754 | 0.0014734502545674 | up | NA |
| CCL20 | 3.96745669669846 | 0.000217793877768999 | up | Pathway |
| TMEM184A | 3.9716979494808 | 0.00336189646931736 | up | NA |
| CXCL13 | 3.97648455550977 | 0.02601879802371 | up | Pathway |
| WISP1 | 3.97910570117269 | 0.00182453898514806 | up | NA |
| TROAP | 3.98328620045191 | 0.000148855646673274 | up | NA |
| SPDEF | 3.99419657963587 | 0.0271985415793516 | up | Pathway |
| RRM2 | 3.99873565565564 | 0.000164493266543036 | up | Pathway |
| NLRP2 | 4.0126886515256 | 0.0195792023221121 | up | NA |
| ILDR1 | 4.01399948995433 | 0.00130531727742229 | up | Pathway |
| TRIM59 | 4.02028781675901 | 0.0000594803468915483 | up | NA |
| HJURP | 4.04084895475553 | 0.000205838892437834 | up | Pathway |
| DEPDC1 | 4.04572468106916 | 0.00152837552553578 | up | NA |
| SLC7A11 | 4.05872602485958 | 0.000433136280562955 | up | Pathway |
| SUSD4 | 4.06290845985171 | 0.00179279225386599 | up | NA |

| | | | | |
|---|---|---|---|---|
| CORIN | 4.06551999230361 | 0.000187646059955679 | up | Pathway |
| CCNB2 | 4.08192223054799 | 0.000102385843801771 | up | Pathway |
| TOP2A | 4.08749080792726 | 0.000343693861847958 | up | Pathway |
| PLA2G4F | 4.09283475999268 | 0.0401368491550261 | up | Pathway |
| DLGAP5 | 4.09611700245789 | 0.00110509301084367 | up | Pathway |
| RGS1 | 4.10004335118271 | 0.00124880008254324 | up | Pathway |
| AIM1L | 4.10649457292245 | 0.00035511239248199 | up | NA |
| EVPL | 4.14418288634796 | 0.000347671560518047 | up | Pathway |
| C1orf172 | 4.15601424087294 | 0.00383985220207522 | up | NA |
| AP1M2 | 4.15965148088803 | 0.00624888804618131 | up | Pathway |
| NEIL3 | 4.1701988197952 | 0.000931600112341234 | up | NA |
| BMP8A | 4.17038953668279 | 0.000113657142831213 | up | Pathway |
| E2F8 | 4.1712785585092 | 0.000556422056111695 | up | Pathway |
| ZNF695 | 4.18327999300977 | 0.0000948332101410045 | up | NA |
| FXYD3 | 4.18775009995743 | 0.00762910226193442 | up | Pathway |
| EXO1 | 4.18865913694722 | 0.000233023681866618 | up | Pathway |
| RAD54L | 4.19118388967238 | 0.0000293106134078413 | up | Pathway |
| B3GALT5 | 4.19191513814152 | 0.033278857202715 | up | Pathway |
| MAPK15 | 4.19467516100246 | 0.00271742959675979 | up | Pathway |
| TDO2 | 4.20267264576092 | 0.00601521096434389 | up | Pathway |
| ESM1 | 4.2129155663698 | 0.00138944610105397 | up | Pathway |
| SPINT1 | 4.21634370464848 | 0.000664053732291982 | up | Pathway |
| COMP | 4.21697757319679 | 0.00828525701461702 | up | Pathway |
| S100P | 4.22150964115942 | 0.0125803689236678 | up | Pathway |
| KIAA1324 | 4.22992940403575 | 0.0150178978224102 | up | NA |
| KLK4 | 4.23201111968415 | 0.00501499406465926 | up | Pathway |
| BIRC5 | 4.23975586765353 | 0.000533320369262864 | up | Pathway |
| B4GALNT4 | 4.2554732728037 | 0.000915276692190789 | up | NA |
| C9orf152 | 4.26755411866275 | 0.0221255765870114 | up | NA |
| FUT2 | 4.26888056269836 | 0.00984875232019564 | up | Pathway |
| EDN2 | 4.28597146923453 | 0.00451789014201225 | up | Pathway |
| PKMYT1 | 4.28987814276348 | 0.000402033618715887 | up | Pathway |
| CDH1 | 4.29250934629162 | 0.00879568643309762 | up | Pathway |
| C5orf46 | 4.29581410372717 | 0.00203073193953047 | up | NA |
| KIF14 | 4.29792912959661 | 0.0011393710066944 | up | Pathway |
| KIF4A | 4.30031785204686 | 0.000422655761427861 | up | Pathway |
| WNT7B | 4.31144970797847 | 0.0279094018088086 | up | Pathway |
| TMEM52B | 4.31957573875663 | 0.000216329881928068 | up | NA |
| ATP6V0D2 | 4.32606394470569 | 0.00272612419835538 | up | Pathway |
| UBE2C | 4.32837661314579 | 0.000840027060867341 | up | Pathway |
| MMP3 | 4.32855007877641 | 0.00617849064155119 | up | Pathway |
| CELF5 | 4.33210377550953 | 0.00972254968752351 | up | NA |
| SDC1 | 4.34541390162561 | 0.000362782043348919 | up | Pathway |
| MISP | 4.35057042109413 | 0.00165254468618578 | up | Pathway |
| PRR15L | 4.36029222962226 | 0.0123659286727766 | up | NA |
| KRT80 | 4.36534267911231 | 0.00150230821724179 | up | Pathway |
| C1orf116 | 4.37233640357646 | 0.0104564655945482 | up | NA |

| | | | | |
|---|---|---|---|---|
| KLC3 | 4.39152877488641 | 0.00389636423202183 | up | Pathway |
| CPNE7 | 4.40626808519386 | 0.00460648456323948 | up | Pathway |
| OVOL2 | 4.41133610542074 | 0.0121694911558815 | up | Pathway |
| TNNT1 | 4.41620006175927 | 0.0232111884782677 | up | Pathway |
| F2RL2 | 4.42063853242277 | 0.00471885773183127 | up | Pathway |
| INHBA | 4.42645439509597 | 0.000063970729728925 | up | Pathway |
| RASEF | 4.42876288845529 | 0.00798078523132244 | up | NA |
| SGOL1 | 4.44547595712331 | 0.000295212942885313 | up | NA |
| ERCC6L | 4.45008199505872 | 0.000384763980417933 | up | Pathway |
| MKI67 | 4.45688938633244 | 0.0000835210703882731 | up | Pathway |
| NEK2 | 4.45690554308659 | 0.000123180519229478 | up | Pathway |
| E2F7 | 4.49467618968536 | 0.000116145582440322 | up | Pathway |
| CEACAM6 | 4.49984743978936 | 0.0163191461395652 | up | Pathway |
| IGFL2 | 4.52837230805188 | 0.000751117582664914 | up | NA |
| SMIM22 | 4.54164350471305 | 0.00901198948306945 | up | Pathway |
| TMPRSS2 | 4.54834961762209 | 0.0119069572102935 | up | NA |
| CAPN13 | 4.55170350966035 | 0.00775700125981247 | up | NA |
| ADAMTS14 | 4.56440070954162 | 0.000422655761427861 | up | Pathway |
| CLEC5A | 4.57103561992426 | 0.000248598917320596 | up | Pathway |
| ARMC3 | 4.57525389993196 | 0.00392715490930315 | up | NA |
| CAPN9 | 4.58107020715432 | 0.00265375791816514 | up | Pathway |
| CACNG4 | 4.61013506837501 | 0.00643910889005658 | up | Pathway |
| CLDN4 | 4.61678796196774 | 0.00205820296556223 | up | Pathway |
| LRRC15 | 4.63584260614476 | 0.00138641690390131 | up | Pathway |
| OVOL1 | 4.63632405045167 | 0.00237750733161645 | up | Pathway |
| MARVELD3 | 4.65505206187843 | 0.0030742513166216 | up | Pathway |
| ATP2C2 | 4.66618735834296 | 0.00426390831046671 | up | Pathway |
| EPN3 | 4.67267052116451 | 0.00712630531540568 | up | NA |
| TICRR | 4.69113915533089 | 0.0000907487497202508 | up | Pathway |
| IQGAP3 | 4.70191100284228 | 0.000162610489219813 | up | Pathway |
| ST6GALNAC5 | 4.71327057855147 | 0.00121455867467012 | up | Pathway |
| CAMSAP3 | 4.71901339830188 | 0.00735443587241716 | up | Pathway |
| C1orf210 | 4.73678013665102 | 0.00454529270740616 | up | NA |
| RAB25 | 4.76013211831843 | 0.00165780572876931 | up | Pathway |
| CBLC | 4.77569065167934 | 0.00258052237930075 | up | Pathway |
| ELF3 | 4.78378137193348 | 0.00162988989398329 | up | Pathway |
| PVRL4 | 4.78966662713827 | 0.00167836697251438 | up | NA |
| S100A14 | 4.79764366444056 | 0.012194856171773 | up | Pathway |
| GRHL2 | 4.8174384287864 | 0.00345806640655465 | up | Pathway |
| PRSS22 | 4.82844903596401 | 0.00356982149049716 | up | NA |
| KIF18B | 4.83946376549738 | 0.000141818334379598 | up | Pathway |
| LAD1 | 4.89095767396952 | 0.0024543271402838 | up | NA |
| AUNIP | 4.89571494059834 | 0.0000976214217055977 | up | Pathway |
| KIAA1199 | 4.89725468326449 | 0.00721812611063696 | up | NA |
| COL22A1 | 4.92750596337662 | 0.00142304010132895 | up | Pathway |
| CLDN3 | 4.93232830712026 | 0.0100751328258592 | up | Pathway |
| OLR1 | 4.93363625945041 | 0.000118560170296314 | up | NA |

| | | | | |
|---|---|---|---|---|
| EPCAM | 4.96662532110052 | 0.00197239006890241 | up | Pathway |
| FAM83H-AS1 | 4.97021721578249 | 0.00039694894959709 | up | NA |
| ADAMDEC1 | 4.97927727053638 | 0.00270055283298547 | up | Pathway |
| KIF26B | 5.02397906548448 | 0.0000352912930170513 | up | Pathway |
| RBBP8NL | 5.05007184734798 | 0.00138046603939258 | up | NA |
| CHRNA1 | 5.09530693226678 | 0.0000292644696970919 | up | Pathway |
| CCDC64B | 5.10364734535457 | 0.0098802366493417 | up | NA |
| PRSS8 | 5.16774053310092 | 0.00491317584754662 | up | Pathway |
| ALDH3B2 | 5.22277994299483 | 0.0131197189562952 | up | Pathway |
| SOX11 | 5.33885721191544 | 0.00023427802547337 | up | Pathway |
| GRM4 | 5.6900012100117 | 0.000960026707087691 | up | Pathway |
| SLC24A2 | 5.8307316055099 | 0.000104447813318692 | up | Pathway |
| PPAPDC1A | 5.89441794332459 | 0.0000640331291124177 | up | NA |
| MMP11 | 5.93725084558441 | 4.93078487897174E-06 | up | Pathway |
| SYT13 | 5.99051358896842 | 0.00717527616884998 | up | Pathway |
| ESRP1 | 5.99542029991487 | 0.00123650385139297 | up | Pathway |
| IBSP | 6.14704576164679 | 0.0000180908232180651 | up | Pathway |
| MMP1 | 6.48157335078994 | 0.000565906742003079 | up | Pathway |
| COL11A1 | 6.64567184111418 | 6.42952632029787E-06 | up | Pathway |
| STRA6 | 6.77790701215419 | 0.000795808191345553 | up | Pathway |
| MMP13 | 6.81066822017292 | 0.0000115988018709341 | up | Pathway |
| COL10A1 | 7.47891707111377 | 0.0000110323197280703 | up | Pathway |
