## Supplementary material for "A recurrent pathogenic *BRCA2* truncating variant reveals a role for BRCA2-PCAF complex in modulating NF-κB-driven transcription": Table S5-S7

**Table S5**. Oligonucleotides used for gene editing and HR assay

| Sequence (5’-3’) | PURPOSE | LAB IDENTIFIER |
| --- | --- | --- |
| TGTGGGATTTTTAGCACAG | Target sequence for BRCA2-delT mutation (TALEN1) |  |
| TGAAGCATCTGATACCTG | Target sequence for BRCA2-delT mutation (TALEN2) |  |
| GACAGTGAAAACACAAATCA | Target sequence for BRCA2-del5 mutation (CRISPR) |  |
| GCACAGCAAGTGAAAATCTGTCC | Target sequence to correct BRCA2-delT mutation (CRISPR) |  |
| GACAGTGAAAACACAAAGAG | Target sequence for BRCA2-rev del5 mutation (CRISPR) |  |
| GGGGCCACTAGGGACAGGAT | sgRNA to generate a DSB in the AAVS1 locus (CRISPR) for HR assay |  |
| GCTGACATTCAGAGTGAAGAAATTTTACAAC | Genotyping of +/delT and +/revdelT (F) | oAC337 |
| GCAGATGAATTTACCACATTATATGAAAAGCC | Genotyping of +/delT and +/revdelT(R) | oAC338 |
| GGACAAAGGGATGATTCATGTCCCAAG | Genotyping of del5 (F) | oAC525 |
| GGTGAAATGCCATCTCTACTAAAAATCAATAGTTG | Genotyping of del5 (R) | oAC517 |
| GTCCTGCAACTTGTTACACAAATCAGTCC | Forward oligonucleotide to amplify by PCR the homology flanking sequence including the delT mutation from CAPAN-1 cells for TALEN editing | oAC350 |
| CCCCAAACTGACTACACAAAAATGGCTG | Reverse oligonucleotide to amplify by PCR the homology flanking sequence including the delT mutation from CAPAN-1 cells for TALEN editing | oAC378 |
| T·T·CAACTAAACAGAGGACTTACCATGACTTGCAGCTTCTCTTTGTGTTTTCACTGTCTGTCACAGAAGCGATAAATCT·A·T | Oligonucleotide containing the sequence of homology including the BRCA2 999del5 mutation for CRISPR gene editing | oAC513 |
| T·T·CAACTAAACAGAGGACTTACCATGACTTGCAGCTTCTCTTTGATTTGTGTTTTCACTGTCTGTCACAGAAGCGATAAATCTATA·G·T | Oligonucleotide containing the sequence of homology including the BRCA2 999del5 reverted to WT for CRISPR gene editing | oAC994 |
| AGCGTCCGTGGCCACTG (F)  ACTTCGGTTTGGGCGCAG (R) | To check the expression of *CXCL1* | oAC1209 and 1210 |
| TTGCCTGGTGAAAATCATCAC (F)  AGGAACTGGATCAGGACTTTTG (R) | To check the expression of *IL6* | oAC1207 and 1208 |
| TGTCCAAATCGATGTGGATGTTTC (F)  TTGTACCATTCTTCTGCCTCCTG (R) | To check the expression of *VIM* | oAC1135 and 1136 |
| TGACTTCGGTTTGGGCGC (F)  GGAGCGTCCGTGGTCAC (R) | To check the expression of *CXCL3* | oAC1223 and 1224 |
| GCCAAGGAGTGCTAAAGAACT (F)  TAATTTCTGTGTTGGCGCAGTG (R) | To check the expression of *CXCL8* (or *IL8*) | oAC1221 and 1222 |
| TCAGGTGGCAGTGGCTGC (F)  CTGAAGCACCTGGGAGTG (R) | To check the expression of *NFKBIZ* | oAC1225 and 1226 |
| CTGGCAGGTACTGGAATTCC (F)  CTGATGTGCACCGACAAGTG (R) | To check the expression of *RelA (NFKB3)* | oAC1227 and 1228 |
| GGTGTGAATAACTTTGTGCAGTA (F)  TCATTGGGAGATCGCAGTCTT (R) | To check the expression of *PCAF* | oAC1229 and 1230 |
| TAATGGCACTGGTGGCAAGTCC (F)  CACATGCTTGCCATCCAACCAC (R) | To check the expression of *PPIA* | oAC985 and 986 |

(·) Denotes a phosphorothioate linkage modification

| Construct | Source | Identifier |
| --- | --- | --- |
| TALEN BRCA2 delT fok F pcDNA6.2/N-EmGFP-DEST GW | Life technologies | #V35620 |
| TALEN BRCA2delT fok R pcDNA6.2/N-YFP-DEST GW | Life technologies | #V35820 |
| pCas9-GFP | Addgene | #44719 |
| pBS U6 sgRNA CRISPR | Addgene | #43860 |
| pU6-pegRNA-GG-acceptor | Addgene | #132777 |
| pcDNA6.2 N-EmGFP | Kyle Miller’s lab |  |
| pcDNA6.2 N-EmGFP-PCAF | Kyle Miller’s lab |  |
| pCMV1-EGFP-MBP-2xNLS | Alvaro-Aranda *et al.,* NAR 2023 | pAC344 |
| pCMV1 EGFP-MBP-BRCA2 | Von Nicolai *et al.,Nature Commun.* 2016 | pAC037 |
| pCMV1 MBP-BRCA2-WT-RFP | Alvaro-Aranda *et al.,* NAR 2023 | pAC337 |
| pCMV1 EGFP-MBP-delT | This study (GenScript) | pAC105 |

**Table S6**. Plasmids

**Table S7**. Barcodes RNAseq and WGS

| Barcode | ID RNAseq | ID WGS |
| --- | --- | --- |
| Ctrl #1 | B255T01 |  |
| Ctrl #2 | D1471T148 |  |
| Ctrl #3 | D610T170 |  |
| del5 #1 | D956T267 |  |
| del5 #2 | D956T268 |  |
| del5 #3 | D1342T008 |  |
| delT #1 | D956T269 |  |
| delT #2 | D610T171 |  |
| delT #3 | D610T172 |  |
| 452087H_H | D454T17 | D458R01 |
| 462295H_H | D454T18 | D458R02 |
| 549025H_H | D454T19 | D458R03 |
| 619138H_H | D454T20 | D458R04 |
| 631699H_H | D454T21 | D458R05 |
| 638538H_H | D454T22 | D458R06 |
| H091731_H | D454T23 | D458R07 |
| H150105_H | D454T24 | D458R08 |
| 514207H_H | D454T26 | D458R09 |
| 371480H_H | D454T27 | D458R10 |
| 452087H_T | D454T28 | D458R11 |
| 462295H_T | D454T29 | D458R12 |
| 549025H_T | D454T30 | D458R13 |
| 619138H_T | D454T31 | D458R14 |
| 631699H_T | D454T32 | D458R15 |
| 638538H_T | D454T33 | D458R16 |
| H091731_T | D454T34 | D458R17 |
| H150105_T | D454T35 | D458R18 |
| 514207H_T | D454T37 | D458R20 |
| 371480H_T | D454T38 | D458R21 |
| R01_T |  | D1320-D1317R01 |
| R03_T |  | D1320-D1317R03 |
| R01_H |  | D1320-D1317R04 |
| R03_H |  | D1320-D1317R06 |

*H: healthy tissue; T: tumors
